# Classification of Urticaceae based on morphology and phylogenetic inference

**DOI:** 10.1101/2025.05.16.651835

**Authors:** Alexandre K. Monro, Olivier Maurin, Long-Fei Fu, Tom Wells, Melanie Wilmot-Dear, Juliet Beentje, D.J. Nicholas Hind, Ib Friis, Yi-Gang Wei, Grace Brewer, Robyn Cowan, Steven Dodsworth, Jia Dong, Niroshini Epitawalage, Izai A.B. Sabino Kikuchi, Isabel Larridon, Alison Moore, Hervé Sauquet, Jacquie Ujetz, Zeng-Yuan Wu, Felix Forest, William J. Baker, Elliot Gardner

## Abstract

The Urticaceae (ca. 2600 species) were first formally recognized by Jussieu in the 18th century and last comprehensively monographed by Weddell in the 19th century. Since Weddell’s work, family delimitation has been modified and many genera described in a fragmented manner. Over the past two decades, numerous molecular studies have supported the inclusion of Cecropiaceae within Urticaceae and identified paraphyly in several genera, notably *Laportea, Urera, Boehmeria, Parietaria, Pellionia,* and *Pouzolzia*. However, few studies have translated these molecular insights into a revised taxonomy.

This study aimed to provide a robust, updated classification for Urticaceae by: a) increasing taxon and genomic locus sampling through the integration of newly generated sequence data with previously published datasets; and b) incorporating morphological data to support a revised delimitation of tribes and genera, and to establish a new linear sequence for the family. We also sought to identify remaining taxonomic challenges.

Using Sanger and Angiosperms353 sequence data, we constructed a phylogenetic framework for 57 out of 59 currently accepted genera. We also assessed the phylogenetic informativeness of 57 morphological characters by mapping them onto the phylogeny.

Our analyses support the delimitation of 61 monophyletic genera and an infrafamilial classification comprising seven tribes, two of which we describe as new: Myriocarpeae and Leukosykeae. We provide a revised linear sequence for the family. Our classification reinstates several names previously treated as synonyms (*Fleurya, Leptocnide, Margarocarpus, Polychroa, Scepocarpus, Sceptrocnide*), places several genera in synonymy (*Hemistylus* and *Rousselia* under *Pouzolzia*; *Hesperocnide* under *Urtica*; *Gesnouinia* and *Soleirolia* under *Parietaria*), and proposes the recognition of two new genera, *Muimar* gen. nov. and *Pouzolziella* gen. nov., to accommodate *Boehmeria nivea* and *Pouzolzia australis,* respectively.

Mapping morphological characters onto the phylogeny indicates that while most states are homoplastic at the family level, their combination is valuable for recognizing genera. Geographic character mapping suggests a high degree of spatial conservatism at the genus rank. Our dated ultrametric tree suggests an origin for Urticaceae in Indomalaya during the mid-Cretaceous, followed by establishment in the Laurasian boreotropical flora and subsequent dispersal to the neotropics and Africa.

Once classified within an evolutionary framework we believe that the Urticaceae represent a valuable study system in evolutionary biology for investigating transitions across biomes, the drivers of floral trait evolution, and intrinsic speciation mechanisms.

*New tribes*: —Leukosykeae, Myriocarpeae

*New genera*: —*Muimar, Pouzolziella*

## Introduction

According to the last conspectus (Friis, 1989) of the family, the Urticaceae Juss. comprise ca 900 species grouped into 48 genera. With the inclusion of the Cecropiaceae, as supported by several studies (see taxonomic history below) and a revised assessment of species numbers the Urticaceae currently comprises 54 genera and 2,625 species (Christenhusz & Byng, 2016) making it a large flowering plant family (Willis, 2017). Approximately two-thirds of the species diversity of the family is accounted for by two species-rich genera, *Pilea* with ca 715 spp (Monro, 2004) and *Elatostema* with ca 800 spp (Fu, Wei & Monro, pers obs). Urticaceae are found in all major plant biogeographic regions and biomes except for Antarctica. Greatest species and genus diversity is observed in the wet and moist tropics and subtropics at elevations between 500 and 2000 masl in mountainous landscapes, especially those associated with karst. Urticaceae include hemi-epiphytes (*Poikilospermum*), herbs (Boehmerieae, Elatostemateae, Parietarieae, Urticeae), vines (Urticeae), shrubs (Boehmerieae, Elatostemateae, Parietarieae, Urticeae) and trees (Boehmerieae, Cecropieae, Urticeae). The tribes can be crudely divided into disturbance-adapted taxa that are common in early, to mid-succession (including riparian) vegetation and agricultural landscapes (Boehmerieae, Cecropieae, Parietarieae, Urticeae), and deep-shade climax vegetation specialists (Elatostemateae) (Wu et al., 2018). The main economic uses of the family are as a fibre (*Boehmeria nivea, Dendrocnide* spp., *Girardinia diversifolia, Obetia tenax, Scepocarpus spp.* (Wells et al., 2021), *Touchardia latifolia, Urtica dioica* (Sfiligoj Smole et al., 2019), with minor medicinal (Bansal et al., 2012) (Lin et al., 1998) (Modarresi Chahardehi et al., 2010), and occasional nutritional use (Diazgranados et al., 2020). *Cecropia,* locally dominant in low to mid elevation early succession forest in the Neotropics represents an important food source for several mammal and bird species of importance for habitat conservation (sloths, toucans, bats) (Charles-Dominique, 1986; Dejean et al., 2012; Lobova et al., 2003)

Urticaceae flowers begin development as bisexual, unisexuality being achieved by the much slower pace of development of one of the sexes, whose vestiges are retained as staminodes or pistilodes. The former may play a role in achene dispersal within the Elatostemateae. Within the tribe Parietarieae and Forrskaoleae, however, there is a notable trend to bisexuality. The Parietarieae include both truly bisexual flowers (*Parietaria officinalis*) and flower-like inflorescences. For example, in *Forrskaolea,* pistillate and staminate flowers are borne in reduced flower-like inflorescences, the male flowers being reduced to a single stamen and tepal in the process. Within all other tribes, flowers and most inflorescences are unisexual, or where inflorescences are bisexual, they do not form flower-like configurations.

The Urticaeae have ‘dry’ stigmas, pollen accessing the pistil via papillae whose morphology does not vary across the family and which appear almost identical to those found in *Arabidopsis* (Edlund, 2004). There is, however, great variation in pistil morphology (Fig. 1), and this has played an important role in genus delimitation (Blume, 1857;Weddell, 1854, 1856, 1869). For example, the sister tribes Elatostemateae and Urticeae are characterised by pin-cushion shaped pistils over which the stigma papillae are uniformly spread; in contrast most members of the Boehmerieae mostly have extended pistils over which stigma papillae may be uniformly distributed or restricted to a specific surface; both types of pistil are exhibited in the Parietarieae. This contrasts with relatively uniform male flower- and stamen- morphology across the family, where the male sepal is characterised by a sub-apical appendage that may be equal in length to the sepal or reduced to a ridge The ovaries are functionally unicarpellate but begin development bicarpellate, occasionally giving rise to bi-ovulate gynoecia. The fruit are achenes which are usually associated with the vestiges of the perianth or a modification of the pedicel or bracts that in many cases are fleshy and/ or brightly coloured.

**Figure 1.**
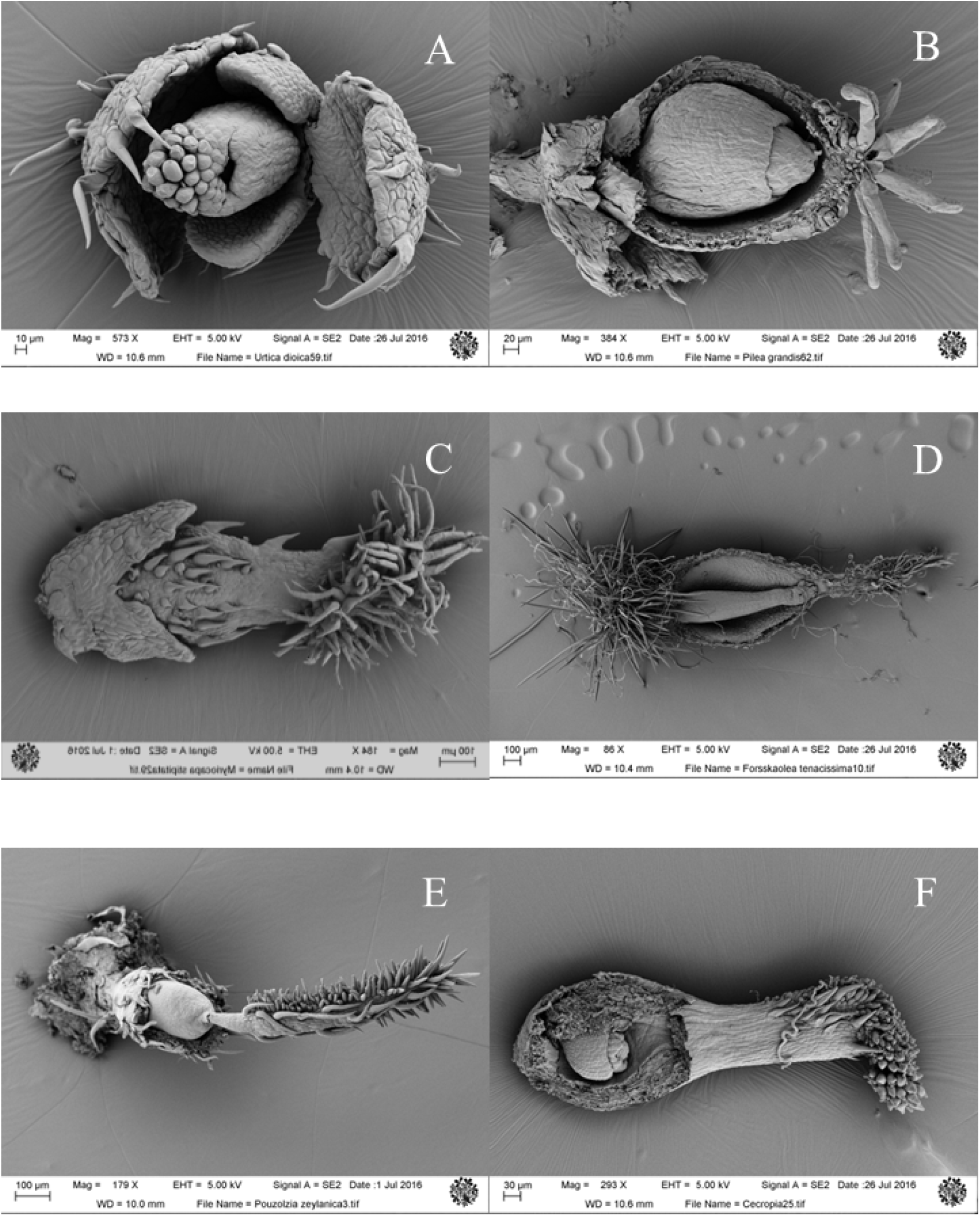
Variation in stigma morphology across the Urticaceae, A) Fleurya aestuans, Urticeae; B) Pilea grandis, Elatostemateae; C) Myriocarpa stipitate, Myriocrapeae; D) *Forsskaolea* tenacissima, Parietareeae; E) Pouzolzia zeylanica, Boehmerieae; F) *Cecropia* obtusifolia, Cecropieae. Images made by Jia Dong and Louis Ronse De Craene

### History of Urticaceae circumscription

The main revisions of the family are summarised in Table 1. The Urticaceae were first described as a coherent group by Jussieu (Jussieu, 1789), as the order Urticae [Urticales], whose circumscription included genera now assigned to the Cannabaceae, Moraceae and Urticaceae together with a small number currently assigned to the Caryophyllaceae (*Pteranthus*), Monimiaceae (*Ambora* [*Tambourissa*]*, Hedycarya*) and Rubiaceae (*Theligonum*). Urticae genera were assigned to three subgroupings based on ovule and embryo disposition. Of these groupings, the second corresponds moderately well to the Urticaceae as currently circumscribed (APG IV)(The Angiosperm Phylogeny Group, 2016), with the exceptions of *Morus, Artocarpus, Cannabis* and *Theligonum.* Later, Gaudichaud ((Gaudichaud, 1830), see Conn & Hadiah (Conn & Hadiah, 2009) for a detailed discussion) treated Jussieu’s Urticae as a family, ‘les Urticées’ which he subdivided into three un-named groupings of unspecified rank, between family and tribe, based on ovule position and embryo anatomy. Gaudichaud’s subdivisions were largely incongruent with Jussieu’s classification. Gaudichaud recognized eleven tribes within these three groupings: i. Elatostemateae, ii. Urereae, iii Boehmerieae, iv. Parietarieae, v. Forsskaoleeae, vi. Cecropieae, vii. Celtideae, viii. Cannabineae, ix. Broussonetieae, x. Moreae, and xi. Ficeae, of which i-vi correspond to the Urticaceae as currently circumscribed (APG IIV). As was the case with Jussieu’s classification, Gaudichaud included several genera that are now assigned to other families (Gunneraceae, Piperaceae, Lacistemataceae and Gnetaceae).

**Table 1.**
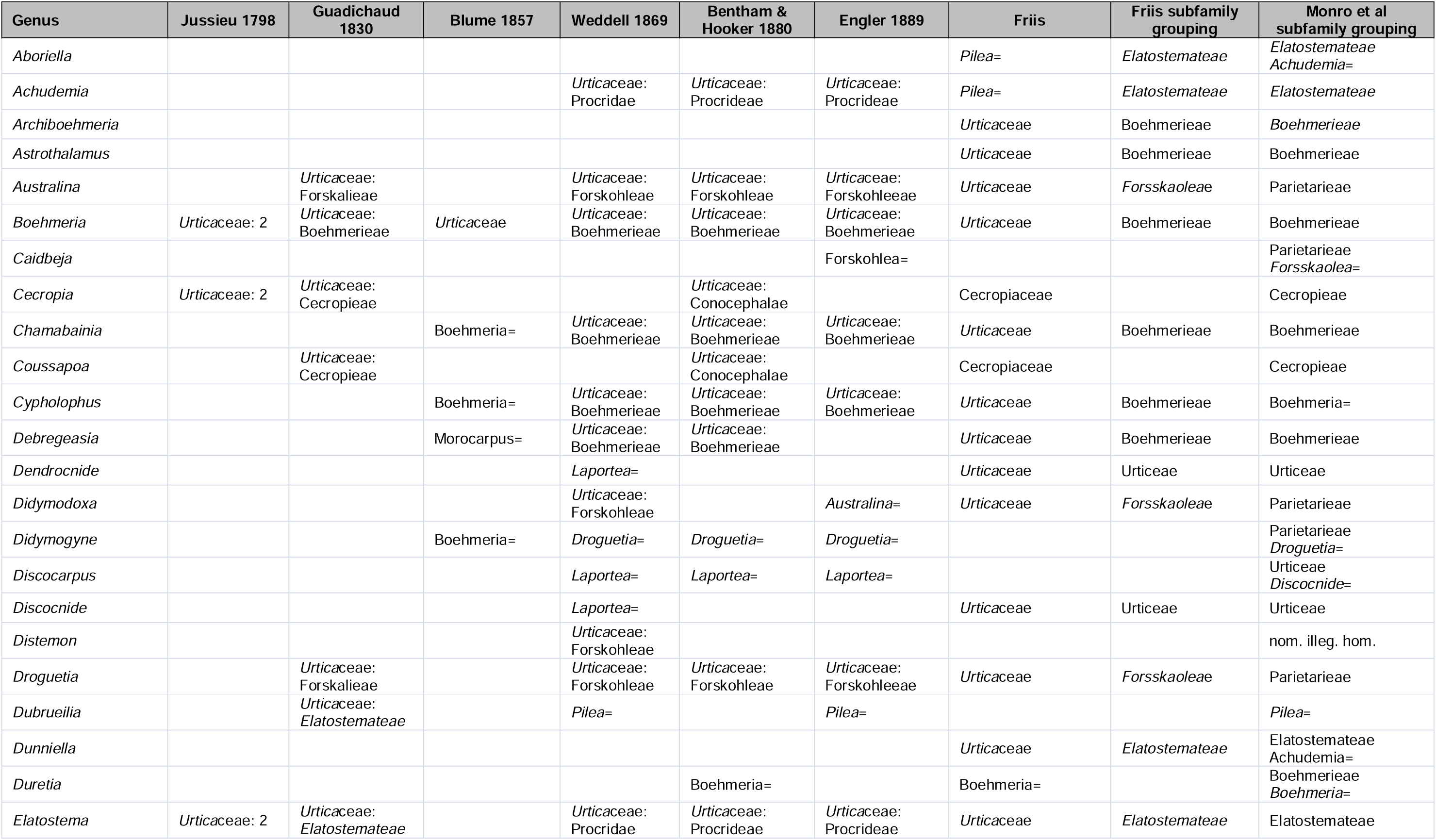
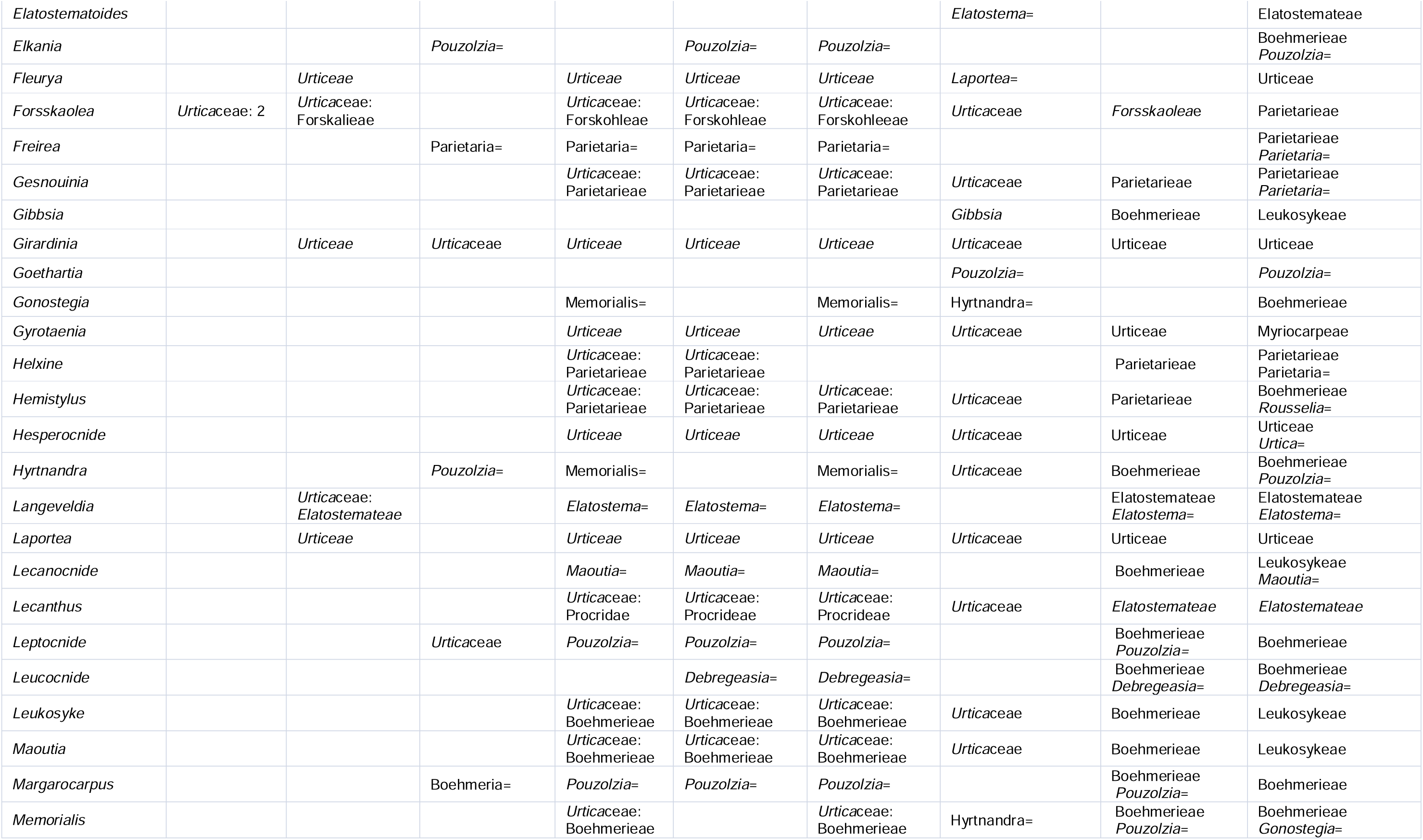
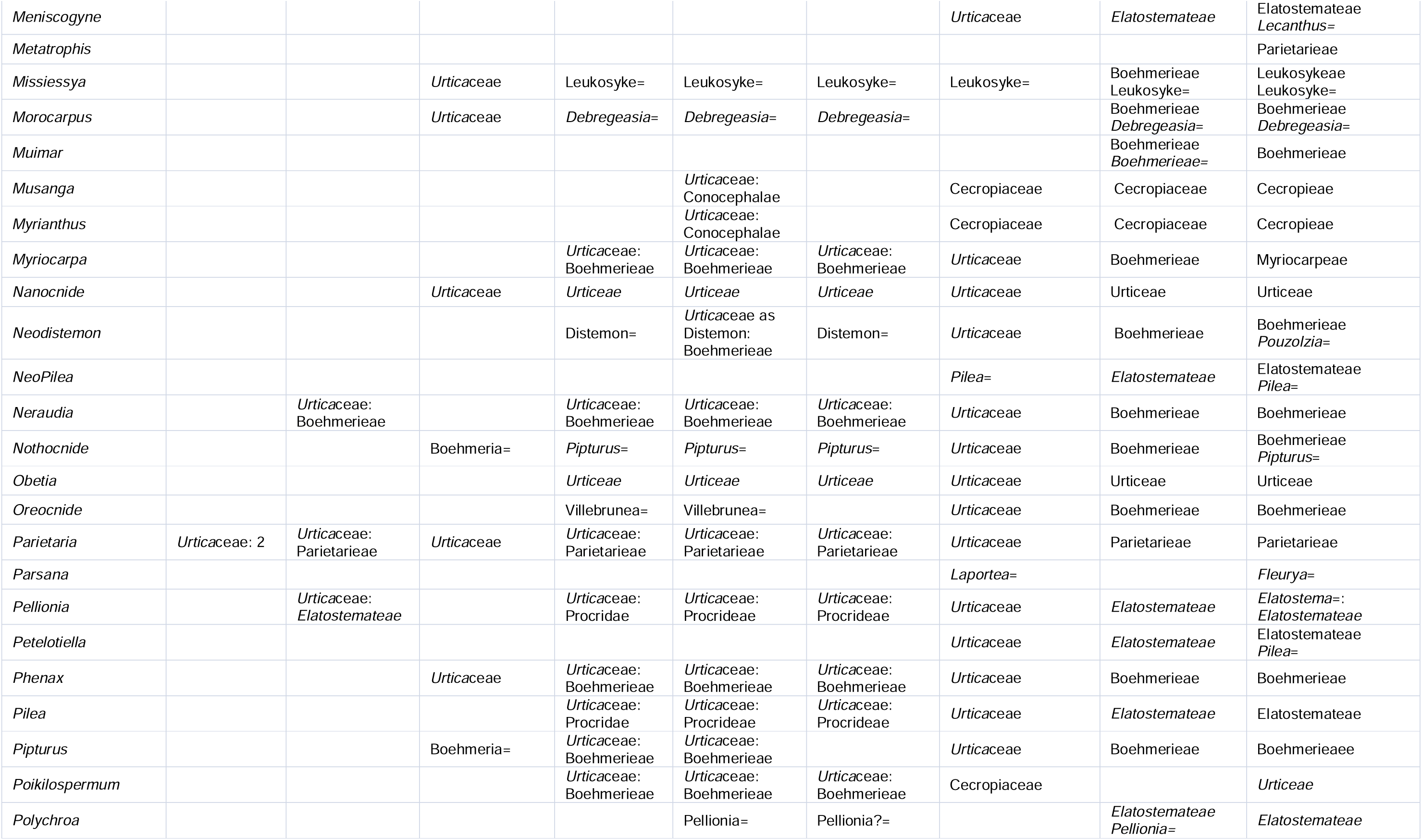
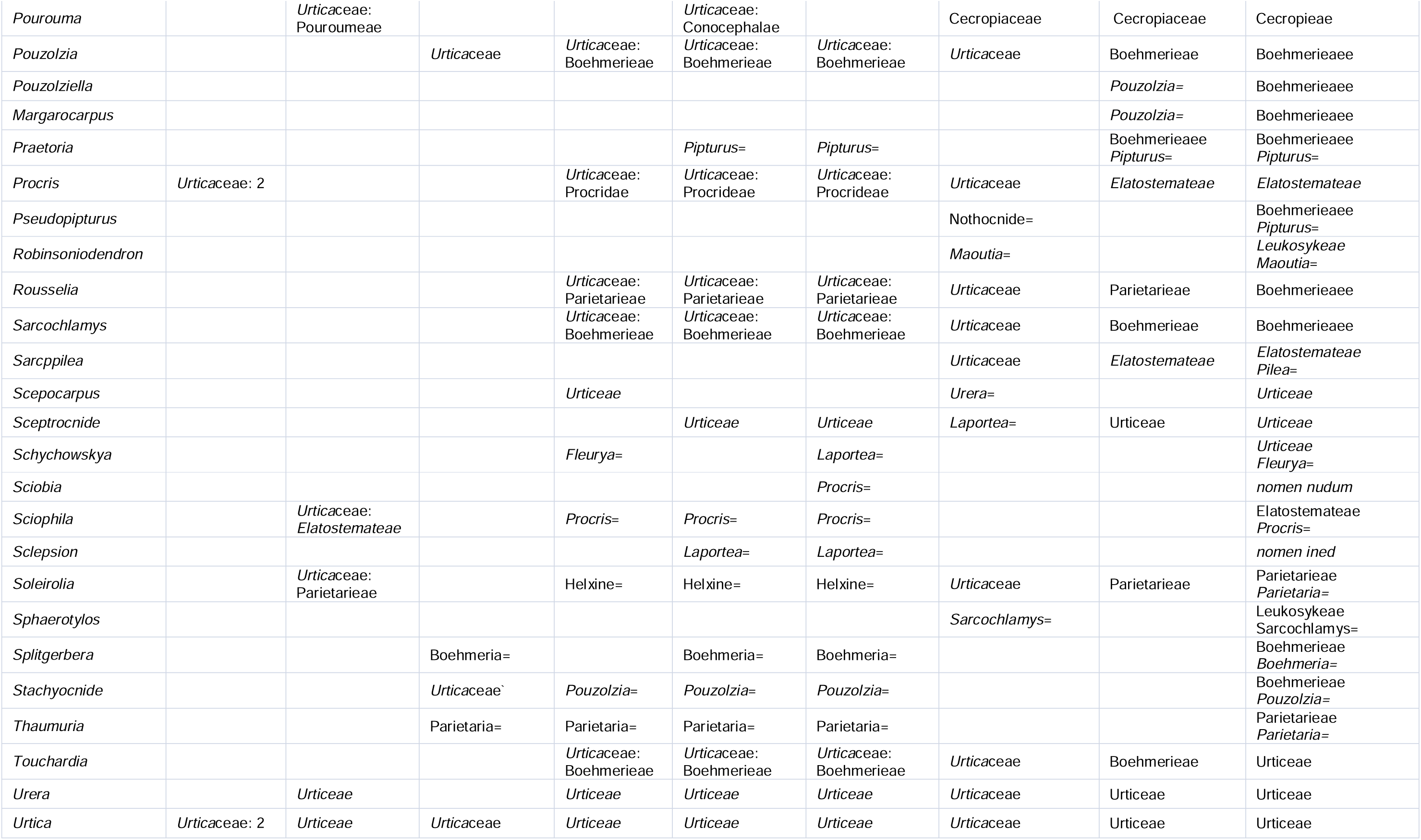
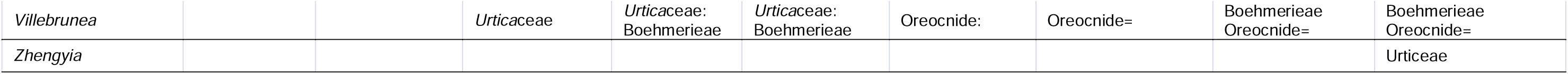
Synopsis of genera and their classification within the Urticaceae.

Between 1854 and 1869 Weddell published three revisions of the Urticaceae (1854, 1856, 1869). These remain, to this day, the most comprehensive, detailed and thoughtful discussions of the family and its circumscription. Weddell (Weddell, 1856) divided the Urticaceae into five tribes (Urereae, Procrideae =Elatostemateae, Boehmerieae, Parietarieae and Forsskaoleae) based on the number of divisions in the staminate and pistillate perianths, the presence or absence of stinging hairs, the degree of fusion and symmetry of the pistillate perianth, stigma morphology and the prominence of bracts within the inflorescence. Weddell’s circumscriptions were congruent with both Jussieu’s (Jussieu, 1789) and Gaudichaud’s (Gaudichaud, 1830), the main difference being that he restricted the family circumscription to the first five of Gaudichaud’s tribes, Elatostemateae to Forrskaoleeae, thereby excluding the Cecropieae. Weddell’s 1854 revision ((Weddell, 1854)) recognized 34 genera and renamed the Elatostemateae tribe Lecantheae (Conn & Hadiah, 2009). In 1856 (Weddell, 1856) Weddell recognized 39 genera, maintained the same five tribes and again changed the name of the Elatostemateae, this time to Procrideae. In 1857 ((Blume, 1857)) Blume published a parallel revision of the Urticaceae in ignorance of Weddell’s 1856 classification and this resulted in a number of now superfluous generic names (see the footnote to the title page of Blume 1857 (Blume, 1857)). Blume did not include a tribal or equivalent classification of the genera and like Weddell, excluded the Cecropieae from his circumscription of the family. In 1869 (Weddell, 1869) Weddell published his final revision of the family in DeCandolle’s *Podromus systematis naturalis regni vegetabilis,* increasing the number of genera to 43 but maintaining his 1856 tribal arrangement.

Except for the subtribes, it is very much Weddell’s classification (Weddell 1856), characterised by five tribes and the exclusion of the Cecropieae, that has been followed by subsequent authors for the rest of the 19th and 20thC (Bentham & Hooker, 1883) (Engler, 1894) (Berg, 1973, 1978a) (Friis, 1989)(see Table 1). Friis (Friis, 1989) followed the same subdivisions but included the diagnostic characters of pistillode ejecting the achene or not and of the pistillode pubescent or not. The exception being Corner (Corner, 1962) and Chew (Chew, 1963) who maintained the Cecropieae within the Urticaceae. Early phylogenetic analysis of morphological characters (Humphries & Blackmore, 1989) (Fig. 14.1) also suggested a monophyletic grouping of Urticaceae, *Poikilospermum* and Cecropieae (as Cecropiaceae) but it was not until the 21stC, and the analyses of DNA sequence data that authors again began to question the exclusion of the Cecropieae (Datwyler & Weiblen, 2004; Hadiah et al., 2008; Monro, 2006; Wu et al., 2013) and in the 2003 APG II, after nearly 180 years, the Cecropieae returned to their position within the Urticaceae.

[Add in review of molecular phylogenetic contribution relevant to taxonomy]

From Weddell’s treatments onwards, a number of genera have remained problematic with respect to their tribal placement. These include *Myriocarpa,* which was tentatively placed within the Boehmerieae by Weddell (Weddell, 1869); *Touchardia,* whose position in the Boehmerieae has been questioned by Friis (Friis, 1989), *Poikilospermum,* variously included (Bentham & Hooker, 1883; Engler, 1894; Weddell, 1869) and excluded (Berg, 1978a; Friis, 1989) from the Urticaceae, and *Metatrophis,* initially ascribed to the Moraceae (Brown, 1935)(Brown 1935) but which Florence (Florence, 1997) proposed should be included in the Urticaceae. These have to some extent been resolved through the analysis of DNA sequence data (Datwyler & Weiblen, 2004; L. Fu et al., 2022; Hadiah et al., 2008; Kim et al., 2015a; Monro, 2006; Sytsma et al., 2002; Wells et al., 2021; Wu et al., 2013). In addition, approximately 1/4 of the genera (*Urera, Meniscogyne*, *Elatostema, Elatostematoides, Laportea, Pellionia, Procris, Achudemia, Arboriella, Pouzolzia, Boehmeria, Parietari* a and *Leukosyke*) have been shown to be para- or polyphyletic (Chen, 1982; Kim et al., 2015b; Monro, 2006; Schüßler et al., 2019; Tseng et al., 2019; Wells et al., 2021; Wu et al., 2013).

### The taxonomic challenge

Weddell (Weddell, 1856: 4) wrote in the introduction to his monograph of the family: “… there is no plant family where the species are more variable, none, in other words, where it is more difficult to recognise the species and even the genera by a simple inspection of appearance…”. This, to some extent, has been reflected by the abundant homoplasy amongst the characters traditionally used to delimit taxa (Kim et al., 2015c; Tseng et al., 2019; Wells et al., 2021; Wu et al., 2015) in the family but despite this Weddel’s 19thC classifications (Weddell, 1854, 1856, 1869) the tribal classification remains little changed to this day.

The Urticaceae have been consistently recovered as sister to the Moraceae within a subclade of the Rosales that also comprises Ulmaceae and Cannabaceae (Stevens, 2017). The first published phylogeny of relationships within the Urticaceae was that of Friis (Friis, 1989: Fig. 16.9A). This suggested a partially resolved phylogeny with five tribes arranged in an unresolved trichotomy: the Urticeae united by stinging hairs, the Elatostemateae (as Lecantheae) by the staminodes actively ejecting the achene from the fruit and the Boehmerieae, Parietarieae and Forsskaoleae united by the presence of a lanate or pilose pistillode. Sytsma *et al*. (Sytsma et al., 2002) published the first phylogeny based on analyses of DNA sequence data which suggested the inclusion of *Poikilospermum* in the Urticaceae. This was followed by Datwyler *et al*. (Datwyler & Weiblen, 2004) whose analysis of DNA data suggested the inclusion of the Cecropiaceae into the Urticaeae, and by Monro (Monro, 2006) who confirmed the inclusion of *Poikilospermum* within the Urticaceae, suggesting its placement within the Urticeae. In 2008, Hadiah *et al*. (Hadiah et al., 2008) confirmed the inclusion of the Cecropiaceae and the placement of *Poikilospermum* and proposed the placement of *Myriocarpa* within the Elatostemateae. In 2013, Wu *et al*. (Wu et al., 2013) with the most comprehensive sampling of genera and genomes to date recovered four monophyletic groups broadly congruent with Friis’s 1989 phylogeny except for the inclusion of the Cecropiaceae as sister to the Boehmerieae, Parietarieae and Forsskaoleae. From 2013 there has been support for an Urticaceae comprising five tribes (Z. Y. Wu et al., 2013: Fig. 2) which corresponds closely to Gaudichaud’s 1830 classification (Gaudichaud, 1830) but differs from those of Weddell (Weddell, 1854, 1856, 1869) and Friis (Friis, 1989) with respect to the inclusion of the Cecropiaceae. Despite the increased taxon sampling by Wu et al. (Wu et al., 2013) and the resolution of some of the basal branches in the phylogenies, many clades were recovered with weak support. For example, Huang *et al*. (Huang et al., 2019) were unable to resolve the relationship between species of *Urera* in the Urticeae clade despite the inclusion of data from four chloroplast regions and increased taxon sampling. Wells *et al*. (Wells et al., 2021), however, were able to resolve this node through the application of Angiosperms353 nuclear sequence data (Johnson et al., 2019a).

**Figure 2.**
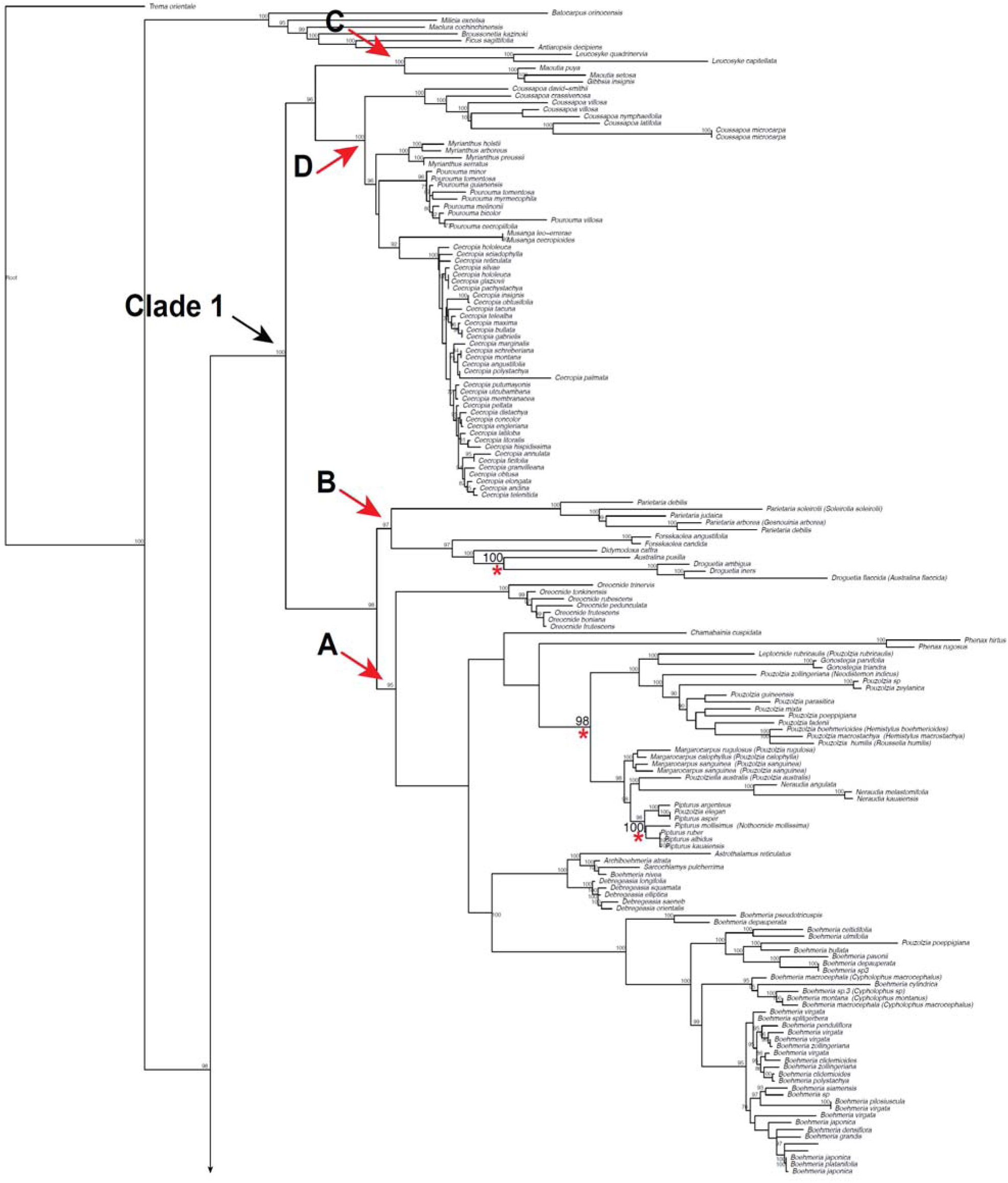
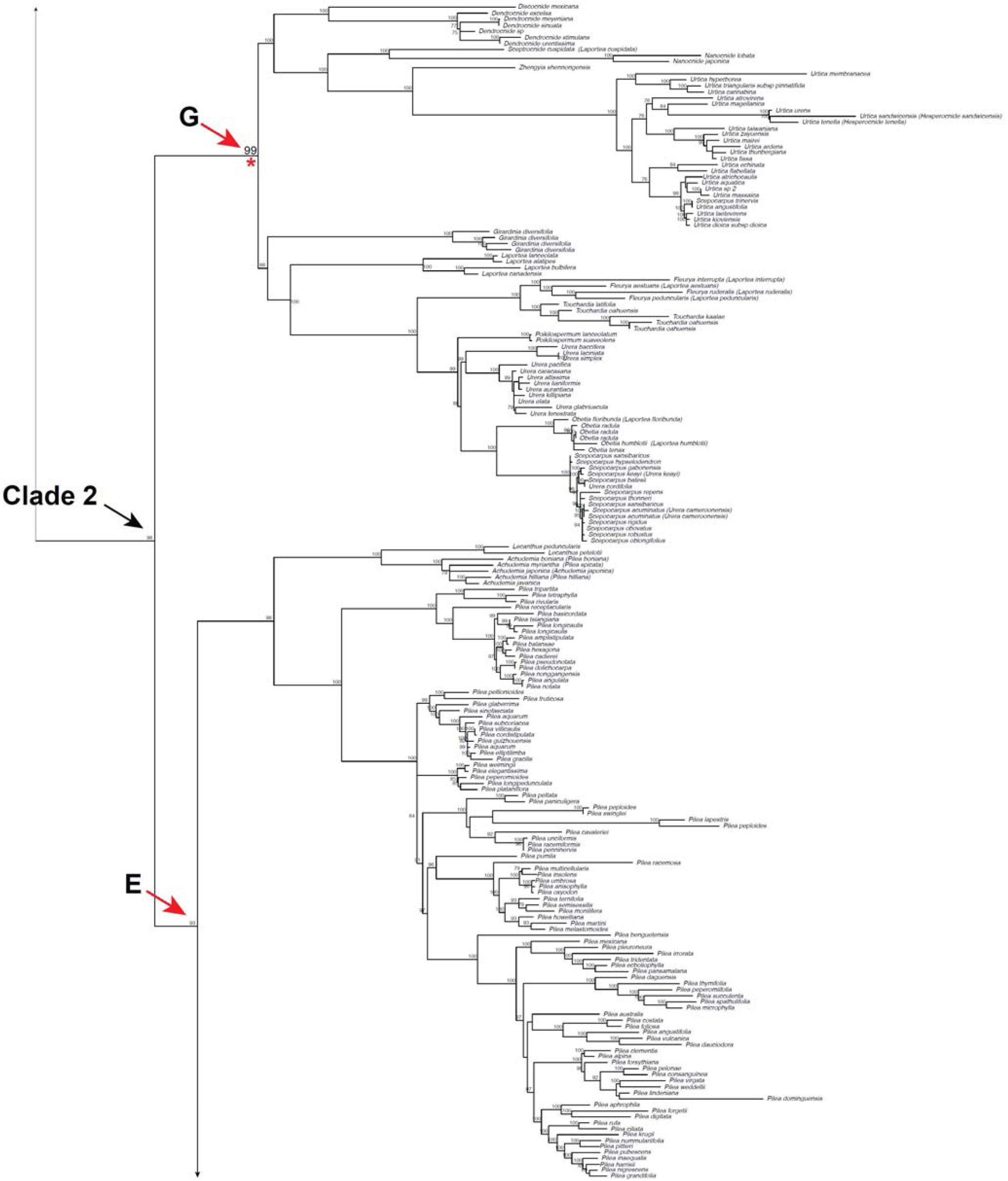
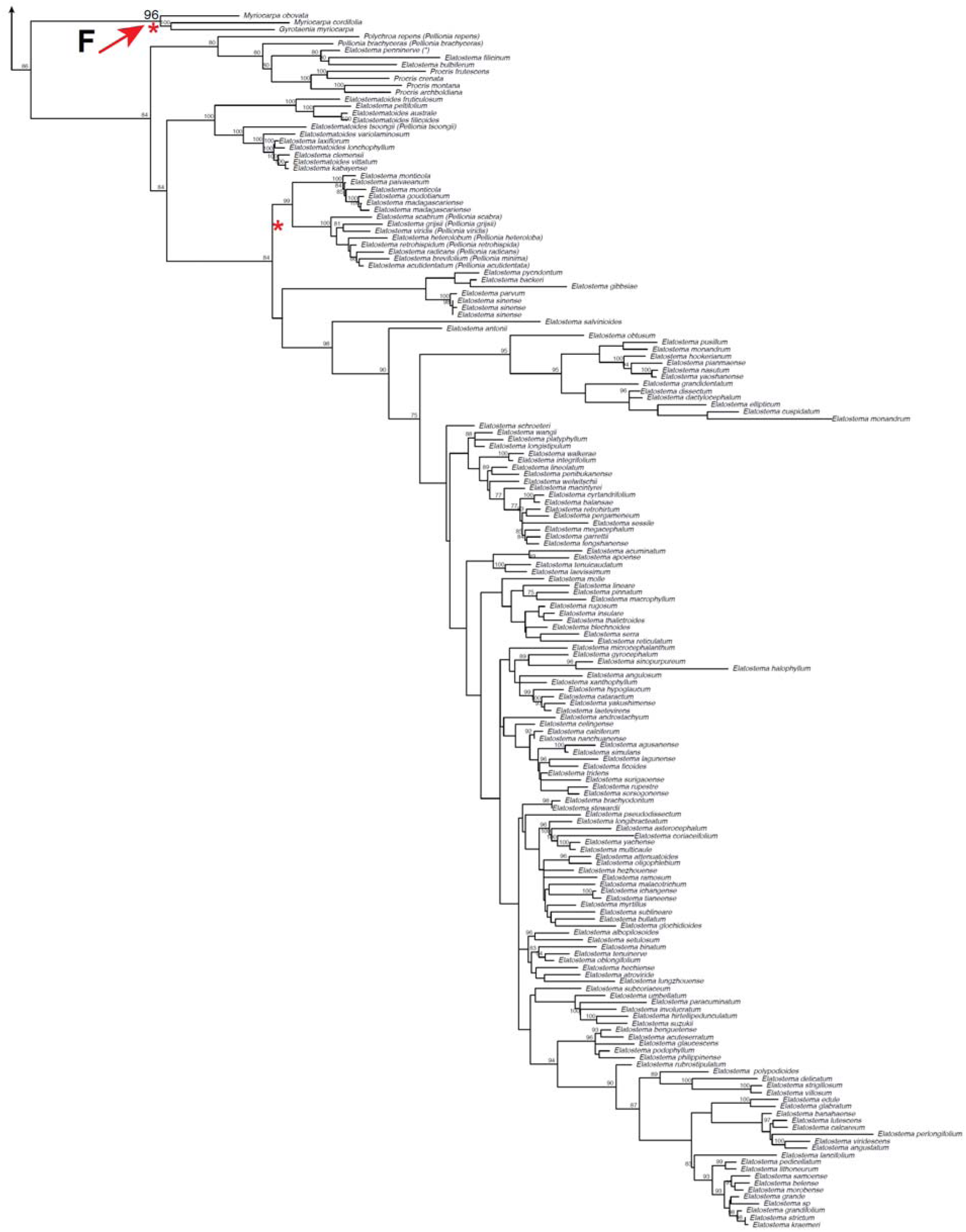
Bayesian analysis of Sanger sequence dataset constrained by the topology of an Astral analysis of the Angiosperms353 HybeSeq low copy nuclear gene dataset. Clades 1 and 2 represent the first order subdivision of the family. Clades A to G represent lower order subdivisions corresponding to tribes. Values above branches are bootstrap percentage above 75. Red asterisks indicated branches with high support where genera were not recovered as monophyletic.

### Morphological diagnosis of the Urticaceae

The exclusion of the morphologically distinct Conocephaloideae (*Cecropia, Coussapoa, Pourouma, Musanga,* and *Myrianthus*) from the Urticaceae simplified the diagnosis of the family although it left the status of the former ambiguous (Corner, 1962). Jussieu (Jussieu, 1789) had originally circumscribed the Urticaceae based on the arrangement of flowers in the inflorescence and the basal and orthotropous (upright) position of the ovule in the ovary. Gaudichaud (Gaudichaud, 1830) employed a similar circumscription delimiting the Urticaceae by the presence of an upright ovule, initially attached at both its ends to the ovary wall and the embryo in the achene being reversed. With the exclusion of the Cecropieae, additional diagnostic characters included elastic and simultaneously reflexing stamens (Friis, 1989), non-amplexicaulous and simple leaf laminae and the presence of cystoliths in the leaf lamina. With the inclusion of the Cecropieae into the Urticaceae (Datwyler & Weiblen, 2004; Hadiah et al., 2008; Monro, 2006; Sytsma et al., 2002; Wu et al., 2013), however, previously used diagnostic characters to distinguish the family from the fig family, Moraceae, (erect orthotropous ovule, absence of amplexicaulous stipules) are lost leaving only the absence of a white latex.

### Morphological basis for higher classification in Urticaceae

Several transitions to perenniality of height (trees, shrubs, lianas), fusion of inflorescence branches, fusions of the perianth, arrangement of the stigma and fleshy fruits have made the application of morphological characters to generic and tribal delimitation problematic (Monro, 2006; Wells et al., 2021; Wu et al., 2018). For example, the absence of a visible perianth tube can be the product of its fusion to the carpel, as in *Astrothalamus* and *Oreocnide*, or a product of its complete reduction, as in *Phenax.* Similarly, some characters, such as caudiform versus capitate stigma types, that are relatively constant within genera, can both occur within the same *Parietaria* inflorescence, or different species of *Poikilopsermum.* The availability of DNA sequence data has therefore been of great help in revising generic and tribal delimitation and of course in evaluating the phylogenetic informativeness of morphological characters.

Gaudichaud, Blume, Bentham and Weddell were generally in agreement with respect to the characters to be used for generic delimitation although they occasionally disagreed as to their interpretation. Weddell (Weddell, 1856) is the most expansive of the authors with respect both to character selection and interpretation, making frequent reference to how decisions were made by himself and contemporary authors (Blume, Bentham, Gaudichaud) in his introductory chapters and the ‘*Obs.*’ sections after each genus and we have relied heavily on this work.

## Aims

The aim of this study was to establish a fully resolved and strongly supported hypothesis of phylogenetic relationship for all genera within the Urticaceae, and to use these to propose a revised and stable tribal classification that could form the basis of a linear sequence for the family. In the process we sought to evaluate the monophyly of the genera and phylogenetic informativeness of the morphological characters that have defined genera and tribes to date.

## Materials & Methods

Our strategy was to use four data sources iteratively and reciprocally, to inform the application of tribal and genus names, an approach successfully applied to the Urticeae (Wells et al., 2021) and Elatostemateae (L. Fu et al., 2022). These data sources were: Sanger sequence data for up to eight non-coding plastid (chloroplast) and one coding (18S) nuclear region (data source 1), used to evaluate the monophyly of genera and inform taxon sampling for Angiosperms353 targeted enrichment (HybSeq) data of up to 350 nuclear genes (data source 2) (Johnson et al., 2019b), which in turn was used to evaluate relationships between genera and assess the phylogenetic informativeness of morphology (data source 3) and geography (data source 4). In light of assessments of their phylogenetic informativeness, the morphological and geographical data were then used to redelimit the genera in conjunction with a nomenclatural review, which was subsequently translated into taxonomic actions (see Taxonomy section).

### Taxon sampling

For all data sets we sampled all the genera recognized by Friis (Friis, 1989) as belonging to the Urticaceae and by Berg (Berg, 1978a) as belonging to Cecropiaceae. In addition, we included genera described, or allocated to the family since that time, *Haroldiella* J.Florence (Florence, 1997), *Metatrophis* F.Br. (Florence, 1997), and *Zhengyia* T.Deng, D.G.Zhang & H.Sun (Deng et al., 2013). *Metapilea* W.T.Wang and *Parsana* Parsa & Maleki were excluded from this study as their taxonomic status is doubtful (see below) and herbarium material was not available for sampling.

We also sampled putative unrecognized genera recovered in previous studies (Kim et al., 2015c; Tseng et al., 2019; Wells et al., 2021; Wu et al., 2013, 2015). These included *Boehmeria nivea* L., *Laportea aestuans* (L.) Chew, *L. alatipes* Hook. f. *L. bulbifera* (Siebold & Zucc.) Wedd., *L. canadensis* (L.) Wedd., *L. cuspidata* (Wedd.) Friis, *L. floribunda* (Baker) Leandri, *L. interrupta* (L.) Chew, *L. humblotii* (Baill.) Friis, *L. mooreana* (Hiern) Chew, *L. ovalifolia* (Schumach. & Thonn.) Chew, *L. peduncularis, L. ruderalis* (G. Forst.) Chew, *Pellionia tsoongii (Merr.) Merr., Pouzolzia rugulosa* (Wedd.) Acharya & Kravtsova (as *Boehmeria rugulosa* in Wu *et al*. (Wu et al., 2018)), *Pouzolzia sanguinea* (Blume) Merr., *Pouzolzia* calophylla W.T.Wang & C.J.Chen, and *Procris repens* (Lour.) B.J.Conn & Hadiah.

We verified accession determination using type collections housed at BM and K, complemented by images of type collections available through Global Plants (https://plants.jstor.org/), the Muséum National d’Histoire Naturelle, Paris (https://science.mnhn.fr/institution/mnhn/collection/p/item/search), The Linnean Herbarium of the Swedish Museum of Natural History (http://linnaeus.nrm.se/botany/fbo/welcome.html.en), Naturalis Bioportal (https://bioportal.naturalis.nl/?language=en) and the Natural History Museum of Denmark digitalized assets (http://www.daim.snm.ku.dk/search-in-types). As many of the type assignations in these portals are erroneous, we used the original published descriptions (protologues) to verify type status.

#### For Sanger sequencing

For the collection of Sanger sequence data we aimed to sample at least three species from each putative genus. Where a genus was monospecific, a single accession was used. Outgroups were chosen based on a phylogeny of the Rosales (L. B. Zhang & He, 2011) and 14 genera selected from the Ulmaceae, Cannabaceae and Moraceae.

We combined sequence data from previous studies (Wu et al., 2013, 2015) and fully vouchered accessions on NCBI GenBank with newly generated data. We generated additional sequences for 21 ingroup taxa using herbarium collections at BM, HAJB and K, focussing on genera that were absent in previous studies (*Musanga, Cypholophus*), poorly sampled (*Hemistylus, Rousselia, Obetia, Nothocnide, Myrianthus, Gyrotaenia, Phenax*) or problematic in previous analyses (*Boehmeria, Laportea, Pouzolzia, Urera*).

#### Angiosperms353 sequencing

We sampled one accession of all monophyletic groupings recovered in previous studies that could be equated to the rank of genus. This resulted in a total of 111 accessions. *Trema orientale* (Cannabaceae) and six species of Moraceae were selected as outgroups.

### Data generation

#### DNA extraction and amplification for Sanger sequencing

Total genomic DNA was mostly isolated from silica–gel dried or fresh leaves, with some from herbarium material supplied by BM and K. All material was extracted using the CTAB method (Doyle, 1987) with minor modifications. PCR was conducted in a total volume of 25 μL containing 2.5 μL 10× PCR buffer, 2.5 μL MgCl2 (25 mM), 2.0 μL dNTP mixture (2.5 mM), 0.75 μL each primer (10 μM), 0.125 μL Taq polymerase (5 U/μL) (TaKaRa, Dalian, China), 1–2 μL template DNA (containing 5–50 ng genomic DNA), and finally distilled deionized water to give a final volume of 25 μL. The PCR profiles for trnL– trnF, rpll4–rps8–infA–rpl36, matK, rbcL, ITS included an initial denaturation step at 94 °C for 1 min, followed by 30 cycles of 50 s at 94 °C, 1 min at 52 °C (55 °C for ITS), 80 s at 72 °C and a final extension at 72 °C for 10 min. The PCR parameters for 18S were as follows: a 97 °C initial hot start for 4 min, followed by 35 cycles of 94 °C for 30 s, 52 °C for 30 s, 72 °C for 70 s, finished with an extension step of 7 min at 72 °C. The PCR conditions for matR were below: initial denaturation at 95 °C for 3 min, 16 cycles of 20 s at 94 °C, 40 s at 65 °C, and degrade 1 °C per cycle, 90 s at 72 °C. Additionally, 20 cycles of 20 s at 94 °C, 40 s at 50 °C, 90 s at 72 °C, and with a final extension period of 5 min at 72 °C. The purified PCR products were directly used for cycle sequencing with the ABI Prism BigDye Terminator Cycle Sequencing Kit (Applied Biosystems, Foster City, California, USA) following the manufacturer’s recommendations. Both strands of the resulting products were sequenced by using an ABI 377 and ABI 3730xl and automated sequencer ((Applied Biosystems, Foster City, California, USA)).

#### Target-enrichment DNA extraction & library preparation

Targeted sequencing data was generated as part of the Plant and Fungal Trees of Life (PAFTOL) project at RBG Kew. DNA extractions were performed using a modified CTAB protocol (Doyle and Doyle, 1987) and purified using AMPure XP beads (Beckman Coulter, Indianapolis, IN, USA). Purified DNA extracts were run on a 1.5% agarose gel to assess the average fragment size. Samples with very low concentration (not visible on a 1.5% agarose gel), were assessed on a 4200 TapeStation System using Genomic DNA ScreenTapes (Agilent Technologies, Santa Clara, CA, USA). The quality of the DNA was evaluated based on agarose gel and TapeStation images and were then quantified using a Qubit® 3.0 Fluorometer (Life Technologies, Carlsbad, CA, USA). DNA samples with the average fragment sizes above 350 bp were sonicated using a Covaris M220 Focused-ultrasonicator with Covaris microTUBES AFA Fiber Pre-Slit Snap-Cap (Covaris, Woburn, MA, USA) with varied shearing times depending on the DNA fragment size profile, to obtain an average of 350-bp insert sizes. Dual-indexed libraries for Illumina® sequencing were prepared using the DNA NEBNext® Ultra II Library Prep Kit at half the recommended volume, with Dual Index Primers Set 1, NEBNext® Multiplex Oligos for Illumina® (New England BioLabs, Ipswich, MA, USA). Quality of libraries were evaluated on an Agilent Technologies 4200 TapeStation System using High Sensitivity D1000 ScreenTape (Agilent Technologies, Santa Clara, CA, USA). Libraries were subsequently quantified using a Qubit® 3.0 Fluorometer. Equimolar (10 nM) libraries were pooled and a total of 1 μg DNA in each pool was enriched using the target capture kit Angiosperms-353 v1, Catalog #308196; (Johnson et al., 2018) following the manufacturer’s protocol v4 (4.0; http://www.arborbiosci.com/mybaits-manual). Hybridizations were performed at 65 ◦C for 28–32 hrs in a Hybex™ Microsample Incubator and using red Chill-out™ Liquid Wax (Bio-Rad, Hercules, CA, USA) to prevent evaporation. Enriched products were amplified from the bait-bound templates, with KAPA HiFi 2X HotStart ReadyMix PCR Kit (Roche, Basel, Switzerland) for 10 cycles. PCR products were then cleaned using the QIAquick PCR purification kit (Qiagen). Products were quantified with a Qubit™ 3.0 Fluorometer and in some cases reamplified (and repurified) a second time between 3 and 8 cycles due to low DNA concentrations. Final products were run on an Agilent Technologies 4200 TapeStation System using High Sensitivity D1000 ScreenTape to assess quality and average fragment size. Two pools of 25 hybridised libraries each, were multiplexed together and sequenced on an Illumina MiSeq with v2 (300-cycles, 150 bp paired-end reads) and v3 (600-cycles, 300 bp paired-end reads) chemistry (Illumina, San Diego, CA, USA) at the Royal Botanic Gardens, Kew.

#### Sampling of morphological characters

Following Scotland et al.’s (Scotland et al., 2003) discussion on the best way to integrate morphological and molecular data in a phylogenetic context we evaluated the congruence of each morphological character state with nodes on a fully resolved ultrametric Hyb-Seq phylogenetic tree with the aim of identifying phylogenetically informative characters. For the purpose of coding morphological characters for use in phylogenetic analyses, Hawkins *et al*. (Hawkins et al., 1997) demonstrated that morphological characters and character states should be distinguished and treated separately, which we did using a two-step process whereby, (i) a comparative study of organismal variation was used to define discrete characters, and (ii) those characters were partitioned and coded as different states that were assigned to the analysed taxa. The character states were then mapped onto a fully resolved ultrametric phylogenetic tree and their distribution across the tree used to classify them as informative or not at any particular rank according to Scotland (Scotland, 1992).

Due to the large number of Urticaceae ingroup taxa for which character coding was required, we used a two-stage iterative approach to morphological sampling. The first stage comprised the evaluation of the phylogenetic informativeness of a ‘long list’ of 73 morphological characters drawn from the discussions of Weddell (Weddell, 1856), Friis (Friis, 1989), Monro *et al*. (Monro et al., 2015), Wells *et al*. (Wells et al., 2021, Urticeae), Kim *et al*. (Kim et al., 2015a, Urticeae), Treiberg *et al*. (Treiber et al., 2016, Cecropieae), Tseng *et al*. (Tseng et al., 2019, Elatostemateae) and Wu *et al*. (Wu et al., 2018) and the first authors own field and herbarium experience of the family. ‘Long list’ characters evaluated, and their states are listed in Table S1. Phylogenetic informativeness was assessed visually by reconstructing ancestral states (see ‘Ancestral state reconstructions’ below) across the HybSeq ultrametric (dated) tree (Wells et al., 2021). We selected characters which were informative for at least one node equivalent to the rank of tribe in the phylogeny. Once evaluated in this way, we selected 21 morphological characters for evaluation on the combined Sanger - Target-enrichment data tree, to which we added an additional morpho-ecological character, ‘*fruit fleshy’* versus ‘*fruit not fleshy*’ (Table 2, character #21) to reflect a bird facilitated dispersal syndrome (Bolmgren & Eriksson, 2005).

**Table 2.**
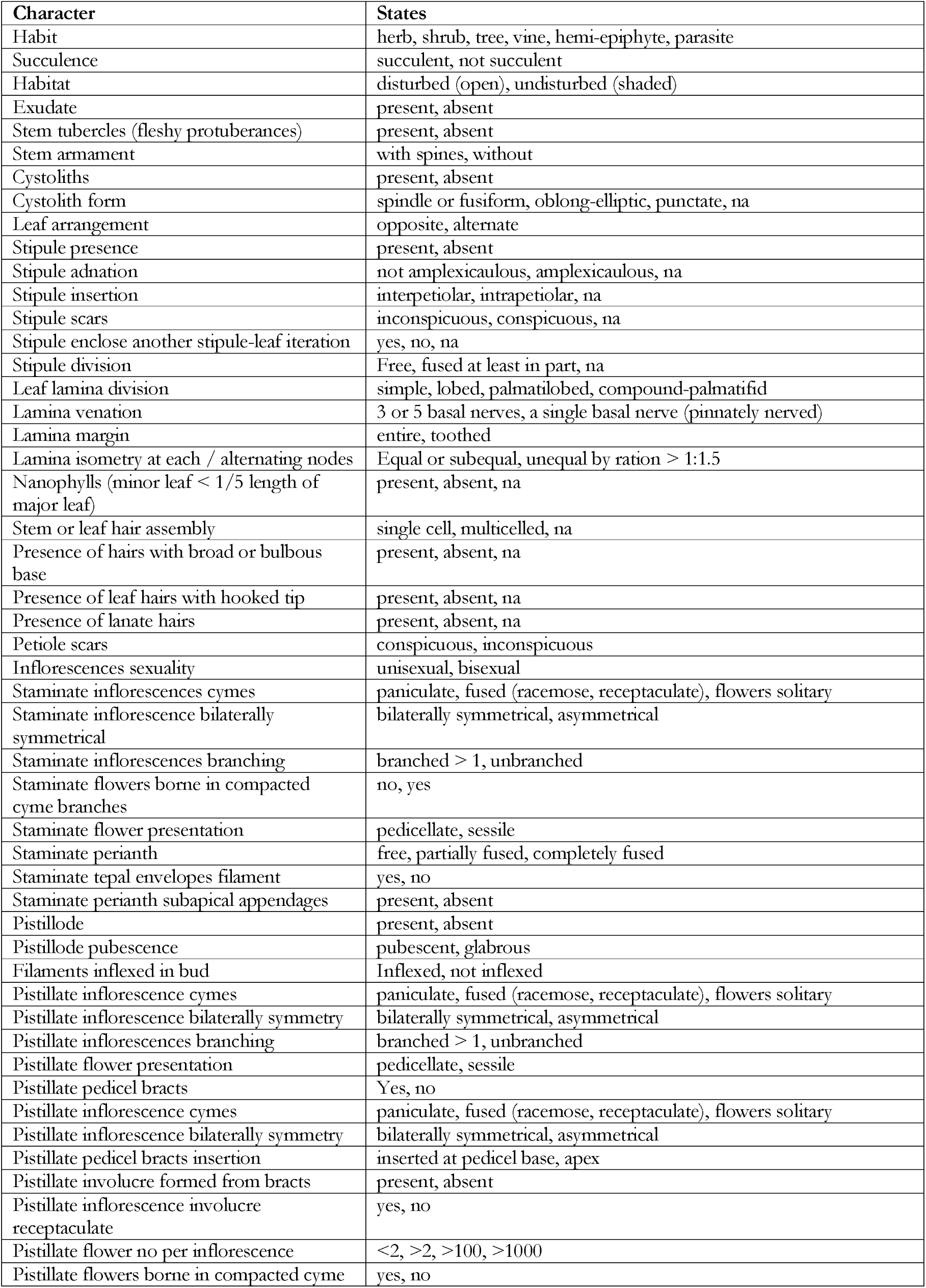
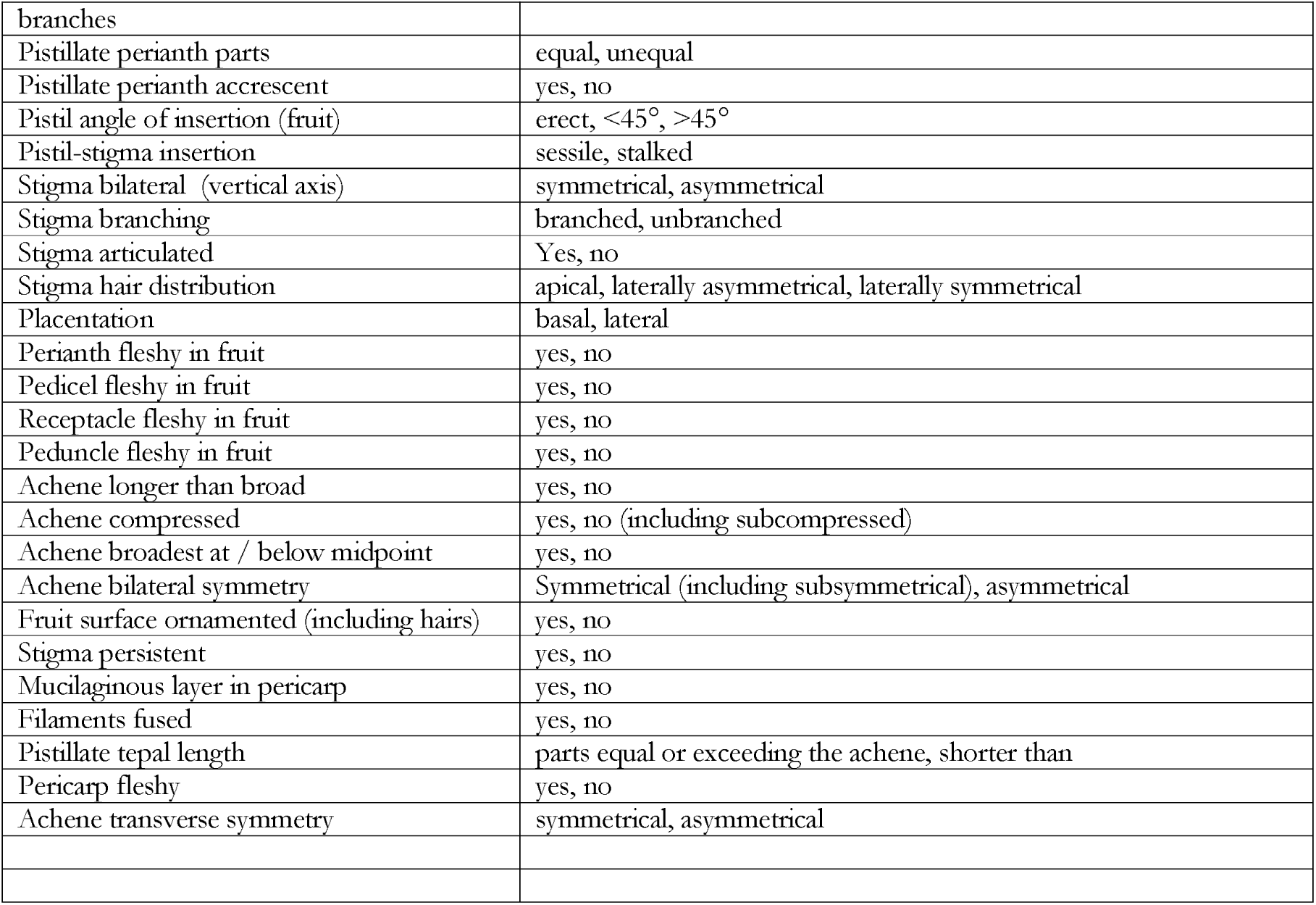
Morphological characters and their states.

#### Sampling and analysis of geographical characters

Phylogenetic relationships have been consistently shown to be closely aligned with geography within the Urticaceae (Wells et al., 2021; Wu et al., 2013, 2018), and for this reason we included geographic distribution as a phylogenetic character in our analysis. We used the following Ecozones as defined by Olson & Dinerstein (Olson & Dinerstein, 2002), Afrotropic (Af), Australasia (Au), Indomalaya (In), Nearctic (Na), Neotropic (Nt), Pacific Islands (Pf), and Palearctic (Pr). Distribution data was gathered from specimen labels. A Dispersal-Extinction-Cladogenesis (DEC) analysis was carried out using BioGeoBears (Matzke, 2013), allowing a maximum of two areas per node, on 32 CPU threads.

### Phylogenetic analyses

#### Sanger sequence alignment and phylogenetic analysis

Sanger sequences were initially aligned using MAFFT with manual adjustments made in Bioedit v7.2 (Hall, 2011)

#### Estimates of support

We adopted the same criteria of in BI, ML and MP analyses as Tseng & al. (Tseng et al., 2019): “For MP and ML analyses, 70–79%, 80–90%, and 90–100% bootstrap supports were considered as weak, moderate, and strongly supported, respectively, and values lower than 70% were considered as providing insufficient support. For BI analyses, the posterior probabilities of <0.9, 0.9–0.94, 0.95–0.99, and 1.0 were considered as providing no, weak, moderate, and strong support, respectively”.

#### HybSeq sequence assembly and phylogenetic analysis

We trimmed reads using Trimmomatic v.0.36 (ILLUMINACLIP: TruSeq3-PE.fa:2:30:10 HEADCROP:3 LEADING:30 TRAILING:25 SLIDINGWINDOW:4:25 MINLEN:20) (Bolger et al., 2014) and assembled them with HybPiper 1.2, which produces gene-by-gene, reference-guided *de novo* assemblies (Johnson et al., 2016), using the reference described by Johnson & al. (Johnson et al., 2019b). To ensure good alignment across the entire family, all analyses were carried out on exonic sequences only, the default output for HybPiper. For each gene, we first discarded sequences shorter than 25% of the average length for the gene. We aligned each gene with MAFFT v.7.453 (Katoh & Standley, 2013) and used trimAl v.1.4 (Capella-Gutiérrez et al., 2009) to trim columns containing more than 75% gaps. An initial set of gene trees was inferred using FastTree, and these were used to prune outlier sequences using TreeShrink. We then ran trimAl once more using the same parameters as before, mainly to remove any all-gap columns resulting from the removal of outlier sequences. We used IQ-TREE to estimate a maximum-likelihood tree for each gene under the best-fit model, with 1000 ultrafast bootstrap replicates, collapsing nodes with less than 30% support using TreeCollapserCL4 (Hodcroft, 2013). We then used ASTRAL-III, a summary-coalescent method, to estimate a species tree from all gene trees estimating node support using local posterior probability (LPP), which reflects quartet tree concordance. Alignment, trimming, and estimation of gene trees were parallelized using GNU Parallel v.20191122 (Tange, 2018).

#### Combined analysis of Sanger and HybSeq data

We combined the Sanger and HybSeq data sets to generate a supertree as follows. First, Sanger loci were assembled from off-target reads in HybSeq samples using HybPiper, with select sequences from the Sanger data set used as references. Sanger loci were then aligned in three sets: (1) Elatostemateae; (2) Boehmerieae, Urereae, and Urticeae; and (3) Cecropieae. To ensure correct placement of the sub-clades, the Sanger loci assembled from the HybSeq samples were included in all three sets of alignments. Each set of alignments was merged to create a supermatrix, and a maximum-likelihood tree was inferred using IQ-TREE. The resulting three trees were then combined with the HybSeq gene trees, and ASTRAL was run on the entire set to infer a supertree.

### Divergence time estimation

For the ASTRAL HybSeq tree (Fig. 3), branch lengths in substitutions per site were estimated on a supermatrix of all loci using RAxML (Stamatakis, 2014) under the GTR+CAT model, with 100 bootstrap replicates. The resulting trees were used to infer a set of 100 time-calibrated trees using treePL (Smith & O’Meara, 2012). Fossil age constraints were derived from the angiosperm calibration dataset of (Ramírez-Barahona et al., 2020), updated for this study. Specifically, the following minimum age constraints were imposed based on fossils: crown Boehmerieae, 41.2 Ma (Collinson, 1989); stem *Boehmeria,* 23.03 Ma (Takhtajan, 1982); stem *Pilea,* 23.03 Ma (Takhtajan, 1982); stem Urticeae, 48.7 Ma (DeVore et al., 2020). Outgroup constraints included stem Artocarpeae (Collinson, 1989), 64 Ma; stem Dorstenieae, 33.9 Ma (Chandler, 1961); stem Ficeae, 56 Ma (Chandler, 1963); and crown Moraceae, 47.8 Ma (Chandler, 1963). Full details for these age constraints are provided in Supplementary Data X. The maximum age of the tree (Moraceae+Urticaceae crown) was constrained to 81.7–93.3 Ma based on Zhang *et al*. (Q. Zhang et al., 2019). Parameters for treePL were optimized following Maurin (Maurin, 2020). 95% HPD of age estimates were summarized on the maximum likelihood tree using TreeAnnotator.

**Figure 3.**
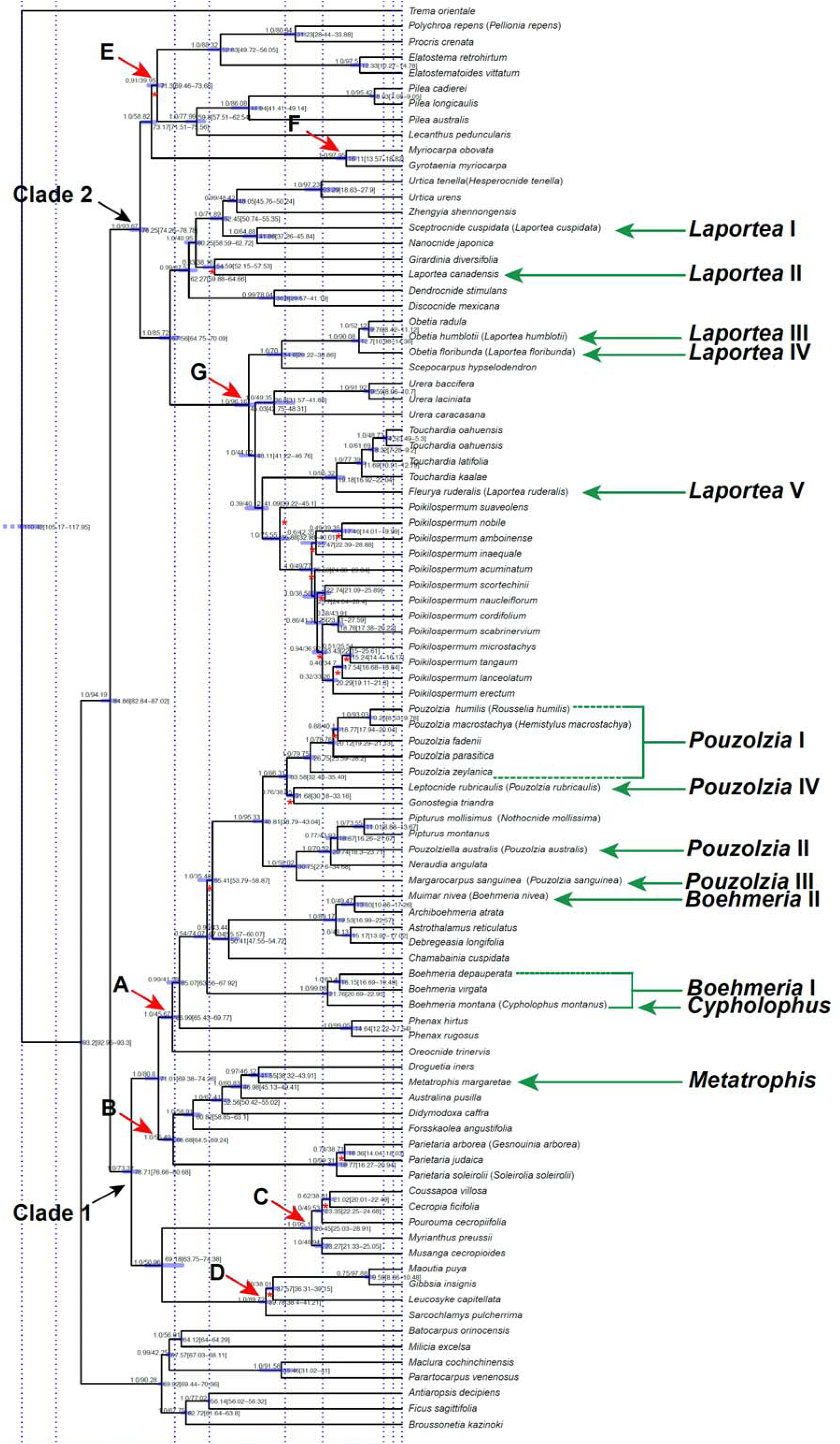
Time-calibrated ASTRAL analysis of Angiosperms353 HybSeq low copy nuclear gene dataset. Clades 1 and 2 represent the first order subdivision of the family. Clades A to G represent lower order subdivisions corresponding to tribes. Estimated average age is provided on the right of each node, with age range in bracket and represented with blue bars. Above branches are local posterior probability values followed by normalised quartet support or percentage gene trees that agree with a branch. Red asterisks below branches indicate the null hypothesis of a polytomy could not be rejected with the polytomy test (p > 0.05).

The HybSeq+Sanger supertree (Fig. 5) was calibrated as follows. To calculate branch lengths in substitutions (for later use in estimating divergence times), the sub-clade Sanger aligments were combined using MAFFT –merge, and then all HybSeq and Sanger loci were combined into a supermatrix. We then used IQ-TREE to infer branch lengths on the existing supertree based on the supermatrix, under the best-fit model for each locus. We then ran treePL on this tree, using the age constraints and parameter optimization method described above.

### Nomenclatural review

With the aim of underpinning a stable classification, a review of the classifications of Jussieu (Jussieu, 1789), Gaudichaud (Gaudichaud, 1830), Blume (Blume, 1857), Weddell (Weddell, 1869), Bentham & Hooker (Bentham & Hooker, 1883), Engler (Engler, 1894) and Friis (Friis, 1989) was undertaken (Table 1). This was accompanied by a review of all the genus names assigned to the Urticaceae and Cecropiaceae, their generotypes and their types. Names were sourced from Jussieu (Jussieu, 1789), Gaudichaud (Gaudichaud, 1830), Blume (Blume, 1857), Weddell (Weddell, 1869), Bentham & Hooker (Bentham & Hooker, 1883), Engler (Engler, 1894), Friis (Friis, 1989) and Deng (Deng et al., 2013), from The World Checklist of Vascular Plants (Govaerts et al., 2021). For each genus name the original publication was located and checked. Generotypes were sourced from Index Nominum Genericorum (Farr & Zijlstra, 1996), verified using the Biodiversity Heritage Library (http://www.biodiversitylibrary.org/) and Botanicus (http://www.botanicus.org/). The validity and priority of names was undertaken in accordance with Articles 6, 20, 21, 38 and 40 of the Shenzhen International Code of Nomenclature for algae, fungi, and plants (Turland et al., 2018).

### Generic and tribal delimitation

As with any rank, tribes provide a means of subdividing a larger group into smaller ones whilst highlighting coinciding patterns and discontinuities in morphology and evolution. They can also make the identification process more manageable. As in Wells *et al*. (Wells et al., 2021) and Fu *et al*. (L. Fu et al., 2022), we aimed to provide a classification in which tribes and genera are morphologically diagnosable and monophyletic. The justification for this is that we want to deliver a classification which supports identification both *in situ* or in biological collections, is predictive and also provides an evolutionary framework that supports the generation and testing of evolutionary hypotheses. We therefore sought to avoid taxa that could only be diagnosed using molecular data or proposing groupings that were recovered as poly- or para- phyletic with strong support. This approach was problematic where genera recovered as para- or polyphyletic were well-defined by unique combinations of morphological characters. For example, the genera *Pouzolzia* and *Laportea* are both fairly easy to recognise based on characters of habit, leaves, perianth, and fruit, but both have been recovered as paraphyletic ((Kim et al., 2015c; Wells et al., 2021; Wu et al., 2018)). In these circumstances we looked for additional morphological characters and used the pronounced phylogeographic signal within the family (Huang et al., 2019; Wells et al., 2021; Wu et al., 2018) to inform our decision-making. Where a taxon recovered as paraphyletic, comprising several monophyletic groupings which co-occur in the same biogeographic domain and were morphologically indistinguishable, then we would consider such a situation intractable and not propose any taxonomic action.

#### Linear sequence of the tribes and genera

A phylogeny is essentially a three-dimensional representation of putative relationships, branches at each node being free to rotate around each other (Haston et al., 2007). Conversion to a two-dimensional sequence is, however, frequently needed for the physical organisation of biological collections, or printed lists in monographs, catalogues, and field guides. By using a systematic arrangement that incorporates basic information of relationships, the resulting arrangement or sequence adds significant value to the uses of the collection (Funk, 2003) and, in theory, stability of the classification, assuming that estimates of relationships are correct. Converting a three-dimensional organisation of information into a two-dimensional one requires some arbitrary decisions around the ordering of terminals at a node. For example, Haston *et al*. (Haston et al., 2007) decided to place first the terminal with the fewer number of taxa in a rotating node.

Because we are proposing a tribal classification, we have opted to align the order of the linear sequence with the existing status-quo (Friis, 1989; Weddell, 1869), whilst respecting the hierarchy of relationships recovered by the phylogenetic analyses. Where arbitrary decisions needed to be made between two genera at a node then we used the approach of Haston (Haston et al., 2007) and (Christenhusz et al., 2011) to place the clade with the smaller number of extant species first, which at least may facilitate the identification-by-matching process in the herbarium.

### Missing sequence data

A single genus sampled for the Sanger sequence data set that is absent from the Angiosperms353 sequence dataset, *Neodistemon* (Boehmerieae). A single genus sampled for the Angiosperms353 sequence data is absent from the Sanger sequence data, *Metatrophis* (Parietarieae) (Appendix 1).

## Results

### Taxon sampling

We obtained Sanger sequence data for a total of 532 ingroup and seven outgroup accessions, and HybSeq data for 82 ingroup and 6 outgroup accessions (Fig.S1. plot of the loci against a phylogram).

### Phylogenetic analyses

#### Sanger sequence data

[Figure 2. Bayesian analysis of the Sanger sequence dataset constrained by the topology of an ASTRAL analysis of the Angiosperms353 HybSeq low copy nuclear gene dataset. Clades 1 and 2 represent the first order subdivision of the family. Clades A to G represent lower order subdivisions corresponding to tribes. Values above branches are bootstrap percentage above 75. Red asterisks indicated branches with high support where genera were not recovered as monophyletic.]

The combined analyses of Urticaceae Sanger sequence data (Fig. 2) divides into two strongly supported (PP=1), clades, Clades I & 2). Clade 1 comprises all of the ingroup taxa from the Cecropiaceae, Boehmerieae, Forsskaolae and Parietariae *sensu* Friis (1989) (with the exception of *Gyrotaenia, Myriocarpa* and *Touchardia*); and Clade 2 comprisies all of the taxa in the Urticeae and Elatostemateae, with the addition of *Gyrotaenia, Myriocarpa, Poikilospermum,* and *Touchardia.* Clades 1 and 2 are subsequently subdivided into five (Clades IA-D1, D2) and three (Clades IIE-G) strongly supported (PP=100) subclades respectively (see Fig.1a-e), each of which can be morphologically delimited (see below).

Within Clade 1 (Boehmerieae, Cecropieae, Forsskaoleae, Parietarieae) we recovered accessions of *Leukosyke, Maoutia* and *Gibbsia* (Clade D) as sister to the Cecropieae (Clade C), with *Maoutia* recovered as polyphyletic with respect to *Gibbsia* with strong support. The Boehmerieae (Clade A) and Cecropiaeae were recovered as moderately supported (90%, 88% respectively), whilst the Forsskaoleae (Clade B) were recovered with weak support (BS 57%), and the Parietarieae (Clade B) excluding *Rousselia* and *Hemistylus*) with strong support (100%). *Sarcochlamys* was recovered sister to *Boehmeria nivea* with strong support (BS100%).

Within the Boehmerieae, *Pouzolzia* was recovered as polyphyletic (*Gonostegia*, Neodistemon, *Rousselia, Hemistylus*) and paraphyletic (*P. australis, P. calophylla, P. elegans, P. poeppigiana, P. rubricaulis, P. rugulosa, P. sanguinea*). *Pipturus* was recovered as polyphyletic (*Nothocnide mollissima, Pouzolzia elegans*). *Boehmeria* was recovered as paraphyletic (*B. nivea, B. virgata*) and polyphyletic (*Cypholophus*).

Within the Forsskaoleae (Clade B), *Australina* was recovered as polyphyletic with respect to *Droguetia* and *Metatrophis.* Within the Parietariae (Clade B), *Parietaria* was recovered as polyphyletic with respect to *Gesnouinia* and *Soleirolia*.

Within the Cecropieae (Clade C), the accession of *Musanga* was recovered within *Cecropia* with strong support (100%), and one of four accessions of *Pourouma* was recovered within *Cecropia* accessions but with weak support (50%).

Within Clade 2, *Gyrotaenia* and *Myriocarpa* (Clade F) were recovered as sister to Elatostemateae (Clade E), with *Myriocarpa* polyphyletic with respect to *Myriocarpa* with strong support (BS 100%). Within the Urticeae (Clade G), *Laportea* was recovered as paraphyletic, forming four groups sister to *Nanocnide* (weak support), *Touchardia* (strong support), within *Obetia* (strong support) and an unplaced grouping (strong support). Within the Elatostemateae, *Procris repens* was recovered sister to a clade comprising all accessions of *Procris* and *Elatostema* penninerve, *E. bulbiferum* and *E. filicinum,* the latter rendering *Elatostema* polyphyletic with respect to *Elaostematoides* and *Procris*.

#### Angiosperms353 low copy nuclear gene data

Urticaceae divide into two strongly supported clades, Clade I) comprising all of the ingroup taxa from the Cecropiaceae, Boehmerieae, Forsskaolae and Parietariae (PP 1.0; QS 73.32), with the exception of *Gyrotaenia, Myriocarpa* and *Touchardia*; and Clade 2) comprising all of the taxa in the Urticeae and Elatostemateae (sensu Friis 1989) (PP1.0, QS 93.67) with the addition of *Gyrotaenia, Myriocarpa, Touchardia* and the unplaced genus *Metatrophis* (Florence 1997).

Clade I divides into four strongly supported subclades corresponding to the tribes Cecropieae (Fig. 3, C, PP 1.0, QS 95.1), Forsskaoleae + part of the Parietarieae (Fig. 3, B, PP 1.0; QS 56.49), Boehmerieae (Fig.3, A, PP 1.0, QS 45.67) and a clade (Fig. 3, D, PP 1.0, QS 89.72) comprising *Gibbsia, Leukosyke, Maoutia* and *Sarcochlamys*. The Parietarieae *sensu* Friis (1989) was recovered as paraphyletic, the genera *Hemistylus* and *Rousselia* being recovered within a clade comprising mostly *Pouzolzia* species (Fig. 3, ‘Pouzolzia I’, PP 1.0, QS 79.75).

Clade 2 divides into three strongly supported subclades corresponding to the tribes Urticeae (Fig. 3, G, PP 1.0, QS 96.16), Elatostemateae (Fig. 3, E, PP 0.91, QS 39.95) and a clade comprising the genera *Gyrotaenia* and *Myriocarpa* (Fig. 3, F, PP 1.0, QS 97.95), the latter clade being recovered as sister to the Elatostemateae (PP 1.0, QS 58.82).

In addition, we found evidence of paraphyly for *Boehmeria, Pouzolzia, Pellionia* and *Laportea. Boehmeria* species were recovered in three positions in the tree, 1) sister to *Archiboehmeria* + *Astrothalamus* + *Debregeasia* (Fig. 3, ‘Boehmeria II’, PP 1.0, QS 89.17), 2) sister to *Cypholophus* (Fig.4, ‘Cypholophus’, PP 1.0, QS 99.08), and 3) sister to *Phenax* (PP 0.99, QS 41.58). *Pouzolzia* was recovered in four positions in the tree, 1) sister to a clade comprising *Neraudia, P. australis, Pipturus* and *Nothocnide* (Fig. 3, ‘Pouzolzia III’, PP 1.0, QS 58.02) within a clade comprising *Neraudia, Pipturus* and *Nothocnide* (Fig. 3, ‘Pouzolzia II’, PP1.0, QS 70.52), 3) sister to *Gonostegia* (weakly supported, Fig. 3, ‘Pouzolzia IV’, PP 0.76, QS 38.05), and 4) in a clade comprising *Rousselia* and *Hemistylus* (Fig. 3, ‘Pouzolzia I’,BS 100%, 100/99.36).

**Figure 4.**
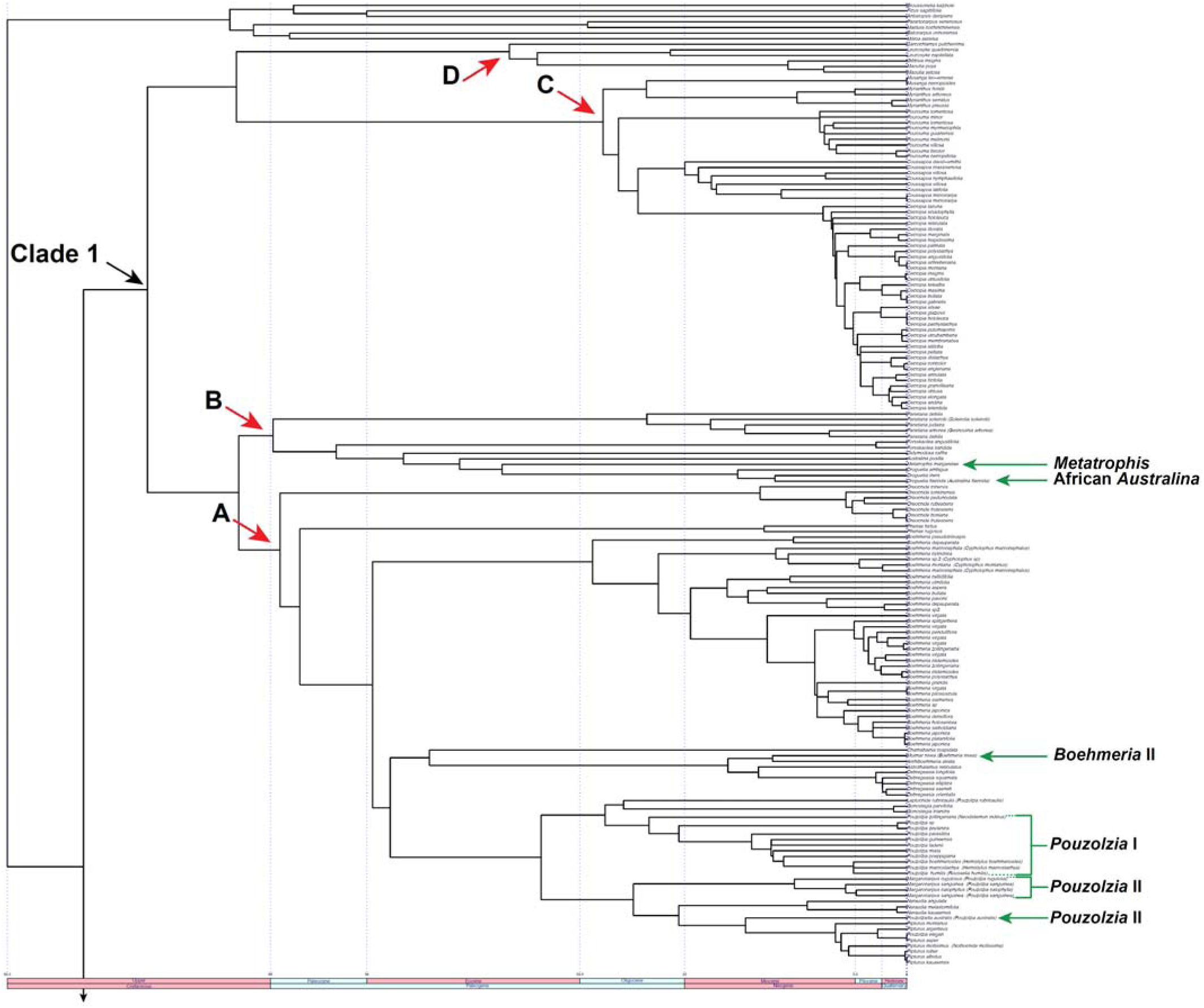
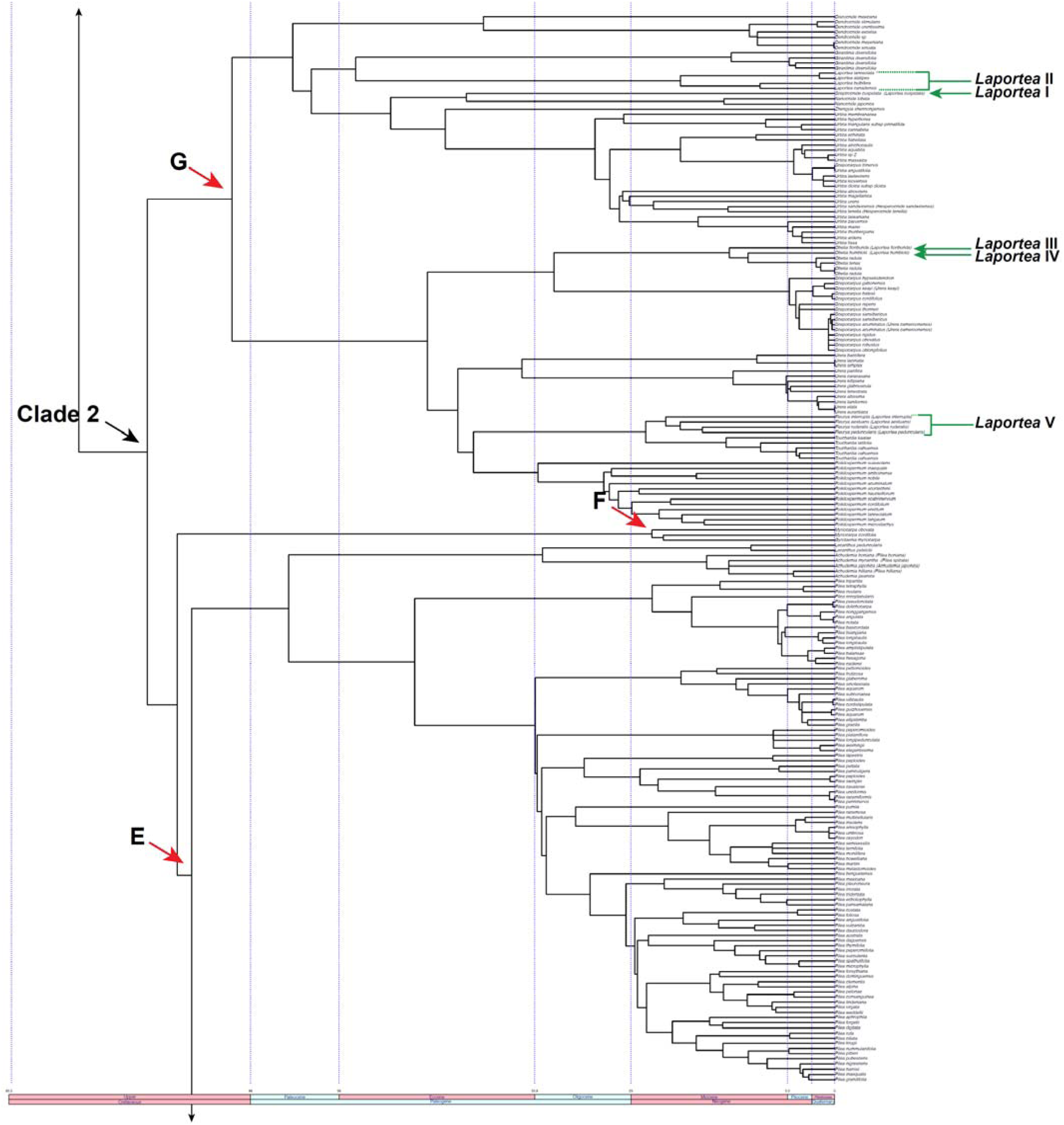
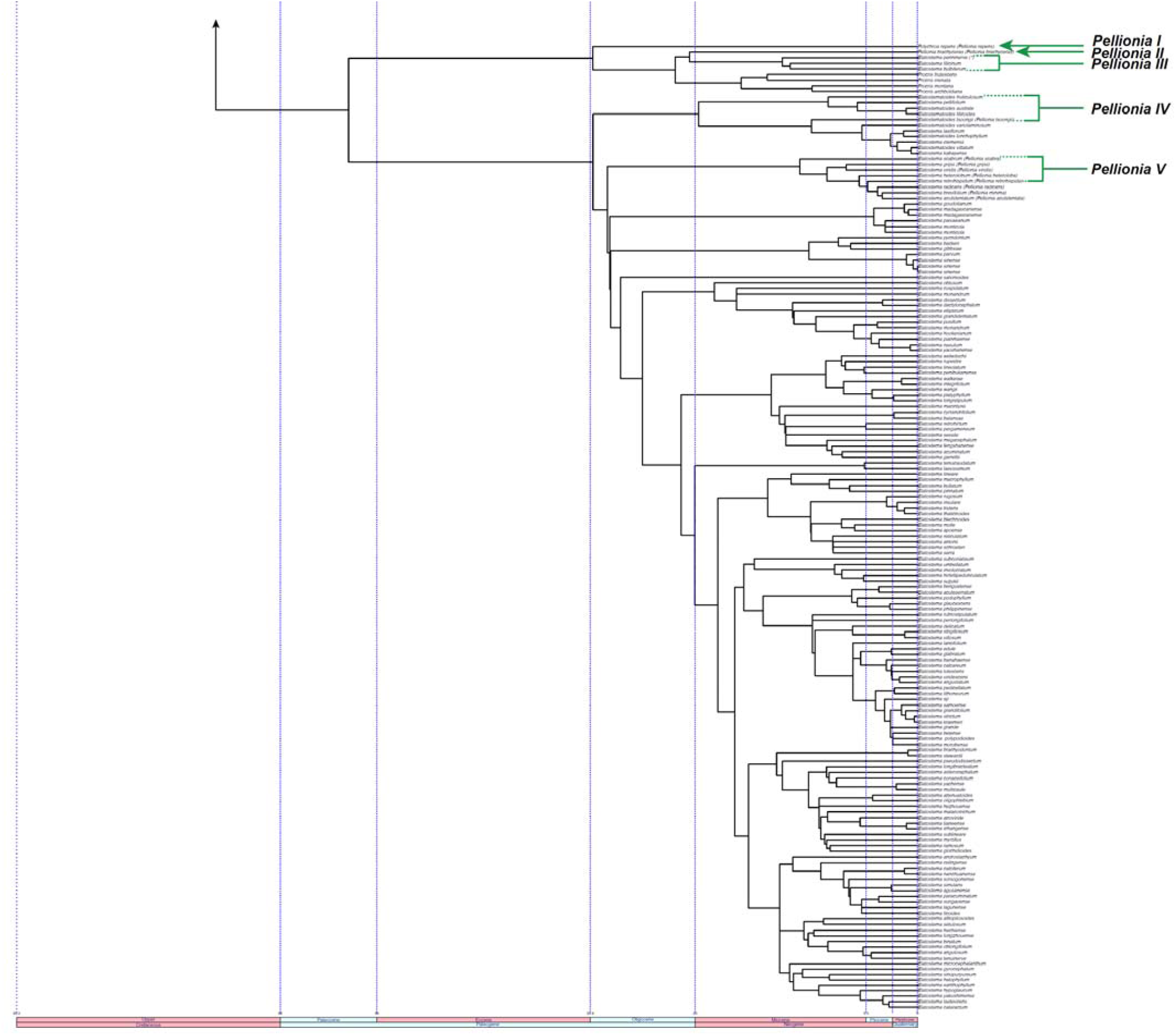
Time-calibrated supertree of Sanger sequence and Angiosperms353 HybSeq low copy nuclear gene datasets. Clades 1 and 2 represent the first order subdivision of the family. Clades A to G represent lower order subdivisions corresponding to tribes.

*Pellionia* was recovered in four positions (Fig.s 2.3, 4.3), 1) within *Elatostema* (Fig. 2.3, BS 100%), 2) sister to *Procris, Elatostematoides, Elatostema* and two un-named clades (Fig. 2.3, BS 800%) (*Pellionia repens*), 3) sister to *Elatostematoides* (Fig. 2.3, BS 100%) (*Pellionia tsoongii*), and 4) sister to *Procris* (Fig. 2.3, BS 100%) (*Pellionia brachyceras, Elatostema penninerve*).

*Laportea* was recovered in five positions in the tree, 1) sister to *Girardinia* (Fig. 3, ‘Laportea II’ PP 0.83, QS 38.16) or a clade comprising *Nanocnide, L. cuspidata, Zhengyia, Hesperocnide* & *Urtica* (Fig.3, ‘Laportea I’ PP 1.0, QS 40.95),, 2) sister to *Nanocnide* japonica (Fig. 3, ‘Laportea I’, BS 1.0, QS 64.88), 3) sister to a clade comprising *Obetia* & *L. humblotii* (Fig. 3, ‘Laportea IV’, PP 1.0, QS 90.08), 4) sister to *Obetia* (Fig. 3, ‘Laportea III’, PP 1.0, QS 52.12), and 5) sister to *Touchardia* (Fig. 3, ‘Laportea V’, PP 1.0, QS 95.32).

We also recovered the unplaced genus, *Metatrophis* sister to *Droguetia* within Clade B (Fig. 3, PP 0.97, QS 46.12).

#### Combined analysis of Sanger and HybSeq data

[Fig. 4.1-4.3 Time-calibrated supertree of Sanger sequence and Angiosperms353 HybSeq low copy nuclear gene datasets. Clades 1 and 2 represent the first order subdivision of the family. Clades A to G represent lower order subdivisions corresponding to tribes.]

The supertree recovers major groupings (Clades 1 & 2, A through to G) congruent with the Angiosperms353 ASTRAL tree.

### Conflict between Sanger and HybSeq data

Combined Sanger and HybSeq data

Comparing topologies recovered by analyses of the Sanger *versus* Angiosperms353, sequence data (Figs. 2.1, 2.2 & 3) we identified three genera whose position was both strongly supported and in conflict between the respective datasets.

#### Sarchochlamys

In the Angiosperms353 sequence analyses *Sarchochlamys* is recovered within the Leukosykeae with strong support (Fig. 3, PP 1.0, QS 89.72). Within the Sanger data analyses, *Sarchochlamys* is recovered within the Boehmerieae (Fig. 2.1, PP 1.0, BS 100%).

#### Laportea canadensis

In the Angiosperms353 sequence analyses, *Laportea canadensis,* and sister *Laportea* species, is recovered within the Urticaeae and sister to *Girardinia* (Fig. 3: Laportea II, PP 0.83, QS 38.16). In the Sanger data analyses (Fig. 2.2), however, *Laportea canadensis* is recovered sister to a clade comprising *Laportea* accessions recognised as *Fleurya, Urera, Poikilospermum, Touchardia, Obetia* and *Scepocarpus* (BS 100%), which is itself recovered sister to *Girardinia*.

### Phylogenetic informativeness of morphological characters

Reconstructions of 22 morphological phylogenetically informative characters and 73 states useful for the delimitation of tribes and genera are shown in Fig. S3. Of these, all were recovered as homoplastic at the scale of family or tribe. In combination, however, character states were useful in delimiting both tribes and genera. For example, the combination of stem releasing an exudate (character state 2), amplexicaulous stipules (character state 8), and absence of a pistillode (character state 10) distinguished the putative tribe Cecropieae from all other tribes. The unique combinations of character states which delimit the proposed tribes and genera are listed below (Taxonomy section).

Our results, therefore, in combination with phylogenetic analyses, enable the recognition of seven major phylo-morphological groupings (A to G, Fig. 3, 4 & 5).

### Divergence time estimates and ancestral area reconstruction

Table 3 summarises the stem ages for all Urticaceae generic level groupings. Figure 3 presents divergence time estimates for the Angiosperms353 dataset and Figure 5 a time calibrated ASTRAL tree with reconstructed ancestral areas predictions.

**Table 3.**
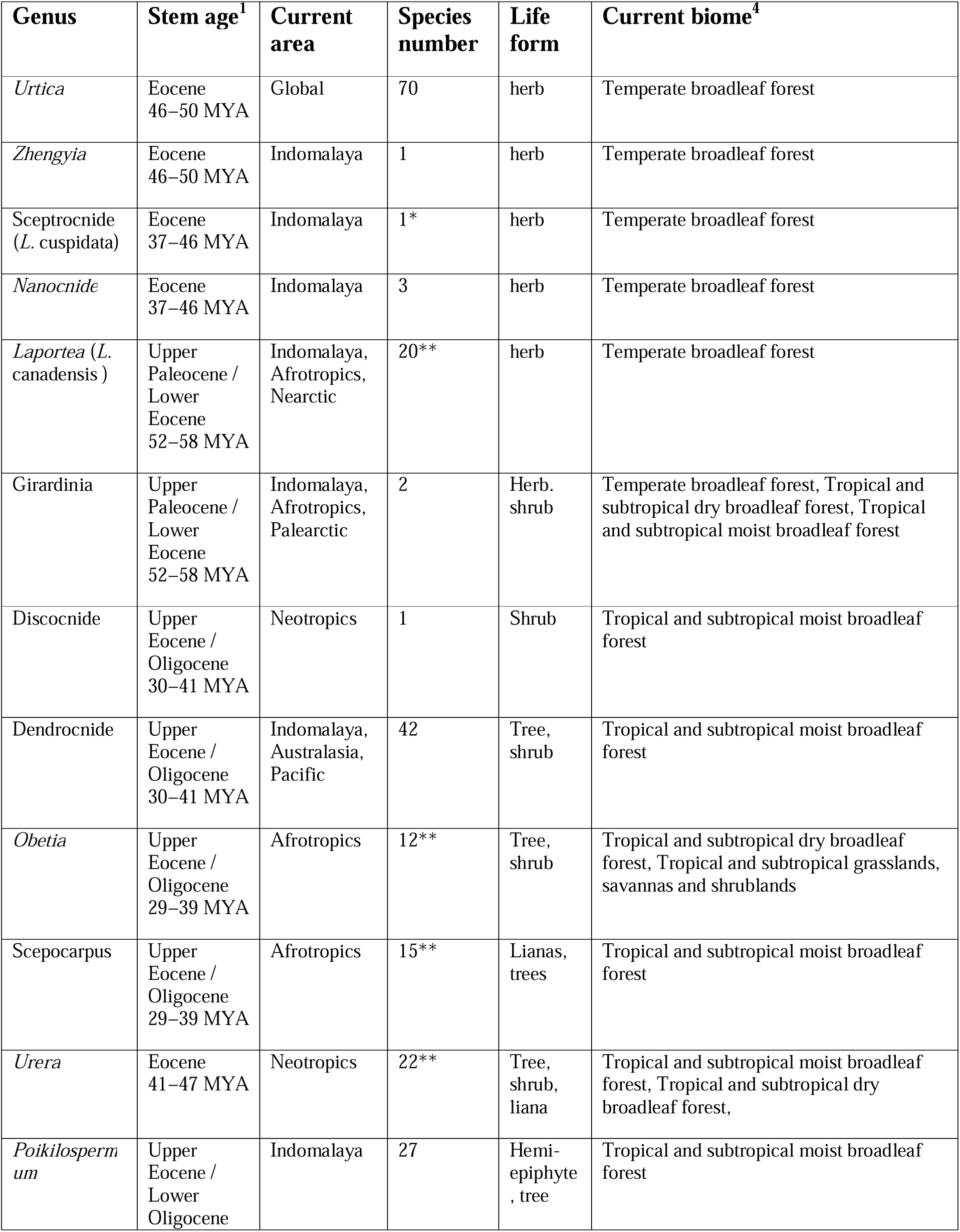
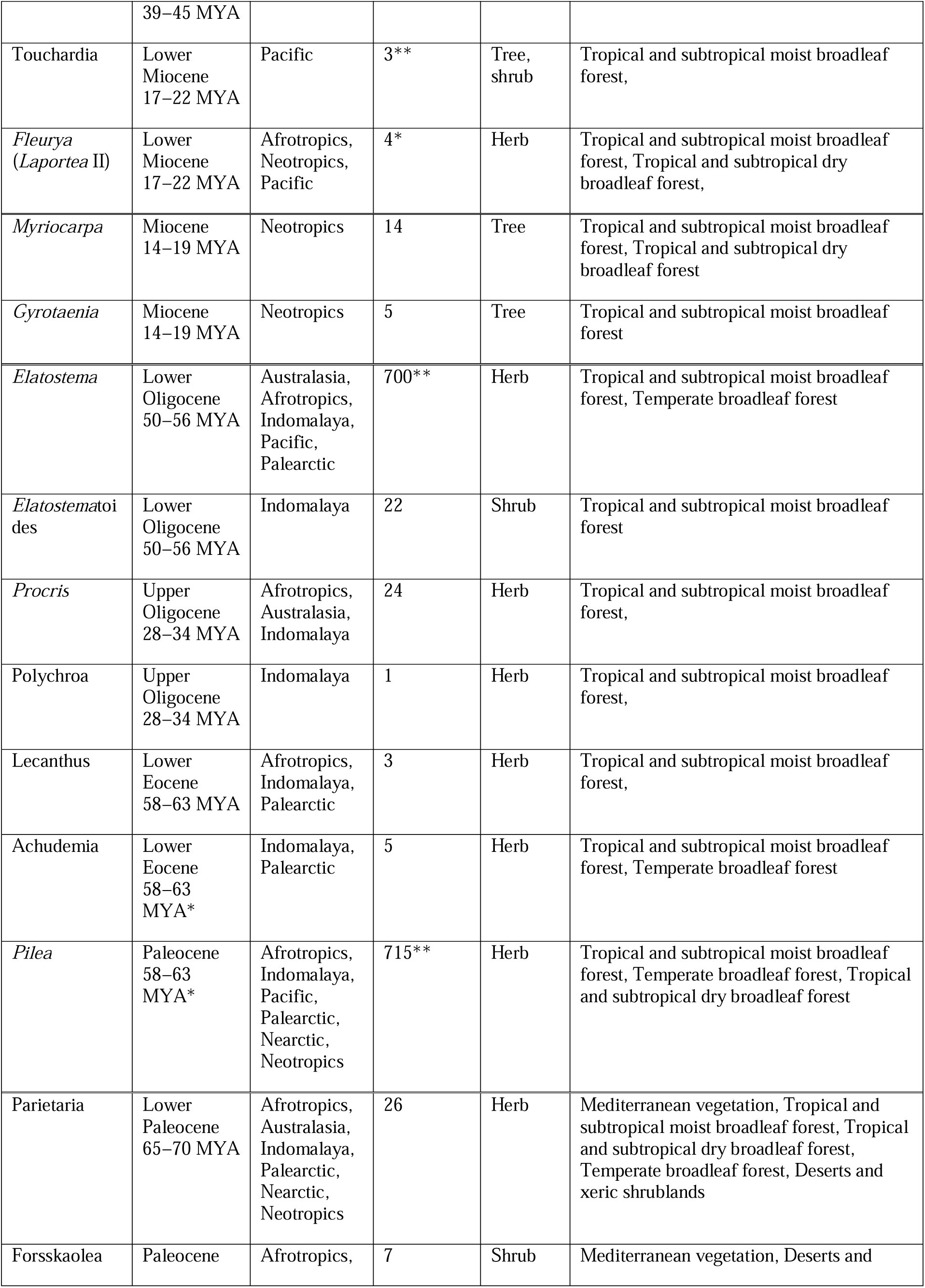
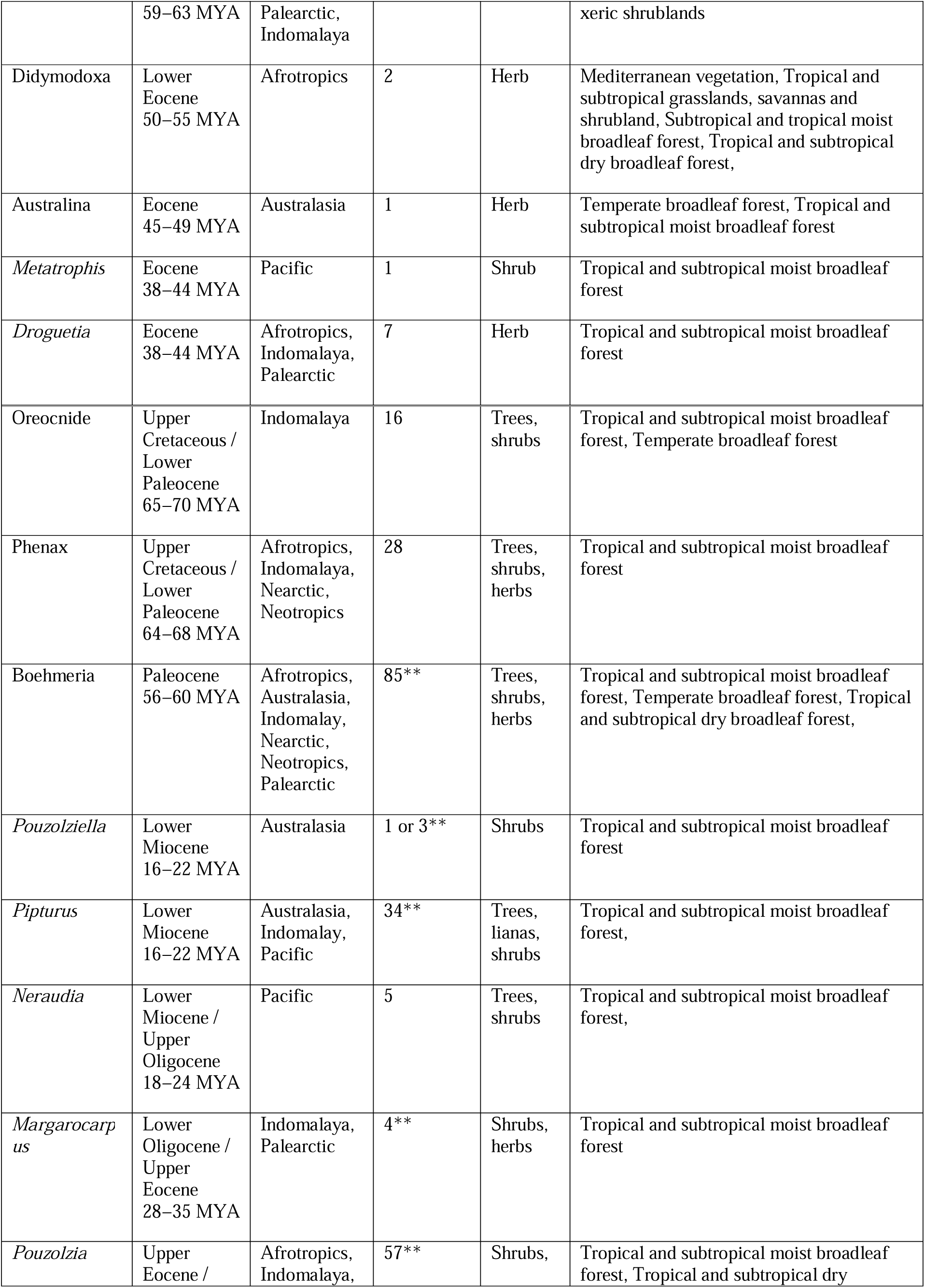
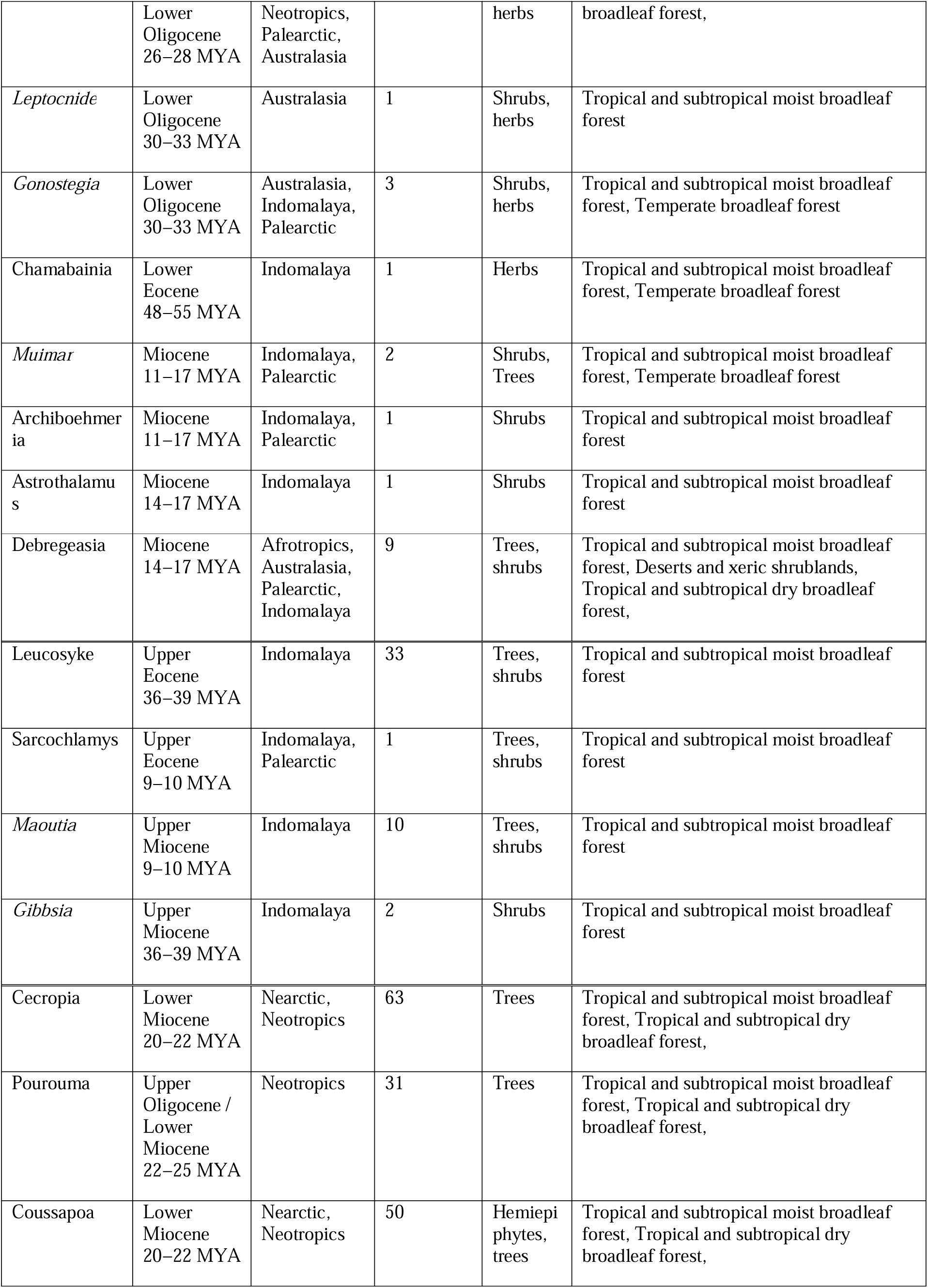
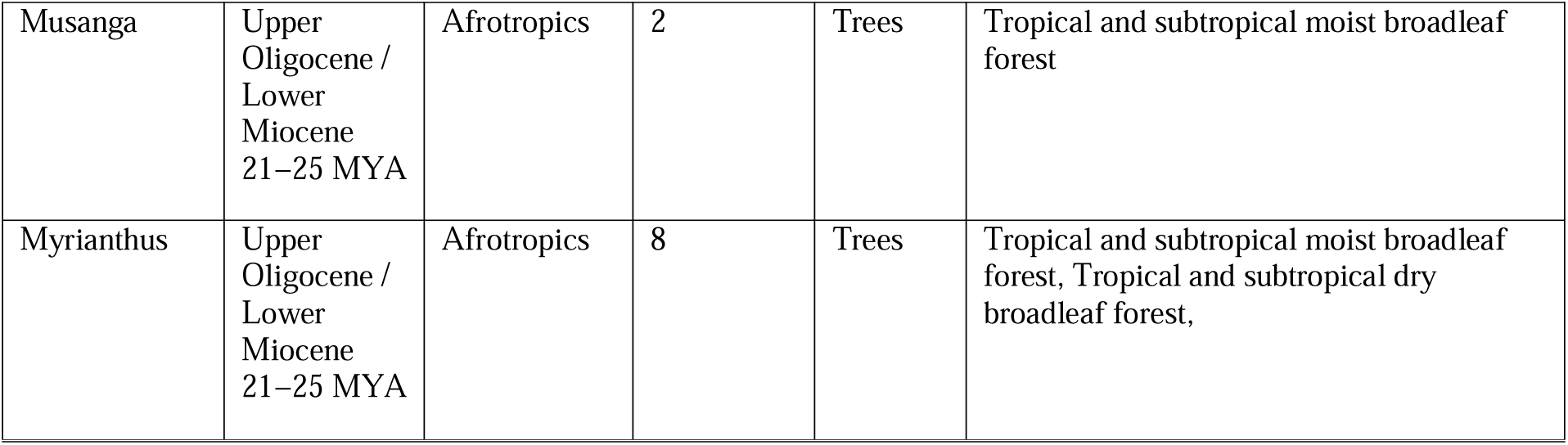
Estimated stem lineage ages, current distribution, species number, life form and biomes for Urticaeae genera.

We propose a stem age of 105−118 million years and a crown age of 93−93.3 million years for the Urticaceae. Of the seven major phylo-morphological groupings, four, Urticeae, Elatostemateae, Boehmerieae, Forrskaoleae and Parietarieae (Fig. 3 (clades A to E) are proposed to have originated at the end of the Cretaceous Period (Maastrichtian). Clade F (*Myriocarpa, Gyrotaenia*) is proposed to have originated in the Cretaceous (Fig. 3); Clade C (Cecropieae) in the Oligocene (clade C, Fig. 3); and clade D (*Gibbsia, Maoutia, Leukosyke* and *Sarcochlamys*) in the Eocene (D, Fig. 3).

Our results suggest that most genera originated in the Eocene, followed by the Miocene.

[Figure 5. Ancestral area estimates projected onto a time-calibrated supertree of Sanger sequence and Angiosperms353 HybSeq low copy nuclear gene datasets. Clades A to G represent lower order subdivisions corresponding to tribes.]

We recover a strong geographical signal within our analyses. Indomalaya is recovered as the most likely ancestral area for six of the seven major phylo-morphological groupings: that comprise *Gibbsia* + *Maoutia* + *Leukosyke* + *Sarcochlamys* clade; the Cecropieae, Boehmerieae, Urticeae, and Elatostemateae. The Palearctic was recovered as the most likely ancestral area for the Parietarieae. In contrast, the higher-order clades into which the tribes group lack strong geographical structure, several biogeographic regions being equally probable ancestral areas. Within tribes there is also major geographical structure. For example, within the Cecropieae, *Pourouma, Coussapoa* and *Cecropia* represent a monophyletic group of ancestrally neotropical taxa; within the Parietarieae, *Didymodoxa, Australina, Metatrophis* and *Droguetia* form a monophyletic group of Afrotropical taxa; within the Urticeae, *Obetia* (plus sister unplaced *Laportea* segregate) and *Scepocarpus* form a monophyletic group of Afrotropical taxa); and all of the extant taxa in the Myriocarpeae comprise neotropical taxa. Of the 53 taxa delimited as genera, 51 (96%) were recovered as having an ancestral area restricted to a single biogeographical region and most of these to Indomalaya. Exceptions were *Forsskaolea, Laportea* V (*Fleurya*), *Boehmeria, Elatostema, Fleurya, Pilea, Phenax, Poikilospermum, Pouzolzia* and *Urtica.* Of the exceptions, only *Fleurya* occurred in non-adjacent regions (Neotropics + Afrotropics, Pacific). Within species-rich genera that occurred across more than one biogeographic region (*Elatostema, Pilea, Urtica*) only *Urtica* was not recovered to have strong geographical phylostructure.

## Discussion

We were able to obtain a well resolved and strongly supported phylogeny for the Urticaceae that included all genera and ca 1/3 of the species. We found that the Angiosperms353 low copy nuclear data provided a well resolved and strongly supported backbone phylogeny of the Urticaceae, which when used as a constraint on a combined analysis with Sanger sequence data provided a robust framework for testing the monophyly of the genera, organising these into tribes, reconstructing morphological character ancestral states and allowing the identification of monophyletic morphological groups. This enabled us to recognise 53 groupings at the rank of genus and based on our evaluation of morphology and phylogenetic relationships we arrange these into seven tribes and provide a linear arrangement. The lack of resolution of previous studies may have resulted from a reliance on plastome sequence data (Gonçalves et al., 2019; Rokas et al., 2003), and or, less complete taxon-sampling.

### Experimental Error

#### Phylogenetic informativeness of morphological characters

Establishing hypotheses of homology for Urticaceae genera can be challenging due to the reduced nature of the flowers, on which much of past classifications were based, which can result in convergence in character states and or saturation of morphological space (Lee & Palci, 2021), leading to systematic errors in their coding and interpretation (Dávalos et al., 2014). Because we evaluated phylogenetic informativeness across the whole family (Fig. S3), morphological characters that would have been recovered as informative (plesiomorphic, apomorphic) within a subset of the Urticaceae may be classed as uninformative (homoplastic non-homologous) in the context of the whole family. The converse, however, would not be true, a character recovered as homoplastic in a subset of the tree could not be recovered as synapomorphic across broader tree. Our approach is therefore very conservative and character states that we have recovered as non-homologous at the family level should not be ignored by future studies focussing on subsets of the family.

### Tribal classification

Weddell’s tribal classification, based on Jussieu’s (Jussieu, 1789), Gaudicahud’s (Gaudichaud, 1830) and his own detailed and comprehensive reviews of the genera (Weddell, 1856, 1869), later updated by Friis (Friis, 1989), is largely congruent with recent family-level phylogenies of the Urticaceae. The exceptions are the Cecropieae (Fig.s 4 & 5, Clade C) and two clades, one comprising the Indomalayan genera,

*Gibbsia, Leukosyle, Maoutia* and *Sarchochlamys* (Fig.s 4 & 5, Clade D, Leukosykeae); the other, the neotropical genera, *Myriocarpa* and *Gyrotaenia* (Fig.s 4 & 5, Clade F, Myriocarpeae). Weddell’s classification of the family into five tribes is intuitive and using a small number of morphological traits it is easy to place taxa within it. For these reasons we have decided to adapt Weddell’s classification rather than replace it. To do so we propose two new tribes, the Leukosykeae, restricted to Indomalaya, and the Myriocarpeae, restricted to the neotropics.

Weddell (Weddell, 1856) had placed *Leukosyke* (as *Missiessya*), *Maoutia* and *Myriocarpa* (*Gyrotaenia* and *Gibbsia* had yet to be described) within the Boehmerieae subtribe Maoutieae based on the absence of a perianth in the female flowers (Weddell, 1856: 341). *Sarcochlamys* was placed in another subtribe of the Boehmerieae, together with several other genera, based on the female flowers having a perianth comprising free parts and a weakly succulent fruit. We recovered the Maoutieae, excluding *Myriocarpa* and with the addition of *Sarcochlamys,* as sister to the Cecropieae (Fig. 4, Clade C), and both as sister to a clade comprising Boehmerieae (Fig. 4, Clade A) + Parietarieae (Fig. 4, Clade B) + Forsskaoleae (Fig. 4, Clade B). Our evaluation of the phylogenetic informativeness of perianth absence suggests that the absence of perianth is homoplastic at this rank, occurring in *Australina, Didymodoxa, Droguetia, Forsskaolea, Maoutia* and *Phenax.* Our options were to place the genera in this clade into the Cecropieae or to propose a new tribe. We chose the latter, under the name Leukosykeae as these taxa are morphologically distinct from the Cecropieae, having non-amplexicaulous stipules, stipules not enclosing another clearly visible iteration of leaf and stipule, and possessing visible cystoliths in their leaf laminae.

*Myriocarpa* and *Gyrotaenia* were recovered as sister to the Elatostemateae (Fig.s 2.3, 4.2, 5, Clades E & F), and both as sister to the Urticeae (Fig.s 2.3, 3, 4.2 Clade G). *Myriocarpa* and *Gyrotaenia* are morphologically distinct from the *Elatostemateae* by their tree habit, the release, when cut, of a semi-opaque exudate from the stem that oxidises dark grey or black, conspicuous stipule scars, and the absence of subapical appendages in the male flowers. The two genera share the presence of fusiform cystoliths with the Elatostematatae, and conspicuous stipules and staminate flowers lacking subapical appendages with the Urticeae. We could have placed *Myriocarpa* and *Gyrotaenia* within an expanded Elatostemateae, or combined the Elatostemateae with the Urticeae tribes, or recognised them as a distinct grouping. We chose the latter option as either combination of taxa would have resulted in tribes that were very heterogenous and a challenge to delimit morphologically, and very difficult to diagnose in the field or herbarium.

The Forsskaoleae and Parietarieae were recovered as sister to each other (Fig.s 2.1, 3, 4.1, Clade B). Friis (Friis, 1989) expressed doubts as to their monophyly and indeed we recovered *Rousselia* and *Hemistylus* as embedded within the Boehmerieae. Schüßler *et al*. (Schüßler et al., 2019) showed clearly that *Parietaria* is polyphyletic with respect to *Gesnouinia* and *Soleirolia* and we have included them in *Parietaria* (see discussion on genus delimitation in the Taxonomy section below). The Parietarieae is therefore reduced to a single genus (*Parietaria*). Both Forsskaoleae and Parietarieae share complex and atypical flower arrangements for the Urticaceae. *Parietaria* includes species with truly bisexual flowers whilst the Forsskaoleae, are characterised by genera with flower-like inflorescences or flower clusters approximating to bisexual flowers (see Fig. S3: Trait 19), and associated with this, a transition to male flowers reduced to a single stamen (see Fig. S3 Trait 16). Both tribes are characterised by the presence of uncinate (hooked) hairs (see Fig. S3 Trait 4), one or no stipule (see Fig. S3 Trait 7), bisexual inflorescences (see Fig. S3 Trait 19), and an association with xeric environments. Given these shared characteristics we decided to combine the two tribes.

The Cecropieae is delimited as proposed, albeit at the rank of family, by Berg (Berg, 1978b), with the exception of *Poikilospermum,* which was recovered by us, and several previous studies as within the Urticeae (Huang et al., 2019; Kim et al., 2015c; Monro, 2006; Wells et al., 2021; Wu et al., 2013). The remaining tribes remain unchanged as summarised by Friis (Friis, 1989). For a morphological delimitation of the tribes see the ‘Key to the Tribes’ and circumscriptions of the tribes below (Taxonomy section).

### Phylogeography

Whilst not the main focus of this study we feel that it is worth noting the strong geographical signal across the Urticaceae (Fig. 5), which may contradict the proposed role of long-distance dispersal (LDD) (Wu et al., 2018) in the diversification of the Urticaceae. Patterns of relationships and stem ages of the genera and clades recovered by us suggest (Fig. 5) a boreotropical distribution with divergence along Laurasian dispersal routes, coupled with relatively slow dispersal, resulting in an Indomalayan centre of origin for the taxa ancestral to most of the tribes and many of the genera, with continent-restricted tropical distributions for the terminal genera, in Indomalaya, Africa or the neotropics. For example, the ancestral taxon to the Urticeae likely diverged in the Cretaceous in Indomalaya, but the present-day resulting genera now all have distributions restricted to a single biogeographic region, with the exception of the cosmopolitan genus *Urtica* (Fig.5, Table 3).

**Figure 5.**
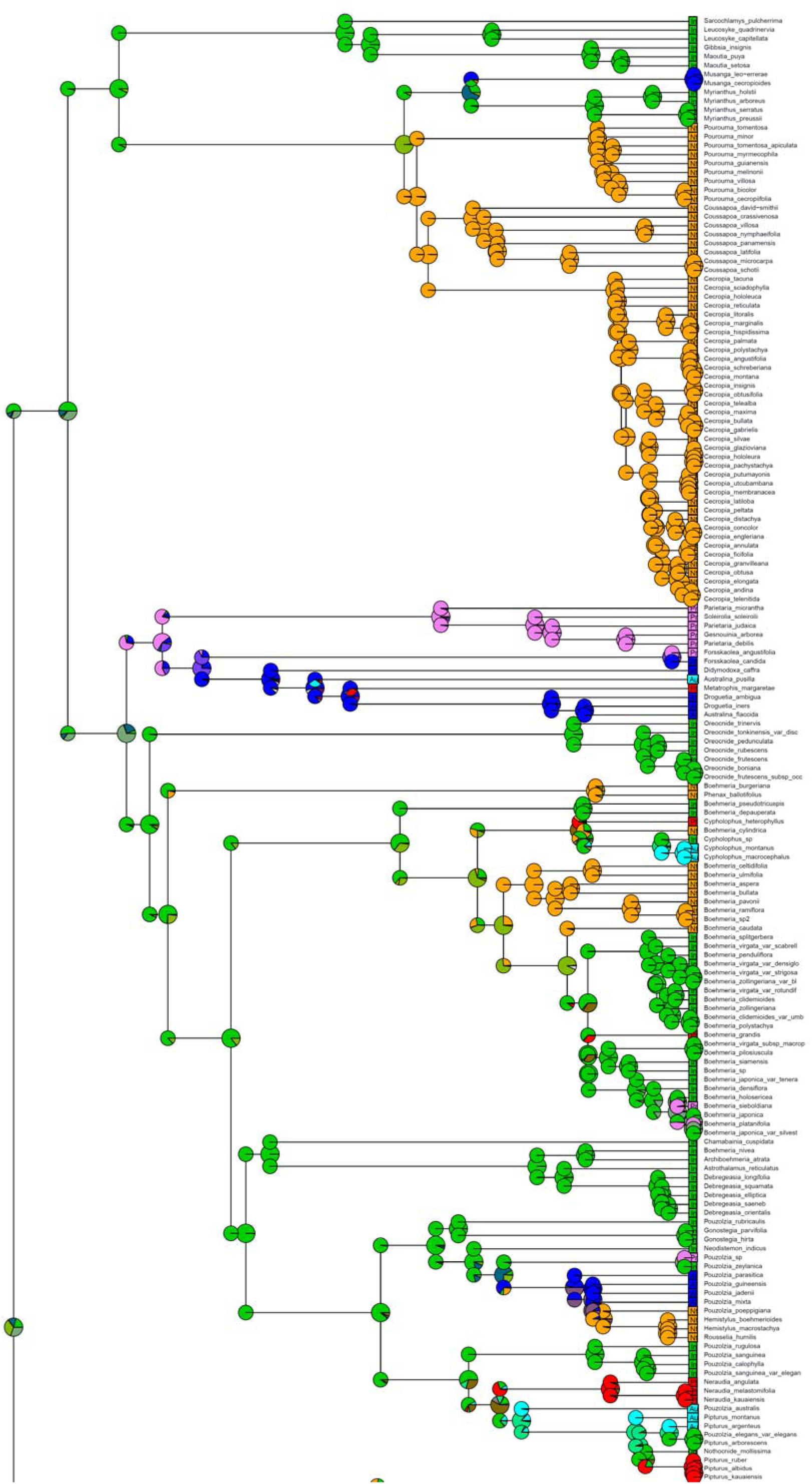
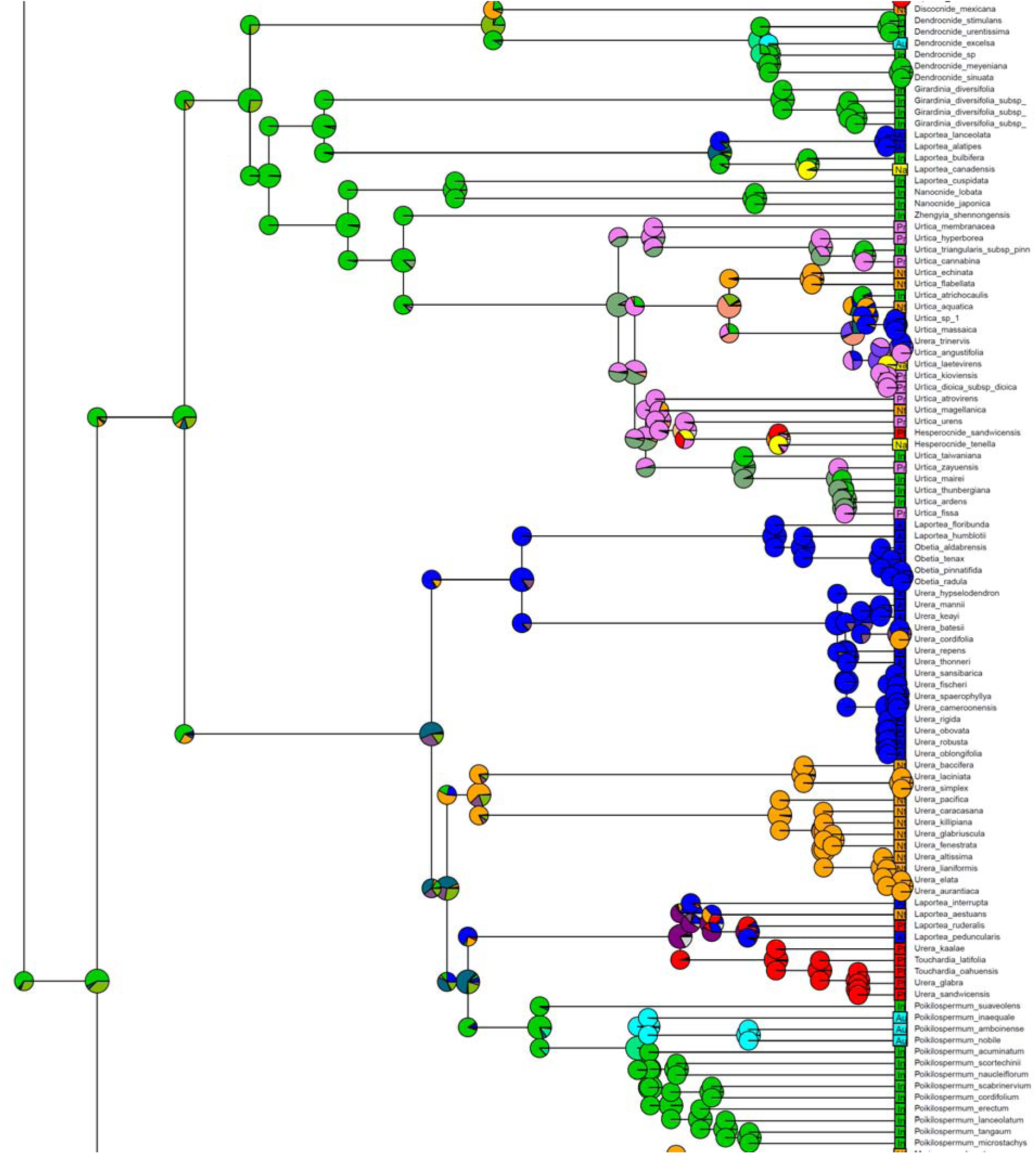
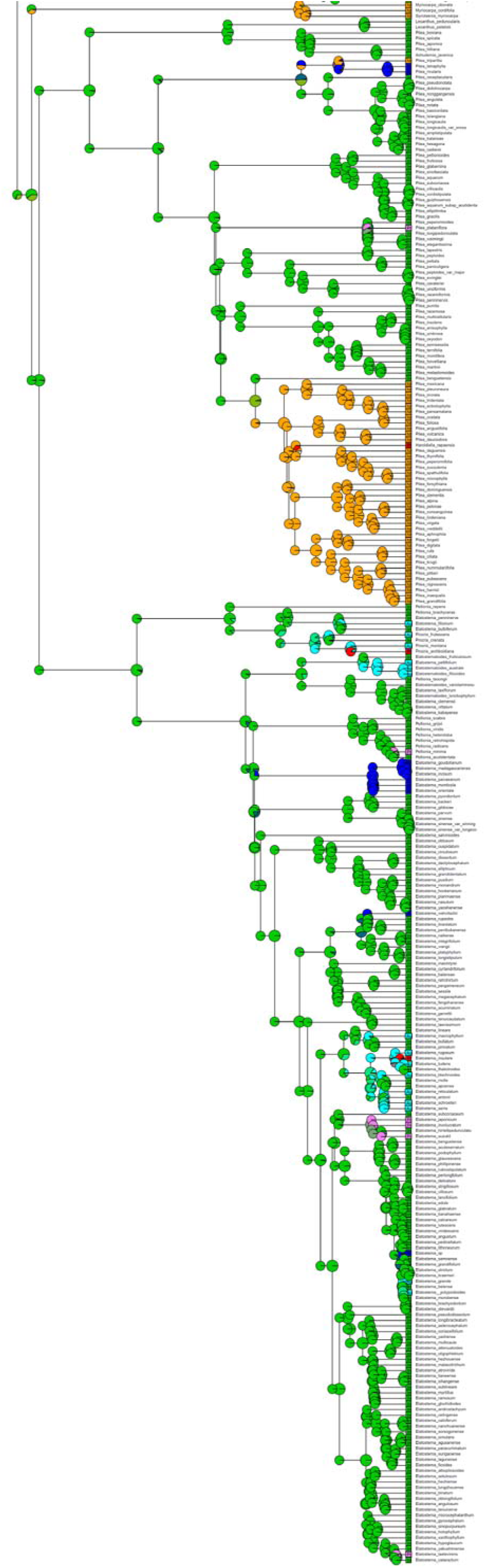
Ancestral area estimates projected onto a time-calibrated supertree of Sanger sequence and Angiosperms353 HybeSeq low copy nuclear gene datasets. Clades A to G represent lower order subdivisions corresponding to tribes.

Our results support a scenario whereby the Urticaceae originated in Indomalaya during the mid Cretaceous (Fig.s 4 & 5), becoming well represented in the Laurasian boreotropical flora (Dick & Pennington, 2019), 2019), dispersing from there to the neotropics and Africa during this time and subsequently becoming fragmented as the world cooled and tropical climates receded towards the equator during the Oligocene. The latter is supported by our recovery of the Afrotropics and neotropics as Upper Eocene / Oligocene / Miocene ancestral area for the stem node of several genera: *Cecropia, Coussapoa, Discocnide, Gyrotaenia, Musanga, Myrianthus, Myriocarpa, Obetia, Pourouma,* and *Scepocarpus*. Exceptions to this scenario are the proposed Paleocene, Lower Eocene or mid Eocene and Afrotropical origin for the Parietarieae and Forsskaoleae (Fig. 5).

### The delimitation of genera

In his ‘Théorie élémentaire de la botanique’, De Candolle (Candolle, 1813) wrote: “il résulte qu’un jour les limites des familles seront beaucoup mieux prononcées que celles des genres, et que ceux-ci ne pourront être établis définitivement qu’après les familles elles-mêmes; pour le moment il en est tout autrement, [translation: it follows that one day family limits will be much better articulated than those of genera, whose limits will only be definitively established after the families themselves; for the moment it is quite different, … ‘]. This is clearly the case with the Urticaceae at the present time, the integration of molecular data demonstrates many genera to be poly- and para- phyletic and in the process disrupting the morphological groupings developed by taxonomists over the last two centuries.

Adopting the criteria of monophyly and morphological diagnosability we recognize 53 genera within the Urticaceae. With the inclusion of the Cecropiaceae (five genera) and *Zhengyia* (Deng et al., 2013) into the family, this represents no increase on the total of 48 genera recognized by Friis (Friis, 1989. The delimitation of the genera, however, differs significantly with respect to *Boehmeria, Elatostema, Laportea, Neraudia, Parietaria, Pouzolzia,* with *Fleurya, Gonostegia*, *Leptocnide, Margarocarpus,* and *Sceptrocnide* being resurrected, and two new genera, *Muimar* and *Pouzolziella* being described (see Taxonomy section).

### A linear sequence for the genera

By aligning our linear sequence as much as possible with those used to organise herbarium collections ((Bentham & Hooker, 1883; Friis, 1989; Weddell, 1869),whilst reflecting phylogenetic relationships we seek to propose a sequence that will minimise the number of curatorial changes required should it be used for collections’ management. The major differences to existing sequences (Bentham & Hooker, 1883; Friis, 1989; Weddell, 1869) will be the establishment of the Myriocarpeae and their placement prior to the *Elatostemateae*; the placement of the Parietarieae between the *Elatostemateae* and the Boehmerieae. The establishment of the Leukosykeae does not affect the linear sequence as the genera that comprise it were placed at the end of the Boehmerieae, which they now follow. The re-delimitation and arrangement of many genera in the Boehmerieae, however, will lead to a number of changes: *Cypholophus* is incorporated into *Boehmeria, Muimar* is described (formerly *Boehmeria*) and placed adjacent to *Archiboehmeria, Leptocnide* and *Margarocarpus* (both formerly *Pouzolzia*) are resurrected and placed adjacent to *Gonostegia* and *Neraudia* respectively, *Pouzolziella* (formerly *Pouzolzia*) is described and placed prior to *Pipturus.* The sequence within the remaining tribes is, however, little affected: in the Urticeae, *Poikilospermum* and *Touchardia* are included and placed adjacent to *Fleurya* and *Laportea* sensu Chew is split into four resurrected genera. In the Parietarieae, *Metatrophis* is included and placed adjacent to *Droguetia*.

### Opportunities for future research

The aim of our research was to provide not only a stable classification but an evolutionary framework to facilitate further research in the family. We feel that the Urticaceae offers many exciting opportunities to ask broader taxonomic and evolutionary questions, and that in this respect the family has been overlooked. Of these we would like to highlight the following:

#### Transitions across biomes

Our dated estimations of ancestral states (Fig. S2) suggest that transitions from herbaceous to woodiness and non-fleshy to fleshy fruits occurred synchronously several times (Fig. S2, traits 2 & 21) in the diversification of the Urticaceae. Wells *et al*. (Wells et al., 2021), proposed that such a transition in the Urticeae may have been associated with major biome shifts and specifically the colonisation of closed-canopy forest. Closed canopies would have required lineages to increase in height to reach direct sunlight, either by becoming trees or lianas, or to adapt to the reduced light levels in the forest understory. Fleshy fruit would have enabled animal dispersal of the seeds. Within the Urticaceae both strategies can be observed. For example, the tree and liana genera such as *Dendrocnide* and *Scepocarpus*, or the deep shade specialists, *Elatostema* and *Pilea.* Fleshy fruit, however, are largely restricted to large shrubs and trees.

The shift to an arborescent habit and fleshy fruits occurred several times. Notably, during the upper Cretaceous / lower Palaeocene for the Cecropieae, arborescent Urticeae and *Oreconide,* the time at which closed canopy forests are proposed to have first developed (Carvalho et al., 2021). This was also the time at which there was a shift from mid / late successional vegetation to early successional vegetation by the Cecropieaea, and *Cecropia* is now one of the dominant ecological pioneer tree species in the Neotropics. It was also the epoch during which the Elatostemateae shifted to deep shade habitats, characterised by succulent herbaceous habit and distichous heteromorphic leaves. Taxa in these genera, notably *Pilea* and *Elatostema* are common components of the tropical montane forest groundstory, and in southeast Asia of the twilight zone in caves (Monro et al., 2018).

Within the Parietarieae there has been a shift to dry biomes, all of the genera included here being associated, or have species associated with deserts and xeric shrublands, the stem of this clade was also dated to the Cretaceous / lower Palaeocene.

We believe that the above warrant further study, incorporating greater taxon and trait sampling and that doing so would make a valuable contribution to macro-ecological studies that seek to address biome shifts and the origin of the forest biome.

#### Identifying the drivers of floral morphology

There are two aspects of floral morphology that merit greater study, the trend towards gynomonoecy in the *Parietarieae* (Fig. S2, trait 19), and the great variation in stigma morphology and associated degree of sepal fusion (Figure I).

The trend to bisexuality within the Parietarieae clade is noteworthy because it is restricted to this clade and has resulted in truly gynomonoecious flowers in *Parietari*.*a*We did not, however, recover bisexual inflorescences as ancestral in this clade (Fig. S2, trait 19) and so the most parsimonious explanation for it would be that it represents a shared response to a common driver. The observation that all taxa in the clade share a perennial herbaceous habit and an association with xeric, often rock environments may indicate an environmental rather than biological driver. Bertin & Kerwin (Bertin & Kerwin, 1998) in their study of gynomonoecy in *Aster* found no association with environmental or physiological variables and concluded that gynomonoecy could be a mechanism for reducing pollen-pistil interference or attracting pollinators. Gregg (Gregg, 1975), however, suggested that gynomonoecy could be a strategy for prioritising seed production in high light environments, whereby plants take advantage of abundant light and airflow to maximise seed production at the expense of pollen production.

We found some correlation between sepal fusion and pistil extension, taxa where the sepals are fused to form a tubular perianth having extended pistils, whilst those with divided free sepals generally lack extended pistils (FigS2, trait 12 & 17). The association of extended pistils and tubular flowers is well documented in animal pollinated flowers ((doi: 10.2307/annurev.ecolsys.34.011802.30000015, https://doi.org/10.3732/ajb.0900182) but less so in wind-pollinated species, especially where gynoecious. The broad variation in perianth and pistil morphology across the Urticaceae, combined with morphological arrangement characteristic of zoophily may suggest such a mechanism of pollination in ancestral lineages

#### Species documentation in Boehmeria and Cecropia

Our findings of strong phylogeographic structure within Urticaceae suggest that *Boehmeria,* which includes many species with broad geographical ranges and morphological variation (e.g. *B. virgata* occurs in 32 countries and has 14 varieties, (Wilmot-Dear & Friis, 2013)) likely harbours many more species, many of which could be considered cryptic, than currently recognised. For this reason, a DNA-first approach involving much greater taxon sampling followed by an evaluation of the phylogenetic informativeness of morphological characters is a priority.

The genus *Cecropia* comprises ca 63 species of pioneer tree species. Species delimitation, however, is unstable (Berg et al., 2005) and based on relatively few collections, often of a single sex. A lack of collections is due to the fact that the species are dioecious, but it is not obvious, combined with their occupation by aggressive stinging ants that can undermine the enthusiasm of the most stoic collector. *Cecropia* are, however, important components of neotropical tropical forest, both because of their role as pioneer species, but also as important sources of food for many large frugivorous birds and mammals (Stevenson et al., 2015). A stable and comprehensive classification will thus have a broad impact on the science of ecology.

#### Intrinsic drivers of speciation and substrate as drivers of speciation in the Elatostemateae

The two genera where species radiations have unequivocally occurred are *Pilea* (ca 715 spp versus sister genus Lecanthus, 5spp), and *Elatostema* (ca 700 spp, versus sister genus, *Elatostema* toides, ca 20 spp). Both are characterised by high numbers of point-endemics, especially on limestone and ultramafic substrates. For example, in the case of *Elatostema,* 184 out of 280 species native to China are associated with and mostly restricted to limestone substrates (Wang, 2014). We have found evidence that the colonization of limestone has occurred very rarely within *Elatostema* and that it is associated with species radiation, apomixis and increased rates of diversification (Fu et al. *in prep.*). Both *Pilea* and *Elatostema* occupy shaded microhabitats in montane forest and have wind-pollinated flowers and poorly mechanically dispersed seeds. As such they seem ill-suited for dispersal and vulnerable to allopatry. A study by (Fu et al., 2017) based on a small sample of limestone associated species suggested that 1/3 of the species reproduce by apomixis and this is also suggested to be the case in *Pilea* (L. Fu et al., 2022). Apomixis can provide a mechanism whereby founding individuals can colonise new habitats, reproduce, disperse and benefit from sexual isolation (Hörandl, 2010) enabling well-adapted lineages to persist..

## Conclusion

We use a combination of Sanger sequence data from the plastome, 353 low copy nuclear genes and morphology to revise the classification of the Urticaceae and delimitation of many of the genera. We aimed to do so by recognising groupings where monophyly and morphological discontinuities coincide. This resulted in the recognition of seven tribes, two of which we describe here as new, 53 genera, two of which we describe here as new, and a linear sequence for the family. Our analyses suggest strong congruence between geography and phylogenetic signal. We also identify several foci for future research and suggest that the family could be of value in researching several broader questions in evolutionary biology.

## Taxonomy

Tribal arrangement and linear sequence of the *Urtica* ceae

**Urticeae** Lam. & DC. (1806: 184). Fig. 6.

**Fig. 6.**
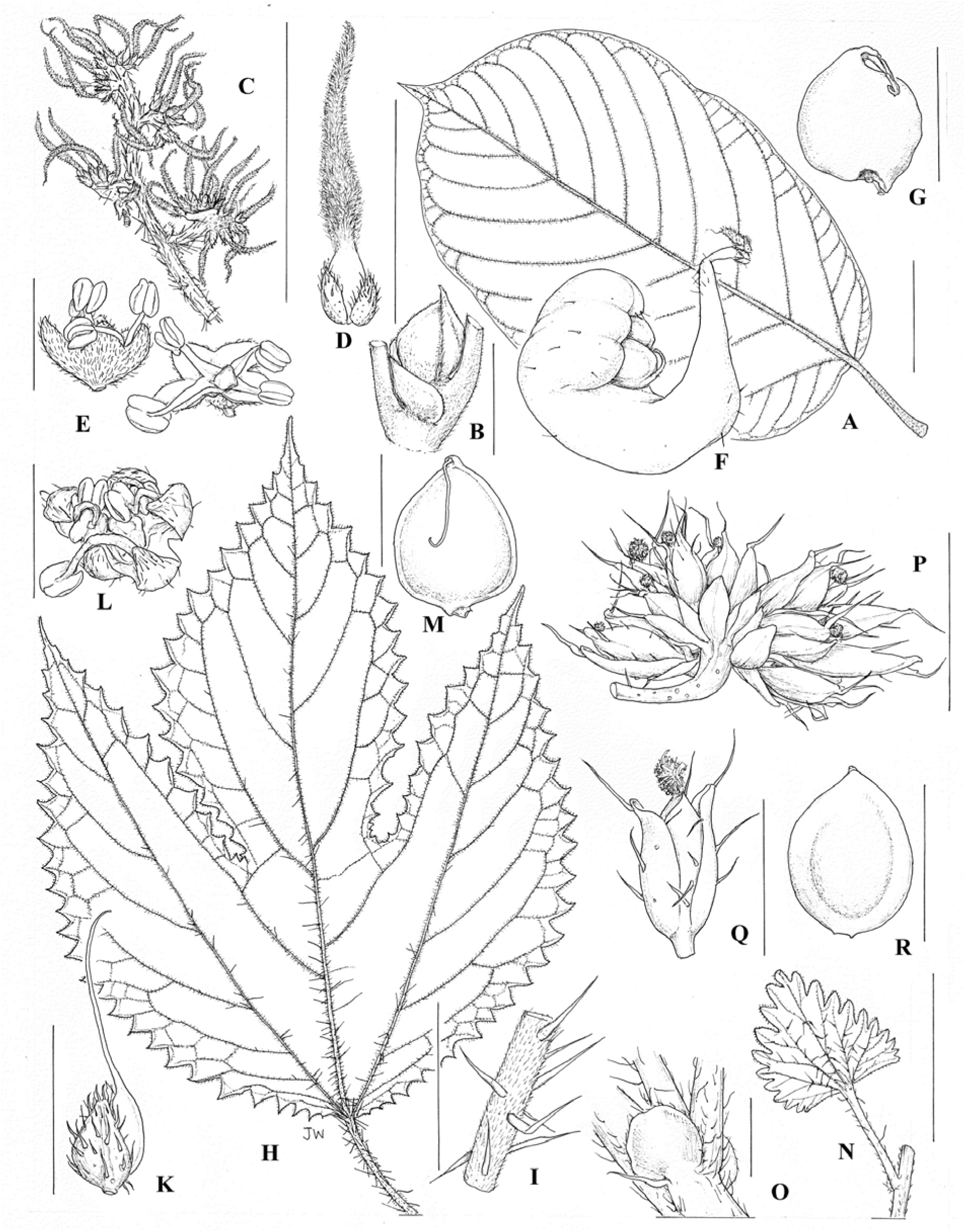
Urticeae Tribe. *Dendrocnide meyerianum*: **A** leaf lower surface based on xxx 5008 (scale bar = 3 cm); **B** stipule based on xxx 26044 (scale bar = 1 cm); **C** female flower cluster based on xxx 5008 (scale bar = 1 cm); **D** female flower based on xxx 5008 (scale bar = 2 mm); **E** male flowers based on xxx 5008 (scale bar = 2 mm); **F** fruit based on image xxx (scale bar = 2 mm); **G** achene based on xxx 6892 (scale bar = 2 mm). *Girardinia diversifolia*: **H** leaf lower surface based on Biye 124 (scale bar = 3 cm); **I** section of stems with bulbed hairs based on Reekmans 6129 (scale bar = 1 cm); **J** female flower based on Biye 124 (scale bar = 1 mm); **K** male flower based on Reekmans 6129 (scale bar = 2 mm); **L** achene based on Biye 124 (scale bar = 2 mm). *Nanocnide japonica*: **M** leaf lower surface based on Yao 8878 (scale bar = 3 cm); **N** female inflorescence based on Togashi 7840 (scale bar = 2 mm); **O** female flower based Togashi 7840 (scale bar = 1 mm); **P** stipule based on Yao 8878 (scale bar = 2 mm); **Q** achene based on Furusa 5632 (scale bar = 1 mm). Illustration by Juliet Beentje.

*Urtica*

*Zhengyia*

Sceptrocnide (*Laportea* cuspidata clade)

*Nanocnide*

*Laportea* (*Laportea* canadensis clade)

*Girardinia*

*Discocnide*

*Dendrocnide*

*Obetia*

*Scepocarpus*

*Urera*

*Poikilospermum*

*Touchardia*

*Fleurya* (*Laportea* II)

**Myriocarpeae** tribe nov., A.K.Monro, I. Friis & Wilmot_Dear. Type: ***Myriocarpa*** Benth., Bot. Voy. Sulphur 168. t. 55. 1844. Fig. 7.

**Fig. 7.**
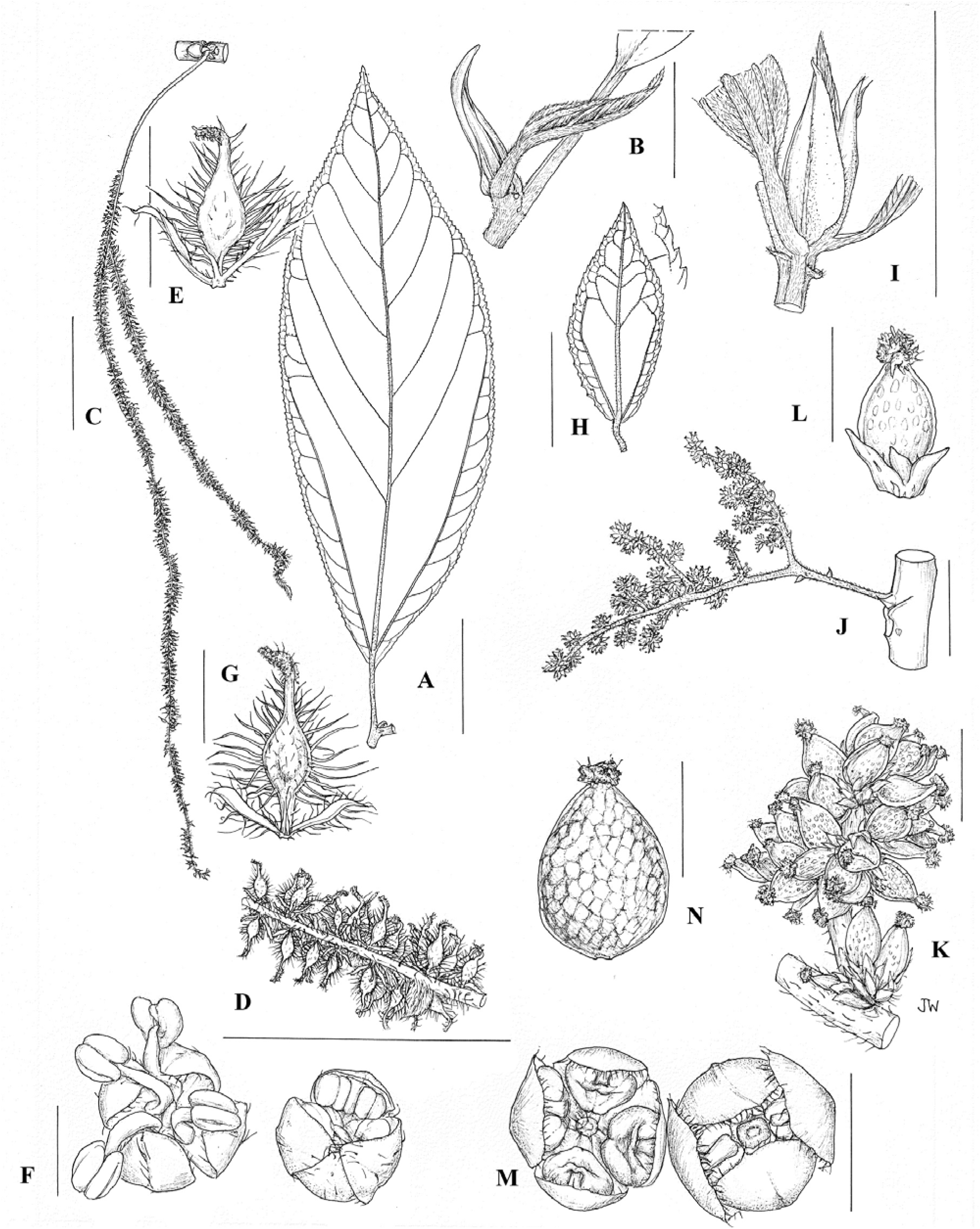
Myriocarpeae Tribe. Myriocarpa 79tipitate:**A** leaf lower surface based on *Huallparimachi* 3052 (scale bar = 3 cm); **B** stipule () based on *Huallparimachi* 3052 (scale bar = 1 cm); **C** female inflorescence based on *Huallparimachi* 3052 (scale bar = 3 cm); **D** female flower cluster based on *Huallparimachi* 3052 (scale bar = 1 cm); **E** female flower based on based on *Huallparimachi* 3052 (scale bar = 1 mm); **F** male flowers in bud, and open, based on *Ule* 6507 (scale bar = 1 mm); **G** fruit and achene based on *Huallparimachi* 3052 (scale bar = 1 mm). *Gyrotaenia macrocarpa*: **H** leaf lower surface based on *Monro et al.* 7479 (scale bar = 3 cm); **I** stipules based on *Breedlove* 68859 (scale bar = 1 cm); **J** female inflorescence based on *Hinton* 757 (scale bar = 1 cm); **K** female flower cluster based on *Hinton* 757 (scale bar = 1 mm); **L** female flower based on *Hinton* 757 (scale bar = 0.5 mm); **M** male flowers based on *Monro et al.* 7479 (scale bar = 1 mm); **N** achene based on *Breedlove* 68859 (scale bar = 1 mm). Illustration by Juliet Beentje.

*Myriocarpa*

*Gyrotaenia*

**Elatostemateae** Gaudich. (1830: 493). Fig. 8

**Fig. 8.**
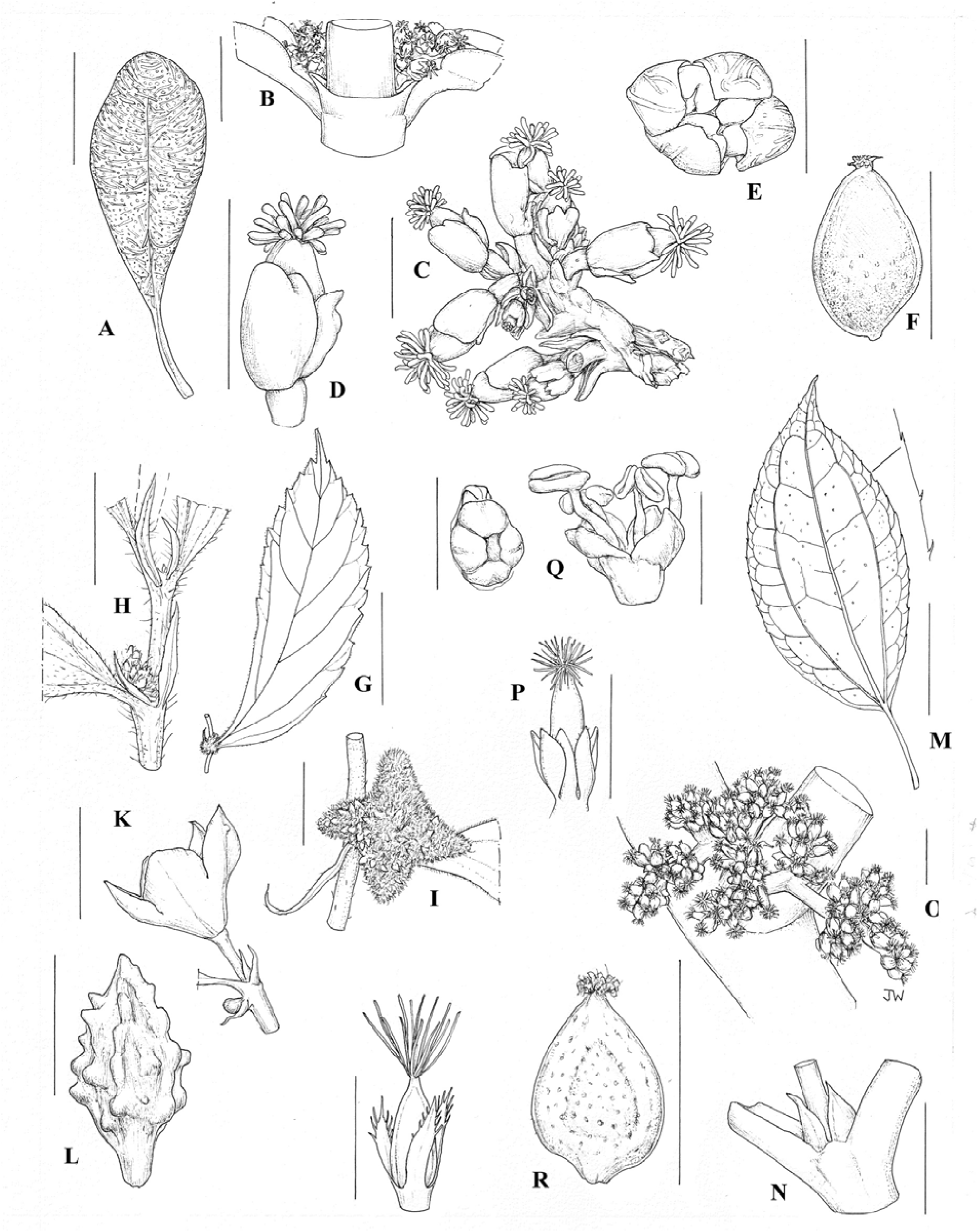
Elatostemateae Tribe. *Pilea microphylla*: **A** upper leaf surface based on *Ekman* 695 (scale bar = 2 mm); **B** stipule based on photograph by P. York (BM) (scale bar = 0.5 mm); **C** female inflorescences based on *Ekman* 695 (scale bar = 0.5 mm); **D** female flower based on *Ekman* 695 (scale bar = 0.5 mm); **E** male flower based on based on *Ekman* 695 (scale bar = 0.5 mm); **F** achene based on *Britton & Shafer* 300 (scale bar = 0.5 mm). *Elatostema XX*: **G** leaf outline based on *Du* HNK2960 (scale bar = 3 cm); **H** stipules based on *Du* HNK2960 (scale bar = 5 mm); **I** female inflorescence based on *Du* HNK2960 (scale bar = 5 mm);**J** female flower based on Weddell (1856: plate X, 10 (XX); **K** male flower based on *Parnell* 95-661 (scale bar = 2 mm); **L** achene based on *Du* HNK2960 (). *Achudemia boniana*: **M** leaf outline based on *Henry* 9771 (scale bar = 3 cm); **N** stipules based on *Henry* 9771 (scale bar = 2 mm); **O** female inflorescence based on *Monro* 6704 (scale bar = 5 mm); **P** female flower based on *Monro* 6704 (); **Q** male flower and bud based on *Monro* 6704 and *Morse* 495 (scale bar = 2 mm); **R** achene based on *Henry* 9771 (scale bar = 2 mm). Illustration by Juliet Beentje.

*Elatostema*

*Elatostematoides*

*Procris*

*Polychroa (Pellionia repens)*

*Lecanthus*

*Achudemia*

*Pilea*

**Parietarieae** Gaudich. (1830: 501). Fig. 9.

**Fig. 9.**
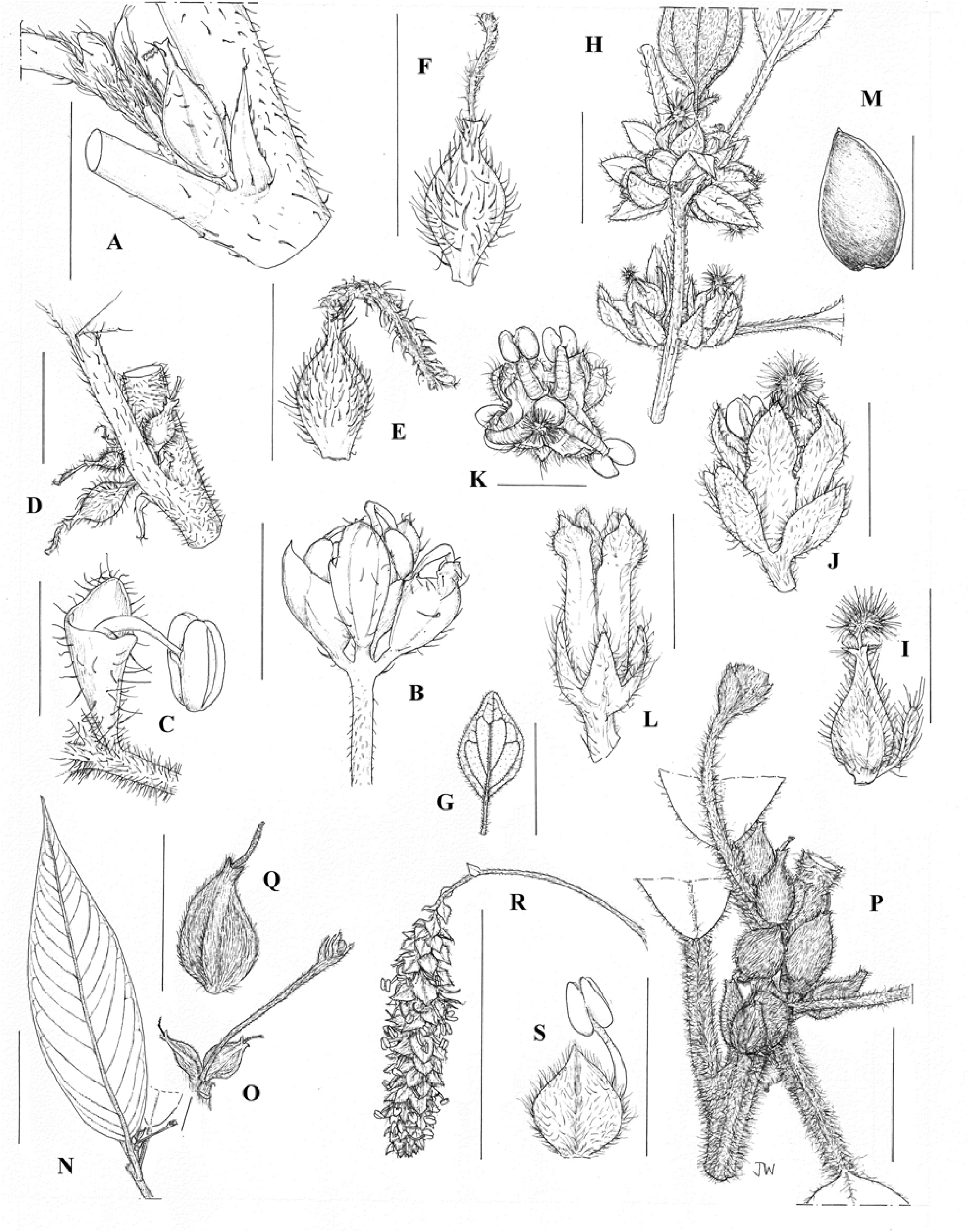
ParietarieaeTribe. *Australina pusilla*: **A**. stipule based on *Constable* 5468 (scale bar = 2 mm); **B** male inflorescence based on *Constable* 5468 (scale bar = 4 mm); **C** male flower based on *Adams* 766 (scale bar = 2 mm); **D** female inflorescence showing bracts based on *Adams* 766 (scale bar = 2 mm); **E** female flower based on *Adams* 766 (scale bar = 1 mm); **F** achene based on *Adams* 766 (scale bar = 2 mm). *Parietaria Judaica*: **G** leaf outline and lower surface based on *Carter* 893 (scale bar = 3 cm); **H** flower clusters comprising female and bisexual flowers based on *Carter* 893 (scale bar = 4 mm); **I** female flower based on *Carter* 893 (scale bar = 2 mm); **J** bisexual flowers (side-view) based on *Carter* 893 (scale bar = 2 mm); **K** bisexual flowers (top-view) based on *Carter* 893 (scale bar = 2 mm); **L** bisexual fruiting perianth bisexual flowers (side-view) based on *Carter* 893 (scale bar = 2 mm); **M** achene bisexual flowers (side-view) based on *Carter* 893 (scale bar = 1 mm). *Metatrophis margaretae*: **N** leaf with inflorescence in the axil based on *Fosberg* 11422 (scale bar = 3 cm); **O** inflorescence detail (female flowers towards the base, male flowers towards the apex) based on *Stokes* 123 (scale bar = XX); **P** cluster of female flowers and fruits based on *Stokes* 123 (scale bar = 4 mm); **Q** female flower based on *Fosberg* 11422 (scale bar = 4 mm); **R** male inflorescence based on *Fosberg* 11422 (scale bar = 3 cm); **S** male flower based on *Fosberg* 11422 (scale bar = 4 mm). Illustration by Juliet Beentje.\

*Parietaria*

*Forsskaolea*

*Didymodoxa*

*Australina*

*Metatrophis*

*Droguetia*

**Boehmerieae** Gaudich. (1830: 499). Fig. 10.

**Fig. 10.**
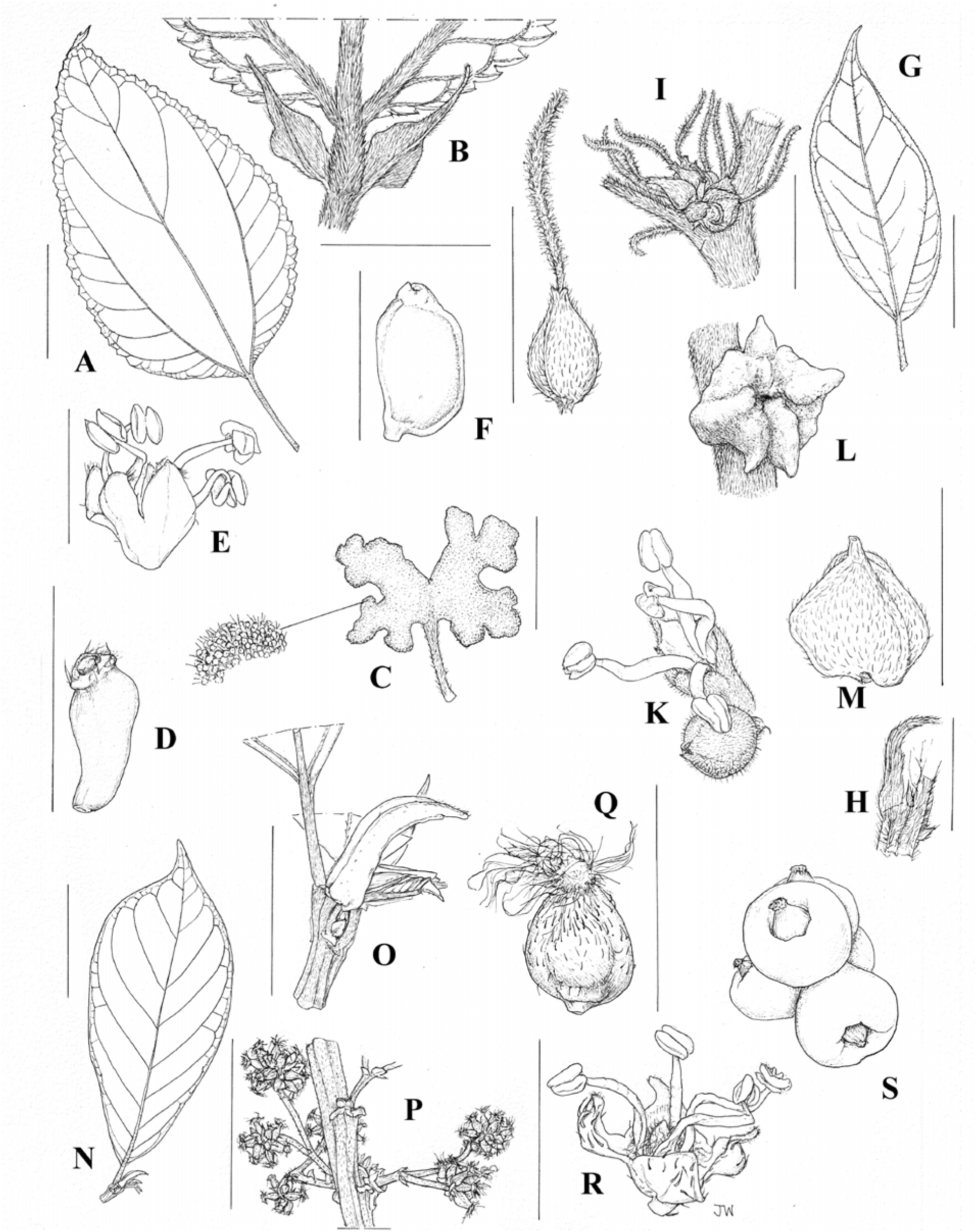
Boehmerieae Tribe. *Astrothalamus reticulatus*: **A.** leaf outline based on *Argent* 981987 (scale bar = 2 mm); **B** stipules based on *Argent* 981987 (scale bar = 4 mm); **C** female inflorescence based on SAN 78665 (scale bar = 2 mm); **D** female flowers based on SAN 78665 (scale bar = 2 mm); **E** male flower based on *Argent* 981987 (scale bar = 1 mm); **F** fruit based on *Argent* 981987 (scale bar = 2 mm). *Neraudia melastomifolia*: **G** leaf outline based on *Cowan* 836 (scale bar = 3 cm); **H** stipules based on *N. angulata* and *Greenwell* 20832 (scale bar = 4 mm); **I** female inflorescence based on *Cowan* 836 (scale bar = 2 mm); **J** female flower based on *Cowan* 836 (scale bar = 2 mm); **K** male flower based on field image (scale bar = 2 mm); **L** fruit based on field image of *N. angulata* (scale bar = 2 mm); **M** fruit based on *Cowan* 836 (scale bar = 2 mm)*. Oreocnide xxx.* **N** leaf outline based on *Gardette* 957 (scale bar = 3 cm); **O** stipules based on *Gardette* 957 (); **P** female inflorescence based on *Gardette* 957 (scale bar = 4 mm); **Q** female flower based on *Gardette* 957 (scale bar = 4 mm); **R** male flower based on *Lörzing* 13751 (scale bar = 3 cm); **S** fruits based on field image (scale bar = 4 mm). Illustration by Juliet Beentje.

*Oreocnide*

*Phenax*

*Boehmeria*

*Pouzolziella* gen. nov.

*Pipturus*

*Neraudia*

*Margarocarpus*

*Pouzolzia*

*Leptocnide*

*Gonostegia*

*Chamabainia*

*Muimar gen. nov*.

*Archiboehmeria*

*Astrothalamus*

*Debregeasia*

**Leukosykeae** tribe nov., A.K.Monro, I. Friis & Wilmot_Dear. Type: Leukosyke Zoll. & Moritzi, Syst. Verz.: 100. 1845. Fig. 11.

**Fig. 11.**
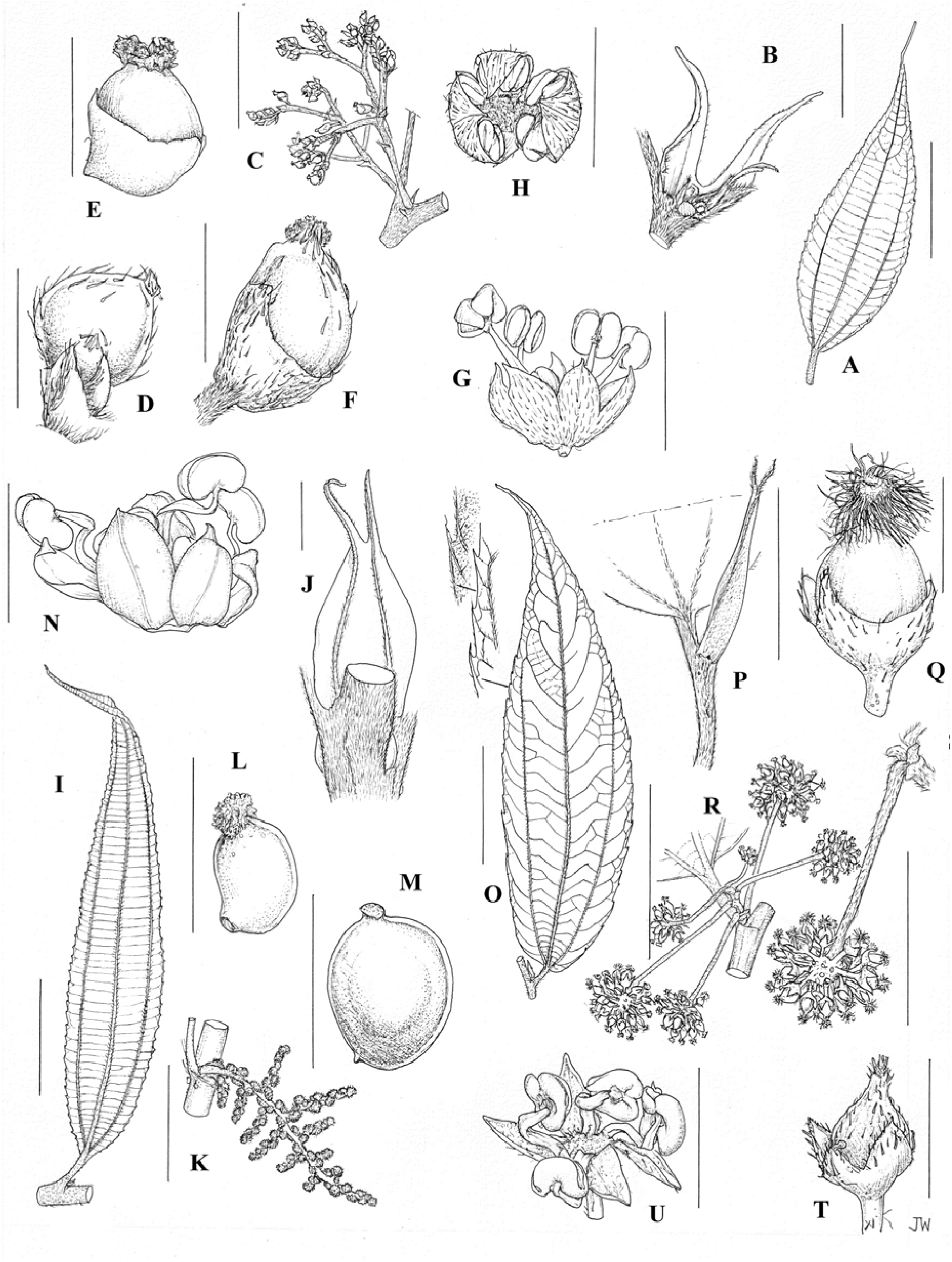
LeukosykeaeTribe. *Gibbsia insignis.* **A** leaf lower surface based on *Brass* 12279 (scale bar = 3 cm); **B** stipule based on *Lae* 59516 (scale bar = 2 mm); **C** female infructescence based on *Brass* 12279 (scale bar = 1 cm); **D** female flower lateral view based on *Brass* 12279 (scale bar = 1 mm); **E** female flower based on *Lae* 59516 (scale bar = 1 mm); **F** fruit and achene based on *Brass* 12279 (scale bar = 1 mm); **G** male flower lateral view based on *Brass* 11075 (scale bar = 2 mm); **H** male flower apical view based on *Brass* 12279 (scale bar = 2 mm). *Sarcochlamys pulcherrima*: **I** leaf lower surface based on *Hossain* 1911 (scale bar = 3 cm); **J** stipule based on *Grierson* 1427 (scale bar = 2 mm); **K** female inflorescence based on *Grierson* 1525a (scale bar = 3 cm); **L** female flower based on *Hossain* 1911 (scale bar = 0.5 mm); **M** achene based on *Grierson* 1525a (scale bar = 0.5 mm); **N** male flower lateral view based on *Grierson* 1427 (scale bar = 1 mm). *Leucosyke corymbulosa*: **O** leaf lower surface based on *Greenwood* 43 (scale bar = 3 cm); **P** stipule based on *Parham* 47 (scale bar = 1 cm); **Q** female flower based on *Smith* 184 (scale bar = 1 mm); **R** infructescence based on *Smith* 184 (scale bar = 3 cm); **S** infructescence detail based on *Smith* 184 (scale bar = 1 cm); **T** achene based on *Greenwood* 43 (scale bar = 1 mm); **U** male flower based on *Meebold* 16699 (scale bar = 2 mm). Illustration by Juliet Beentje.

*Leukosyke*

*Sarcochlamys*

*Maoutia*

*Gibbsia*

The tribal name is derived from “syce”, Greek for “fig tree”. If treated as Latin and following the third declension, it should be declined as a consonant in singular nominative and in singular genitive -is: Leukosyke (genitive singular => Leukosykis; tribal name ==> Leukosykeae).

**Cecropieae** Gaudich. (1830: 506). Fig. 12.

**Fig. 12.**
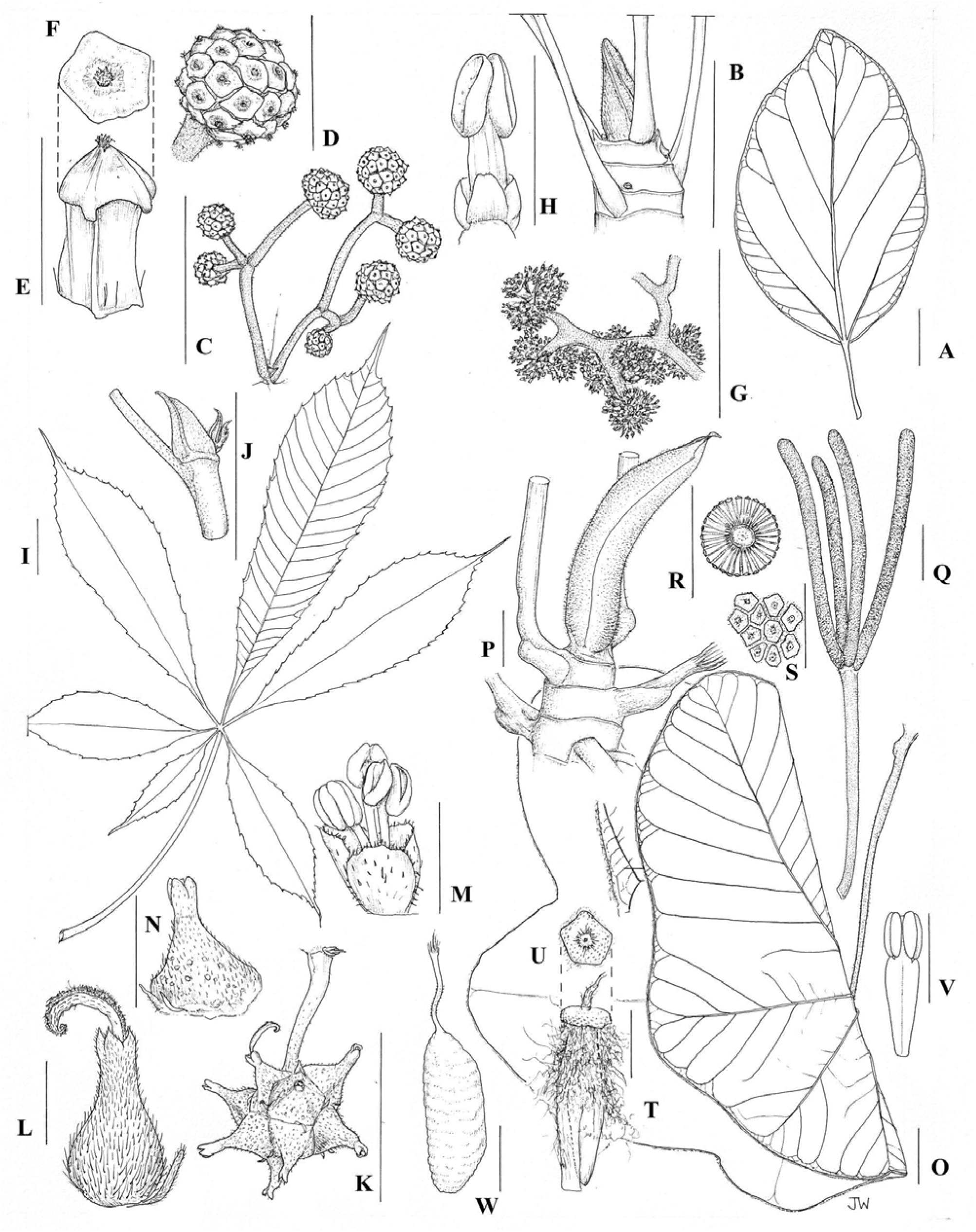
CecropieaeTribe. *Coussapoa latifolia*: **A** leaf based on *Lindeman* 249(?) (scale bar = 3 cm); **B** stipule based on *Lindeman* 249(?) (scale bar = 3 cm); **C** infructescence based on *Lindeman* 249(?) (scale bar = 1 cm); **D** female flower cluster based on *Lindeman* 249(?) (scale bar = 3 mm); **E** female flower lateral based on *Lindeman* 249(?) (scale bar = 1 mm); **F** female flower apical view based on *Lindeman* 249(?) (); **G** small section of male inflorescence based on *Cid* 8852 (scale bar = 1 cm); **H** male flower based on *Cid* 8852 (scale bar = 0.5 mm). *Myrianthus* xx: **I** leaf based on *Osborne* 326 (scale bar = 3 cm); **J** stipule based on *Etuge* 5161 (scale bar = 3 cm); **K** fruiting inflorescence based on *Leuuwenberg* 5190 (scale bar = 3 cm); **L** female flower based on *Osborne* 326 (scale bar = 3 mm); **M** male flower based on *Etuge* 5161 (scale bar = 1 mm); **N** single unit of fruit based on *Leuuwenberg* 5190 (scale bar = 1 cm). *Cecropia* XX: **O** leaf outline based on *Benson* 10345 (scale bar = 3 cm); **P** stipule based on *Benson* 10345 (scale bar = 3 cm); **Q** infructescence based on *Nee* 42895 (scale bar = 3 cm); **R** cross-section of female inflorescence based on *Nee* 42895 (scale bar = 1 cm); **S** surface detail of female inlorescence based on *Nee* 42895 (scale bar = 3 mm); **T** female flower lateral view based on *Nee* 42895 (scale bar = 1 mm); **U** female flower apical view based on *Nee* 42895 (scale bar = 1 mm); **V** male flower based on Flora Neotropica: (scale bar = 1 mm); **W** fruit based on Flora Neotropica: (scale bar = 1 mm). Illustration by Juliet Beentje.

*Cecropia*

*Pourouma*

*Coussapoa*

Musanga

Myrianthus

### Key to the tribes

1. Stipules amplexicaulous and conical, enclosing a well-formed leaf which itself encloses another well-formed stipule; cystoliths absent from mesophyll; secondary veins of leaf laminae parallel, conspicuously closely and regularly spaced **Cecropieae**

1. Stipules not amplexicaulous or amplexicaulous, where amplexicaulous (*Zhengyia*) not conical and not enclosing a well-formed leaf; cystoliths present in the mesophyll (only visible in dried material); secondary veins of leaf laminae parallel or not, conspicuously closely and regularly spaced or not **2**

2. Shrubs or small trees; trichomes present, trichomes neither hooked or with bulbous bases; fruit fleshy, **achene and ovary borne at an angle to the main axis of the perianth**, broadest above the midpoint (exception: *Maoutia*); Indomalaya, Australasia, Oceania **Leukosykeae**

2. Shrubs, trees, *lianas, hemi-epiphytes* or *herbs*; trichomes present or absent, where present trichomes not hooked or hooked, with or without bulbous bases; fruit fleshy or not, **achene and ovary aligned with the main axis of the perianth,** broadest below the midpoint (exceptions: Chamabainia, *Phenax, Debregeasia*, *Pipturus*); Palearctic, Afrotopics, Indomalaya, Australasia, Oceania, Nearctic or Neotropics **3**

3. Stems not releasing an exudate when cut, plant with **hooked trichomes** and never with a bulbous base; pistillode pubescent **4**

3. Stems releasing a clear or weakly opaque exudate when cut, or not; plant **with trichomes not hooked** and with or without a bulbous base; pistillode glabrous **5**

4. Stipules two at each node, occasionally one (*Astrothalamus, Debregeasia*, *Margarocarpus* sanguinea, *Neraudia, Pouzolzia*); inflorescences, or at least flower-clusters unisexual (rarely some *Boehmeria* species with some bisexual clusters also present), never forming reduced flower-like structures **Boehmerieae**

4. Stipules one at each node, or absent (*Parietaria, Metatrophis*); inflorescences or flowers bisexual, often forming a reduced flower-like structure (*Australina, Didymodoxa, Droguetia, Forsskaolea*) **Parietarieae**

5. Stems releasing a clear or weakly opaque exudate when cut**; cystoliths in the mesophyll punctiform, or occasionally elliptic**; some of the trichomes on stem and or leaf or inflorescences with a bulbous base or not (*Poikilospermum, Scepocarpus*, *Touchardia*) but where not then always a tree or vine; female sepals unequal or completely fused forming a tube **Urticeae**

5. Stems releasing a clear or weakly opaque exudate when cut, or not, where releasing a clear exudate male inflorescence racemose; **cystoliths in the mesophyll fusiform**; trichomes on stem and or leaf or inflorescences never with a bulbous base; female sepals equal or not (*Pilea*), never completely fused to form a tube **6**

6. Herbs, occasionally shrubs or epiphytes, never trees; **stem not releasing exudate when cut**; male inflorescence paniculate or capitate, rarely racemose (*Achudemia myriantha*), **male flower sepals bearing a subapical appendage;** Palearctic, Afrotopics, Indomalaya, Australasia, Oceana, Nearctic, or Neotropics **Elatostemateae**

6. Woody shrubs or small trees; **stems releasing clear exudate when cut**; male inflorescence racemose; **male flower sepals not bearing a subapical appendage**; Neotropics **Myriocarpeae**

### Keys to genera

#### Urticeae

1. Trees or hemi-epiphytes; **leaves entire or weakly and irregularly sinuate** 2

1. Herbs, shrubs, or trees; **Leaves toothed, lobed or laciniate** 5

2. Plant bearing trichomes with bulbous bases, or not; Inflorescence a branched panicle, the female sepals free for most of their length, never tubular ***Dendrocnide***

2. Plant often glabrous, where pubescent, lacking trichomes with bulbous bases, the female inflorescence one or more densely capitate and often brightly coloured heads, the female sepals fused for all or most of their length forming a tube ***Poikilospermum***

3. Cystoliths fusiform 3

3. Cystoliths punctiform

4. Herbs; stems not woody; stipules intrapetiolar, undivided; perianth not fleshy in fruit ***Nanocnide***

4. Trees, shrubs or lianas; stems woody; stipules interpetiolar, divided, at least at apex; perianth fleshy in fruit ***Urera***

5. Climbers or repent herbs ***Scepocarpus***

5. Herbs, trees or shrubs 6

6. Trees 7

6. Shrubs or herbs 9

7. Afrotropics; leaves deeply lobed, laciniate or profoundly serrate, where serrate teeth of fully developed leaves ≥ 5 mm, cystoliths punctiform ***Obetia***

7. Indomalaya, Australasia or Oceania; leaf margin toothed, teeth < 5 mm deep, cystoliths punctiform or fusiform 8

8. Hawaii; glabrous; inflorescence ***Touchardia***

8. Indomalaya, Australasia or Oceania (but not Hawaii); pubescent or glabrous, where pubescent hairs strongly *Urtica* ting ***Dendrocnide***

9. Stipules amplexicaulous and leaf-like ***Zhengyia***

9. Stipules never amplexicaulous, never leaf-like 10

10. Stipules intrapetiolar (borne in axis of petiole) 11

10 Stipules interpetiolar (borne at the side of the petiole) 12

11. Leaves alternate, <30 cm tall, single undivided stipule at each node ***Nanocnide***

11. Leaves opposite, > 20 cm tall, two stipules at each node ***Urtica***

12. Leaves serrate and deeply divided, lobed, laciniate or undivided; at least some trichomes > 5 mm; female sepals 3, partially fused ***Girardinia***

12. Leaves serrate, never deeply divided, lobed, laciniate; trichomes ≤ 5 mm; female sepals 4, free 13

13. Female inflorescence raceme-like, unbranched or with <4 branches; filaments partially adnate to the adjacent sepal; sepals equal or longer than the achene in fruit ***Sceptrocnide***

13. Female inflorescence a branched panicle with > 4 branches; filaments not adnate to the adjacent sepal; sepals shorter than the achene in fruit 14

14. Stipule scars inconspicuous; pedicels of female flowers articulated; stigma capitate or sub-caudiform; male sepals partially fused (at base) ***Fleurya***

14. Stipule scars conspicuous; pedicels of female flowers not articulated; stigma caudiform; male sepals free ***Laportea***

#### Myriocarpeae

1. Laminae not asperous; female inflorescences few branched and raceme-like, > 6 cm in length; female flowers with a foot-shaped stigma; no part of infructescence fleshy ***Myriocarpa***

1. Laminae asperous; female inflorescences not raceme-like, < 6 cm in length; female flowers with a penicillate a stigma, peduncle branches becoming fleshy in fruit ***Gyrotaenia***

#### Elatostemateae

1. Leaves opposite or spirally arranged; Palearctic, Afrotopics, Indomalaya, Australasia, Oceania, Nearctic or Neotropics 2

1. Leaves alternate or opposite, where opposite one of the leaves in each pair minute and soon lost, never spirally arranged; Paleactica, Afrotropics, Indomalaya, Australasia, Oceania, but never Nearctic or Neotropics 4

2. Achene bearing a crescent-shaped protuberance; Indomalaya ***Lecanthus***

2. Achene bearing a crescent-shaped protuberance; Palearctic, Afrotopics, Indomalaya, Australasia, Oceania, Nearctic or Neotropics

3. Male and female flowers 5-parted; Palearctic, Indomalaya ***Achudemia***

3. Male and female flowers 2-, 3- or 4-parted; Palearctic, Afrotopics, Indomalaya, Australasia, Oceania, Nearctic or Neotropics ***Pilea***

4. Stipules intrapetiolar (borne in axis of petiole); female flower sepals equal, inflorescence bracts absent; peduncle fleshy in fruit; achene surface smooth ***Procris***

4. Stipules interpetiolar (borne at the side of the petiole); female flower sepals equal or unequal, inflorescence bracts present; peduncle not fleshy in fruit; achene surface ornamented, never smooth 5

5. Male inflorescence symmetrical; female inflorescence a panicle, not involucrate ***Polychroa***

5. Male inflorescence asymmetrical; female inflorescence involucrate or not 6.

6. Stem succulent; leaves 3-nerved or penninerved, female inflorescence involucrate, achene not compressed ***Elatostema***

6. Stem woody or succulent; leaves 3-nerved, female inflorescence not involucrate, achene compressed ***Elatostematoides***

#### Leukosykeae

1. Stipules not forked; female inflorescence fused to form a coral-like structure, bearing > 1000 flowers, flowers sessile ***Sarcochlamys***

1. Stipules forked; female inflorescence a panicle, partially fused or not, but never fully fused or forming a coral-like structure, bearing < 1000 flowers, flowers sessile female flowers pedicellate **2**

2. Stipule scars conspicuous; male and female inflorescences once dichotomously branched, comprising two tight heads bearing < 500 flowers ***Leukosyke***

2. Stipule scars inconspicuous; male and female inflorescences branched dichotomously once or more, or paniculate, where dichotomously branched flowers borne in loose clusters and not tight heads **3**

3. Filaments inflexed in bud; female inflorescence involucrate, bearing <50 flowers, stigma sessile; achene compressed ***Gibbsi***

3. Filaments not inflexed in bud; female inflorescence not involucrate, bearing > 50 flowers, stigma stalked; achene not compressed ***Maoutia***

#### Cecropieae

1. Afrotropics, Indomalaya **2**.

1. Neotropics **4**

2. Stipules not enclosing another shoot iteration; leaf margin prominently serrate ***Myrianthus***

2. Stipules enclosing another shoot iteration which itself includes a visible stipule; leaf margin entire or irregularly sinuate **3**

3. Leaves palmately compound, petiolules winged; male inflorescence a panicle of fused capitae ***Musanga***

3. Leaves palmately lobed; male inflorescence a fascicle of spadices ***Cecropia***

4. Leaves palmately lobed; female and male inflorescence a fascicle of fused spadices ***Cecropia***

4. Leaves palmately lobed or entire; female inflorescence paniculate, male inflorescences paniculate or spicete, flowers borne in capitae, panicles or spikes **5**

5. Hemi-epiphytes; both female and male flowers borne in fused capitae borne on dichotomously branching panicles ***Coussapoa***

5. Trees or hemi-epiphytes; both female and male flowers free, borne in compact irregularly branched panicles or spikes ***Pourouma***

#### Parietarieae

1. Shrubs; leaves subcoriaceous, lanate below; male sepals lacking a sub-apical appendage; Afrotropics, Palearctic, Oceania **2**

1. Herbs; leaves membranous to chartaceous; glabrous or pubescent below but never lanate; male sepals with a sub-apical appendage; Afrotropics, Indomalaya, Australasia **3**

2. Cystoliths fusiform; stipules absent; Oceania (Austral Islands)

2. Cystoliths punctiform; stipules present; Afrotropics, Palearctic ***Metatrophis Forsskaolea***

3. Stipules absent; leaf margin entire

3. Stipules present; leaf margin crenate, serrate or entire

4 Parietaria

4. Cystoliths punctiform; stipules two at each node; inflorescence bracts fused almost to apex forming involucre ***Droguetia***

4. Cystoliths punctiform or fusiform; stipule one at each node; inflorescence bracts free **5**

5. Cystoliths punctiform, Afrotropics, Indomalaya ***Didymodoxa***

5. Cystoliths fusiform or oblongiform, Australasia (Australia, New Zealand) ***Australina***

#### Boehmerieae

1. Achenes associated with a fleshy structure derived from the perianth, pedicel or peduncle; Afrotopics, Indomalaya, Australasia, Oceania **2**

1. Achenes not associated with a fleshy structure derived from the perianth, pedicel or peduncle **6**

2. Infructescence a panicle with the fused sub-terminal and terminal branches, resembling a branching coral; Southeast Asia (Borneo, Philippines). ***Astrothalamus***

2. Infructescence a panicle or fascicle, where paniculate, branches not fused. Afrotopics, Indomalaya, Australasia, Oceania **3**

3. Stem not releasing a clear or greyish exudate when cut; hooked hairs present at shoot tips, petiole, lamina or inflorescence; tree or shrub; Oceania (Hawaii) ***Neraudia***

3. Stem releasing a clear or greyish exudate when cut; hooked hairs absent, or glabrous from any part of plant; tree, vine or shrub; Afrotopics, Indomalaya, Australasia, Oceania **4**

4. Fruit bright orange or red; only persistent stigma visible ***Debregeasia***

4. Fruit white, pale grey or cream; most or some part of achene clearly visible, stigma visible or not **5**

5. Trees or shrubs; stipule scars inconspicuous; female flowers pedicellate, perianth accrescent; infructescence clearly branched or sessile, fruit borne in compact heads bearing < 10 units; achene broadest at or below midpoint ***Oreocnide***

5. Trees, shrubs or vines; stipule scars conspicuous; female flowers sessile, perianth not accrescent; infructescence unbranched or sessile, fruit borne in compact heads bearing >15 units; achene broadest above midpoint ***Pipturus***

6. Achene white, cream or pale brown, shiny **7**

6. Achene brown, dark brown or black, shiny or matt **11**

7. Single undivided stipule associated with each leaf; leaf lamina toothed or entire; male sepals with subapical appendages or not **8**

7. Two stipules associated with each leaf; leaf lamina toothed; male sepals bearing a sub-apical appendage **9**

8. Stipules intrapetiolar; leaf lamina toothed; male flowers with subapical appendages; female flower with stigma articulated; achene bilaterally asymmetrical ***Margarocarpus***

8. Stipules interpetiolar; leaf lamina entire or toothed; male sepals lacking subapical appendages; female flower with stigma not articulated; achene bilaterally symmetrical***Pouzolzia***

9. Stipules broader than long; leaf lamina toothed; fruit surface neither ribbed nor winged; achene bilaterally asymmetrical ***Leptocnide***

9. Stipules longer than broad; leaf lamina toothed or entire; fruit surface ribbed or winged; achene bilaterally symmetrical **10**

10. Stipules scars conspicuous; lamina toothed; pubescence never comprising hooked hairs; male flowers sessile; Australasia (Lord Howe Islands, Kermadec Islands) ***Pouzolziella***

10. Stipules scars inconspicuous; lamina toothed or entire; pubescence comprising hooked as well as not hooked hairs, sometimes hooked hairs restricted to inflorescence or young stems; male flowers pedicellate ***Pouzolzia***

11. Leaf margin entire; achene shiny, easily detached from the perianth in fruit **12**

11. Leaf margin toothed; achene not shiny **13**

12. Leaves alternate or opposite, where opposite alternate leaves also present, leaves one or two at each node; lateral ‘primary’ veins extending ½ to ¾ the lamina length; achene brown ***Pouzolzia***

12. Leaves opposite, two or three at each node; lateral ‘primary’ veins extending the full length of the lamina, including the apex; achene black ***Gonostegia***

13. Hooked hairs absent from plant; stipules intrapetiolar **14**

13. Hooked hairs present on leaves, young stems, inflorescences or perianth; stipules interpetiolar **15**

14. Leaves lanate below; female perianth fused, extending for most of the length of the flower, hairs borne only on one side of the stigma; male flowers sessile, sepals not bearing sub-apical appendages; achene subcompressed, bilaterally symmetrical ***Muimar***

14.Leaves pubescent but never lanate below; female perianth absent; hairs borne the full circumference of the stigma; male flowers pedicellate, sepals bearing sub-apical appendages; achene compressed, bilaterally asymmetrical ***Phenax***

**15.** Small herbs; female flowers borne in paniculate cymes; stigma capitate, male flowers solitary, achene broadest above the midpoint ***Chamabainia***

15. Shrubs, small trees or herbs; female flowers in compacted paniculate cymes that appear fasciculate; stigma extended, liguliform, corkscrew-like or filiform; male flowers borne in paniculate or fused cymes; achene broadest below or at the midpoint **16**

16. Leaves alternate or opposite; female and male flowers borne in sessile, dense fascicle-like cymes either along the stem or along modified branches; stigma corkscrew-like or filiform, hairs borne the full length of the extended stigma ***Boehmeria***

16. Leaves alternate; female and male flowers borne in capitae in paniculate cymes; stigma ligulate, hairs borne the full length of the extended stigma ***Archiboehmeria***

### Synopsis of the genera

#### URTICEAE. Fig. 6

1. ***Urtica*** L. in Sp. Pl. 2: 983. 1753 – Type (designated by Green, 1929): *Urtica dioica* L. *= Hesperocnide* Torr. in Pacif. Rail. Rep. iv. 139. 1856 – Type: *Hesperocnide tenella* Torr. **Species and distribution**: 70 spp., global.

**Notes**: *Hesperocnide* has been repeatedly recovered within *Urtica* (Wu & al., 2013; Kim & al., 2015). As one of its two species lack a combination with *Urtica,* we make it here.

***Urtica tenella*** (Torr.) A.K.Monro **comb. nov. ≡** *Hesperocnide tenella* Torr. in Pacif. Railr. Rep. Whipple, Bot. 4(5; 4): 139. 1857. Holoype: USA, California, Napa valley, April 25 1853, *Bigelow s.n.* (NY barcode NY00284326; isotypes, K barcode K000708597, GH barcode GH 00035109).

2. ***Zhengyia*** T.Deng, D.G.Zhang & H.Sun in Taxon 62: 94. 2013 – Type: *Zhengyia shennongensis* T.Deng, D.G.Zhang & H.Sun.

**Notes** Indomalaya; 1 sp.

3. **Sceptrocnide** Maxim. in Bull. Acad. Imp. Sci. Saint-Pétersbourg 22: 238. 1876 – Type: *Sceptrocnide macrostachya* Maxim. in Bull. Acad. Imp. Sci. Saint-Pétersbourg 22: 240. 1876.

***Sceptrocnide cuspidata*** (Wedd.) A.K.Monro & I. Friis ***comb. nov.*** ≡ *Laportea cuspidata* (Wedd.) Friis in Kew Bull. 36: 156. 1981. ≡ *Girardinia cuspidata* Wedd. in Prodr. [A. P. de Candolle] 16(1): 103. 1869 – **Lectotype (designated here)**: China, Pekin [Beijing], *R.P. David 21* [2278] (P barcode P06816094; isotype: P barcode P06816093).

***Sceptrocnide interrupta*** (L.) A.K.Monro & I. Friis ***comb. nov.*** ≡ *Laportea interrupta* (L.) Chew in Gard. Bull. Singapore 21: 200 (1965). ≡ *Schychowskya interrupta* (L.) W.Wight in Contr. U.S. Natl. Herb. 9: 371 (1905). ≡ *Boehmeria interrupta* (L.) Willd. in Sp. Pl., ed. 4, 4: 340 (1805) ≡ *Urtica interrupta* L. in Sp. Pl.: 985 (1753). – Lectotype (selected by Corsi *et al*. 1999 ??): “Habitat in India.” [India], Herb. Hermann 3: 1, No. 336 (BM barcode BM-000594643).

**Notes** Indomalaya; 1 spp.

Molecular studies have consistently recovered *Laportea* as paraphyletic (Kim et al., 2015c; Wells et al., 2021; Wu et al., 2013). Whilst the taxa are morphologically similar, they may be distinguished based on combinations of stipule, inflorescence, stigma and fruit characters as summarised in Table 4.

**Table 4.**
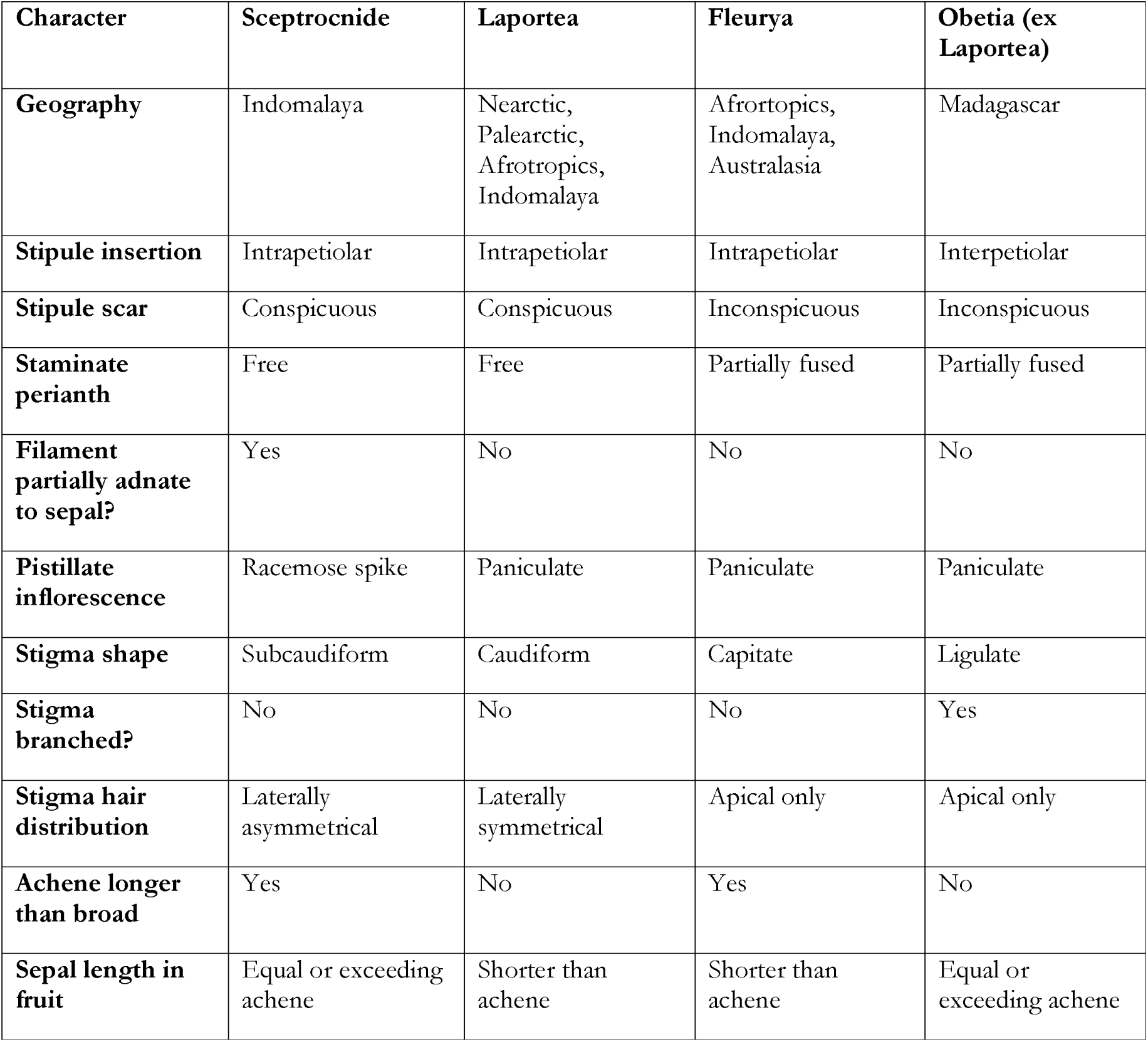
Diagnostic characters for *Sceptrocnide*, *Laportea, Fleurya* and Madagascan *Obetia* previously ascribed to *Laportea*.

[Table 4. Diagnostic characters for *Sceptrocnide*, *Laportea, Fleurya* and Madagascan *Obetia* previously ascribed to *Laportea.*]

4. ***Nanocnide*** Blume in Mus. Bot. 2: 154. 1856 – Type: *Nanocnide japonica* Blume in Mus. Bot. 2: 155. 1856. Type: Blume, *Mus. Bot.* 2: t. 17. 1856.

**Notes** Indomalaya, Palearctic; 3 sp.

5. ***Laportea*** Gaudich. in Voy. Uranie: 498. 1830, nom. cons. – Type: *Laportea canadensis* (L.) Wedd. in Ann. Sci. Nat. Bot., sér. 4. 1: 181. 1854. ≡ *Urtica canadensis* L. in Sp. Pl. 2: 985. 1753.

*Oblixilis* Raf. in Fl. Tellur. 3: 49. 1837 – **Lectotype (selected here)**: *Oblixilis canadensis* (L.) Raf. in Fl. Tellur. 3: 49. 1837.

*Urticastrum* Heist. ex Fabr. in Enum.: 204. 1759, nom. rej.

*Sclepsion,* Raf., ined.

**Notes** Nearctic, Palearctic, Afrotropics, Indomalaya; ca 5 spp. Several studies have consistently recovered ***Laportea*** as paraphyletic across the Urticeae tribe (Kim et al., 2015c; Wells et al., 2021; Wu et al., 2013). Whilst the taxa are morphologically similar, they may be distinguished based on combinations of stipule, inflorescence, stigma and fruit characters as summarised in Table 4.

We recovered as paraphyletic (Laportea I — V, Fig.s 3 & 4). Four of these groupings had also been recovered by previous authors (Kim et al., 2015c; Wells et al., 2021; Wu et al., 2018). To some extent, the paraphyly of Laportea is an artefact of Ohwi’s (Ohwi, 1936) decision to combine Sceptrocnide and Laportea, and Chew’s (Chew, 1965) decision to combine Fleurya and Laportea. Laportea I, II and V (, Fig.s 3 & 4) can be diagnosed morphologically based on inflorescence, flower and fruit morphology (Table 4, also the see key to the Urticeae below). We therefore decided to revert to the earlier delimitations of these genera, resulting in groupings that are congruent with morphology, monophyly and geography. In the case of Laportea IV (L. humblotii) a morphological review suggested that it can be considered congeneric to Obetia, notably in the shared development of a large papery fruit.

6. ***Girardinia*** Gaudich. in Voy. Uranie: 498. 1830 – Type: *Girardinia leschenaultiana* Decne. in Voy. Inde 152. 1844 – **Lectotype (selected here)**: India, *Leschenault 54* (P barcode P00601956).

**Notes** Palearctic, Afrotropics, Indomalaya; 2 spp.

7. ***Discocnide*** Chew in Gard. Bull. Singapore 21: 207. 1965 – Type: *Discocnide mexicana* (Liebm.) Chew in Gard. Bull. Singapore 21: 208. 1965 – Holotype: Mexico, *Liebmann14222* (C barcode C10017123) *= Discocarpus* Liebm., Kongel. Danske Vidensk. Selsk. Naturvidensk. Math. Afh., ser. 2, 2: 308. 1851, nom. illegit., non Discocarpus Klotzsch (1841: 201).

**Notes** Neotropics; 1 sp.

8. **Dendrocnide** Miq. in Pl. Jungh. 29: 1851 – Type: *Dendrocnide costata* Miq. in Pl. Jungh. 30: 1851.

**Notes** Indomalaya, Australasia; ca 36 spp.

9. ***Obetia*** Gaudich. in Voy. Bonite, Bot., Atlas: t. 82. 1844 – Type: *Obetia ficifolia* Gaudich. in Voy. Bonite, Bot., Atlas: t. 82. 1844.

***Obetia floribunda*** (Baker) A.K. Monro & T. Wells, **comb. nov**. ≡ Laportea floribunda (Baker) Leandri in Ann. Inst. Bot.-Géol. Colon. Marseille 7–8: 24. 1950 ≡ *Pilea floribunda* Baker in Bull. Misc. Inform., Kew 1897: 280. 1897 – Holotype: Madagascar, Tanala, Ambohimitombo, *F. Major 270* (K barcode K000242862; isotypes: G barcode G00023431, G barcode G00023542)]

***Obetia humblotii*** (Baill.) A.K. Monro & T. Wells, **comb. nov**. ≡ Laportea humblotii (Baill.) Friis in Nordic J. Bot. 2: 233. 1982. ≡ *Urera humbloti* i Baill. in Bull. Mens. Soc. Linn. Paris 1: 479. 1885 – **Lectotype (selected here)**: Madagascar, Antsianaka, *M. Humblot 484* (P barcode P06457704; isolectotypes: LD barcode LD1216815, P barcode P06457701, P barcode P06457702, P barcode P06457703).

***Obetia oligoloba*** (Baker) A.K. Monro & T. Wells, **comb. nov**. ≡ Laportea oligoloba (Baker) Friis in Nordic J. Bot. 2: 233. 1982. ≡ Urera oligoloba Baker in J. Linn. Soc., Bot. 20: 265. 1883 – **Lectotype (selected here)**: Madagascar (Centre) [Imerina], *R. Baron* s.n. ‘Aug. 1881’ (K barcode K000242866; isolectotypes: K barcode K000242863, P barcode P00524247, P barcode P00435850).

***Obetia perrieri*** (Leandri) A.K. Monro & T. Wells, **comb. nov**. ≡ *Laportea perrieri* Leandri in Ann. Inst. Bot.-Géol. Colon. Marseille 7–8: 20–21. 1950 – **Lectotype (designated here)**: Madagascar, Diego-Suarez Province, l’Ankarana limestone plateau, *H. Humbert 18884,* (P barcode P00435843; isolectotypes: P barcode P00435842, P barcode P00435841).

**Notes** Afrotropics; 12 spp. Madagascan species ascribed to *Laportea* exhibit the combination of stipules adjacent to the petiole base, leaves with ‘box’ venation and fruit with papery rather than fleshy sepals.

Two representatives of this group (*L. humblotii* (Baill.) Friis, *L. floribunda* (Baker) Leandri) were included in this study and recovered as congeneric or sister to *Obetia.* Morphological examination of *Obetia* species, *L. humblotii, L. oligoloba* (Baker) Friis, *L. perrieri* Leandri suggests that they could be considered congeneric with *Obetia* which we propose here. The presence of intrapetiolar stipules and absence of cystoliths and urticating hairs in *L. floribunda,* however, suggests that it differs from *Obetia* and the other Madagascan species ascribed to *Laportea* by Chew. Further research may conclude that *L. floribunda* represents a distinct and undescribed genus but in the present classification we have included it in *Obetia*.

10. ***Scepocarpus*** Wedd. in Prodr. [A. P. de Candolle] 16: 98. 1869 – Type: *Scepocarpus mannii* Wedd. in Prodr. [A. P. de Candolle] 16: 98. 1869.

**Notes** Afrotropics; ca 15 spp. Distinguished from *Urera* by the asymmetrical pistillate inflorescences, pistillate sepals fused to a greater degree and Afrotropical distribution (Wells & al., 2021).

***Scepocarpus acuminatus*** (Poir.) T.Wells & A.K.Monro in Mol. Phylogenet. Evol. 158: 12. 2021. ≡ Urera acuminata (Poir.) Gaudich. ex Decne., Nouv. Ann. Mus. Hist. Nat. 3: 490. 1834. ≡ *Urtica* acuminata Poir., Encycl. [J. Lamarck & al.] Suppl. 4. 226. 1816. Validation of *Scepocarpus acuminatus* (Poir.) T.Wells & A.K.Monro described in Molec. Phylogen. Evol. 158: 12. 2020.

***Scepocarpus keayi*** (Letouzey) T.Wells & A.K.Monro **comb. nov.** ≡ Urera keayi Letouzey in Adansonia, n.s., 7: 299. 1967. **Lectotype (designated here)**: Ivory Coast, Lamé (40 km NE of Abidjan), 6 November 1958, *A.J.M. Leeuwenberg 1901* (WAG barcode WAG0003460; isotypes: K barcode K000242888, WAG barcode WAG0003461, WAG barcode WAG0003462, WAG barcode WAG0003463).

11. ***Urera*** Gaudich. in Voy. Uranie: 496. 1830, nom. cons. – Lectotype (selected by Britton & P.Wilson. in Scient. Surv. Porto Rico 1: 243, 1924): *Urera baccifera* (L.) Gaudich. ex Wedd. in Ann. Sci. Nat. Bot., ser. 3, 18: 199. 1852 ≡ *Urtica baccifera* L. in Sp. Pl., ed. 2, 2: 1398. 1763.

***Urera*** subgenus ***capitata*** [add authors] **subgen. nov**. Type: Urera capitata Wedd., Ann. Sci. Nat., Bot. sér. 3, 18: 201 (1852). Validation of *Urera* subgen. *Capitata* T.Wells & A.K.Monro described in Molec. Phylogen. Evol. 158: 14. 2020.

**Notes** The African species have been placed in the resurrected genus *Scepocarpus* (see above), and the Hawaiian species moved to *Touchardia* (below) as proposed by Wells & al. (Wells et al., 2021). Last monographed by Weddell (1856).

12. ***Poikilospermum*** Zipp. ex Miq. in Ann. Mus. Bot. Lugduno-Batavi 1: 203. 1864 – Type: *Poikilospermum amboinense* Zipp. ex Miq. in Ann. Mus. Bot. Lugduno-Batavi 1: 203. 1864.

=*Conocephalus* Blume in Bijdr. Fl. Ned. Ind.: 483. 1825 – Type: *Conocephalus suaveolens* Blume in Bijdr. Fl. Ned. Ind.: 483. 1825.

=*Balansaephytum* Drake in Bull. Soc. Bot. France 43: 82. 1896 – Type: *Balansaephytum tonkinense* Drake in Bull. Soc. Bot. France 43: 82. 1896.

=*Conocephalopsis* Kuntze in Revis. Gen. Pl. 3: 136. 1898, nom. nud.

**Notes** Indomalaya, Australasia; 27 spp. Mainly hemi-epiphytes. Morphologically very similar to *Touchardia.* Last monographed by Chew (Chew, 1963).

13. ***Touchardia*** Gaudich. in Bot. Voy. Bonite: t. 94. 1844 – Type: *Touchardia latifolia* Gaudich. in Bot. Voy. Bonite: t. 94. 1844.

***Touchardia kaalae*** (Wawra) T.Wells & A.K. Monro, **comb. nov.** ≡ *Urera kaalae* Wawra in Flora 57: 542. 1874.

**Notes** Nearctic (Hawaii); 3 spp.

14. ***Fleurya*** Gaudich. in Voy. Uranie: 497. 1830 – Lectotype (designated by Britton & Millspaugh, 1920: 106): *Fleurya paniculata* Gaudich. in Voy. Uranie: 497. 1830.

*=Schychowskya* Endl. in Ann. Wiener Mus. Naturgesch. 1: 187. t. 13. 1836 – Type: *Schychowskya ruderalis* (G. Forst.) Endl. in Ann. Wiener Mus. Naturgesch. 1: 187. t. 13. 1836 ≡ *Urtica ruderalis* G. Forst. in Fl. Ins. Austr. 66. 1786.

=*Parsana* Parsa & Maleki in Fl. Iran [Parsa] Suppl. Gén.: 548. 1952.

**Notes** Afrotropic, Indomalaya, Australasia; 4 spp. Chew (Chew, 1965) cited the generitype as *Fleurya spicata* Gaudich., but a generitype had already been designated by Britton & Millspaugh (1920).

Molecular studies have consistently recovered *Laportea* as paraphyletic (Kim et al., 2015c; Wells et al., 2021; Wu et al., 2013). Whilst the taxa are morphologically similar, they may be distinguished based on combinations of stipule, inflorescence, stigma and fruit characters as summarised in Table 4.

#### MYRIOCARPEAE. Fig. 7

15. ***Myriocarpa*** Benth. in Bot. Voy. Sulphur: 168. t. 55. 1844 – Type: *Myriocarpa stipitata* Benth. in Bot. Voy. Sulphur: 168. t. 55. 1844 – Type: Bot. Voy. Sulphur: t. 55. 1844. **Epitype (selected here)**: Colombia, Chocó, Nuquí, *A. Gómez 451* (HUA; isoepitype: MO).

**Notes** Neotropics; ca 14 spp. The Sanger molecular data presented here suggests that *Myriocarpa* may be polyphyletic with respect to *Gyrotaenia.* The two taxa are united by spike-like racemes but are morphologically distinct, *Myriocarpa* having long filament-like extended infructescences and dry fruit, *Gyrotaenia* having short (< 5 cm) infructescences and fleshy fruit. We have decided to maintain the taxa as distinct until additional samples have been analysed.

16. ***Gyrotaenia*** Griseb. in Pl. Wright. 1: 174. 1860 – Type: *Gyrotaenia myriocarpa* Griseb. in Pl. Wright. 1: 174. 1860.

**Notes** Neotropics; 5 spp. see above.

#### ELATOSTEMATEAE. Fig. 8

17. ***Elatostema*** J.R.Forst. & G.Forst. in Char. Gen. Pl., ed. 2. [105]. 1776. nom. con., typ. cons. (Sprague, 1929; Cambridge Code 1930: 70. 1988) – Type: *Elatostema sessile* J.R. Forst. & G. Forst. in Char. Gen. Pl., ed. 2.: 106. 1776. Type: Char. Gen. Pl., ed. 2., p. 53. 1776.

=*Langeveldia* Gaudich. in Voy. Uranie: 494. 1830 – Type: *Langeveldia acuminata* (Poir.) Gaudich. in Voy. Uranie: 495. 1830 ≡ *Procris acuminata* Poir. in Encycl. [J. Lamarck & al.] 5(1): 629. 1804. Type: [Indonesia] Isle du Java, *Commerson s.n.* (P-JU 16858!, P barcode P00663015).

**Notes** Afrotropics, Palearctic, Indomalaya, Australasia; *ca* 700 spp.

A clade comprising *Pellionia brachyceras, Elatostema penninerve, Elatostema filicinum* and *Elatostema bulbiferum* were recovered sister to *Procris* in our supertree of Sanger and Angiopserms353 data (Fig. 4.3). This clade comprises penninnerved leaved species that possess nanophylls, and female flowers whose sepals lack a sub-apical appendage that are in these respects different from *Elatostema* and species descibed under *Pellionia,* and similar to *Procris* (L. Fu, 2021; Wang, 2016). This clade differsdiffer morphologically from *Procris* in the absence of the subglobose capitate female inflorescence that is the main diagnostic character for the genus. The generitype of *Pellionia* (*P. elatostematoides*) shares the combination of penninnerved leaves and the presence of nanophylls, and should maybe be moved to this clade, which would then be named as *Pellionia,* and excluded from *Elatostema,* to which it was moved by Tseng et al. (Tseng et al., 2019). Material of *P. elatostematoides* has not, however, been sampled for sequence data by this or previous studies and so whilst we hypothesise that were it to be sequenced it would be recovered within this clade, in the absence of the molecular data we have left it un-named.

18. ***Elatostematoides*** C.B. Rob. in Philipp. J. Sci., C, 5: 497. 1911 – Type: *Elatostematoides manillense* (Wedd.) C.B. Rob. in J. Sci., C, 5: 501. 1911.

*Elatostematoides tsoongii* (Merr.) A.K.Monro **comb. nov.** ≡ Elatostema tsoongii (Merr.) H. Schroet. in Repertorium Specierum Novarum Regni Vegetabilis, Beihefte 83(2): 21. 1936. ≡ Pellionia tsoongii (Merr.) Merr. in Lingnan Science Journal 6(4): 325. 1928. ≡ Polychroa tsoongii Merr. in Philipp. J. Sci. 21: 493. 1922. Lectotype (designated here): China, Kwangtung Province [Guangdong], Tung Sing, June 22 1918, *K.K. Ts’oong 1870* (PE)

**Notes** Indomalaya, Australasia; ca 20 spp. Outside of the Philippines, *Elatostematoides* has been considered congeneric with *Elatostema* (Rodda & al., 2020) but based on Tseng & al.’s (Tseng et al., 2019) molecular phylogeny and morphological revision of the delimitation of *Elatostema* is recognized as a distinct genus here.

19. ***Procris*** Comm. ex Juss. in Gen. Pl. 403. 1789, nom. cons. – Type: *Procris cephalida* Comm. ex Poir. in Encycl. [J. Lamarck & al.] 5(1): 629. 1804, nom. cons.

*Sciophila* Gaudich., Voy. Uranie, Bot..: 493. 1830, nom. illegit., non Wibel 1799.

**Notes** Afrotopics, Indomalaya, Australasia; ca 20 spp.

20. ***Polychroa*** Lour. in Fl. Cochinch. 2: 559. 1790 – Type: *Polychroa repens* Lour. in Fl. Cochinch. 2: 559. 1790. ≡ *Elatostema repens* (Lour.) Haller f. in Ann. Jard. Bot. Buitenzorg 13: 316. 1896 ≡ *Pellionia repens* (Lour.) Merr. in Lingnan Sci. J. 6(4): 326. 1928 ≡ *Procris repens* (Lour.) B.J.Conn & Hadiah in Gard. Bull. Singapore 63(1-2): 160. 2011 – Type: *J. de Loureiro s.n.,* China] (type material not traced). **Neotype (selected here)**: China, Hainan, Lam Ko District, LinFa Shan, Siu Shui Kau, *W.-T. Tsang 262* (Lingnan University #15761) (K).

**Notes** Indomalaya; 1 sp. We have recovered *Polychroa* as sister to a clade comprising both *Elatostema, Elatostematoides* and *Procris* (Fig. xx), whilst previous studies (Fig. X, Tseng et al., 2019; Z. Y. Wu et al., 2013) recover it as sister, and so potentially congeneric with *Procris*. Morphologically *Polychroa* is distinct from *Elastostema* and *Procris* in its repent habit, the presence of prominent orange-brown papery stipules and the non-capitate / receptaculate pistillate inflorescences. Of note are the very few known collections bearing female inflorescences or infructescences.

We could not locate any material at BM corresponding to a type collection (Merrill, 1935; Wajer, pers. comm. 2021) and so designate a neotype from the type locality.

21. ***Lecanthus*** Wedd. in Ann. Sci. Nat., Bot., sér. 4, 1: 187. 1854 – Lectotype (designated by Hutchinson in Gen. Fl. Pl. 2: 186. 1967), *Lecanthus wightii* Wedd. in Ann. Sci. Nat., Bot., sér. 4, 1: 187. 1854. nom. illeg.

=*Meniscogyne* Gagnep. in Bull. Soc. Bot. France 75: 99. 1928 – Type: *Meniscogyne thorelii* Gagnep. in Bull. Soc. Bot. France 75: 99. 1928 – **Lectotype (designated here)**: Laos, *Thorel 2391* P barcode P00602029).

**Notes** Afrotropics, Indomalaya; 5 spp.

22. ***Achudemia*** Blume in Mus. Bot. Lugd.-Bat. 2: 57. 1856. Type: *Achudemia javanica* Blume in Mus. Bot. Lugd.-Bat. 2: 57. 1856.

=*Smithiella Dunn.* in Bull. Misc. Inform., Kew 1920: 210, 1920 – Generitype: *Smithiella myriantha* Dunn., nom. illeg., non *Smithiella* H. Perag. & Perag. ≡ *Aboriella* Bennet in Indian Forester 107: 437, 1981

≡ *Dunniella* Rauschert in Taxon 31: 562, 1982 – **Lectotype (designated here)**: India, *Burkill 37636* (K barcode K000708616).

**Notes** Palearctic, Indomalaya; five spp. We believe that the type material of *Metapilea* may be of an immature *Achudemia boniana* specimen (see L. Fu et al., 2022). Due to the sampling policy of the herbarium where the type collection is stored it is not possible to sample this material for DNA and so its status and position remain unknown.

**23 Pilea** Lindl. in Coll. Bot. ad t. 4. 1821, nom. cons. – Type: *Pilea muscosa* Lindl., nom. illeg. superfl. = *Parietaria microphylla* L. = *Pilea microphylla* (L.) Liebm.

*Adicea* Raf. in First Cat. Gard. Transylv. Univ.: 13. 1824, nom. nud.

*Adicea* Raf. ex Britton & A. Br. in Ill. Fl. N. U.S. 1: 533. 1896. nom. illeg., homonym.

*=Adike* Raf. in New Fl. 1: 63. 1836 – Type: *Adike pumila* (L.) Raf. in New Fl. 1: 63. 1836.

*=Chamaecnide* Nees & Mart. ex Miq., in Martius Fl. Bras. 4: 203. 1853 – Type: *Chamaecnide microphylla* Nees ex Miq.

*=Dubrueilia* Gaudich. in Voy. Uranie: 495. 1830 – Type: *Dubrueilia peploides* Gaudich. – **Lectotype (designated here)**: Hawaii, *Gaudichaud s.n.* (P, barcode P00636991).

*=Haroldiella* J.Florence in Fl. Polynésie française 1: 218. 1997 – Type: *Haroldiella sykesii* J. Florence in Fl. Polynésie française 1: 221. 1997.

*=Neopilea* Leandri in Ann. Mus. Colon. Marseille, sér. 6, 7-8: 46. 1950 – Type: *Neopilea tsaratananensis* Leandri.

*=Sarcopilea* Urb. in Symb. Antill. 7: 201. 1912. Generitype: *Sarcopilea domingensis* Urb. – **Lectotype (designated here)**: Dominican Republic, *Tuerckheim 3392,* (BR barcode BR0000005623624).

**Notes** Afrotropics, Palearctic, Indomalaya, Australasia (absent from Australia and New Zealand), Nearctic, Neotropics; *ca* 715 spp. The delimitation of *Pilea* and a subgeneric classification was recently proposed by Fu & al. (L. Fu et al., 2022). PARIETARIEAE. Fig. 9.

24. ***Parietaria*** L. in Sp. Pl. 2: 1052. 1753. Type: *Parietaria officinalis* L. in Sp. Pl. 2: 1052. 1753.

=*Freirea* Gaudich. in Voy. Uranie: 502. 1830 – **Lectotype (selectded here)**: *Freirea australis* Nees in Pl. Preiss. 1: 638. 1845 – **Lectotype (designated here)**: Australia, *Preiss 2399* (MEL 115271; isolectotypes: L barcode L0420597, LD barcode LD1513730, LE barcode LE00011441, LE barcode LE00011442, LE barcode LE00011455, MEL barcode MEL115373).

=*Thaumuria* Gaudich. in Voy. Uranie:502. 1830 – **Lectotype (designated here)**: *Thaumuria cretica* Gaudich. ex Wedd. in Ann. Sci. Nat., Bot., sér. 4, 1: 210. 1854 ≡ *Parietaria cretica* L. in Sp. Pl. 2: 1052. 1753.

=*Soleirolia* Gaudich. in Voy. Uranie: 504. 1830. Type: *Soleirolia repens* Kuntze in Revis. Gen. Pl. 2: 633. 1891. ≡ *Parietaria soleirolii* (Req.) Spreng. in Syst. Veg. ed. 16, 4(2): 318 (1827).

=*Gesnouinia* Gaudich. in Voy. Uranie: 502. 1830. Type: *Gesnouinia arborea* (L. f.) Gaudich. in Voy. Uranie: 502. 1830 ≡ *Boehmeria arborea* (L.f.) Desf. in Tabl. École Bot., ed. 3: 348. 1829 ≡ *Parietaria arborea* (L.f.) L’Hér. in J. Phys. Chim. Hist. Nat. Arts 33: 55. 1788 ≡ *Urtica arborea* L.f. in Suppl. Pl.: 417. 1782 – **Lectotype (designated here)**: Spain, *Masson* s.n. (BM barcode BM000829399).

**Notes** Global; 26 spp. *Parietaria* was recovered as polyphyletic (Schüßler et al., 2019) *Parietaria* was recovered as polyphyletic with respect to the island endemics *Gesnouinia* and *Soleirolia* (Fig. 3 & 4.1). The main differences between these three genera according to Friis and Weddell (Friis, 1989; Weddell, 1856), relate to habit and flower sexuality. *Parietaria* comprising pantropical erect or suberect herbs with bisexual flowers; *Gesnouinia,* a large woody shrub with unisexual flowers, endemic to Macaronesia; and *Soleirolia,* a minute creeping herb with unisexual flowers and winged fruits, endemic to Corsica in the Mediterranean. A closer evaluation of the taxa reveals a number of shared morphological characters such as, an absence of stipules, prominently bracteate inflorescences and the bracts associated with pistillate flowers forming tubular lobed involucres that transform markedly in fruit. Applying our criteria for genus delimitation, therefore, we treat all three genera as *Parietaria*.

*Parietaria debilis* G.Forst. and *P. officinalis* are remarkable in the Urticaceae for being gynomonoecious, bearing unisexual and bisexual flowers, which can be found on the same inflorescence, and which have dimorphic stigmas and fruiting perianths. Gynomonoecy has only been documented in a few *Parietaria* species but understanding how it is determined could enable the elaboration of testable hypotheses on the evolution of monoecy in the Urticales. *P. debilis* (including *P. pensylvanica*) is notable for its global distribution which encompasses temperate climates in both the temperate and tropical latitudes of the southern and northern hemispheres.

25. ***Forsskaolea*** L. in Opobalsamum: 17. 1764 ≡ *Chamaedryfolia* Kuntze in Revis. Gen. Pl. 2: 625. 1891 – Type: *Forsskaolea tenacissima* L. in Opobalsamum: 18. 1764.

=*Caidbeja* Forssk. in Fl. Aegypt.-Arab.: 82. 1775 – Type: *Caidbeja adhaerens* Forssk. in Fl. Aegypt.-Arab.: 82 (1775) – **Lectotype (designated here)**: Egypt, *Forsskål* s.n. (C barcode C10001905).

**Notes** Afrotropics, Indomalaya; 7 spp. Last revised by Friis & Wilmot-Dear (Friis & Wilmot-Dear, 1988).

26. ***Didymodoxa*** E. Mey. ex Wedd. in Monogr. Fam. Urtic.: 547. 1856 – Type: *Didymodoxa acuminata* (Wedd.) Wedd. in Monogr. Fam. Urtic.: 549. 1856.

**Notes** Afrotropics; 2 spp. Last revised by Friis & Wilmot-Dear (Friis & Wilmot-Dear, 1988).

27. ***Australina*** Gaudich. in Voy. Uranie: 505. 1830. Type: *Australina pusilla* (Poir.) Gaudich. in Voy. Uranie: 505. 1830.

=*Didymotoca* E.Mey. in Zwei Pflanzengeogr. Docum. (Drège): 178 1843, nom. nud.

=*Anaganthos* Hook.f. in Fl. New Zealand 1: 226. 1854, nom. inval.

**Notes** Australasia; 1 sp. Last revised by Friis & Wilmot-Dear (1988). *Australina,* as delimited by Weddell (Weddell, 1869) and Friis (Friis, 1989), included two species, *A. flaccida* (A. Rich.) Wedd., and *A. pusilla* that occurr in east Africa and Australasia respectively. *Australia flaccida* are opposite-leaved and *A. pusilla* alternate-leaved. Our analyses (Fig. xx) recover *A. flaccida* as sister to the Afrotropical *Droguetia,* to which we have moved it (see *Droguetia* below), and the type, *A. pusilla* as sister to a clade comprising *Droguetia* and *Metatrophis*.

28. ***Metatrophis*** F.Br. in Bull. Bernice P. Bishop Mus. 130: 34. 1935. Type: *Metatrophis margaretae* F.Br. in Bull. Bernice P. Bishop Mus. 130: 35. 1935.

**Notes** Australasia (Austral islands, Polynesia); 1 sp. First ascribed to Moraceae (Brown, 1935) based on the catkin-like inflorescences. Florence (1997) correctly moved it to the Urticaceae based on the basal placentation of the ovary and the position of the stamens in bud. With respect to flower morphology and arrangement, *Metatrophis* is very similar to *Australina* (as delimited here), the main difference being the arrangement of the male glomerules into spikes in *Metatrophis,* as opposed to free in *Australina.* The two genera, however, differ greatly morphologically. Further research may decide that *Metratrophis* and *Australina* be combined into a single genus.

Whilst both Brown (Brown, 1935) and Florence (Florence, 1997) describe *Metatrophis* as having caducous stipules we could find neither stipules or stipule scars on the single sheet at K, or images available at JSTOR Global Plants (BISH10052652022, BISH1005268, BISH1005269, P00636979). We believe, therefore, that Brown and Florence may have mistaken very immature leaves for stipules. The presence of stipules should be resolvable from the examination of living material. If confirmed, the absence of stipules is a trait shared with *Parietaria*.

29. ***Droguetia*** Gaudich. in Voy. Uranie: 505. 1830. Type: *Droguetia leptostachys* Gaudich. in Voy. Uranie: 505. 1830.

=*Didymogyne* Wedd. in Ann. Sci. Nat., Bot., sér. 4, 1: 207. 1854, nom. illeg., superfl. (Friis & Wilmot-Dear, 1988) – Type: *Didymogyne abyssinica* Wedd in Ann. Sci. Nat., Bot., sér. 4, 1: 207. 1854.

***Droguetia flaccida*** (A.Rich.) A.K. Monro, I. Friis & Wilmot-Dear, **comb. nov**. ≡ *Australina flaccida* (A.Rich.) Wedd. in Prodr. [A.P.de Candolle] 16(1): 23560. 1869 ≡ *Pouzolzia flaccida* A.Rich. in Tent. Fl. Abyss. 2: 259. 1850 – **Lectotype (designated here)**: Abyssinia [Ethiopia], *R. Quartin-Dillon* s.n. ‘Sept. 1839’ (P barcode P00435811; isolectotypes: P barcode P00435810, P barcode P00435812).

**Notes** Afrotropics, Indomalaya; 8 spp. Last revised by Friis & Wilmot-Dear (Friis & Wilmot-Dear, 1988). In the light of changes to the delimitation of *Australina* and *Droguetia* suggested here, *Droguetia* can now be distinguished from *Australina* based on the presence of punctiform cystoliths versus fusiform or oblongiform in the latter.

#### BOEHMERIEAE. Fig. 10

30. ***Oreocnide*** Miq. in Pl. Jungh. 39.1851 – **Lectotype (designated here)**: *Oreocnide silvatica* (Blume) Miq. in Miq., Pl. Jungh.: 40. 1851. ≡ *Urtica sylvatica* Blume in Bijdr. Fl. Ned. Ind. 506. 1825.

=*Villebrunea* Gaudich. in Voy. Bonite, Bot., Atlas: t. 91. 1844 – **Lectotype (designated here)**: *Villebrunea integrifolia* Gaudich. in Voy. Bonite, Bot., Atlas: t. 91. 1844. **Notes**ϑ Palearctic, Indomalaya, Australasia; 16 spp.

31. ***Phenax*** Wedd. in Ann. Sci. Nat. Bot. sér. 4. 1: 191. 1854 – Type: *Phenax vulgaris* Wedd., Ann. Sci. Nat. Bot., sér. 4. 1: 191. 1854 ≡ *Gesnouinia boehmerioides* Miq. in Martius, Fl. Bras. 4: 194. 1853.

**Notes** Neotropics, Afrotropics (Madagascar); 28 spp. *Phenax* remains very poorly studied with no comprehensive revision since Weddell (Weddell, 1869). The disjunct distribution of the Neotropics and Madagascar, together with very few species having been sampled for DNA suggest that *Phenax* should be a high priority for future study.

32. **Boehmeria** Jacq. in Enum. Syst. Pl. 9, 31. – Type: *Boehmeria ramiflora* Jacq. in Enum. Syst. Pl. 9: 31. 1760.

=*Cypholophus* Wedd. in Ann. Sci. Nat., Bot., sér. 4, 1: 198. 1854 – Type: *Cypholophus macrocephalus* (Blume) Wedd. in Ann. Sci. Nat., Bot., sér. 4, 1: 198. 1854.

*=Duretia* Gaudich. in Voy. Uranie: 500. 1830, nom. nud.

*=Splitgerbera* Miq. in Comm. Phytogr.: 133. t. 14. 1840. Type: *S. japonica* Miq. in Comm. Phytogr.: 134. 1840.

***Boehmeria brunneola*** (Elmer) A.K.Monro & Wilmot-Dear **comb. nov.** *≡ Cypholophus brunneolus* Elmer in Leafl. Philipp. Bot. 3: 896. 1910 ― Holotype: Philippines, Mindanao, Davao District, Todaya (Mt. Apo), 1200 masl, Sept. 1909, *A.D.E. Elmer 11641* (A barcode A00039518*).

***Boehmeria chamaephyton*** (H.J.P.Winkl.) A.K.Monro & Wilmot-Dear **comb. nov.** *≡ Cypholophus chamaephyton* H.J.P.Winkl. in Bot. Jahrb. Syst. 57: 582. 1922. Type: Papua New Guinea, Madang Province] Mount Hellwig summit, 2500 m, Nov. 1909, *L.S.A.M. von Römer 1213* (B†).

***Boehmeria decipiens*** (H.J.P.Winkl.) A.K.Monro & Wilmot-Dear **comb. nov.** *≡ Cypholophus decipiens* H.J.P.Winkl. in Bot. Jahrb. Syst. 57: 573. 1922 – Lectotype (designated by Wilmot-Dear & Friis, 1998: 922): Papua New Guinea, Kaiser Wilhelmsland, 1889ℒ1891, *K. Weinland 169* (L barcode L0281467*).

***Boehmeria engleriana*** (H.J.P.Winkl.) A.K.Monro & Wilmot-Dear **comb. nov.** *≡ Cypholophus englerianus* in H.J.P.Winkl. in Bot. Jahrb. Syst. 57: 578. 1922 – Syntypes: Indonesia, Papua, Mt. Hellwig summit, 2500 m, 5 Jan. 1913, *A.A.Pulle 919, 920* (B†, WRSL*).

***Boehmeria friesiana*** (K.Schum.) A.K.Monro & Wilmot-Dear **comb. nov.** *≡ Pilea friesiana* K.Schum. in K.M.Schumann & C.A.G.Lauterbach in Fl. Schutzgeb. Südsee, Nachtr.: 252. 1905 ≡ *Cypholophus friesianus* (K.Schum.) H.J.P.Winkl. in Bot. Jahrb. Syst. 57: 584. 1922 – Holotype: [Papua New Guinea, Morobe Province, Huon Peninsula] Kaiser Wilhelmsland, Sattelberg, June 1899, *E.O.A. Nyman 420* (B†; isotype: L!, UPS, WRSL*?).

Material corresponding to Nyman 420 was seen at WRSL but it lacks collector labels. Winkler deposited a set of duplicatesupplicate type collections at WRSL for names that he described from New Guinea (1922).

***Boehmeria gjellerupii*** (H.J.P.Winkl.) A.K.Monro & Wilmot-Dear **comb. nov**. *≡ Cypholophus gjellerupii* H.J.P.Winkl. in Bot. Jahrb. Syst. 57: 580. 1922 ― Holotype: Papua New Guinea, upper Tami river between Eti and Arfom rivers, 80 m, 27 Mar. 1910, *K. Gjellerup 8* (B†; isotype: WRSL*).

***Boehmeria harveyi*** Seem. Fl Vitiens. t. 62 (1868). Replaced name: Cypholophus heterophyllus (Wedd.) Wedd. in Prodr. [A. P. de Candolle] 16(1): 23518 (1869). *≡* Cypholophus macrophyllus var. heterophyllus Wedd (1857:435.], non *Boehmeria heterophylla* Wedd. ― **Type:** Fl Vitiens. t. 62 (1868).

Seemann’s name has only a colour plate as description and type (1868). A specimen collected by Harvey in Fiji in 1855 (*s.n.,* BM, K) is annotated ‘Gov.Miss Fiji 1st Append.’. This would suggest that there may be an earlier place of publication. We have, however, been able to locate this publication

***Boehmeria hochreutineri*** A.K.Monro & Wilmot-Dear **nom. nov.** Replaced name: *Cypholophus stipulatus* Hochr. in Candollea 2: 348 (1925) non *Boehmeria stipularis* Wedd. in Ann. Sci. Nat., Bot., sér. 4, 1: 200. 1854 ― Holotype: Samoa, Upola Island, road to Lanuto Lake, 500 m, Mar. 21 1905, *B.P.G. Hochreutiner 3248* (G barcode 00354082*).

***Boehmeria integer*** (H.J.P.Winkl.) A.K.Monro & Wilmot-Dear **comb. nov.** *≡ Cypholophus integer* H.J.P.Winkl. in Bot. Jahrb. Syst. 57: 584. 1922 – Holotype: [Indonesia, Sandaun, East Sepik] Sepik area, Felsspitzec station, 1400 to 1500 m, Aug. 4 1913, *C.L. Ledermann 12587* (B†, WRSL*)

***Boehmeria kerewensis*** (P.Royen) A.K.Monro & Wilmot-Dear **comb. nov.** *≡ Cypholophus kerewensis* P.Royen in Alp. Fl. New Guinea 3: 2137. 1982 ― Holotype: Papua New Guinea, South Highland District, Tari Subdistrict, N slopes of Mt. Kerewa, 6°S 143°E, 2940 m, June 29 1966, *C. Kalkman 4720* (BISH barcode BISH1005216; isotype: K!).

***Boehmeria lauterbachiana*** A.K.Monro & Wilmot-Dear **nom. nov.** Replaced name: *Cypholophus velutinus* H.J.P.Winkl. in Bot. Jahrb. Syst. 57: 578. 1922, non *Boehmeria velutina* Decne. in Herb. Timor.: 163 (1834) ― **Lectotype (designated here)**: Northeast New Guinea [Papua New Guinea], Toliba, 300 m, Dec. 15 1908, *F.R.F. Schlechter 18966,* (K barcode K000741427; isolectotype BR barcode BR0000005623464*)). [Syntypes: NE New Guinea [Papua New Guinea], Shigaun, 400 m, Jun. 9 1896, *C.A.G. Lauterbach 2298* (B†, BO, K); Toliba, 300 m, Dec. 15 1908, *F.R.F. Schlechter 18966* (B†, isosyntypes BR barcode BR0000005623464, K barcode K000741427]).

***Boehmeria ledermannii*** (H.J.P.Winkl.) A.K.Monro & Wilmot-Dear **comb. nov**. *≡ Cypholophus ledermannii* H.J.P.Winkl. in Bot. Jahrb. Syst. 57: 581. 1922 ― Holotype: Indonesia, Sandaun, East Sepik] Sepik area, Felsspitzec station, 1400 to 1500 m, Jul. 30 1913, *C.L. Ledermann 12383* (B†; isotype: WRSL).

***Boehmeria melanocarpoides*** (H.J.P.Winkl.) A.K.Monro & Wilmot-Dear **comb. nov.** *≡ Cypholophus melanocarpoides* H.J.P.Winkl. in Nova Guinea 14: 129. 1924 ― Holotype: Indonesia, Papua], Mount Hellwig, 1800 m, Dec. 18 1912, *A.A.Pulle 755* (B†; isotype: WRSL).

***Boehmeria microphylla*** (Elmer) A.K.Monro & Wilmot-Dear **comb. nov.** *≡ Cypholophus microphyllus* Elmer in Leafl. Philipp. Bot. 3: 895. 1910 – Holoype: Philippines, Mindanao, Davao District, Mt. Apo, Todaya, [1800 m], Aug. 1909, *A.D.E. Elmer 11588* (A barcode A00039519; isotypes: BISH barcode BISH1005217, E barcode E00504430, G barcode G00354073, G barcode G00354081, HBG barcode HBG513214, K barcode K000741437!, MO barcode MO204580, NY barcode NY00284304, NY barcode NY00284305, US barcode US00090906).

***Boehmeria montana*** (Ridl.) A.K.Monro & Wilmot-Dear **comb. nov.** *≡ Cypholophus montanus* Ridl. in Trans. Linn. Soc. London, Bot. 9: 157. 1916 ―Holotype: Dutch New Guinea [Indonesia, Papua], Camp 10-11, Jan 27 1912, *C. Boden-Kloss s.n.* (BM barcode 000951675; isotype: K! barcode K000741433).

***Boehmeria nummularis*** (H.J.P.Winkl.) A.K.Monro & Wilmot-Dear **comb. nov.** *≡ Cypholophus nummularis* H.J.P.Winkl. in Bot. Jahrb. Syst. 57: 582. 1922 ― **Lectotype (designated here)**: New Guinea [Papua New Guinea], Madang Province, Kani Mountains, 1000 m, Dec. 5 1907, *F.R.F. Schlechter 16959* (B†; isolectotype WRSL) [Note: Two B collections were cited by Winkler, *Schlechter* 16959 and *Warburg* 20780. The Berlin duplicates were destroyed in WWII/ *Warburg* 20780 could not be traced, but a duplicate of *Schlechter* 16959, annotated as’original’ was located in WRSL and selected as lectotype.]

***Boehmeria pachycarpa*** (H.J.P.Winkl.) A.K.Monro & Wilmot-Dear **comb. nov.** *≡ Cypholophus pachycarpus* H.J.P.Winkl. in Bot. Jahrb. Syst. 57: 574. 1922 ― Syntypes: [Indonesia, Sandaun, East Sepik] Sepik, Schraderberg, 2070 m, May 29 1913, *C.L. Ledermann 11764* (B†); May 28 1913, *C.L. Ledermann 11695* (B†, WRSL).

***Boehmeria patens*** (H.J.P.Winkl.) A.K.Monro & Wilmot-Dear **comb. nov.** ≡ ― *Cypholophus patens* H.J.P.Winkl. in Bot. Jahrb. Syst. 57: 577. 1922. ― Holotype: Indonesia, Irian Jaya, South slope of the Hellwig Mountains, 1750 m, Dec. 20. 1912, *A.A.Pulle 786* (B†, WRSL).

***Boehmeria pulleana*** (H.J.P.Winkl.) A.K.Monro & Wilmot-Dear **comb. nov.** ≡ *Cypholophus pulleanus* H.J.P.Winkl. in Bot. Jahrb. Syst. 57: 584. 1922 ― Holotype: [Indonesia, Papua] South slope of the Hellwig Mountains, 1700-1800 m, Dec. 10. 1912, *A.A.Pulle 756* (B†, WRSL).

***Boehmeria radicans*** (H.J.P.Winkl.) A.K.Monro & Wilmot-Dear **comb. nov.** ≡ *Cypholophus radicans* H.J.P.Winkl. in Bot. Jahrb. Syst. 57: 576. 1922 ― Holotype: Indonesia, Papua, South slope of the Orange Mountains, Parameles bivouac, 750 m, Dec. 3. 1912, *A.A.Pulle 541* (B†, WRSL).

***Boehmeria reticulata*** (H.J.P.Winkl.) A.K.Monro & Wilmot-Dear **comb. nov.** ≡ *Cypholophus reticulatus* H.J.P.Winkl. in Bot. Jahrb. Syst. 57: 577. 1922 ― Holotype: [Indonesia, Sandaun, East Sepik] Sepik, Schraderberg, 2070 m, Jun. 2 1913, *C.L. Ledermann 11946* (B†, WRSL).

***Boehmeria rudis*** (Ridl.) A.K.Monro & Wilmot-Dear **comb. nov.** ≡ *Cypholophus rudis* Ridl. in Trans. Linn. Soc. London, Bot. 9: 157. 1916 ― **Lectotype (designated here)**: Dutch New Guinea [Indonesia, Papua], Camp 9-10, 1500-2000 m, Jan 26 1912, *C. Boden-Kloss s.n.* (K barcode 000741430; isolectotype: BM barcode 000951677).

***Boehmeria schlechtendahliana*** A.K.Monro & Wilmot-Dear **nom. nov**. Replaced name: non *Boehmeria rotundifolia* D.Don. in Prodr. Fl. Nepal.: 60. 1825 ≡ *Cypholophus rotundifolius* H.J.P.Winkl. Bot. Jahrb. Syst. 57: 571 (1922), ― Holotype: New Guinea [Papua New Guinea], Finisterre Mountains, 1300 m, Jan. 10 1909, *F.R.F. Schlechter 19051* (B†; isotypes: K barcode 000741432, BR barcode BR0000005623792*, WRSL).

***Boehmeria trapula*** (H.J.P.Winkl.) A.K.Monro & Wilmot-Dear **comb. nov.** ≡ *Cypholophus trapula* H.J.P.Winkl. in Bot. Jahrb. Syst. 57: 579. 1922 ― Holotype: [Papua New Guinea] Northeastern New Guinea, Jul. 14 1907, *F.R.F. Schlechter 16263* (B†; isotypes: A barcode 00035011,BISH barcode 1005218, E barcode 00301048, G barcode 00354086, K barcode 000741428, K barcode 000741429, S (S07-9824), SING barcode 0062080, SING barcode 0062081).

***Boehmeria treubii*** (H.J.P.Winkl.) A.K.Monro & Wilmot-Dear **comb. nov.** ≡ *Cypholophus treubii* H.J.P.Winkl. in Bot. Jahrb. Syst. 57: 583. 1922 ― Holotype: Southwest New Guinea [Papua New Guinea] North slope of the Treub Mountains, 2400 m, Feb. 17 1913, *A.A.Pulle 1087* (B†).

***Boehmeria vaccinioides*** (H.J.P.Winkl.) A.K.Monro & Wilmot-Dear **comb. nov.** ≡ *Cypholophus vaccinioides* H.J.P.Winkl. in Bot. Jahrb. Syst. 57: 583. 1922 ― Holotype: [Indonesia, Papua], between Mount Hellwig and Alkmaar camp, *L.S.A.M. von Römer 724* (B†); summit of Mount Hellwig, 2600 m, 1700-1800 m, Jan. 2. 1913, *A.A.Pulle 877* (B†). ― **Lectotype (selected here)**:

***Boehmeria vriesiana*** A.K.Monro & Wilmot-Dear **nom. nov.** Replaced name: *Cypholophus ellipticus* Wedd. in Prodr. [A.P.de Candolle] 16(1): 235(12) (1869), *Cypholophus ellipticus* Wedd. in Prodr. 16(1): 235(12) (1869), non *Boehmeria elliptica* Wedd. in Ann. Sci. Nat., Bot., sér. 4, 1: 200 (1854). Holotype: ― [Indonesia, Sulawesi, Celebes] Java [see note below], *G.H. de Vriese 23* (P barcode 06873943).

There are four specimens collected by *de Vriese* at L, one of which is numbered’23’,’Celebes’ and’type duplicate’ (L barcode 2070508), all of which were collected in the Celebes Islands, Sulawesi. The presumed holotype in P (P06873943) is labelled *De Vriese*’23’ and’Java’, P. The L duplicate has the more complete label and indicates the collectors as *’G.H. de Vriese & J.E. Teijsmann* 23’, the P specimen label indicates’23’, *’de Vriese’* and’Java’. Given that Weddell described the species in P, the latter is the holotype. The locality given is, however, likely incorrect, given that all of the De Vries collections of this taxon in L include the Celebes as the locality.

***Boehmeria warburgiana*** (Lauterb.) A.K.Monro & Wilmot-Dear **comb. nov.** ≡ *Cypholophus warburgianus* Lauterb. in K.M.Schumann & C.A.G.Lauterbach in Fl. Schutzgeb. Südsee, Nachtr.: 255. 1905 ― Holotype: Kaiser-Wilhelmsland [Papua New Guinea], Bismark Mountains, 1800 m, Jan. 1902, *F.R.F. Schlechter 14007* (B†).

**Notes** Nearctic, Neotropics, Palearctic, Afrotropics, Indomalaya, Australasia; ca 85 spp.

Our increased taxon and genome sampling confirmed the findings of previous studies (Wu et al., 2013, 2018) that Boehmeria is both poly- and para- phyletic (Fig. 3, 4.1). Translating our results into a phylogenetic classification required us to choose one of several scenarios: 1) treat all species of *Boehmeria,* together with those genera with which they form a monophyletic clade (see Fig.s 3, 4.1), as a single genus. This would require treating *Archiboehmeria, Astrothalamus, Chamabainia, Cypholophus, Debregeasisa*, *Gonostegia*, *Neraudia, Phenax, Pipturus,* and *Pouzolzia* as a single genus and establishing a grouping with no diagnostic morphological characters; or 2) retain the morphologically distinct genera above but place *Cypholophus* in synonymy with the core of *Boehmeria* and propose a new genus to account for *Boehmeria ourantha* and *B. nivea*, or 3) maintain *Cypholophus* as a distinct genus and propose two new genera, one to accommodate *Boehmeria ourantha* and *B. nivea* (Boehmeria II,Fig. 3, 4.1), and another to accommodate *B. depauperata* and *B. pseudotricuspis.* We have chosen scenario 2 as we believe that it represents the best compromise between morphologically diagnosable and monophyletic groupings.

*Boehmeria* needs much greater taxon sampling, and its morphology requires a critical re-evaluation. Although we sampled ca 25% of the described species and much of the geographical distribution, there is a large number of subspecies and varieties in the list of accepted species in Wilmot-Dear & Friis (Wilmot-Dear & Friis, 1996, 2013) that warrant further investigation. Given the strong geographical signal within the family and genus, and the broad distribution of some of these taxa, we would not be surprised if many of the varieties and subspecies should be raised to the rank of species following a detailed taxon sample including several accessions of each subspecies and variety and close morphological evaluation.

33. ***Pouzolziella*** A.K.Monro, Friis & Wilmot-Dear, **gen. nov**. Type: *Pouzolziella australis* (Endl.) A.K.Monro, Friis & Wilmot-Dear, **comb. nov**. ≡ *Pouzolzia australis* (Endl.) Friis & Wilmot-Dear in Nordic J. Bot. 24: 15. 2006. ≡ *Ramium australe* Kuntze in Revis. Gen. Pl. 2: 633. 1891. ≡ *Boehmeria australis* Endl. in Prodr. Fl. Norfolk.: 38. 1833 – Holotype: “Crescit in insula Norfolk, mense Octobri florens”, *F.L. Bauer s.n.* (W destroyed; isotype: K). Fig. 13.

**Fig. 13.**
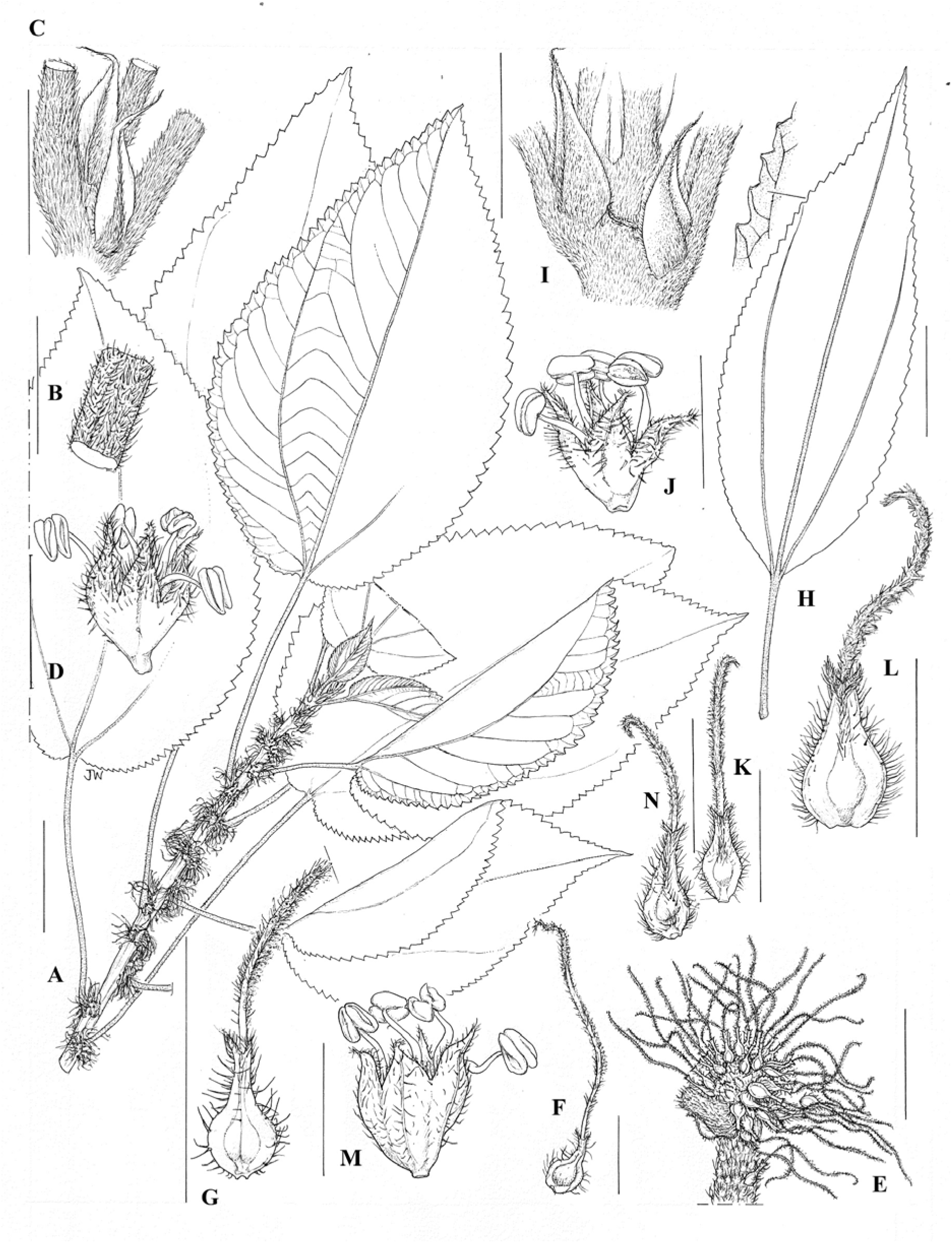
*Pouzolziella* Plate. *Pouzolziella australis.* Norfolk Islands population: **A** habit with female inflorescence based on *Cunningham s.n.* (scale bar = 3 cm); **B** stem pubescence based on *Cunningham s.n.* (scale bar = 5 mm); **C** stipule based on *Cunningham s.n.* (scale bar = 5 mm); **D** male flower based on *Green* 1404 (scale bar = 2 mm); **E** female flower cluster based on *Cunningham s.n.* (scale bar = 5 mm); **F** female flower based on *Cunningham s.n.* (scale bar = 2 mm); **G** fruit based on *Cunningham s.n.* (scale bar = 2 mm). Kermadec Islands population: **H** leaf outline and margin based on XXX (scale bar = 3 cm); **I** stipule based on XXX (scale bar = 5 mm); **J** male flower based on XXX (scale bar = 2 mm); **K** female flower based on XXX (scale bar = 2 mm); **L** fruit based on XXX (scale bar = 2 mm). Lord Howe Island population: **M** male flower based on *Von Mueller s.n.* (scale bar = 2 mm). Sunday Islands population: **N** female flower based on *Oliver s.n. ‘10/10/1908’* (scale bar = 2 mm). Illustration by Juliet Beentje.

**Fig. 14.**
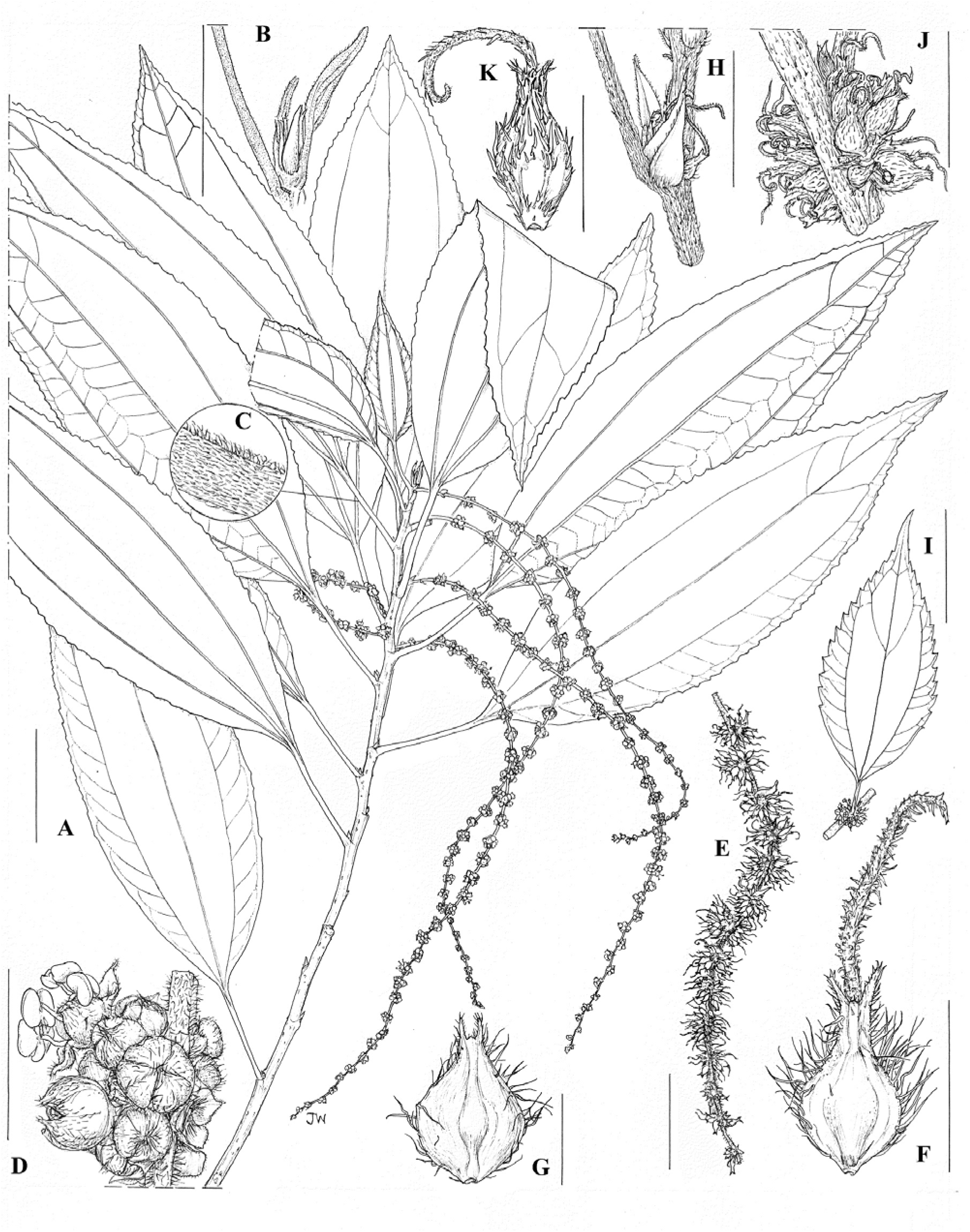
*Margarocarpus* Plate. *Margarocarpus rugulosus*: **A.** habit with male inflorescences based on *Williams & Stainton* 8289 (scale bar = 3 cm); **B** stipules based on *Williams & Stainton* 8289 (scale bar = 1 cm); **C** uncinate hairs on petiole based on *Williams & Stainton* 8289 (); **D** male flower clusters based on *Williams & Stainton* 8289 (scale bar = 3 mm); **E** female inflorescences based on *Wall* 4597 (scale bar = 1 cm); **F** female flowers based on *Wall 4597* (scale bar = 1 mm); **G** achene based on *Wall 4597* (scale bar = 1 mm). *Margarocarpus sanguineus*: **H** stipule based on *Hiep* 3867 (scale bar = 3 mm); **I** leaf outline based on *Hiep* 3867 (scale bar = 3 cm); **J** female inflorescence based on *Hiep* 3867 (scale bar = 3 mm); **K** female flower based on *Hiep* 3867 (scale bar = 1 mm). Illustration by Juliet Beentje.

Diagnosis: most similar to *Pouzolzia* Gaudich. but lacking uncinate hairs and possessing the combination of serrate leaves with dense white and shiny tomentose pubescence below and fruiting perianths that are distinctly winged.

Densely branched shrub or small tree with spreading crown; young stems with abundant pubescence; stipules soon deciduous. Leaves alternate, slightly or moderately dimorphic, broadly or narrowly ovate to rhombic-ovate; margin serrate; chartaceous; basal veins extending just into upper third of lamina, upper lateral veins 3-6 but often scarcely distinguishable from coarser tertiary venation, lowermost arising near middle, veins slender but at least basal veins visible above, lateral veins and coarser reticulation visible beneath; upper surface glabrous or with scattered hairs, scabrid with extremely dense prominent cystoliths, lower surface with dense shining white or greyish tomentum, this absent from main veins and often from even fine reticulation. Petiole densely pubescent. Inflorescences unisexual or bisexual axillary clusters, bracteate, with few to < 50 flowers, clusters sometimes so large as to be almost continuous along stem. Male flowers 4-merous, sessile, with sparse adpressed or dense fine spreading hairs. Female flowers with perianth not accrescent, fused to form a tube with a pronounced beak, pubescent, compressed-ovoid, narrowly winged; style persistent. Fruiting perianth as in flower but more prominently winged and with 1-2 facial ribs. Seed light brown, shiny, easily detachable from perianth at maturity.

**Notes** Australasia; 1 sp. *Pouzolziella australis,* as delimited by Wilmot-Dear & Friis (Wilmot-Dear & Friis, 1996) had been considered as three distinct island endemics by previous authors (*P. australis* ℒ Norfolk Island; *Boehmeria calophleba* C.Moore & F.Muell. ℒ Lord Howe Island; *B. dealbata* Cheesem. ℒ Kermadec Island). Increased taxon sampling and the resulting morphological and molecular observations may lead to these taxa again being considered as distinct species.

*Pouzolziella* was recovered as relatively distantly related to *Pouzolzia* within the Boehmerieae and sister to the Indomalayan and Australasian genus *Pipturus* (Pouzolzia II Fig. 3, Fig. 4.1) with which it shares a woody habit, densely tomentose lower leaf surfaces and absence of uncinate hairs, alternate serrate leaves, a long stigma and glossy achene surface. Wilmot-Dear & Friis (Wilmot-Dear & Friis, 2004) had already noted that *P. australis* was an atypical member of *Pouzolzia* based on the combination of serrate leaf margins, leaves with densely tomentose lower leaf surfaces and relatively long persistent stigmas and winged fruit. We also observed that it lacks the uncinate (hooked) hairs characteristic of *Pouzolzia.* We considered moving *P. australis* to *Pipturus* but despite the similarities listed above, it can be distinguished by the bracteate inflorescences (*versus* ebracteate), persistent stigma in fruit (*versus* soon caducous) and non-articulated ((*versus* articulated) stigmas that are dorsally glabrous (*versus* pubescent for the full circumference) and winged fruit (*versus* fleshy fruit) (see Table 5 below). For the above reasons we proposed this new genus.

**Table 5.** Diagnostic comparison of *Margarocarpus*, *Neraudia, Pouzolziella* and *Piptrurus.*

| **Character state** | *Margarocarpus* | *Neraudia* | *Pouzolzia* | *Pouzolziella* | *Pipturus* |
| --- | --- | --- | --- | --- | --- |
| **Uncinate hairs** | present | present | present | absent | absent |
| **Stigma articulation** | articulated | articulated | Not articulated | not articulated | articulated |
| **Distribution of papillae on stigma** | asymmetrical, portion of circumference glabrous | radially symmetrical, whole circumference pubescent | asymmetrical, portion of circumference glabrous | asymmetrical, portion of circumference glabrous | whole circumference pubescent |
| **Stigma persistence in fruit** | Not persistent | persistent | Persistent | Persistent | Not persistent |
| **Female inflorescence** | compact cymose | cymose / fasciculate | compact cymose | compact cymose | fused capitate |
| **Fruit winged?** | not winged | winged | Winged or not | winged | not winged |
| **Achene colour** | Cream to white | brown | Pale brown, white | Pale brown | Brown-grey |
| **Geography** | Indomalaya | Hawaii | Afrotropics, Australasia, Indomalaya, Nearctic, Neotropics, Pacific Islands, Palearctic | Australasia | Australasia, Indomalaya, Pacific Islands |

34. ***Pipturus*** Pipturus Wedd. in Ann. Sci. Nat. Bot., sér. 4. 1: 196. 1854 – Type: *Pipturus velutinus* (Decne.) Wedd. in Ann. Sci. Nat. Bot., sér. 4. 1: 196. 1854 – Lectotype designated by Skottsberg in Acta Horti Gothob. 8: 117. 1933): *Boehmeria velutina* Decne. in Nouv. Ann. Mus. Hist. Nat., sér. 3. 3: 491. 1834.

=*Nothocnide* Blume in Mus. Bot. 2: t. 14 – Type: *Nothocnide repanda* Blume in Mus. Bot. 2: t. 14. 1856 ≡ *Urtica repanda* Blume in Bijdr. Fl. Ned. Ind. 10: 501. 1826.

=*Pseudopipturus* Skottsb. in Acta Horti Gothob. 8: 117. 1933. ≡ *Urtica* repanda Blume in Bijdr. Fl. Ned. Ind. 10: 501. 7 Dec 1825-24 Jan 1826. Type: *Pseudopipturus repandus* (Blume) Skottsb. in Acta Horti Gothob. 8: 117. 1933.

**Notes** Indomalaya, Australasia, Pacific Islands; 34 spp. *Pipturus* was recovered as polyphyletic with respect to *Nothocnide.* The taxa differ in habit, *Pipturus* comprises trees and shrubs, *Nothocnide* vines. They are otherwise identical with respect to the morphological characters sampled here (see Table2) and share the same geographical distribution (See Fig. S2).

The perianth of *Pipturus* appears completely fused in flower, the caducous stigma leaving a characteristic hole at the flower apex. In fruit, however, once dried it can be observed that the perianth splits into two, exposing the achene, suggesting that the perianth may not be fully fused. This would benefit from further research.

One of two accessions of *Pouzolzia sanguinea* var. *aspera* (Li) Friis & Wilmot-Dear, extracted from material from Taiwan, was recovered sister to *Pipturus aspera* Wedd., also sampled from Taiwan. Both originated from samples at KUN generated by Wu & al. (Wu et al., 2013). It would be worthwhile revising the identification of this material.

35. ***Neraudia*** Gaudich. In Voy. Uranie: t. 117. 1826. Type: *Neraudia melastomifolia* Gaudich. In Voy. Uranie: t. 117. 1826. *Praetoria* Baill. In Étude Euphorb. 470, t. 11. 1858, nom. Nud.

**Notes** Endemic to Hawaii; 5 spp.

36. ***Margarocarpus*** Wedd. In Ann. Sci. Nat., Bot., sér. 4, 1: 203. 1854 – **Lectotype (designated here)**: *Margarocarpus 56xasperates* Wedd. In Ann. Sci. Nat., Bot., sér. 4, 1: 204. 1854. Fig.14.

***Margarocarpus rugulosus*** (Wedd.) A.K.Monro & Wilmot-Dear **comb. Nov**. ≡ *Pouzolzia rugulosa* (Wedd.) Acharya & Kravtsova in Edinburgh J. Bot. 66: 60. 2009. ≡ *Boehmeria rugulosa* Wedd. In Arch. Mus. Hist. Nat. 9: 378. 1856 – Lectotype (designated by Wilmot-Dear & al. 2009): Nepal, *Wallich 4597* (K-W barcode K001039434; isolectotypes: BR!, G!,K!, M!).

***Margarocarpus sanguineus*** (Blume) A.K.Monro & Wilmot-Dear **comb. Nov**. ≡ *Pouzolzia sanguinea* (Blume) Merr. In J. Straits Branch Roy. Asiat. Soc. Special Number: 233. 1921 ≡ *Boehmeria sanguinea* (Blume) Hassk. In Cat. Hort. Bot. Bogor. Alt.: 79. 1844 ≡ *Urtica sanguinea* Blume in Bijdr. Fl. Ned. Ind.: 501. 1826 – Holotype?: Indonesia, Java, Salak, *Blume 1550* [L barcode L0397592].

***Margarocarpus calophyllus*** (W.T.Wang & C.J.Chen) A.K.Monro & Wilmot-Dear **comb. Nov**. ≡ *Pouzolzia calophylla* W.T.Wang & C.J.Chen in Acta Phytotax. Sin. 17(1): 108. 1979. Holotype: China, Xizang, Chayu, 1900ℒ2000 m, 8 or 9 July 1973, *Qinghai-Xizang Expedition 647* (PE).

**Notes** Indomalaya; 2 spp. Pfeiffer (1873) selected *M. vimineus* as lectotype of *Margarocarpus,* presumably on mechanical grounds, as it is the first of 11 species assigned to the genus at the time of its description (Weddell, 1854). *Margarocarpus vimeus* is however an invalid name as the basionym cited by Weddell is illegitimate being both a *nomen nudum* and a later name for *Pouzolzia borbonica* Wight (≡ *P. sanguinea* var. *sanguinea* (Blume) Merr.). For this reason, we have selected *Margarocarpus aspera* Wedd. as lectotype, the next available name in the sequence.

Wu & al. (Z. Y. Wu et al., 2018, Supplementary Fig.1) (2018, Supplementary Figs1) and we recovered *Pouzolzia sanguinea* and *P. rugulosa* sister to a clade comprising *Neraudia, Pipturus* and *Pouzolziella,* rather than with the other species of *Pouzolzia* (Pouzolzia III Fig. 3, 4.1) sampled which were recovered in a clade comprising *Gonostegia* and *Pouzolzia* (Fig. 3, 4.1). *Pouzolzia sanguinea* and *<u>P. rugulosa</u>* are morphologically distinct from *Neraudia, Pipturus* and *Pouzolziella* as shown in Table 5 (below).

[Table 5. Diagnostic comparison of *Margarocarpus, Neraudia, Pouzolziella* and *Pipturus.*]

Wilmot-Dear & al. (Wilmot-Dear & Friis, 2004) identified *P. sanguinea* and *P. rugulosa* as forming part of a distinct grouping within *Pouzolzia* that also included *P. australis, P. niveotomentosa, P. sanguinea, P. rubricaulis, P. variifolia, P. fadenii* and *P. parasitica,* all of which share toothed leaf margins. We did not, however, recover this group of toothed species as a monophyletic, *Pouzolzia fadenii* and *P. parasitica* being recovered within a clade comprising entire-leaved species of *Pouzolzia,* suggesting that serrate leaf margins are homoplastic across these genera. In light of the above molecular and morphological observations, we decided to resurrect *Margarocarpus* Wedd.

37. ***Pouzolzia*** Gaudich. in Voy. Uranie: 503. 1830 – Type: *Pouzolzia laevigata* Gaudich. in Voy. Uranie: 503. 1830 – **Lectotype (designated here)**: Mauritius, *Anon. s.n.,* (P barcode P-LAM 05238550; isolectotype: P barcode P06818056).

*=Rousselia* Gaudich. in Voy. Uranie, Bot. pt. 12: 503. 1830 – **Lectoype (designated here)**: *Rousselia humilis* (Sw.) Urb. in Symb. Antill. (Urban) 4: 205. 1905 ≡ *Urtica humilis* Sw. in Kongl. Vetensk. Acad. Nya Handl., ser. 2, 6: 34. 1785. Type: ‘Habitat in Jamaicae petrosis vulgatissima.’ ≡ *Urtica lappulacea* Sw. in Kongl. Vetensk. Acad. Nya Handl., ser. 2, 8: 69. 1787, nom. illeg. & superfl. ≡ *Lithocnide lappulacea* (Sw.) Raf. in Fl. Tellur. 3: 48 (1836)[Nov.-Dec. 1837], comb. illeg. ≡ *R. lappulacea* (Sw.) B. D. Jackson in Index Kewensis 2(2 = fasc. 4): 744. 1895.

*Lithocnide* Raf. in Fl. Tellur. 3: 48. 1836[Nov.-Dec. 1837], nom. superfl.

*=Hemistylus* Benth. in Pl. Hartw. [Bentham]: 123. [late Dec. 1843-early Jan. 1844] – Holotype: *Hemistylus boehmerioides* Benth. in Pl. Hartw. [Bentham]: 123. [late Dec. 1843-early Jan. 1844] – **Lectotype (selected here)**: Ecuador, near Guyaquil, April, *Hartweg 696* (K barcode K000575855; isotype: K barcode K000575854).

*=Leucococcus* Liebm. in Kongel. Danske Vidensk.-Selsk. Skr. ser. 5. 2: 311. 1851 – Type: *Leucococcus occidentalis* Liebm. in Kongel. Danske Vidensk.-Selsk. Skr. ser. 5. 2: 311. 1851 – **Lectotype (selected here)**: Nicaragua, 1845, *A.S. Ørsted 14258* (C; isolectotype, MO).

=*Stachyocnide* Blume in Mus. Bot. 2: 227. 1857 – Type: *Stachyocnide luzonica* Blume in Mus. Bot. 2: 227. 1857.

=*Praetoria* Baill. in Étude Euphorb. 470, t. 11. 1858, nom. nud.

=*Elkania* Schlecht. ex Wedd. in Prodr. [A. P. de Candolle] 16(1): 227. 1869 – Type: *Elkania multinervis* Schlecht. ex Wedd. in Prodr. [A. P. de Candolle] 16(1): 227. 1869.

=*Goethartia* Herzog, Meded. Rijks-Herb. 27: 77. 1915 – Type: *Goethartia edentata* (Kuntze) Herzog, Meded. Rijks-Herb. 27: 77. 1915. ≡ *Ramium edentatum* Kuntze, Revis. Gen. Pl. 3: 294. 1898.

*=Neodistemon* Babu & A.N.Henry in Taxon 19: 651. 1970 – Type: *Neodistemon indicus* (Wedd.) Babu & A.N.Henry in Taxon 19: 651. 1970.

***Pouzolzia boehmerioides*** (Benth.) A.K.Monro **comb. nov.** ≡ *Hemistylus boehmerioides* Benth. in Pl. Hartw.: 123. 1843.

***Pouzolzia macrostachya*** (Wedd.) A.K.Monro **comb. nov.** ≡ *Hemistylus* macrostachya in Ann. Sci. Nat., Bot., sér. 4, 1: 208. 1854.

***Pouzolzia zollingeriana*** A.K.Monro **nom. nov.** ≡ *Neodistemon indicus* (Wedd.) Babu & A.N.Henry in Taxon 19: 651 (1970) ≡ *Distemon indicus* Wedd. in Ann. Sci. Nat., Bot., sér. 4, 1: 208. 1854. Monogr. Fam. Urtic.: 551. 1856. **Lectotype (designated here)**: Ins. Javae [Indonesia], *Zollinger 2785* (P barcode 06783132).

A replacement name was selected as the combination *Pouzolzia indica* has previously been published by Wight (1853: 10).

***Pouzolzia humilis*** (Sw.) A.K.Monro**, comb. nov.** ≡ *Rousselia humilis* (Sw.) Urb. in Symb. Antill. (Urban) 4: 205. 1905 ≡ *Urtica humilis* Sw. in Kongl. Vetensk. Acad. Nya Handl., ser. 2, 6: 34. 1785.

**Notes** Neotropics, Afrotropics, Indomalaya, (Australasia); ca 60 spp. We recovered *Pouzolzia* as poly- and para-phyletic (see Discussion). Resolving a classification that reflected both phylogenetic relationships and morphological characters was very challenging. Our solution is to separate out *Pouzolia australis* (= *Pouzolziella* gen. nov.), *P. sanguinea* & *P. rugulosa* (= *Margarocarpus*) and *P. rubricaulis* (= Leptocnide), and to place *Neodistemon, Hyrtanandra, Hemistylus* and *Rousselia* within Pouzolzia. The justification for the latter is that, despite their morphological diagnosability, retaining *Neodistemon, Hyrtanandra, Hemistylus* and *Rousselia* would require the delimitation of several new genera for which morphological diagnoses are currently lacking, rendering the recognition of taxa even more complicated than it already is.

We decided not to place *Gonostegia* into synonymy with *Pouzolzia* as proposed by Wilmot-Dear & Friis (Wilmot-Dear & Friis, 2004). This suggestion had also been discussed, and ultimately rejected by Weddell (Weddell, 1856). *Gonostegia* (cited by Weddell as *Memorialis*) is morphologically very distinct (see below) and phylogenetically, together with *Pouzolzia rubricaulis* (= *Leptocnide,* below), borne on a relatively long branch that is estimated to have diverged from the remainder of the taxa included in Pouzolzia (Pouzolzia I, Fig.s 3, 4.1) >30 MYA. Including *Gonostegia* and *Leptocnide* in *Pouzolzia* would result in a grouping lacking shared diagnostic morphological characters and on this basis we decided to exclude these taxa from *Pouzolzia.*

The combination, *Rousselia lappulacea* (Sw.) Gaudich., was never made by Gaudichaud, – he merely quoted ‘*Urtica lappulacea,* Swartz (Herb. Mérat) and Jackson made the combination retrospectively in Index Kewensis (Ref.).

*Pouzolzia* needs much greater taxon sampling and a critical re-evaluation of its morphology. Although we sampled ca 25% of the described species and much of the geographical distribution, there are a large number of subspecies and varieties concealed in the list of accepted species in Wilmot-Dear & al. (Wilmot-Dear & Friis, 2004). Given the strong geographical signal within the family and genus, and the broad distribution of some of these taxa we would not be surprised if many of the varieties and subspecies were to be raised to the rank of species following a detailed taxon sample including several accessions of each subspecies and variety.

The type for *Pouzolzia laevigata* was incorrectly cited in Wilmot-Dear & Friis (Wilmot-Dear & Friis, 2004). Marais (Marais, 1985) cites’Mauritius, *Anon.* s.n. (P-LA)’ as holotype but did not include any identifying information for the specimen (e.g. folder number or loan number).

38. **Leptocnide** Blume in Mus. Bot. Lugd.-Bat. 2: 193. 1857 ≡ Pouzolzia rubricaulis (Blume) Wedd. in A.P.de Candolle, Prodr. 16(1): 229. 1869. – **Lectotype (designated here)**: *Leptocnide rubricaulis* Blume in Mus. Bot. Lugd.-Bat. 2: 194. 1857. Fig. 15.

**Fig. 15.**
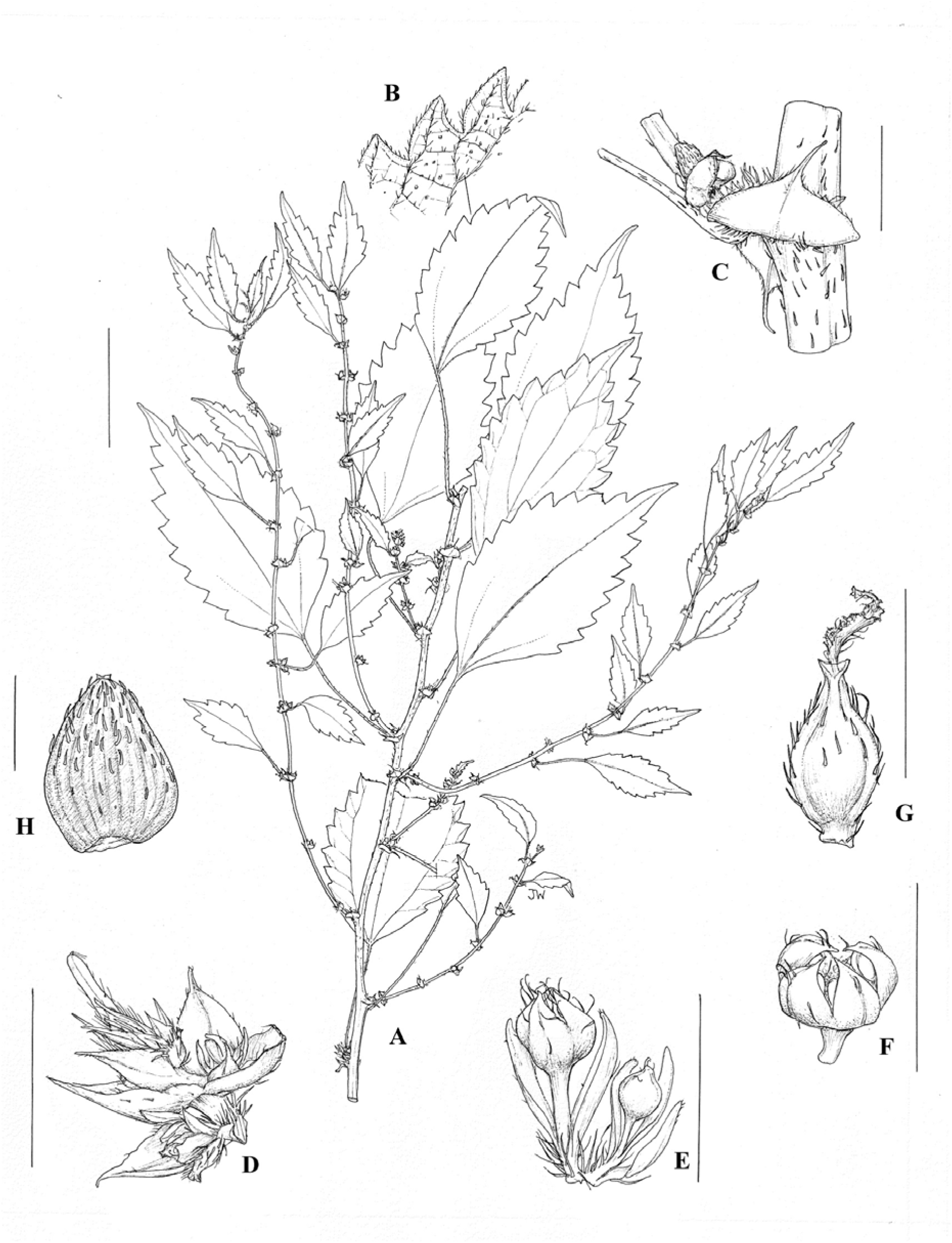
*Leptocnide* Plate. *Leptocnide rubricaulis*: **A**. habit based on *Koorders* 27600B (scale bar = 3 cm); **B** detail of leaf lower surface based on *Koorders* 27600B (scale bar = 3 cm); **C** bract enclosing inflorescence based on *Koorders* 27600B (scale bar = 3 mm); **D** female inflorescence detail based on *Koorders* 27600B (scale bar = 3 mm); **E** male flowers in bud based on *Koorders* 27600B (scale bar = 1 mm);**F** male flowers based on *Koorders* 27600B (scale bar = 1 mm); **G** female flower based on *Koorders* 27600B (scale bar = 1 mm); **H** achene based on *Koorders* 27600B (scale bar = 1 mm). Illustration by Juliet Beentje.

**Notes** Indomalaya, Australasia; 1 sp.

*Leptocnide rubricaulis* Blume was included in *Pouzolzia* by Weddell and later Wilmot-Dear & al. (2004). The latter considered this toothed-leaved species to form part of an anomalous group of *Pouzolzia* which also included *P. sanguinea, P. rugulosa* (≡ *Margarocarpus* above) *P. australis* (≡ *Pouzolziella* above), *P. niveotomentosa*, *P. variifolia, P. fadenii* and *P. parasitica.* We recovered this taxon as sister to *Gonostegia* on a long branch within the Boehmerieae (Pouzolzia IV, Fig.s 3, 4.1). As above, we feel there is strong support for not incorporating *Gonostegia* into *Pouzolzia* and so by association, neither for *L. rubricaulis.* Wilmot-Dear & Friis (2004) noted that *L. rubricaulis* (as *Pouzolzia rubricaulis*) is unique within *Pouzolzia* in its very broad, caudate and thick stipules. *L. rubricaulis* is morphologically distinct from *Gonostegia* with respect to leaf arrangement, margin, stipule, staminate and pistillate flowers as summarised in Table 6 (below). Combining *Gonostegia* and *Leptocnide* would therefore result in a grouping lacking shared diagnostic morphological characters. We therefore decided to treat the taxa as distinct at generic rank and to resurrect *Leptocnide.*

**Table 6.** Diagnostic comparison of *Pouzolzia, Leptocnide* and *Gonostegia*.

| Character state | *Pouzolzia* | *Leptocnide* | *Gonostegia* |
| --- | --- | --- | --- |
| **Stipule** | Longer than broad, lacking an abrupt apex | Broader than long, with abrupt apex | Longer than broad, lacking an abrupt apex |
| **Leaf arrangement** | alternate | alternate | opposite |
| **Leaf margin** | Entire or toothed | toothed | entire |
| **Leaf venation** | Lateral 'primary' veins extending $\frac{1}{2}$ to $\frac{3}{4}$ the lamina length | Lateral 'primary' veins extending ca $\frac{2}{3}$ the lamina length | Lateral 'primary' veins extending the full length of the lamina, including the apex |
| **Staminate flowers in bud** | Sepals fused or not, where fused, for $<1/2$ their length, the free portions not subtended by a crest | Sepals fused or not, where fused, for $<1/2$ their length, the free portions not subtended by a crest | Sepals fused for $>1/2$ their length, free portions subtended by a crest forming a sub-apical circumferential ring |

[Table 6. Diagnostic comparison of *Pouzolzia, Leptocnide* and *Gonostegia.*]

39. ***Gonostegia*** Turcz. in Bull. Soc. Imp. Naturalistes Moscou 19: 509. 1846. Type: *Gonostegia oppositifolia* Turcz. in Bull. Soc. Imp. Naturalistes Moscou 19: 510. 1846.

*Hyrtanandra* Miq. in Pl. Jungh.: 25. 1851 – **Lectotype (designated here)**: *Hyrtanandra javanica* Miq. in Pl. Jungh.: 25. 1851.

*Memorialis* Wedd. in Arch. Mus. Hist. Nat. 9: 415. 1857 – **Lectotype (selected here)**: *M. ciliaris* Wedd. in Arch. Mus. Hist. Nat. 9: 417. 1857 – **Lectotype (designated here)**: Anon s.n. 4598A [ex Herb. Roxburgh], (K-W).

***Gonostegia cymosa*** (Wight) A.K. Monro, Friis & Wilmot-Dear, **comb. nov**. ≡ Pouzolzia cymose Wight in Icon. Pl. Ind. Orient. 6: 42. 1853 – Holotype: India, Tamil Nadu, Nilgiris, *Wight* s.n. (K barcode 000741370; isotypes: K barcode K000741371, L barcode L0484261, MEL barcode MEL2421129, NY barcode NY00063608).

**Notes** Palearctic, Indomalaya; 3 spp. Miquel (Junghuhn & Miquel, 1853) described two species of *Hyrtanandra* at the time of describing the genus and we have elected *H. javanica* on the basis that the description is the more comprehensive of the two. There has been considerable debate as to whether *Gonostegia* should be included within *Pouzolzia* or not (Weddell, 1856; Wilmot-Dear & Friis, 2004). Based on its morphological (see Table 6 above) and molecular distinctiveness we have followed Weddell’s proposal to maintain *Gonostegia* as distinct (cited by Weddell as *Memorialis*), not least because combining the genera would result in a grouping for which there is no morphological diagnosis. *Gonostegia pentandra* (Roxb.) Bennett & Brown is geographically widespread and morphologically variable (Wilmot-Dear & Friis, 2004) and it may be that future research determines it to comprise several species.

40. ***Chamabainia*** Wight in Icon. Pl. Ind. Orient. [Wight] vi. 11. t. 1981. 1853. Type: *Chamabainia cuspidata* Wight in Pl. Ind. Orient. [Wight] vi. 11. t. 19.

**Notes** Indomalaya; 1 sp.

41. ***Muimar*** A.K.Monro, Friis & Wilmot-Dear, **gen. nov.** ϑType Muimar nivea (L.) A.K.Monro, Friis & Wilmot-Dear. Fig. 16.

**Fig. 16.**
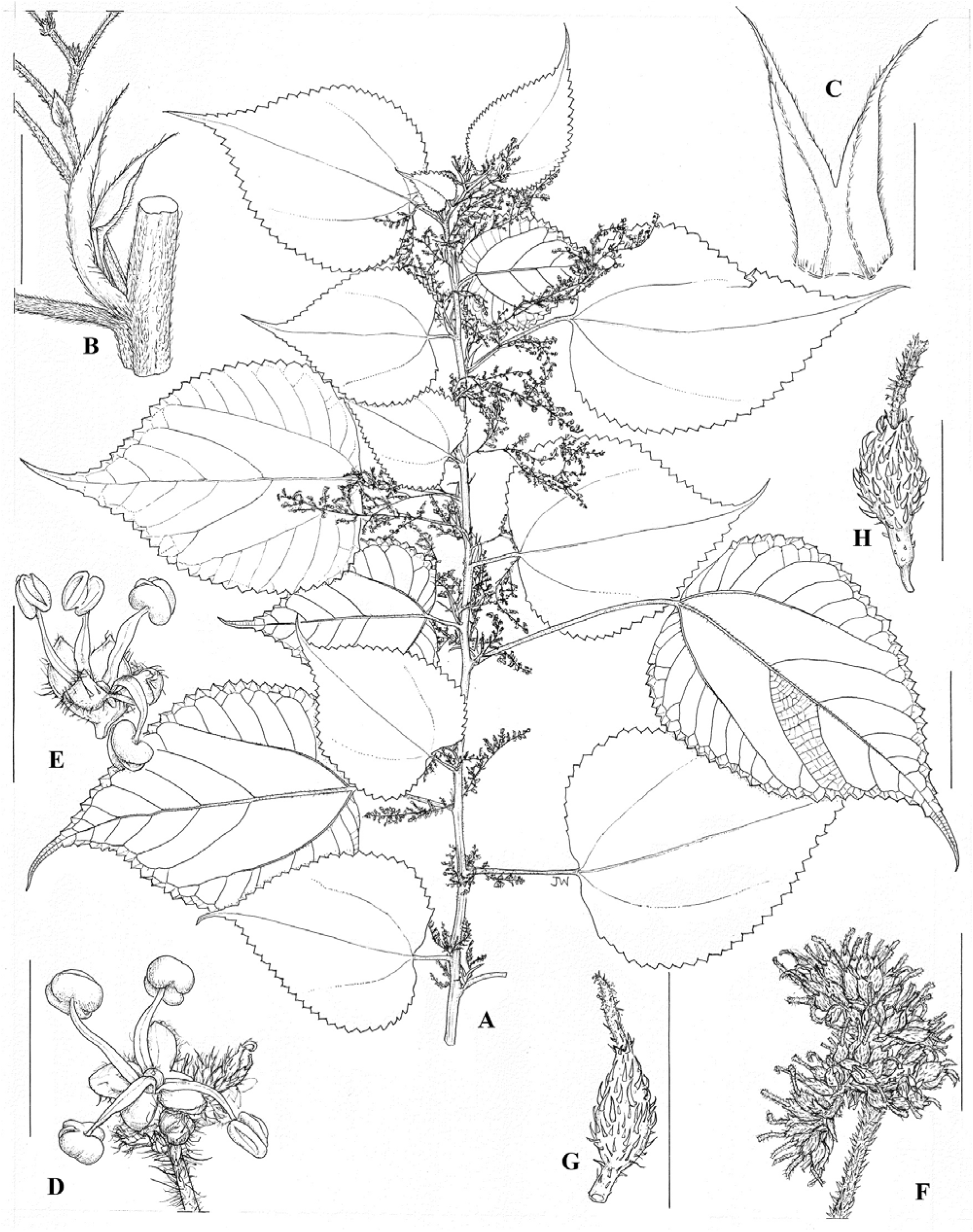
*Muimar* Plate. *Muimar nivea*: **A** habit based on *Guick* 10 (scale bar = 3 cm); **B** stipule position based on *Guick* 10 (scale bar = 5 mm); **C** stipule based on *Guick* 10 (scale bar = 5 mm); **D** male flower positioned within cluster of female flowers based on *Anon* 193 (scale bar = 3 mm); **E** male flower based on *Anon* 193 (scale bar = 3 mm); **F** female flower cluster based on *Hu* 8113 (scale bar = 3 mm); **G** female flower based on *Hu* 8113 (scale bar = 1 mm); **H** fruit based on *Hu* 8113 (scale bar = 1 mm).

Differs from *Boehmeria* by the combination of intrapetiolar stipules and shiny pubescence on the lower leaf surface.

Shrub or subshrub, monoecious or dioecious. Stems pubescent. Stipules interpetiolar, prominent, a pair at each leaf base, partially fused or free. Leaves alternate or opposite; margin serrate; basal pair of lateral veins more prominent than subsequent pairs; adaxial surface sparsely pubescent or glabrous, abaxial surface densely pubescent, the hairs shiny, cystoliths punctiform. Inflorescences paniculate or few-branched raceme-like cymes, unisexual or bisexual, bracts ≤3 mm, visible. Flowers borne in sessile unisexual or bisexual clusters, bracteoles ≤1 m inconspicuous, entirely male clusters small, 2–4 mm diam, bearing 5–15 flowers, entirely female clusters (2–)5–6(–8) mm diam, bearing 10–100 flowers. Male flowers 4-merous, sessile or pedicellate, mature buds globose or depressed-globose, sepals fused for basal 1/3 of their length, with or without dorsal appendages, pubescent. Female flowers ovoid, subcompressed, perianth fused, tubular, rim entire, pubescent; stigma elongated, exerted, curved, hairs borne only on side. Fruiting perianth ovoid, ellipsoid or obovoid, without beak, ± flattened; surface conspicuously pubescent.

### Key to the species

1. Leaves alternate, inflorescences branched, flower clusters discrete, never contiguous *M. nivea*

1. Leaves opposite, inflorescences unbranched, fowers clusterss ± contiguous *M. ourantha*

***Muimar nivea*** (L.) A.K.Monro, Friis & Wilmot-Dear. **comb. nov.** *Urtica nivea* L. in Sp. Pl.: 985. 1753 ≡ *Boehmeria nivea* (L.) Gaudich. in Voy. Uranie: 499. 1830 ≡ *Ramium niveum* (L.) Kuntze in Revis. Gen. Pl. 2: 632. 1891, *nom. ilegit. superfl. ―* Lectoype (designated by Ghafoor, 1977: 18): Anon., Herb. Linn. 1111.19 (LINN 1111.19). Fig. 16.

*Boehmeria candicans* Hassk. in Cat. Pl. Hort. Bot. Bogor.: 79.1844 *≡ Urtica candicans* Blume (1825) 503, nom. illeg., non *Urtica candicans* Burm. f. (1768: 197) *≡ Boehmeria nivea* (L.) Gaudich. var. *candicans* (Hassk.) Wedd. in Prodr. [A. P. de Candolle]: 207. 1869 *―* **Neotype (designated here)**: [Indonesia] Java, *Zollinger 750* (G barcode G00354563*).

We were unable to trace the Blume type material at L or BO and so selected material collected from the type area, Java (Indonesia).

*Urtica tenacissima* Roxb., in [Wissett], Suppl. Observ. Sunn Hemp Bengal: 41. 1811 ≡ *Boehmeria tenacissima* (Roxb.) Blume in Mus. Bot. 2: 211. 1857 ≡ *Boehmeria nivea* var. *tenacissima* (Roxb.) Miq. in Fl. Ned. Ind. 1: 253. 1859 – Lectotype (designated by Turner, 2013: 166): [icon] Roxburgh drawing 1670, K) – Epitype (designated by Turner, 2013: 166): [cultivated in Calcutta Botanic Gardens] *W. Roxburgh s.n.* (BR barcode0000005203147*; ?isoepitypes: BM barcode 000835422*, G barcode G00354054*).

*Boehmeria mollicoma* Miq in H. Zollinger, Syst. Verz. Ind. Archip. 2: 100. 1854. ≡ *Ramium mollicoma* (Miq.) Kuntze in Revis. Gen. Pl. 2: 633. 1891 ― Holotype: [Indonesia] Java, *H. Zollinger 1454* (BO not traced; isotype: U barcode 1749531*).

*Boehmeria compacta* Blume in Mus. Bot. 2: 210. 1857 ―Holotype: [Indonesia] Java, *Anon.* (?Blume) *s.n.* (L barcode L0836071*).

*Boehmeria nivea* (L.) Gaudich. var. *reticulata* Blume in Mus. Bot. 2: 211. 1857 ― **Lectotype (designated here)**: Japan, *P.F. von Siebold s.n.* (L barcode L0836072*)).

*Boehmeria nivea* (L.) Gaudich. var. *concolor* Makino in Bot. Mag. (Tokyo) 23: 251. 1909 ≡ *Boehmeria frutescens* (Thunb.) Thunb. var. *concolor* (Makino) Nakai in Bot. Mag. (Tokyo) 41: 514. 1927 ≡ *Boehmeria nipononivea* Koidz. var. *concolor* (Makino) Ohwi in Fl. Japan: 441. 1953 ≡ *Boehmeria nivea* (L.) Gaudich. forma *concolor* (Makino) Kitam. in Acta Phytotax. Geobot. 20: 208. 1962 – **Lectotype (designated here)**: Japan, near Kyoto, Nov. 7 1893, *Makino* s.n. (MAK(124723*).

*Boehmeria nivea* (L.) Gaudich. var. *viridula* Yamam in J. Soc. Trop. Agric., Taihoku [Taipei] 4: 50. 1932 – Replacement name for *Boehmeria frutescens* (Thunb.) Thunb. var. *viridula* (Yamam.) T.Suzuki in Short Fl. Formos. (ed. Masamune G): 47. 1936 ≡ *Boehmeria nivea* (L.) Gaudich. forma *viridula* (Yamam.) Hatus. in Fl. Ryukyus: 234. 1971 ― Types: *Yamamoto 542* (not traced), Taiwan, Kwarenko Pr., Beisan*; Suzuki 1168* (not traced), Taiwan, between Doba and Taiheizan, 28 July 1928.

*Boehmeria nipononivea* Koidz. in Acta Phytotax. Geobot. 10: 223. 1941.

≡ *Boehmeria nivea* (L.) Gaudich. subsp. *nipononivea* (Koidz.) Kitam. in Acta Phytotax. Geobot. 20: 208. 1962

≡ *Boehmeria nivea* (L.) Gaudich. forma *nipononivea* (Koidz.) Hatus. in Fl. Ryukyus (S. Hatusima ed): 234. 1971 ≡ *Boehmeria nivea* (L.) Gaudich. var. *nipononivea* (Koidz.) W.T.Wang in Acta Bot. Yunnan. 3: 320. 1981 ― Type: No material cited or traced.

*Boehmeria thailandica* Yahara Acta Phytotax. Geobot. 32: 4. 1981 ― Holotype: Thailand, Phitsanulok, Thung Salang Luang, *Tagewa et al. 11233* (KYO barcode KYO00020601*).

***Muimar ourantha*** (Miq.) A.K.Monro, Friis & Wilmot-Dear. **comb. nov.** ≡ *Boehmeria ourantha* Miq. in Pl. Jungh.: 33. 1851 ≡ *Boehmeria caudata* (Burm.f.) J.J.Sm. var. *ourantha* (Miq.) J.J.Sm. in Meded. uitgaande Dep. Landbouw. Bd.12: 714. 1910 ≡ *Boehmeria platyphylla* D.Don var. *ourantha* (Miq.) Hochr. in Candollea 2: 344. 1925 ― Holotype: *Junghuhn s.n*., Indonesia, Jawa [Java], Bantam Dist. (U (U barcode 0256222*).

*Boehmeria pseudotomentosa* Yahara in Acta Phytotax. Geobot. 32: 16. 1981 – Holotype: Thailand, Loei, top area of Phu Kradung, Phu Kradung District, 1220 ― 1280 m, Nov. 17 1979, T. *Shimizu & al. 22957* (KYO barcode 00020600*).

*B. tomentosa* sensu auct., incl. Wedd. (p.p.), non *B. tomentosa* Wedd. s.str.;

*B. macrophylla* Hornem. var. *tomentosa* (Wedd.) D.G.Long, sensu D.G.Long, non var. *tomentosa* (Wedd.) D.G.Long, s.str.

**Etymology** The name *Muimar* is an anagram of *Ramium* an illegitimate name proposed by Kuntze based on the local name, ‘Ramie’.

**Discussion** Within the Boehmerieae, we recover *Muimar* as distantly related to *Boehmeria* and sister or congeneric to *Archiboehmeria* (Fig. 1). Whilst the type, *M. nivea* was recognised as atypical within the genus by Friis & Wilmot-Dear (Wilmot-Dear & Friis, 2013) and the species had been recovered as part of a large paraphyletic and polyphyletic *Boehmeria* (Wu et al., 2013, 2018) identifying synapomorphies has been challenging. In part because of the great morphological variation in *Boehmeria* as currently delimited, which comprises species with alternate or opposite leaves, partially or fully fused stipules, indefinite or definite inflorescences and the presence or absence of hooked hairs. An extensive review of herbarium collections enabled a unique combination of characters to be identified which enable *Muimar* to be distinguished from *Boehmeria* and *Archiboehmeria*

**Notes** Indomalaya, 2 spp. The misapplication of the epithet “tomentosa” to material of *B. ourantha* is common in the literature (Chen et al., 2005). The combination, *B. macrophylla* Hornem. var. *tomentosa* (Wedd.) Long, based on the Afro-Malagasy species *B. tomentosa* Wedd., was first misapplied to Asian material in the Flora of Bhutan (Grierson & Long, 1983: 126). Friis & Wilmot-Dear (Wilmot-Dear & Friis, 2013: 181) distinguish *Boehmeria virgata* subsp. *macrophylla* var. *tomentosa* from *B. ourantha* in the arrangement of male flowers, colour, margin and shape of the leaves, and the spacing of the flowers along the inflorescence.

[Table 7. Morphological characters which distinguish *Muimar* from *Boehmeria* and *Archiboehmeria*]

**Table 7.** Morphological characters which distinguish *Muimar* from *Boehmeria* and *Archiboehmeria*.

|  | *Boehmeria* | *Archiboehmeria* | *Muimar* |
| --- | --- | --- | --- |
| **Stem pubescence** | hairs hooked or not | hairs hooked | hairs not hooked |
| **Degree of stipule fusion** | fully fused to fused for < 1/3 of their length | Stipules fused for > 1/2 of their length | Stipules fused for < 1/3 of their length |
| **Point of stipule insertion** | interpetiolar | interpetiolar | intrapetiolar |
| **Leaf pubescence** | hairs not shiny / glistening | hairs not shiny / glistening | hairs shiny / glistening |
| **Bisexual flowers** | absent | present | absent |
| **Male flowers** | Sepals fused for < 1/3 of their length | Sepals fused for ca 1/2 of their length | Sepals fused for < 1/3 of their length |
| **Female inflorescences** | sessile, pedunculate or on modified stems | pedunculate | pedunculate |

42. ***Archiboehmeria*** C.J.Chen in Acta Phytotax. Sin. 18: 479. 1980 – Type: *Archiboehmeria atrata* (Gagnep.) C.J.Chen in Acta Phytotax. Sin. 18: 479. 1980 ≡ *Debregeasia atrata* Gagnep. in Bull. Soc. Bot. France 75: 556. 1928.

**Notes** Indomalaya (China, Vietnam); 1 sp.

43. ***Astrothalamus*** C.B.Rob. in Philipp. J. Sci., Bot. 6: 19. 1911 – Type: *Astrothalamus reticulatus* (Wedd.) C.B.Rob. in Philipp. J. Sci., Bot. 6: 19. 1911 – **Lectotype (designated here)**: Guam, *Gaudichaud 167* (P barcode 00698510).

**Notes** Indomalaya; 1 sp.

44. ***Debregeasia*** Gaudich. in Voy. Bonite, Bot., Atlas: t. 90. (1844) – Lectotype (selected by Wilmot-Dear 1988): *Debregeasia velutina* Gaudich. in Bot., Voy. Bonite: t. 90. 1844.

*Leucocnide* Miq. in Pl. Jungh.: 36. 1851 – Type: *Leucocnide dichotoma* (Blume) Miq. in Pl. Jungh.: 38. 1851 ≡ *Urtica dichotoma* Blume in Bijdr. Fl. Ned. Ind. 10: 499. 1825-6 – Type: Indonesia, *Blume s.n.* (L barcode L0039694).

*Morocarpus* Siebold & Zucc. in Abh. Math.-Phys. Cl. Königl. Bayer. Akad. Wiss. 4(3): 218. 1846, nom. ilegit., non *Morocarpus* Boehm (1760: 385).

**Notes** AfroTropcs, Palearctic, Indomalaya; 9 spp.

#### LEUKOSYKEAE. Fig. 11

45. ***Leukosyke*** Zoll. & Moritzi in Syst. Verz.: 100. 1845. Type: *Leukosyke javanica* Zoll. & Moritzi in Syst. Verz.: 100. 1845 – **Lectotype (designated here)**: Indonesia, *Zollinger, H. 629,* (G barcode G00354275).

=*Missiessya* Gaudich. ex Wedd. in Ann. Sci. Nat., Bot., sér. 4, 1: 194. 1854 – Type: *Missiessya wallichiana* Wedd. in Ann. Sci. Nat., Bot., sér. 4, 1: 195. 1854.

**Notes** Indomalaya, Australasia; 33 spp.

46. ***Sarcochlamys*** Gaudich. in Voy. Bonite, Bot. t. 89. 1844 – Type: *Sarcochlamys pulcherrima* Gaudich. in Voy. Bonite, Bot. t. 89. 1844.

=*Sphaerotylos* C.J.Chen in Acta Phytotax. Sin. 23: 452. 1985 – Type: *Sphaerotylos medogensis* C. J.Chen in Acta Phytotax. Sin. 23: 453. 1985.

**Notes** Palearctic, Indomalaya; 1 sp.

47. ***Maoutia*** Wedd. in Ann. Sci. Nat., Bot., sér. 4, 1: 193. 1854 – Type: *Maoutia platystigma* Wedd. in Ann. Sci. Nat., Bot., sér. 4, 1: 194. 1854 – **Lectotype (designated here)**: Phillipines, *Cumming 1441,* (P barcode P00698294; isolectotypes: G barcode G00354217, G barcode G00354220, P barcode P00698295).

=*Lecanocnide* Blume in Mus. Bot. 2: t12. 1857 – Type: *Lecanocnide diversifolia* Blume in Mus. Bot. 2: t12. 1857.

=*Robinsoniodendron* Merr. in Interpr. Herb. Amboin.: 204. 1917 – Type: *Robinsoniodendron ambiguum* (Wedd.) Merr. in Interpr. Herb. Amboin.: 204. 1917.

**Notes** Australasia (New Guinea), Indomalaya; 10 spp. Wilmot-Dear & Friis (Wilmot-Dear & Friis, 2013) suggest that *Maoutia* be included within *Leukosyke.* Our results suggest (Fig. 3 & 4.1) that *Maoutia* is instead sister to the New Guinean endemic, *Gibbsia,* which itself is morphologically distinct from both *Maoutia* and *Leukosyke* in its inflorescence and pistillate flower morphology, and the in disposition of the achene relative to the perianth in fruit. We therefore decided to maintain all three as distinct genera.

48. ***Gibbsia*** Rendle in Fl. Arfak Mts.: 129. 1917 – Type: *Gibbsia insignis* Rendle in Fl. Arfak Mts.: 130. 1917 – **Lectotype (designated here)**: Indonesia, *L.S. Gibbs 5961* (BM barcode BM000571389; isotype: K barcode K000741628 (2 leaves and an inflorescence fragment only)).

**Notes** Australasia (New Guinea); 2 spp.

#### CECROPIEAE. Fig. 12

49. ***Cecropia*** Loefl. in Iter Hispan.: 272. 1758 – Type: *Cecropia peltata* L. in Syst. Nat. (ed. 10): 1286. 1759.

=*Ambaiba* Adans. in Fam Pl. 2: 377. 1763 – Type: *Ambaiba peltata* (L.) Kuntze in Revis. Gen. Pl. 2: 623. 1891.

=*Coilotapalus* P. Browne in Civ. Nat. Hist. Jamaica: 111. 1756. nom. nud.

**Notes** Indomalaya, Neotropics; *ca* 65 spp.

50. ***Pourouma*** Aubl. in Hist. Pl. Guiane 2: 891–892, t. 341. 1775 – Type: *Pourouma guianensis* Aubl. in Hist. Pl. Guiane 2: 892, t. 341. 1775.

**Notes** Neotropics; 31 spp.

51. ***Coussapoa*** Aubl. in Hist. Pl. Guiane: 2: 955, t. 362. 1775 – Type: *Coussapoa latifolia* Aubl. in Hist. Pl. Guiane 2: 955. 1775 – **Lectotype (designated here)**: French Guiana, *Aublet s.n.*: (BM barcode BM000993414).

**Notes** Nearctic, Neotropics; 50 spp.

52. ***Musanga*** R.Br. in Pl. Jav. Rar. (Bennett): 49, 1838 – Type: *Musanga smithii* R.Br. in Pl. Jav. Rar. (Bennett): 49, 1838 – **Lectotype (designated here)**: Congo, *Smith s.n.,* (BM barcode BM000911480).

**Notes** Afrotropics; 2 spp.

53. ***Myrianthus*** P.Beauv. in Fl. Oware 1: 16. 1805. Type: *Myrianthus arboreus* P. Beauv. in Fl. Oware 1: 17, t. 11. 1805. Type: Benin, *Palisot de Beauvois* s.n. (lectotype designated here), G barcode G00008686. **Notes**ϑ Afrotropics; 8 spp.

#### Unplaced names

*Metapilea* W.T.Wang in Bull. Bot. Res., Harbin 36: 166. 2016. – Type: *Metapilea jingxiensis* W.T.Wang in Bull. Bot. Res., Harbin 36: 166. 2016. Please see notes under *Pilea*.

*Parsana* Parsa & Maleki Fl. Iran, Suppl. Gen.: 548. 1952. The genus *Parsana* Parsa & Maleki likely belongs within this genus. The description and illustration do not correspond to any known specimens, the type collections having not been deposited where stated in the description (Mahmoud Bidarlord (FAR), Pers. Comm.) could not be located or sampled and no other collections match the description. Based on the locality information and aspects of the illustrations, however, we believe that the description may have been based on material of *Laportea*.

*Pellionia* Gaudich. in Voy. Uranie: 494. 1830 – Type: *Pellionia elatostemoides* Gaudich. in Voy. Uranie: 494. 1830 (typ. cons.). See notes under *Elatostema*.

## Acknowledgements

We would like to thank curators at B, BM, F, GH, K, MO, P, TEX, US for the loan of herbarium collections over the years; Sven Buerki for earlier iterations of the analyses of a subset of the Sanger sequence data; Stephen Russell (Natural History Museum, London) for support generating Sanger sequence data. We would also like to thank the Plant and Fungal Trees of Life Project and the Calleva Foundation for the generation of the Angiosperms353 data.

## Supplementary materials

Table S1 Molecular accessions

Table S2 PROTEUS Fossil calibrations output, 2023-07-19

Fig. S1 Plot of loci sampled by accession

Fig. S2 Comparison of phylograms generated by RAXML and Bayesian analyses of the Sanger sequence data

Fig. S3 Ancestral state reconstructions for 21 morphological characters / traits

