## Supplementary figures and images for "Classification of Urticaceae based on morphology and phylogenetic inference"

### Supplemental Fig 1

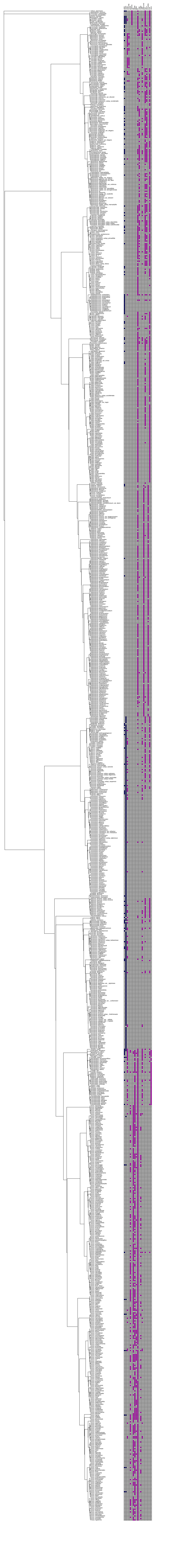
