## Supplemental Fig 3 for "Classification of Urticaceae based on morphology and phylogenetic inference"

**trait 2: Growth.habit,.herb.(0),.woody.climber.(1),.shrub.(2),.tree.(3),.herbaceous.climber.(4),.hemiepiphyte.(5),.epiphyte.(6)**

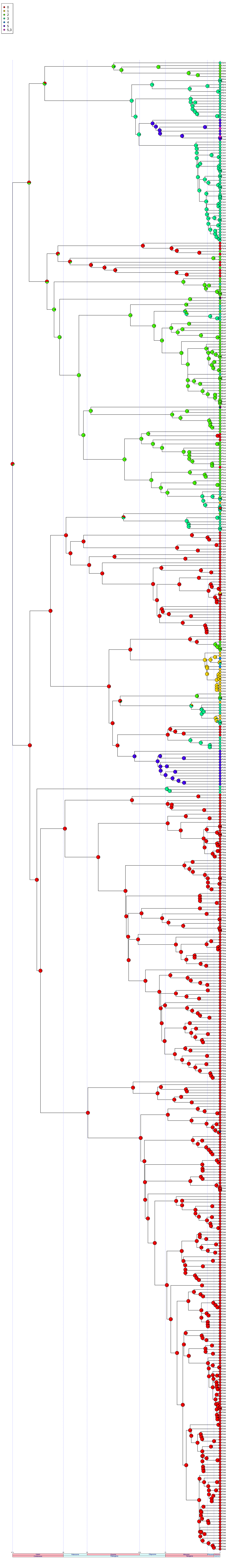

trait 3: Latex.present.(0),.absent.(1)

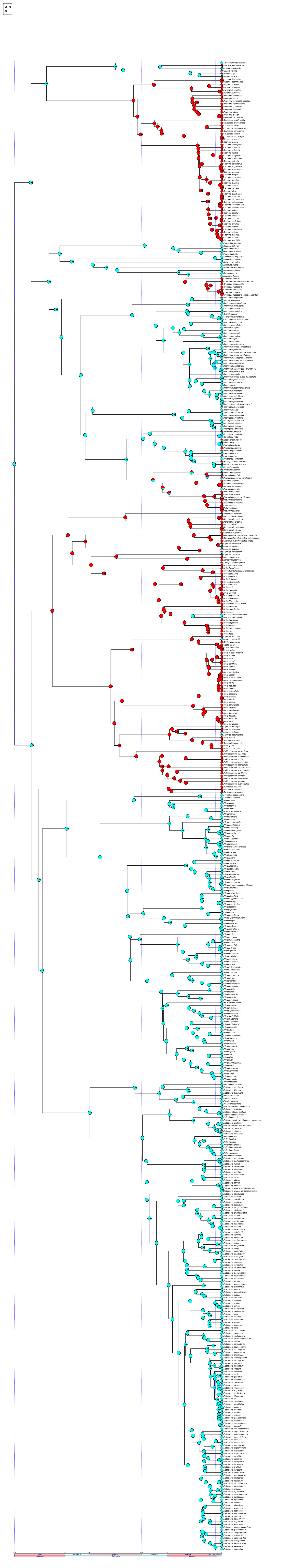

### trait 4: uncinate.hairs.present.(0),.absent.(1)

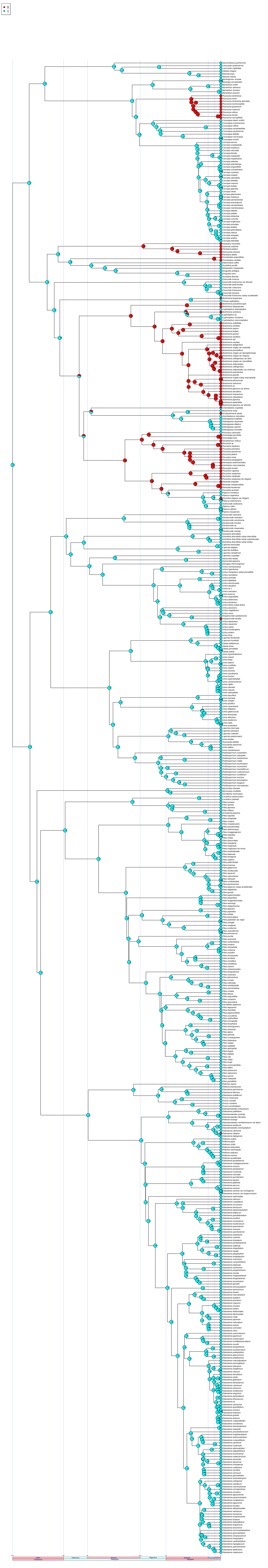

**trait 5: Cystoliths.present.(0),.absent.(1)**

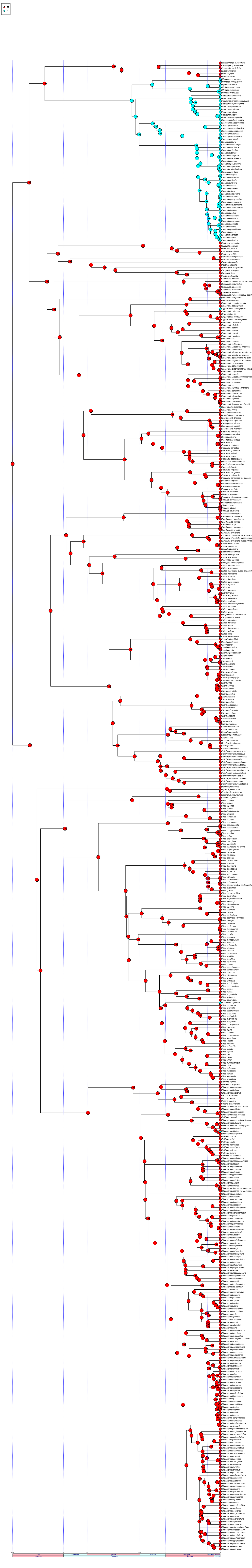

trait 6: Cystolith,shape,,spindle−fusiform,(0),,oblong−elliptic,(1),,punctiform,(2)

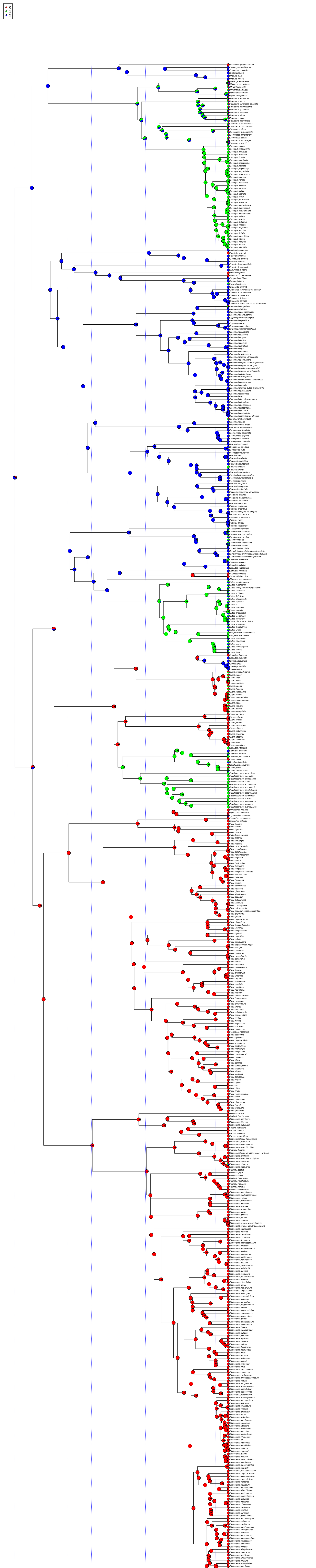

**trait 7: stipule.present.(0),.absent.(1)**

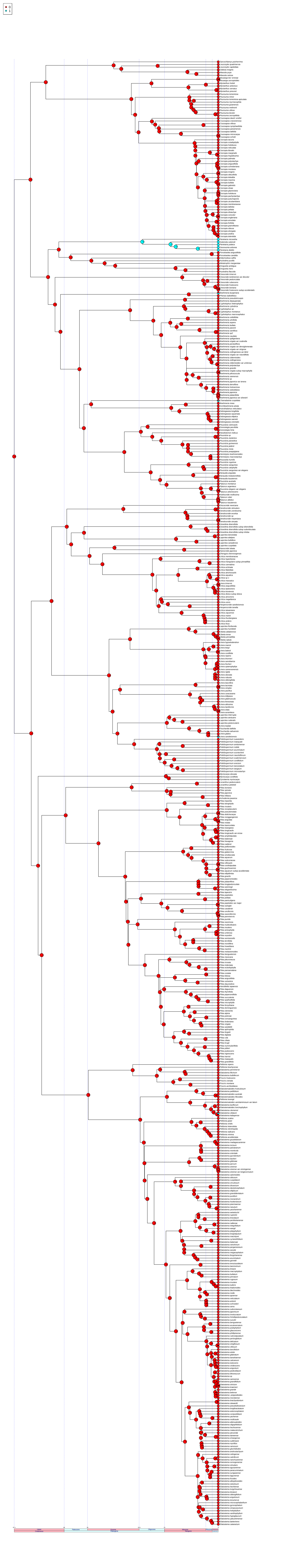

**trait 8: stipule.not.amplexicaulous.(0),.amplexicaulous.(1)**

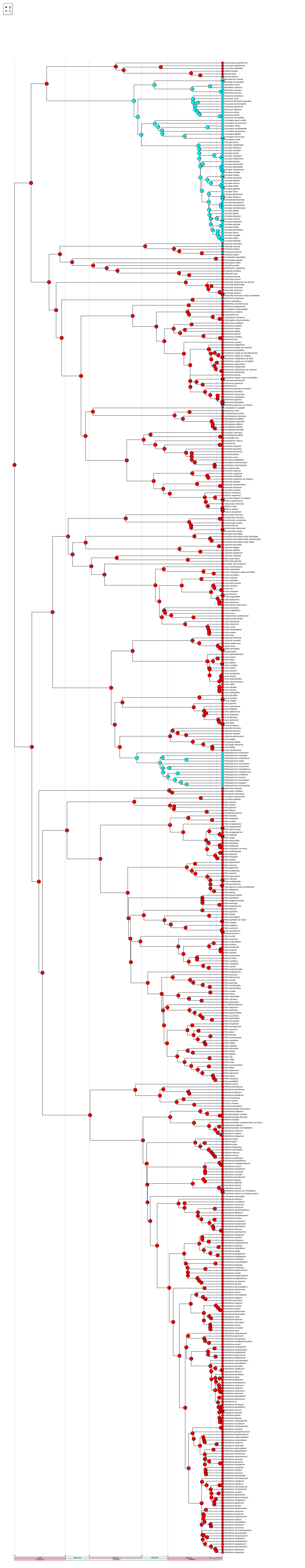

**trait 9: bulbous.based.hairs.present.(0),.lacking.(1)**

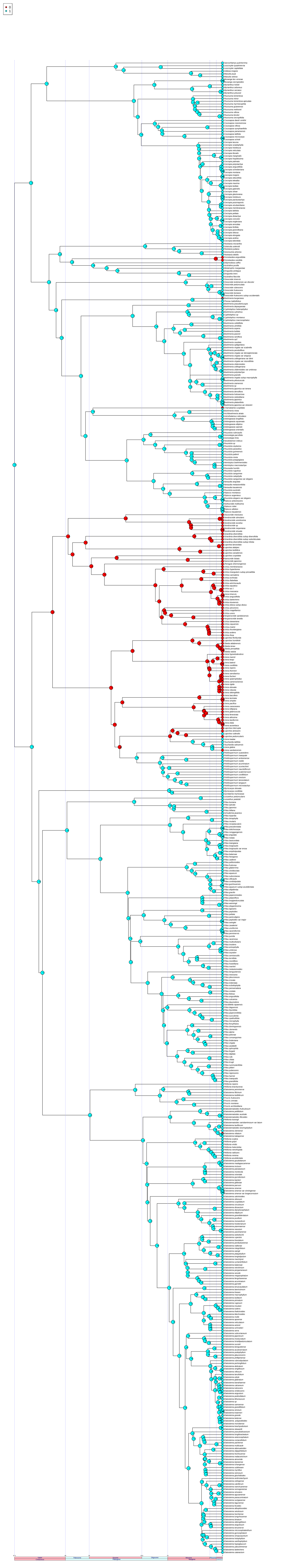

### trait 10: Pistillode.present.(0),.absent.(1)

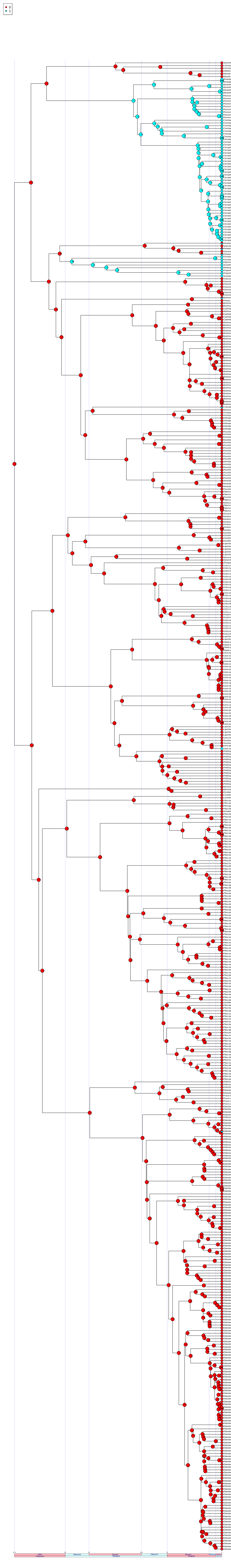

### trait 11: Pistillode.pubescent.(0),.glabrous.(1)

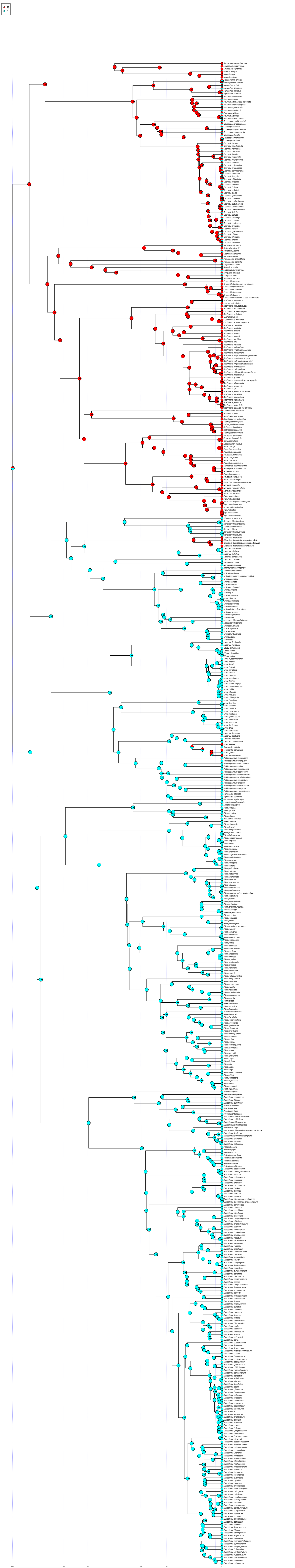

trait 12: P.perianth.parts.free.(0),.partly.or.fully.fused.(1),.absent.(2)

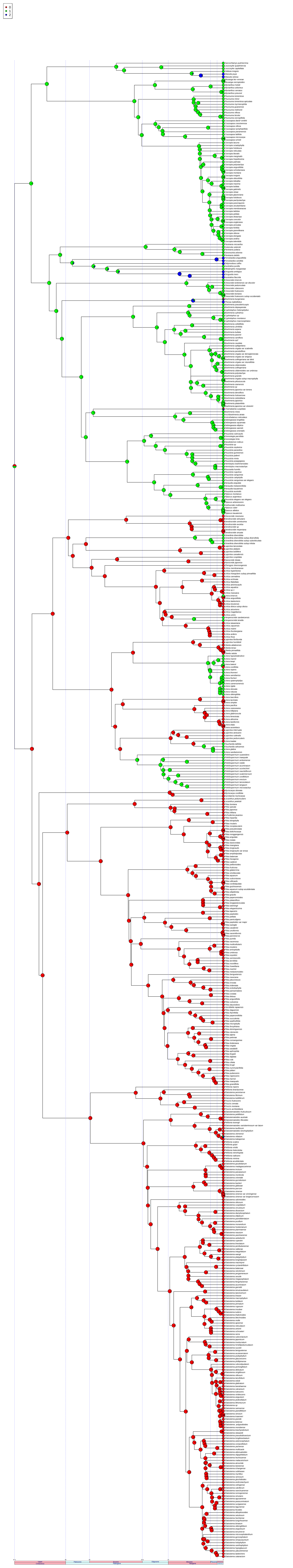

trait 13: Pistil.insertion.angle.erect.(0),.<45.(1),.>45.(2)

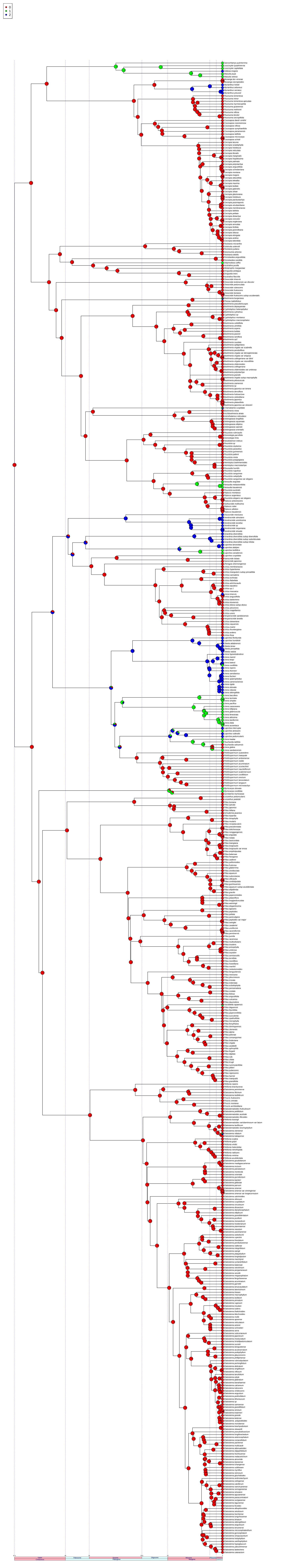

trait 14: Stigma.hair.dist,.apical.only,(0),.lateral.asymm.(1),.lateral.symm.(2)

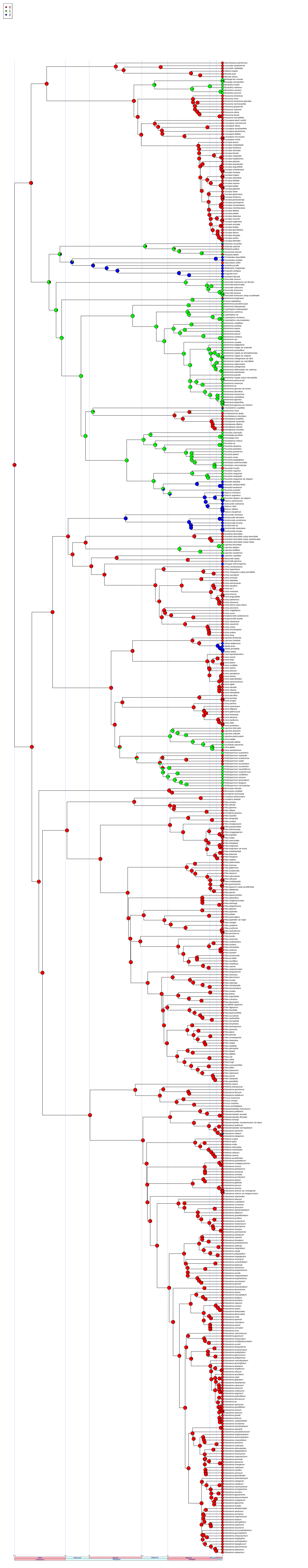

trait 15: Perianth.fleshy.in.fruit.(0),.not.(1)

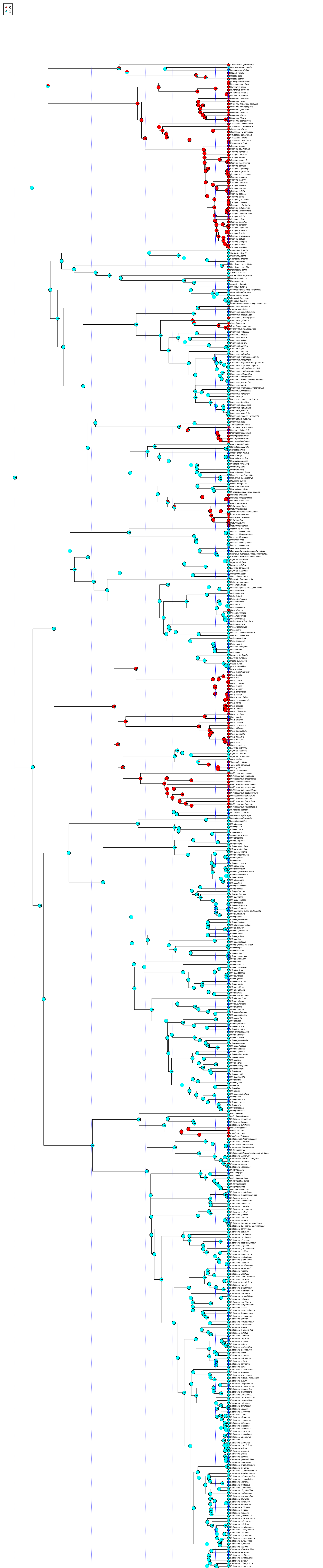

### trait 16: staminate.sepal.envelopes.individual.filament.(0)..not.(1)

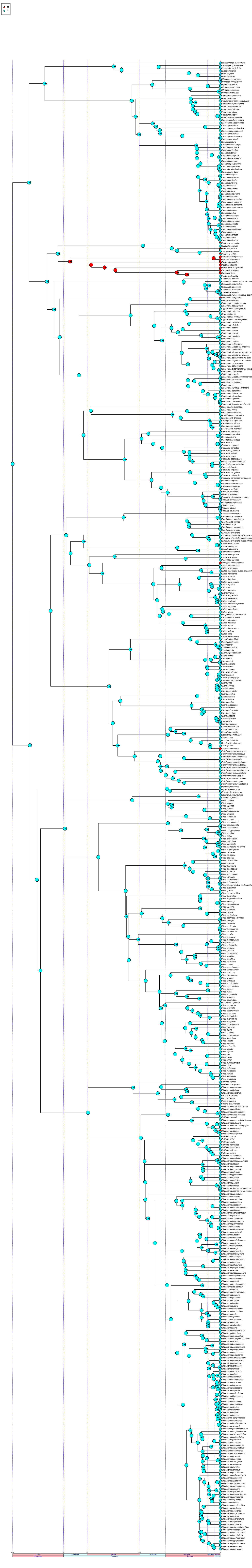

trait 17: stigma.style.spreading/extending.(0),.or.not.(1)

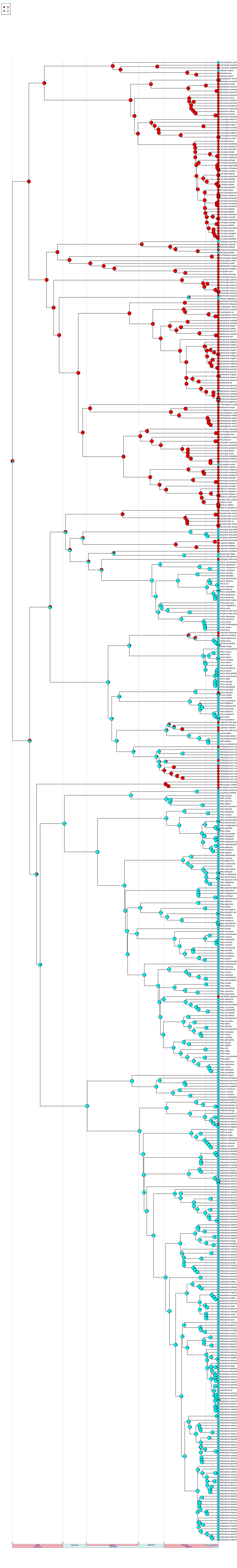

trait 18: stigma.receptive.area.adaxial.+.abaxial.surfaces.and.+/−.confluent.basally.(0),.not.as.before.(1),.NA

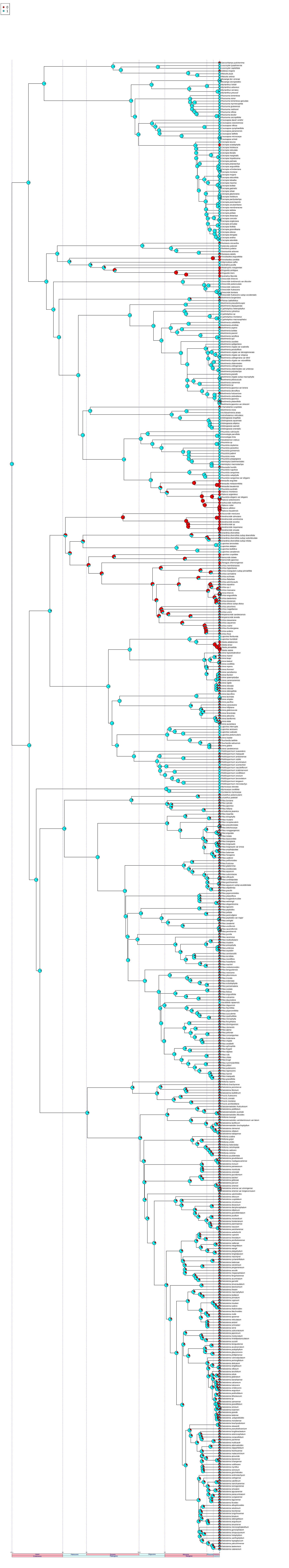

trait 19: Inflorescences.unisexual.(0),.or.bisexual.(1)

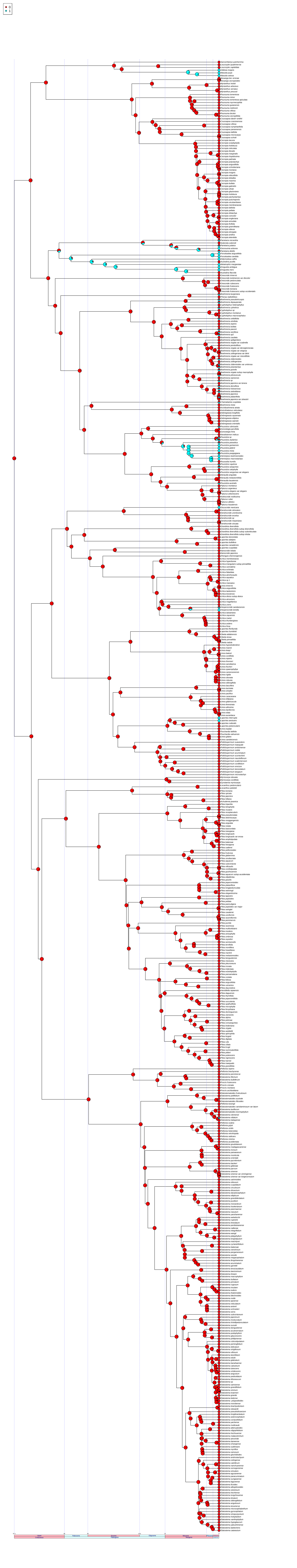

trait 20: filaments.straight.(0).or.inflexed.(1).in.bud

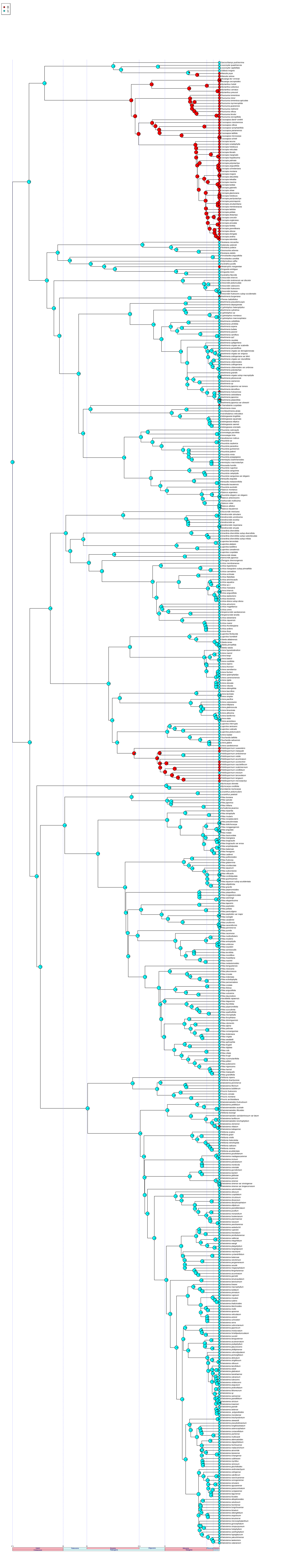

trait 21: fruits.fleshy.(1).or.not.fleshy.(0)

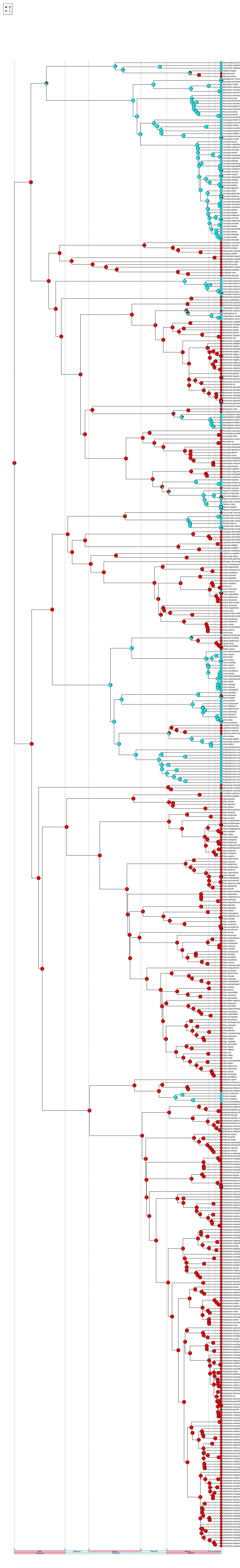

trait 22: Pistillate.inflorescence.branches,.(0).free,.(1).partially.fused,.(2).fully.fused

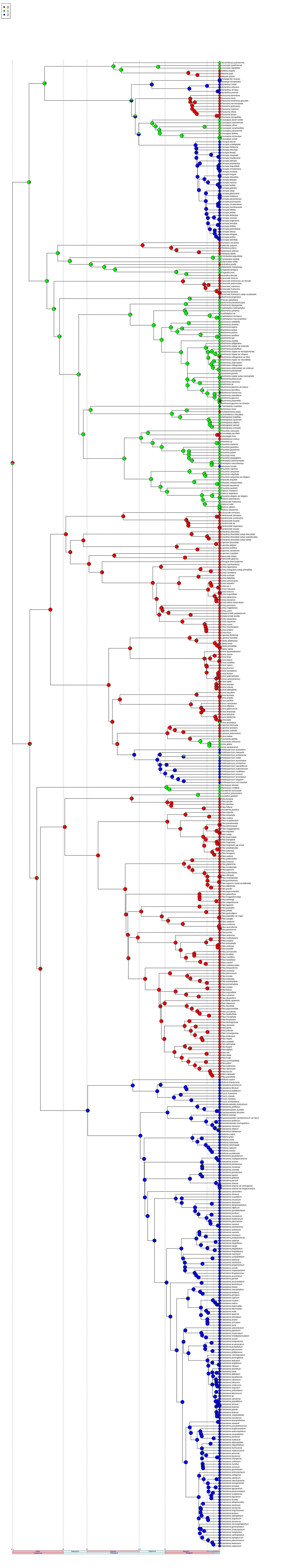

### trait 23: Pistillate.fused.inflorescences.(0).raceme/spike,.(1).syconium,.(2).capitulum,.(3).bivalvate,.(4).NA

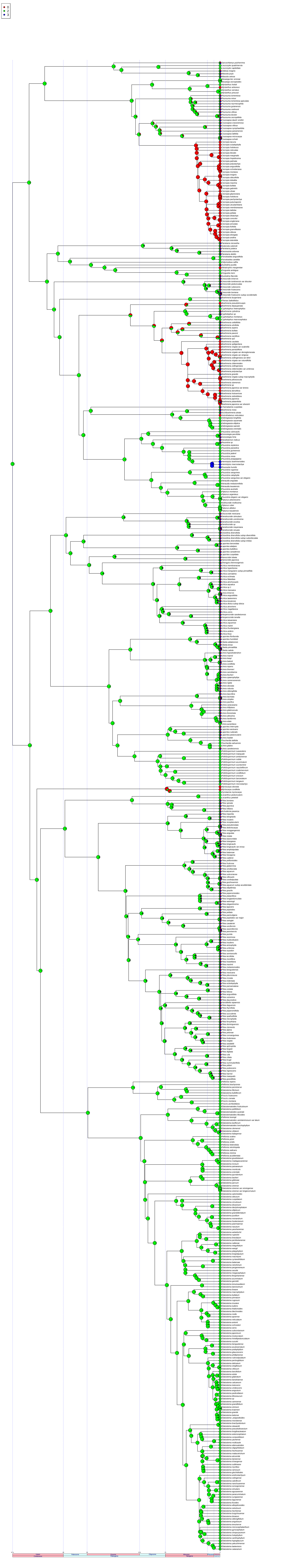

trait 24: Pistillate.fused.inflorescences,.(0).globose.to.ellipsoid,.(1).disc./.flattened,.(2).urceolate,.(3).cylindrical.to.turbinate,.(4).cymose,.(5).NA

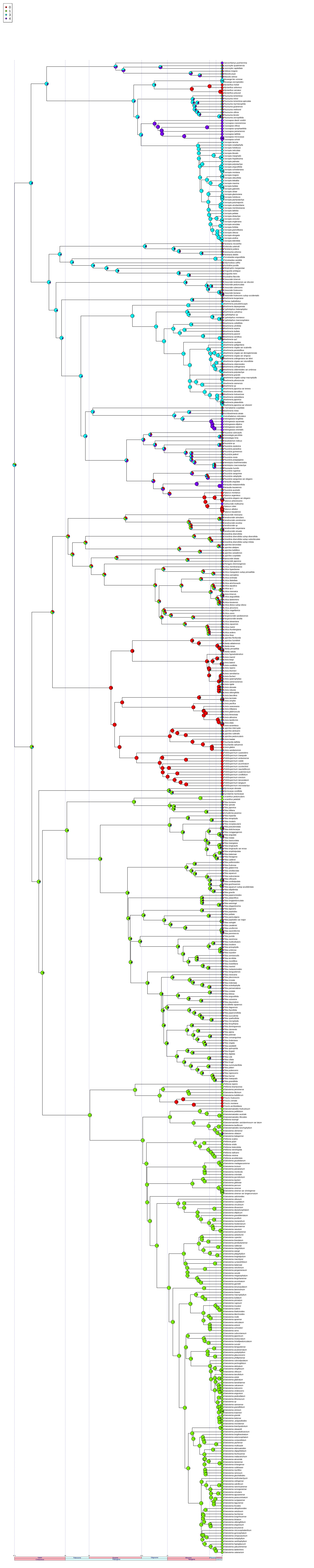
