## Appendix 1 for "Classification of Urticaceae based on morphology and phylogenetic inference"

| **Sequence data** | **Family** | **Taxon** | **Voucher** | **Collector No** | **Herbarium** | **Country** | **RBGKew BankId** | **18S** | **ITS** | **trnLF** | **rbcL** | **rpll4-rps8- infA-rpl36** | **matK** | **Angiosperms3 53 (A353)** |
| --- | --- | --- | --- | --- | --- | --- | --- | --- | --- | --- | --- | --- | --- | --- |
| Angiosperms353 | Cannabaceae | *Trema orientale* | Reeves, G.& | 37 | K | South Africa | 27834 | – | – | – | – | – | – | ERS8701987 |
|  |  |  | Weiblen, G.D.; Montgomery, R.; |  |  | – |  |  |  |  |  |  |  |  |
| Angiosperms353 | Moraceae | *Antiaropsis decipiens* | Isua, B.; Molem, K. | 1865 | K |  | – | – | – | – | – | – | – | ERS4414199 |
| Angiosperms353 | Moraceae | *Batocarpus orinocensis* | Palacios, W. | 3265 | K | – | – | – | – | – | – | – | – | ERS4414228 |
|  |  |  |  |  |  | Korea (the |  |  |  |  |  |  |  |  |
| Angiosperms353 | Moraceae | *Broussonetia kazinoki* | Chase, M.W. | 17827 | K | Republic of) | 17657 | – | – | – | – | – | – | ERS4414202 |
| Angiosperms353 | Moraceae | *Ficus sagittifolia* | Chase, M.W. | 19852 | K | Ivoryt Coast | 19722 | – | – | – | – | – | – | ERS4414205 |
|  |  |  | Wurdack, K.J.; Redden, K.; |  |  | – |  |  |  |  |  |  |  |  |
|  |  |  | Rodriguez, A.; Perry, C.; James, |  |  |  | | | | | | | | |
| Angiosperms353 | Moraceae | *Maquira guianensis* | H.; Simon, H.; Ragnauth, P. | 4570 | K | – – – – – – – To submit | | | | | | | | |
| Angiosperms353 | Moraceae | *Milicia africana* | Williamson, L. | 187 | K | – – – – – – – – ERS4414198 | | | | | | | | |
| Angiosperms353 | Moraceae | *Parartocarpus venenosa* | Renvoize, S.A.; Wilmot-Dear, M.; Saenz, A. | 1518 | K | – | – | – | – | – | – | – | – | ERS5501666 |
| Sanger | Urticaceae | *Achudemia javanica* | Robinson H.C. and Kloss C.B. | 1914 | SING | Indonesia | – | – | MT516339 | MT523094 | MT523050 | – | – | – |
| Sanger | Urticaceae | *Archiboehmeria atrata* | WuZY | 9469 | KUN | China, Guangxi | – | – | KF137798 | KF138269 | KF138106 | KF138434 | KF137946 | – |
| Angiosperms353 | Urticaceae | *Archiboehmeria atrata* | Hiep, N.T. et al. | 189 | K | Viet Nam | 23937 | – | – | – | – | – | – | ERS5501742 |
| Angiosperms353 / Sanger | Urticaceae | *Astrothalamus reticulatus* | Argent, G. et al. | 981987 | K |  | 23941 | – | KF137800 | KF138271 | KF138108 | – | To submit | ERS5501745 |
| Sanger | Urticaceae | *Australina flaccida* | Friis, I. et al. | 12293 | K | Ethiopia | – | KF137745 | KF137801 | KF138272 | KF138109 | KF138436 | MH357978 | – |
| Angiosperms353 | Urticaceae | *Australina pusilla* | Raven, P.H.; et al. | 25912 | K | Australia | 23949 | – | – | – | – | – | – | ERS5501750 |
| Sanger | Urticaceae | *Boehmeria aspera* | H. J. Sarrazola | 937 | HUA | Colombia | – | – | – | MH151324 | – | – | MH151315 | – |
| Sanger | Urticaceae | *Boehmeria bullata* | H. J. Sarrazola | CHG-46 | HUA | Colombia | – | – | – | – | – | – | MH151320 | – |
| Angiosperms353 / Sanger | Urticaceae | *Boehmeria burgerania* | Monro A.K. | 6842 | K | Costa Rica | – | – | To submit | – | – | – | – | ERS5503004 |
| Sanger | Urticaceae | *Boehmeria caudata* |  |  | E | Peru | – | – | – | MH358260 | MH358040 | – | – | – |
|  |  |  |  |  |  | Bolivia |  |  |  |  |  |  |  |  |
|  |  |  |  |  |  | (Plurinational State |  |  |  |  |  |  |  |  |
| Angiosperms353 | Urticaceae | *Boehmeria caudata* | Wood, J.R.I. | 18657 | K | of) | 24013 | – | – | – | – | – | – | ERS5501752 |
| Sanger | Urticaceae | *Boehmeria celtidifolia* | H. J. Sarrazola | 957 | HUA | Colombia | – | – | – | MH151327 | – | – | MH151323 | – |
| Sanger | Urticaceae | *Boehmeria clidemioides* | Nie | 4243 | KUN | China | – | – | KM586402 | KM586574 | KM586488 | MK955173 | MK931191 | – |
| Sanger | Urticaceae | *Boehmeria clidemioides var. umbrosa* | WuZY | 10336 | KUN | China, Yunna | – | – | KF137822 | KF138293 | KF138130 | KF138453 | KF137965 | – |
| Sanger | Urticaceae | *Boehmeria cylindrica* | Abbott | 18035 | FLAS | USA? | – | – | – | – | KJ773314 | – | KJ772586 | – |
| Sanger | Urticaceae | *Boehmeria densiflora* | WuZY | 2012450 | KUN | Taiwan | – | – | KF137806 | KF138277 | KF138114 | – | KF137951 | – |
| Sanger | Urticaceae | *Boehmeria depauperata* |  |  | KUN | Yunnan, China | – | – | KF137807 | KF138278 | KF138115 | KF138440 | – | – |
| Sanger | Urticaceae | *Boehmeria grandis* | Morden 1120 | 1120 | BISH | Hawaii | – | – | – | – | AF500354 | – | – | – |
| Sanger | Urticaceae | *Boehmeria holosericea* | TKMVP | 873 | ?? | Korea | – | – | KT119556 | – | – | – | – | – |
| Sanger | Urticaceae | *Boehmeria japonica* |  | 100024 | KUN |  | – | – | KF137808 | KF138279 | KF138116 | – | – | – |
|  |  |  |  |  | Herbarium of | China? |  |  |  |  |  |  |  |  |
| Sanger | Urticaceae | *Boehmeria japonica var. silvestrii* | Li | 26 | Jiujiang University |  | – | – | FJ750380 | FJ750411 | – | – | – | – |
| Sanger | Urticaceae | *Boehmeria japonica var. tenera* | D. Tao | 84 | ?? | China? | – | – | MK911052 | MK911074 | MK911097 | – | MK931192 | – |
| Sanger | Urticaceae | *Boehmeria nivea* | Liuj | 10645 | KUN | China, Fujian | – | – | KF137815 | KF138286 | KF138123 | KF138447 | KF137958 | – |
| Angiosperms353 | Urticaceae | *Boehmeria nivea* | Hu, S.Y. | 8113 | K | China | 24011 | – | – | – | – | – | – | ERS4414210 |
| Sanger | Urticaceae | *Boehmeria nivea var. tenacissima* | Liuj | 10679 | KUN | China, Zhejiang | – | – | KF137814 | KF138285 | KF138122 | KF138446 | KF137957 | – |
| Sanger | Urticaceae | *Boehmeria pavonii* | H. J. Sarrazola | 933 | HUA | Colombia | – | – | – | MH151326 | – | – | MH151319 | – |
| Sanger | Urticaceae | *Boehmeria penduliflora* | WuZY | 9460 | KUN | China, Guangxi | – | – | KF137816 | KF138287 | KF138124 | KF138448 | KF137959 | – |

|  |  |  |  |  | Herbarium of  Jiujiang University | China? |  |  |  |  |  |  |  |  |
| --- | --- | --- | --- | --- | --- | --- | --- | --- | --- | --- | --- | --- | --- | --- |
| Sanger | Urticaceae | *Boehmeria pilosiuscula* | Li | 17 |  |  | – | – | FJ750372 | FJ750422 | – | – | – | – |
| Sanger | Urticaceae | *Boehmeria platanifolia* |  | AM6715 | NA | South Korea | – | – | – | – | KM218340 | – | – | – |
|  |  |  |  |  | Herbarium of Jiujiang University | China? |  |  |  |  |  |  |  |  |
| Sanger | Urticaceae | *Boehmeria polystachya* | Li | 29 |  |  | – | – | FJ750376 | FJ750421 | – | – | – | – |
|  |  |  |  |  | Herbarium of Jiujiang University | China? |  |  |  |  |  |  |  |  |
| Sanger | Urticaceae | *Boehmeria pseudotricuspis* | Li | 12 |  |  | – | – | FJ750375 | FJ750400 | – | – | – | – |
| Sanger | Urticaceae | *Boehmeria ramiflora* | L0942407 | L0942407 | NHN | Jamaica | – | – | MH357855 | – | MH358042 | – | – | – |
|  |  |  |  |  | Herbarium of Jiujiang University | China? |  |  |  |  |  |  |  |  |
| Sanger | Urticaceae | *Boehmeria siamensis* | Li | 28 |  |  | – | – | FJ750374 | FJ750428 | – | – | – | – |
| Sanger | Urticaceae | *Boehmeria sieboldiana* |  | LiDZ1080 | E | Japan | – | – | MH357859 | – | MH358045 | MH358182 | – | – |
| Sanger | Urticaceae | *Boehmeria sp.* | RC | 1555 | KUN | Nepal, Mayagdi | – | – | KF137818 | KF138289 | KF138126 | KF138450 | KF137961 | – |
| Sanger | Urticaceae | *Boehmeria sp. 1* | Nie | 4249 | KUN | China? | – | – | – | KM586573 | KM586487 | – | – | – |
| Sanger | Urticaceae | *Boehmeria sp.2* |  |  | KUN | Dominica | – | – | MH357861 | MH358261 | MH358049 | – | MH357981 | – |
|  |  |  |  |  | Ramie Repository, Huazhong Agricultural University,  Wuhan, China | China |  |  |  |  |  |  |  |  |
| Sanger | Urticaceae | *Boehmeria splitgerbera* | HZAURS | D9 |  |  | – | – | – | HQ380834 | – | – | – | – |
| Sanger | Urticaceae | *Boehmeria tsaratananensis* | Razanajtovo et al. | MHR 001 | G | Madagascar | – | – | – | – | – | – | – | – |
| Sanger | Urticaceae | *Boehmeria ulmifolia* | H. J. Sarrazola | 929 | HUA | Colombia | – | – | – | MH151325 | – | – | MH151317 | – |
| Sanger | Urticaceae | *Boehmeria virgata subsp. macrophylla* | WuZY | 9196 | KUN | China? | – | – | KF137811 | KF138282 | KF138119 | KF138443 | KF137954 | – |
|  |  |  |  |  | Herbarium of Jiujiang University | China? |  |  |  |  |  |  |  |  |
| Sanger | Urticaceae | *Boehmeria virgata var. densiglomerata* | Li | 24 |  |  | – | – | FJ750378 | FJ750417 | – | – | – | – |
| Sanger | Urticaceae | *Boehmeria virgata var. macrostachya* | HZAURS | X3 | ?? | China? | – | – | – | HQ380837 | – | – | – | – |
| Sanger | Urticaceae | *Boehmeria virgata var. rotundifolia* | Liuj | 10629 | KUN | China, Yunnan | – | – | KF137812 | KF138283 | KF138120 | KF138444 | KF137955 | – |
| Sanger | Urticaceae | *Boehmeria virgata var. scabrella* | WuZY | 9468 | KUN | China, Guangxi | – | – | KF137813 | KF138284 | KF138121 | KF138445 | KF137956 | – |
|  |  |  |  |  | Herbarium of Jiujiang University | China? |  |  |  |  |  |  |  |  |
| Sanger | Urticaceae | *Boehmeria virgata var. strigosa* | Li | 21 |  |  | – | – | FJ750383 | FJ750418 | – | – | – | – |
| Sanger | Urticaceae | *Boehmeria virgata var. tomentosa* | WuZY | 9011 | KUN | China, Yunnan | – | – | KF137820 | KF138291 | KF138128 | KF138452 | KF137963 | – |
|  |  |  |  |  | Herbarium of Jiujiang University | China? |  |  |  |  |  |  |  |  |
| Sanger | Urticaceae | *Boehmeria zollingeriana* | Li | 8 |  |  | – | – | FJ750366 | FJ750424 | – | – | – | – |
| Sanger | Urticaceae | *Boehmeria zollingeriana var. blinii* | LiDZ | 1084 | KUN | China, Guizhou | – | – | KF137824 | KF138295 | KF138132 | – | KF137966 | – |
| Sanger | Urticaceae | *Cecropia angustifolia* | Monro 4424 | 4424 | BM | Panama | – | – | – | – | – | – | – | – |
| Angiosperms353 / Sanger | Urticaceae | *Cecropia ficifolia* | Berg, C.C. et al. | 18418 | K | Brazil | 23955 | – | KF137825 | KF138296 | KF138133 | – | – | ERS4414209 |
| Sanger | Urticaceae | *Cecropia glazioviana* | Bruno | 16 | UFP | Brazil | – | – | MH357864 | MH358262 | MH358053 | – | – | – |
| Sanger | Urticaceae | *Cecropia hololeura* | Bruno | 15 | UFP | Brazil | – | – | – | MH358263 | MH358054 | – | MH357985 | – |
| Sanger | Urticaceae | *Cecropia obtusifolia* | Monro 3767 | 3767 | BM | El Salvador | – | – | – | KF138297 | KF138134 | KF138455 | KF137967 | – |
| Sanger | Urticaceae | *Cecropia pachystachya* | Bruno | 17 | UFP | Brazil | – | – | MH357866 | MH358265 | MH358056 | MH358187 | – | – |
| Angiosperms353 | Urticaceae | *Chamabainia cuspidata* | Zhing-tao, W. | 870226 | K | – | – | – | – | – | – | – | – | ERS5501757 |
| Sanger | Urticaceae | *Chamabainia cuspidata* | WuZY | 10086 | KUN | China, Yunnan | – | – | KF137827 | KF138299 | KF138136 | KF138457 | KF137969 | – |
| Sanger | Urticaceae | *Coussapoa glaberrima* | Br | 2015 | KUN | Brazil | – | – | – | MH358268 | MH358060 | – | MH357987 | – |
| Sanger | Urticaceae | *Coussapoa parvifolia* | A.K. Monro et al. | 6833 | BM | Costa Rica | – | – | – | KF138301 | – | KF138459 | – | – |
| Angiosperms353 | Urticaceae | *Coussapoa villosa* | Pennington, T.D. | 10630 | K | – | – | – | – | – | – | – | – | ERS5501756 |

| Sanger | Urticaceae | *Cypholophus heterophyllus* |  | 6032 | NHN | Fiji | – | – | – | MH358269 | – | – | – | – |
| --- | --- | --- | --- | --- | --- | --- | --- | --- | --- | --- | --- | --- | --- | --- |
| Sanger | Urticaceae | *Cypholophus macrocephalus* |  |  | NHN | Vanuatu | – | – | MH357871 | MH358270 | MH358061 | MH358191 | MH357988 | – |
| Angiosperms353 | Urticaceae | *Cypholophus montanus* | Argent, G. | 535 | K | Indonesia | 23927 | – | – | – | – | – | – | ERS5501737 |
| Sanger | Urticaceae | *Cypholophus montanus* | L0942484 | L0942484 | NHN | Papua, Indonesia | – | – | MH357873 | – | MH358063 | – | – | – |
| Sanger | Urticaceae | *Cypholophus sp.* | L0406219 | L0406219 | NHN | Sulawesi, Indonesia | – | – | MH357874 | MH358273 | MH358065 | MH358192 | MH357991 | – |
| Sanger | Urticaceae | *Cypholophus sp. 1* | L0792369 | L0792369 | NHN | Irian Jaya, Indonesia | – | – | – | – | MH358064 | – | MH357990 | – |
| Sanger | Urticaceae | *Cypholophus sp. 2* | L0792372 | L0792372 | NHN | Bougainville, Papua New Guinea | – | – | – | – | MH358066 | – | – | – |
| Sanger | Urticaceae | *Debregeasia elliptica* | WuZY | 10061 | KUN | China, Yunnan | – | – | KF137830 | KF138303 | KF138139 | KF138461 | KF137972 | – |
| Angiosperms353 | Urticaceae | *Debregeasia longifolia* | Duaneh, J. | 91 | K | Malaysia | 23936 | – | – | – | – | – | – | ERS5501741 |
| Sanger | Urticaceae | *Debregeasia longifolia* | WuZY | 9471 | KUN | China, Guangxi | – | – | KF137831 | KF138304 | KF138140 | KF138462 | KF137973 | – |
| Sanger | Urticaceae | *Debregeasia orientalis* |  | 81482 | KUN | China, Xizang | – | – | KF137834 | KF138307 | KF138143 | KF138465 | KF137976 | – |
| Sanger | Urticaceae | *Debregeasia saeneb* |  | 81107 | KUN | China, Xizang | – | – | KF137835 | KF138308 | KF138144 | KF138466 | KF137977 | – |
| Sanger | Urticaceae | *Debregeasia squamata* | WuZY | 9204 | KUN | Yunnan, China | – | – | KF137837 | KF138310 | KF138146 | KF138468 | KF137979 | – |
| Sanger | Urticaceae | *Dendrocnide excelsa* |  |  | E | Australia | – | – | – | MH358274 | – | – | MH357992 | – |
| Sanger | Urticaceae | *Dendrocnide meyeniana* | Yi 20111184 | 20111184 | KUN | China | – | KF137838 | – | KF138311 | KF138147 | KF138469 | KF137980 | – |
| Sanger | Urticaceae | *Dendrocnide sinuata* | WuZY | 9238 | KUN | China | – | KF137754 | KF137839 | KF138312 | KF138148 | KF138470 | KF137981 | – |
| Sanger | Urticaceae | *Dendrocnide sp.* | WuZY | 9035 | KUN | China | – | – | KF137840 | KF138313 | KF138149 | KF138471 | KF137982 | – |
| Angiosperms353 | Urticaceae | *Dendrocnide stimulans* | Beaman, J.H. | 10369 | K | Malaysia | 23908 | – | – | – | – | – | – | ERS4414128 |
| Sanger | Urticaceae | *Dendrocnide urentissima* | WuZY | 9211 | KUN | China | – | – | KF137841 | KF138314 | KF138150 | KF138472 | – | – |
| Angiosperms353 | Urticaceae | *Didymodoxa caffra* | Friis, I.; Gilbert, M.G.; Vollesen, K. | 3532 | K | – | – | – | – | – | – | – | – | ERS5502184 |
| Sanger | Urticaceae | *Didymodoxa caffra* | Abdallah, R. et al. | 96/92 | K | Ethiopia? | – | KF137755 | – | KF138315 | KF138151 | KF138473 | – | – |
| Angiosperms353 | Urticaceae | *Discocnide mexicana* | Manriquez I., G.; Esquivel Hernadez, K.B. | 6554 | K | – | – | – | – | – | – | – | – | ERS5501982 |
| Sanger | Urticaceae | *Discocnide mexicana* | Gereau *et al* . | 2205 | BM | Mexico | – | – | KF137842 | DQ179369 | KF138152 | KF138474 | KF137983 | – |
| Sanger | Urticaceae | *Droguetia ambigua* | Styles & Styles | 2426 | K | South Africa | – | KF137756 | KF137843 | KF138317 | AM235161 | KF138475 | – | – |
| Angiosperms353 | Urticaceae | *Droguetia iners* | Poulsen, A.D.; et al. | 1000 | K | Uganda | 23947 | – | – | – | – | – | – | ERS5501749 |
| Sanger | Urticaceae | *Droguetia iners* | Liuj | 10621 | KUN | China? | – | KF137757 | KF137844 | KF138318 | KF138154 | KF138476 | KF137984 | – |
|  |  |  |  |  |  | Bagvio. Beugoerc Prov.Lozon.3/1913 |  |  |  |  |  |  |  |  |
| Sanger | Urticaceae | *Elatostema polypodioides* |  | 1119052 | BM |  | – | – | – | – | – | – | – | – |
| Sanger | Urticaceae | *Elatostema acuminatum* | FRI | 29035 | K | Malagsia 22th mile Ginting→Simpsh road | – | – | – | – | – | – | – | – |
| Sanger | Urticaceae | *Elatostema acuteserratum* | Y. H. Tseng | 1121 | TAI | Taiwan | – | – | KP858860 | – | – | – | – | – |
| Sanger | Urticaceae | *Elatostema agusanense* |  | 38445 | BM | Phillipines.Mr.Bulu san,Sorsogon  Prov,Luzon | – | – | – | – | – | – | – | – |
| Sanger | Urticaceae | *Elatostema albopilosoides* | Y. G. Wei & F. Wen | 1083 | IBK | China | – | – | KP858885 | – | – | – | – | – |
| Sanger | Urticaceae | *Elatostema androstachyum* | Y. G. Wei | g120 | IBK | China | – | – | KP858871 | – | – | – | – | – |
| Sanger | Urticaceae | *Elatostema angustum* |  | 13788 | BM | Phillipines Mt.Canumag,Rizal Prov.Luzon | – | – | To submit | – | – | – | – | – |
| Sanger | Urticaceae | *Elatostema angwesnm* |  | 5529 |  |  | – | – | – | – | – | – | – | – |
| Sanger | Urticaceae | *Elatostema apoeuse* |  | 11793 | BM | Phillipines.Lor? 6/7-1917 | – | – | – | – | – | – | – | – |

| Sanger | Urticaceae | *Elatostema asterocephalum* | Y. G. Wei | 623 | IBK | China | – | – | KP858878 | – | – | – | – | – |
| --- | --- | --- | --- | --- | --- | --- | --- | --- | --- | --- | --- | --- | --- | --- |
| Sanger | Urticaceae | *Elatostema atroviride* |  | 36 |  | Sanceng Caue,Qiaotou Village Jialiang Toum,Libo County,Guizhou province | – | – | – | – | – | – | – | – |
| Sanger | Urticaceae | *Elatostema attenuatoides* | Wei Y.G. | 76 | IBK | Dahuadi, Guangxi, China | – | – | – | – | To submit | – | To submit | – |
|  |  |  |  |  |  | Vietnam Cao Bang on lower Slqw lelow bamboo zow on raks |  |  |  |  |  |  |  |  |
| Sanger | Urticaceae | *Elatostema attenuatoides* |  | 4336 | K |  | – | – | To submit | – | – | – | – | – |
|  |  |  |  |  |  | Phillipines Cabadbaraw,Mr.Ur danera,Agosan Prov,Mindanao |  |  |  |  |  |  |  |  |
| Sanger | Urticaceae | *Elatostema auronii* |  | 13894 | BM |  | – | – | – | – | – | – | – | – |
| Sanger | Urticaceae | *Elatostema backeri* | HNK | 2724 |  | Chu Village,Ta Sua, Bac Yen Son La,Vietnam | – | – | – | – | – | – | – | – |
| Sanger | Urticaceae | *Elatostema balansae* | A. K. Monro & Y. G. Wei | 6466 | IBK | China | – | – | KP858847 | – | – | – | – | – |
| Sanger | Urticaceae | *Elatostema banahaense* | C. I. Peng | 23770 | HAST | Philippines | – | – | KP858815 | – | – | – | – | – |
| Sanger | Urticaceae | *Elatostema banahaense* |  | 3680 | K | Thailand. Khao Choug | – | – | – | – | – | – | – | – |
| Sanger | Urticaceae | *Elatostema belense* |  | 11317 | BM | Phillipines.Baraan Prov.Lozon.  Mr.Mariveles 3/1911 | – | – | To submit | – | – | – | – | – |
| Sanger | Urticaceae | *Elatostema beugoereuje* |  | 14274 | BM | New Guinea | – | – | To submit | – | – | – | – | – |
| Sanger | Urticaceae | *Elatostema binatum* |  | 092 | IBK | Banbi, Longzhou County, Guangxi, China | – | – | To submit | – | To submit | – | To submit | – |
| Sanger | Urticaceae | *Elatostema binatum* | A. K. Monro & Y. G. Wei | 6663 | IBK | China | – | – | KP858895 | – | – | – | – | – |
| Sanger | Urticaceae | *Elatostema blechnoides* |  | 6085 | BM | New Guinea.Westem Highlands,Distntv, nr Tomba Village,S.of Mr.Hagen range | – | – | To submit | – | – | – | – | – |
| Sanger | Urticaceae | *Elatostema brachyodontum* | A. K. Monro & Y. G. Wei | 6724 | IBK | China | – | – | KP858888 | – | – | – | – | – |
| Sanger | Urticaceae | *Elatostema bsungletense* |  | 14274 |  |  | – | – | To submit | – | – | – | – | – |
| Sanger | Urticaceae | *Elatostema bulbiferum* |  | 33967 | K | Thailand.Kanchana bun;Erawan | – | – | To submit | – | – | – | – | – |
| Sanger | Urticaceae | *Elatostema bullatum* |  | 10928 | K | Borneo.Sabah.Rana u District nt Posing Hor Spningsi | – | – | – | – | – | – | – | – |
| Sanger | Urticaceae | *Elatostema bwlloifetum* |  | 4315 |  |  | – | – | To submit | – | – | – | – | – |
| Sanger | Urticaceae | *Elatostema calcareum* | R. C. M. Gregor | 432 | BM | Mariana Islands | – | – | KP858817 | – | – | – | – | – |

| Sanger | Urticaceae | *Elatostema calciferum* |  | 83 |  | Yeji Tuo,Limi Town,Yongshun County Hunan province | – | – | – | – | – | – | – | – |
| --- | --- | --- | --- | --- | --- | --- | --- | --- | --- | --- | --- | --- | --- | --- |
| Sanger | Urticaceae | *Elatostema catarctum* |  | 2002131 |  | Jialiang,Libo county,Guizhou province | – | – | To submit | – | – | – | – | – |
| Sanger | Urticaceae | *Elatostema celingense* | FLF | 37 | IBK | Lihu, Nandan County, Guangxi, China | – | – | To submit | – | To submit | – | To submit | – |
| Sanger | Urticaceae | *Elatostema circulosum* |  | 5547 |  | Phillipines.Mr.Mali nao Albng Prov.Lozon. | – | – | – | – | – | – | – | – |
| Sanger | Urticaceae | *Elatostema clemeusii* |  | 20788 | K | Malagsia Sarawak,Kapir,Upp er,Nejang River | – | – | To submit | – | – | – | – | – |
| Sanger | Urticaceae | *Elatostema coriaceifolium* |  | 61 |  | Shuirao village,Libo county Guizhou province | – | – | To submit | – | – | – | – | – |
|  |  |  |  |  |  | Xiao Xi National Natural Reserve,Yongshun County.Hunan province |  |  |  |  |  |  |  |  |
| Sanger | Urticaceae | *Elatostema cuspidatum* |  | 72 |  |  | – | – | To submit | – | – | – | – | – |
| Sanger | Urticaceae | *Elatostema cwpidatum* |  | 8850312 |  | E.Nepal,Koshi Zone,Sankhuwa sabhaDistr,Around Tashi Gaun(Tashigaom)( 2160-2340m) | – | – | – | – | – | – | – | – |
| Sanger | Urticaceae | *Elatostema cyrtandrifolium* | Y. H. Tseng | 1139 | TAI | Taiwan | – | – | KP858844 | – | – | – | – | – |
| Sanger | Urticaceae | *Elatostema dactgiccephcwm* |  | 23678 |  | Gaoagong shan | – | – | To submit | – | – | – | – | – |
| Sanger | Urticaceae | *Elatostema delicarum* |  | 10343 | BM | Phillipines Todaga Mr.Apo Davao District Mindano 9/1909 | – | – | To submit | – | – | – | – | – |
|  |  |  |  |  |  | The 4th Group of Shuichun Village Yuping Town,Libo  County,Guizhou |  |  |  |  |  |  |  |  |
| Sanger | Urticaceae | *Elatostema discolor* |  | 59 |  |  | – | – | To submit | – | – | – | – | – |
| Sanger | Urticaceae | *Elatostema dissectum* | A. J. C. Grierson & D. G. Long | 393 | K | Bhutan | – | – | KP858832 | – | – | – | – | – |
| Sanger | Urticaceae | *Elatostema edule* | Y. H. Tseng | 1119 | TAI | Taiwan | – | – | KP858818 | – | – | – | – | – |
| Sanger | Urticaceae | *Elatostema ellipticum* | FKW | 7951 | K | India | – | – | – | – | – | – | – | – |
| Sanger | Urticaceae | *Elatostema fengshanense* | A. K. Monro & Y. G. Wei | 6651 | IBK | China | – | – | KP858848 | – | – | – | – | – |
| Sanger | Urticaceae | *Elatostema ficoides* |  | 1949 |  | Central Nepal No.1 west | – | – | To submit | – | – | – | – | – |

| Sanger | Urticaceae | *Elatostema filicinum* |  | 0006249871 | BM | New Euinea,Camp  I, | – | – | – | – | – | – | – | – |
| --- | --- | --- | --- | --- | --- | --- | --- | --- | --- | --- | --- | --- | --- | --- |
| Sanger | Urticaceae | *Elatostema garrettii* | P. Suvarnakoses | 38824 | K | Thailand | – | – | KP858849 | – | – | – | – | – |
| Sanger | Urticaceae | *Elatostema gibbsiae* |  | 11312 | K | Sabah.Ranau Disrncr. Mr Kinabalo.Liwago river vallege1700- 1800m. | – | – | – | – | – | – | – | – |
| Sanger | Urticaceae | *Elatostema glabrutum* |  | 16325 | BM | Phillipines.Mr.Bulu san,Sorsogon Prov,Luzon | – | – | To submit | – | – | – | – | – |
| Sanger | Urticaceae | *Elatostema glaucescens* |  | 18397 | BM | Phillipines.Mr.Polis | – | – | – | – | – | – | – | – |
| Sanger | Urticaceae | *Elatostema glochidiodes* |  | 2002121 |  | YongKang,Libo County,Guizhou province Lin-Dong | – | – | To submit | – | – | – | – | – |
| Sanger | Urticaceae | *Elatostema goudotianum* | S. Malcomber | 1723 | BM | Madagascar | – | – | KP858798 | – | – | – | – | – |
| Sanger | Urticaceae | *Elatostema grande* | I. R. Telfor | 10363 | K | Australia | – | – | KP858821 | – | – | – | – | – |
| Sanger | Urticaceae | *Elatostema grandidentatum* | A. J. C. Grierson & D. G. Long | 1999 | K | Bhutan | – | – | KP858833 | – | – | – | – | – |
| Sanger | Urticaceae | *Elatostema grandifolium* | K. L. Rechinger | 1780 | BM | Samoa | – | – | KP858823 | – | – | – | – | – |
| Sanger | Urticaceae | *Elatostema gyrocephalum* | Y. G. Wei | 7101 | IBK | China | – | – | KP858886 | – | – | – | – | – |
| Sanger | Urticaceae | *Elatostema hechiense* | A. K. Monro & Y. G. Wei | 6507 | IBK | China | – | – | KP858875 | – | – | – | – | – |
| Sanger | Urticaceae | *Elatostema hezhouense* | A. K. Monro & Y. G. Wei | 6820 | IBK | China | – | – | KP858881 | – | – | – | – | – |
| Sanger | Urticaceae | *Elatostema hirtellipedunculatum* | Y. H. Tseng | 1123 | TAI | Taiwan | – | – | KP858863 | – | – | – | – | – |
| Sanger | Urticaceae | *Elatostema holophgllum* |  | 23546 | BM | Sorsogou Pror,Luzon,Phillipi nes | – | – | To submit | – | – | – | – | – |
| Sanger | Urticaceae | *Elatostema hookerianum* | Y. G. Wei | 7 | IBK | China | – | – | KP858904 | – | – | – | – | – |
| Sanger | Urticaceae | *Elatostema hypoglaucum* | Y. H. Tseng | 1190 | TAI | Taiwan | – | – | KP858891 | – | – | – | – | – |
| Sanger | Urticaceae | *Elatostema ichangense* |  | 67 |  | Aojiahu Village,Lmoyixi Town Guzhang County,Hunan province | – | – | To submit | – | – | – | – | – |
| Sanger | Urticaceae | *Elatostema incisum* | G. E. Schatx et al. | 3459 | MO | Madagascar | – | – | KP858799 | – | – | – | – | – |
| Sanger | Urticaceae | *Elatostema inregnfolium* |  | 1359 | K | New Euinea.Morobe District.Aseki patrol area near Wengomanga(via Oiwa) | – | – | To submit | – | – | – | – | – |
| Sanger | Urticaceae | *Elatostema insulare* | A. C. Smith | 8145 | K | Fiji | – | – | KP858839 | – | – | – | – | – |
| Sanger | Urticaceae | *Elatostema integrifolium* | E. S. Brown | 1484 | BM | Solomon Islands | – | – | KP858852 | – | – | – | – | – |
| Sanger | Urticaceae | *Elatostema involucratum* | Tsugaru | 30727 | A | Japan | – | – | KP858866 | – | – | – | – | – |
| Sanger | Urticaceae | *Elatostema japonicum* | G. Murata | 72870 | KYO | Japan | – | – | KP858867 | – | – | – | – | – |

| Sanger | Urticaceae | *Elatostema jilicavle* |  | 7627 | BM | Phillipines. Dumagvere. Cuernos MH.Negros Oriemral Prov. Negror Island. 6/1908 | – | – | – | – | – | – | – | – |
| --- | --- | --- | --- | --- | --- | --- | --- | --- | --- | --- | --- | --- | --- | --- |
|  |  |  |  |  |  | Malagsia.Sabah,Kot a Belud District S of Sagap on NW  side of Mt.Kinabalu |  |  |  |  |  |  |  |  |
| Sanger | Urticaceae | *Elatostema kabagense* |  | 9783 | K |  | – | – | To submit | – | – | – | – | – |
| Sanger | Urticaceae | *Elatostema kraemeri* | K. L. Rechinger | 1548 | BM | Samoa | – | – | KP858824 | – | – | – | – | – |
| Sanger | Urticaceae | *Elatostema laetevirens* | S. Tsugarul & T. Takahashi | 25903 | MO | Japan | – | – | KP858893 | – | – | – | – | – |
| Sanger | Urticaceae | *Elatostema laevissimum* | Y. G. Wei | 6393 | IBK | China | – | – | KP858861 | – | – | – | – | – |
| Sanger | Urticaceae | *Elatostema lagunense* |  | 17529 | BM | Phillipines.Los Baüos.Mr.Maguilin g,Laguna Prov,Luzon | – | – | To submit | – | – | – | – | – |
|  |  |  |  |  |  | Indonena,W Sumbawa,Mr.Barula nteh,trailfrom Batuctulang Pusu |  |  |  |  |  |  |  |  |
| Sanger | Urticaceae | *Elatostema lanafohum* |  | 2014 | BM |  | – | – | – | – | – | – | – | – |
|  |  |  |  |  |  | Indonesia.W.Sumba wa,Mr Batulantehtrail fnom Batudulang to Pwsu |  |  |  |  |  |  |  |  |
| Sanger | Urticaceae | *Elatostema lancifolium* |  | 4678 | K |  | – | – | – | – | – | – | – | – |
|  |  |  |  |  |  | Malagsia Sarawak Eat Upper Rejang river Bcre of Mr.Majan low elevnhvn |  |  |  |  |  |  |  |  |
| Sanger | Urticaceae | *Elatostema laxiflorum* |  | 20790 | K |  | – | – | – | – | – | – | – | – |
| Sanger | Urticaceae | *Elatostema lineare* | J & M. S. Clemens | 50589 | BM | Malaysia | – | – | KP858835 | – | – | – | – | – |
| Sanger | Urticaceae | *Elatostema lineolatum* | Y. H. Tseng | 1108 | TAI | Taiwan | – | – | KP858855 | – | – | – | – | – |
| Sanger | Urticaceae | *Elatostema lithoneurum* | J & M. S. Clemens | s.n. | BM | Malaysia | – | – | KP858827 | – | – | – | – | – |
|  |  |  |  |  |  | Cao Bang,Tra Tinh District Quoc Toan Municipality;Trang Hem(Lakes  District) |  |  |  |  |  |  |  |  |
| Sanger | Urticaceae | *Elatostema longibracteatum* |  | 4288 | K |  | – | – | To submit | – | – | – | – | – |
| Sanger | Urticaceae | *Elatostema longistipulum* | A. K. Monro & Y. G. Wei | 6462 | IBK | China | – | – | KP858856 | – | – | – | – | – |
| Sanger | Urticaceae | *Elatostema lungzhouense* | A. K. Monro & Y. G. Wei | 6786 | IBK | China | – | – | KP858868 | – | – | – | – | – |
| Sanger | Urticaceae | *Elatostema lutescens* | C. B. Robinson | 14004 | BM | Philippines | – | – | KP858816 | – | – | – | – | – |
| Sanger | Urticaceae | *Elatostema macintyrei* | A. K. Monro & Y. G. Wei | 6743 | IBK | China | – | – | KP858850 | – | – | – | – | – |

| Sanger | Urticaceae | *Elatostema macnphllium* |  | 001119057 |  | New Euinea,A.F.R.Wolla sta Expeditio Camp V16 | – | – | To submit | – | – | – | – | – |
| --- | --- | --- | --- | --- | --- | --- | --- | --- | --- | --- | --- | --- | --- | --- |
| Sanger | Urticaceae | *Elatostema madagascariense* | Richard Razakamalala et al. | 3199 | MO | Africa | – | – | KP858800 | – | – | – | – | – |
| Sanger | Urticaceae | *Elatostema malacotrichum* | Y. G. Wei | 13 | IBK | China | – | – | KP858876 | – | – | – | – | – |
| Sanger | Urticaceae | *Elatostema megacephalum* |  | 1368 | K | Chingmai,Doi Ka | – | – | To submit | – | – | – | – | – |
| Sanger | Urticaceae | *Elatostema microcephalanthum* | Y. H. Tseng | 1100 | TAI | Taiwan | – | – | KP858874 | – | – | – | – | – |
| Sanger | Urticaceae | *Elatostema molle* |  | 262 | K | Peninsular:Narathi wat Bacho | – | – | – | – | – | – | – | – |
| Sanger | Urticaceae | *Elatostema monandrum* | F. Miyamoto et al. | 9588281 | MO | Nepal | – | – | KP858905 | – | – | – | – | – |
| Sanger | Urticaceae | *Elatostema monticola* | H. A. Osmaston | 1952 | BM | Congo | – | – | KP858803 | – | – | – | – | – |
| Sanger | Urticaceae | *Elatostema morobense* | R. J. Johns | 9209A | K | Indonesia | – | – | KP858820 | – | – | – | – | – |
| Sanger | Urticaceae | *Elatostema multicaule* | A. K. Monro & Y. G. Wei | 6748 | IBK | China | – | – | KP858884 | – | – | – | – | – |
| Sanger | Urticaceae | *Elatostema myrtillus* | Y. G. Wei | 44937 | IBK | China | – | – | KP858880 | – | – | – | – | – |
| Sanger | Urticaceae | *Elatostema nacuturn* |  | 9338 |  | Aalung,Guizhou Y.Tsiang | – | – | To submit | – | – | – | – | – |
| Sanger | Urticaceae | *Elatostema nalkerae* |  | 20916 | K | Laos.Tawrieng,Che ng Kwang | – | – | To submit | – | – | – | – | – |
|  |  |  |  |  |  | Baibi Cave,Qiaotoy Village Jialiang Town,Libo County Guizhou province |  |  |  |  |  |  |  |  |
| Sanger | Urticaceae | *Elatostema nanchuanense* |  | 38 |  |  | – | – | To submit | – | – | – | – | – |
| Sanger | Urticaceae | *Elatostema nasutum* | A. K. Monro & Y. G. Wei | 6401 | IBK | China | – | – | KP858902 | – | – | – | – | – |
| Sanger | Urticaceae | *Elatostema nrangii* |  | 775 |  |  | – | – | To submit | – | – | – | – | – |
| Sanger | Urticaceae | *Elatostema oblongifolium* | A. K. Monro & Y. G. Wei | 6713 | IBK | China | – | – | KP858897 | – | – | – | – | – |
| Sanger | Urticaceae | *Elatostema obtusum* | Y. H. Tseng | 1191 | TAI | Taiwan | – | – | KP858900 | – | – | – | – | – |
| Sanger | Urticaceae | *Elatostema obtusum* | Bouffordet al | al.27415 | A | China | – | – | KP858901 | – | – | – | – | – |
|  |  |  |  |  |  | Indonesia.Sumarm, Dëlëag Singkoet,N of Bërasvagl,Karo Plareau |  |  |  |  |  |  |  |  |
| Sanger | Urticaceae | *Elatostema oglvnnum* |  | 8571 | K |  | – | – | – | – | – | – | – | – |
|  |  |  |  |  |  | Phillipines Luzon.Sorrogon Mr Bulusan about lake Agangag on stores |  |  |  |  |  |  |  |  |
| Sanger | Urticaceae | *Elatostema oimulans* |  | 9654 | K |  | – | – | – | – | – | – | – | – |
| Sanger | Urticaceae | *Elatostema oligophlebium* | Y. G. Wei | g075 | IBK | China | – | – | KP858877 | – | – | – | – | – |
| Sanger | Urticaceae | *Elatostema orienlau* |  | 17156 |  |  | – | – | – | – | – | – | – | – |
| Sanger | Urticaceae | *Elatostema orientale* | C. F. M. Synnerton | 784 | BM | Zimbabwe | – | – | KP858802 | – | – | – | – | – |
| Sanger | Urticaceae | *Elatostema paivaeanum* | Charles Doumenge | 456 | MO | Africa | – | – | KP858801 | – | – | – | – | – |
| Sanger | Urticaceae | *Elatostema paracuminatw* |  | 21016 | K | pu Bia Laos | – | – | – | – | – | – | – | – |

|  |  |  |  |  |  | Baibi care,Qiaotou Village,Jialiang Town,Libo county,Guizhou  province |  |  |  |  |  |  |  |  |
| --- | --- | --- | --- | --- | --- | --- | --- | --- | --- | --- | --- | --- | --- | --- |
| Sanger | Urticaceae | *Elatostema parvum* |  | 41 |  |  | – | – | To submit | – | – | – | – | – |
| Sanger | Urticaceae | *Elatostema parvum* | Y. H. Tseng | 1182 | TAI | Taiwan | – | – | KP858794 | – | – | – | – | – |
| Sanger | Urticaceae | *Elatostema pedicillatum* |  | 31355 | BM | Borneo Mr Kinabalu | – | – | To submit | – | – | – | – | – |
|  |  |  |  |  |  | New Euinea.Indouesra. Mimika Regeng,PF Freenort Indonessa Concesne area:main road 34 mile W side of Aikwa bridge |  |  |  |  |  |  |  |  |
| Sanger | Urticaceae | *Elatostema peltijolium* |  | 290 | K |  | – | – | To submit | – | – | – | – | – |
| Sanger | Urticaceae | *Elatostema penibukanense* | J. H. Beaman | 10466 | K | Malaysia | – | – | KP858854 | – | – | – | – | – |
| Sanger | Urticaceae | *Elatostema peniwkanense* |  | 29737 | BM | Borneo Mr Kinabalu ilau Basin | – | – | – | – | – | – | – | – |
| Sanger | Urticaceae | *Elatostema pennineive* |  | 2430 | K | Biunei Temburong. Batu Apoi;Bukir Galagas | – | – | – | – | – | – | – | – |
|  |  |  |  |  |  | Lang Son Húu Lung District Húu Lien Municipa city Húu Lien proctected Area between Peo Dat |  |  |  |  |  |  |  |  |
| Sanger | Urticaceae | *Elatostema pergameneom* |  | 4121 | K |  | – | – | – | – | – | – | – | – |
| Sanger | Urticaceae | *Elatostema pergameneum* |  | 07298 | IBK | Sanlian, Longzhou County, Guangxi, China | – | – | To submit | – | To submit | – | To submit | – |
| Sanger | Urticaceae | *Elatostema perlongfolium* |  | 16408 | BM | Phillipines Irobin.Mf.Bulasan.S orsogon Prov.Luzon | – | – | – | – | – | – | – | – |
| Sanger | Urticaceae | *Elatostema phillipinense* |  | 6907 | K | Phillipines.Negros. Canlaon Volcauo | – | – | To submit | – | – | – | – | – |
| Sanger | Urticaceae | *Elatostema pianmaense* |  | 24077 |  | Gaoagong shan | – | – | To submit | – | – | – | – | – |
| Sanger | Urticaceae | *Elatostema pinnatum* | J & M. S. Clemens | s.n. | BM | Malaysia | – | – | KP858836 | – | – | – | – | – |
| Sanger | Urticaceae | *Elatostema platyphyllum* | Y. H. Tseng | 1158 | TAI | Taiwan | – | – | KC420490 | – | – | – | – | – |

| Sanger | Urticaceae | *Elatostema podohglum* |  | 19714 | BM | Phillipines  ,Mr.Polos Ifugo Sub prov,Luzoni | – | – | To submit | – | – | – | – | – |
| --- | --- | --- | --- | --- | --- | --- | --- | --- | --- | --- | --- | --- | --- | --- |
|  |  |  |  |  |  | Huangcao ping,Yaolu Town Libo County,Guizhou |  |  |  |  |  |  |  |  |
| Sanger | Urticaceae | *Elatostema pseudissectum* |  | 28 |  |  | – | – | To submit | – | – | – | – | – |
| Sanger | Urticaceae | *Elatostema pusillum* | R. Bedi | 931 | K | Bhutan | – | – | KP858906 | – | – | – | – | – |
| Sanger | Urticaceae | *Elatostema pycndontum* |  | 39 |  | W.T.Wang Baibi Cave Qiaotou Village.Jialiang Town,Libo County.Guizhou province | – | – | To submit | – | – | – | – | – |
| Sanger | Urticaceae | *Elatostema ramosum* | AM | 6800 | IBK | Banbi, Longzhou County, Guangxi, China | – | – | To submit | – | To submit | – | To submit | – |
| Sanger | Urticaceae | *Elatostema reticulatum* | P. I. Forster | 7564 | K | Australia | – | – | KP858834 | – | – | – | – | – |
| Angiosperms353 | Urticaceae | *Elatostema retrohirtum* | Monro, A.K.; et al. | 7601 | K | – | – | – | – | – | – | – | – | ERS4414235 |
| Sanger | Urticaceae | *Elatostema retrohirtum* |  | 236 |  | Doi Chuangdao,Siam,Ta iland 1921.1.4 1500 1800m | – | – | To submit | – | – | – | – | – |
| Sanger | Urticaceae | *Elatostema rubrosvipulatum* |  | 11322 | K | Sabah Nanav Dishnr,Mr Kinabalu Liwago Riov | – | – | – | – | – | – | – | – |
| Sanger | Urticaceae | *Elatostema rugosum* | Hooker | s.n. | BM | New Zealand | – | – | – | – | – | – | – | – |
| Sanger | Urticaceae | *Elatostema rupestre* |  | 245 |  | Chuia Chuli | – | – | – | – | – | – | – | – |
| Sanger | Urticaceae | *Elatostema salvinioides* |  | 4215 | K | Northern Thailand Chiangmai Doi chingdao | – | – | – | – | – | – | – | – |
| Sanger | Urticaceae | *Elatostema samoense* | K. L. Rechinger | s.n. | BM | Samoa | – | – | KP858826 | – | – | – | – | – |
|  |  |  |  |  |  | New Euinea.Morobe Distncr 5m Sw of Wagau Mr.Shungol |  |  |  |  |  |  |  |  |
| Sanger | Urticaceae | *Elatostema schroeteri* |  | 12474 | K |  | – | – | – | – | – | – | – | – |
| Sanger | Urticaceae | *Elatostema serra* | W. Takeuchi et al. | 19898 | MO | Papua New Guinea | – | – | KP858837 | – | – | – | – | – |
| Sanger | Urticaceae | *Elatostema sessile* |  |  | K | Indowesra.Jara | – | – | To submit | – | – | – | – | – |
| Sanger | Urticaceae | *Elatostema setulosum* | Y. G. Wei | g124 | IBK | China | – | – | KP858872 | – | – | – | – | – |
| Sanger | Urticaceae | *Elatostema sinense* | Q. Shao & L. D. Duan | 76 | BM | China | – | – | KP858795 | – | – | – | – | – |
| Sanger | Urticaceae | *Elatostema sinense* | Q. Shao & L. D. Duan | 1 | BM, PE | China | – | – | KP858796 | – | – | – | – | – |
| Sanger | Urticaceae | *Elatostema sinense* | T. T. Chen | s.n. | ? | China | – | – | KP858797 | – | – | – | – | – |
| Sanger | Urticaceae | *Elatostema sinopurpureum* | Y. G. Wei & F. Wen | 1081 | IBK | China | – | – | KP858869 | – | – | – | – | – |

|  |  |  |  |  |  | Phillipines.Irosin Mt.Bulusan,Sorsog  on Prov,Luzon |  |  |  |  |  |  |  |  |
| --- | --- | --- | --- | --- | --- | --- | --- | --- | --- | --- | --- | --- | --- | --- |
| Sanger | Urticaceae | *Elatostema sorsogoncnse* |  | 16322 | BM |  | – | – | To submit | – | – | – | – | – |
| Sanger | Urticaceae | *Elatostema sp.* |  | 143234 | K | Phillipines Luzon.Laguna Pioviuce,Mr. Makiling | – | – | – | – | – | – | – | – |
| Sanger | Urticaceae | *Elatostema sterma* |  | 959 |  | Hendcian, | – | – | To submit | – | – | – | – | – |
|  |  |  |  |  |  | Xiao Xi National Natural Reserve,Yongshun County.Hunan province |  |  |  |  |  |  |  |  |
| Sanger | Urticaceae | *Elatostema stewardii* |  | 77 |  |  | – | – | To submit | – | – | – | – | – |
| Sanger | Urticaceae | *Elatostema strictum* | K. L. Rechinger | 388 | BM | Samoa | – | – | KP858825 | – | – | – | – | – |
| Sanger | Urticaceae | *Elatostema strigillosum* | C. I. Peng | 23775 | HAST | Philippines | – | – | KP858814 | – | – | – | – | – |
| Sanger | Urticaceae | *Elatostema subcoriaceum* | Y. H. Tseng | 1127 | TAI | Taiwan | – | – | KP858894 | – | – | – | – | – |
| Sanger | Urticaceae | *Elatostema sublineare* | A. K. Monro & Y. G. Wei | 6686 | IBK | China | – | – | KP858882 | – | – | – | – | – |
| Sanger | Urticaceae | *Elatostema surigaeense* |  | 9857 | BM | Phillipines  Mt.Banajao,Laguna Prov,Luzon | – | – | To submit | – | – | – | – | – |
| Sanger | Urticaceae | *Elatostema suzukii* | Yasuda | 5170 | KYO | Japan | – | – | KP858865 | – | – | – | – | – |
| Sanger | Urticaceae | *Elatostema tenuicaudatum* | Y. G. Wei & F. Wen | 1050 | IBK | China | – | – | KP858862 | – | – | – | – | – |
| Sanger | Urticaceae | *Elatostema tenuinerve* | A. K. Monro & Y. G. Wei | 6502 | IBK | China | – | – | KP858896 | – | – | – | – | – |
| Sanger | Urticaceae | *Elatostema thalictroides* | J. H. Beaman | 7650 | MO | Malaysia | – | – | KP858838 | – | – | – | – | – |
| Sanger | Urticaceae | *Elatostema tianeense* | A. K. Monro & Y. G. Wei | 6491 | IBK | China | – | – | KP858879 | – | – | – | – | – |
|  |  |  |  |  |  | Papua New Euinea W.Highlands Prov,Bismarck  range,Mt.Oibo |  |  |  |  |  |  |  |  |
| Sanger | Urticaceae | *Elatostema tndens* |  | 10475 | K |  | – | – | – | – | – | – | – | – |
| Sanger | Urticaceae | *Elatostema toidesaustrale* | A. C. Smith | 8131 | K | Fiji | – | – | KP858790 | – | – | – | – | – |
| Sanger | Urticaceae | *Elatostema toidesfilicoides* | H. B. R. Parham | 235a | BM | Fiji | – | – | KP858789 | – | – | – | – | – |
| Sanger | Urticaceae | *Elatostema toidesfruticulosum* | L. S. Brass | 2607 | BM | Solomon Island | – | – | KP858788 | – | – | – | – | – |
| Sanger | Urticaceae | *Elatostema toideslonchophyllum* | J. & M. S. Clemens | 20816 | K | Malaysia | – | – | KP858793 | – | – | – | – | – |
| Sanger | Urticaceae | *Elatostema toidesvariolaminosum var. latum* | Rena George | S38983 | K | Malaysia | – | – | KP858791 | – | – | – | – | – |
| Sanger | Urticaceae | *Elatostema villosum* | Y. H. Tseng | 1275 | TAI | Taiwan | – | – | KP858812 | – | – | – | – | – |
| Sanger | Urticaceae | *Elatostema viridescens* |  | 15371 | BM | Phillipines.Irosin.Pr ov.Sorsogon.Mr.Bul usan.Luzon Island | – | – | To submit | – | – | – | – | – |
| Sanger | Urticaceae | *Elatostema vittatum* | J. H. Beaman | 10976 | K | Malaysia | – | – | KP858792 | – | – | – | – | – |
| Sanger | Urticaceae | *Elatostema welwitschii* | Welwitsch | 6270 | BM | Angola | – | – | KP858841 | – | – | – | – | – |
| Sanger | Urticaceae | *Elatostema xanthophyllum* | Y. G. Wei & V.T. Do | VMN_CN2 4 | IBK | Vietnam | – | – | KP858890 | – | – | – | – | – |
| Sanger | Urticaceae | *Elatostema yachense* | A. K. Monro & Y. G. Wei | 6725 | IBK | China | – | – | KP858883 | – | – | – | – | – |
| Sanger | Urticaceae | *Elatostema yakushimense* | Murata et al. | 15490 | A | Japan | – | – | KP858892 | – | – | – | – | – |
| Sanger | Urticaceae | *Elatostema yaoshanense* | F. Wen | 070509A | IBK | China | – | – | KP858903 | – | – | – | – | – |
| Angiosperms353 | Urticaceae | *Elatostematoides vittatum* | Beaman, J.H. | 10976 | K | – | – | – | – | – | – | – | – | ERS5503005 |
| Angiosperms353 / Sanger | Urticaceae | *Forsskaolea angustifolia* | Chase, M.W. | 16132 | K | Spain | 15957 | KF137762 | KF137861 | KF138334 | KF138170 | KF138493 | – | ERS5501729 |

| Sanger | Urticaceae | *Forsskaolea candida* | unknown collector | 3193 | US | Namibia | – | – | KM586475 | KM586647 | KM586561 | MK955181 | – | – |
| --- | --- | --- | --- | --- | --- | --- | --- | --- | --- | --- | --- | --- | --- | --- |
| Angiosperms353 | Urticaceae | *Gesnouinia arborea* | Wildpret, W.; et al. | s.n. | K | Spain | 23944 | – | – | – | – | – | – | ERS5501746 |
| Sanger | Urticaceae | *Gesnouinia arborea* | C. Evrard | 12088 | BM | Spain, Canary Islands | – | – | KF137862 | DQ179372 | KF138172 | KF138494 | KF138000 | – |
| Angiosperms353 | Urticaceae | *Gibbsia insignis* | Sands, M.J.S. | 7149 | K | Indonesia | 23931 | – | – | – | – | – | – | ERS5501738 |
| Angiosperms353 | Urticaceae | *Girardinia diversifolia* | Abdallah, R.; et al. | 9689 | K | Tanzania, the United Republic of | 23917 | – | – | – | – | – | – | ERS5501731 |
| Sanger | Urticaceae | *Girardinia diversifolia subsp. diversifolia* | Liuj | 10776 | KUN | China | – | KF137763 | KF137863 | KF138337 | KF138174 | KF138496 | KF138001 | – |
| Sanger | Urticaceae | *Girardinia diversifolia subsp. suborbiculata* | WuZY | 10011 | KUN | China | – | – | – | KF138338 | KF138175 | KF138497 | KF138003 | – |
| Sanger | Urticaceae | *Girardinia diversifolia subsp. triloba* | WuZY | 10229 | KUN | China | – | – | – | KF138341 | KF138178 | KF138500 | KF138006 | – |
| Angiosperms353 | Urticaceae | *Gonostegia hirta* | Monro, A.K.; Fu, L.F. | 7504 | K | – | – | – | – | – | – | – | – | ERS5502208 |
| Sanger | Urticaceae | *Gonostegia hirta* | WuZY | 9478 | KUN | Guangxi, China | – | – | KF137865 | KF138342 | KF138179 | – | KF138007 | – |
| Sanger | Urticaceae | *Gonostegia parvifolia* | Liuj | 10649 | KUN | Fujian, China | – | – | KF137867 | KF138344 | KF138181 | KF138502 | KF138009 | – |
| Angiosperms353 | Urticaceae | *Gyrotaenia myriocarpa* | Ekman, E.L. | 7356 | K | Haiti | 23915 | – | – | – | – | – | – | ERS5501730 |
| Sanger | Urticaceae | *Gyrotaenia myriocarpa* | P. AcevedoRdgz. | 12971 | US | Dominican  Republic | – | – | KM586472 | KM586644 | KM586558 | – | – | – |
| Angiosperms353 | Urticaceae | *Haroldiella rapaensis* | Perlman | 18115 | K | – | – | – | – | – | – | – | – | ERS5503008 |
| Sanger | Urticaceae | *Hemistylus boehmerioides* | Schimpff | 445 | G | Ecuador | – | – | – | – | – | – | – | – |
| Angiosperms353 / Sanger | Urticaceae | *Hemistylus macrostachya* | Pittier | 11788 | K | Venezuela (Bolivarian  Republic of) | 23946 | – | KF137868 | KF138346 | KF138182 | KF138503 | – | ERS5501748 |
| Sanger | Urticaceae | *Hesperocnide sandwicensis* | J.B. Gillett | 16765 | EA | Kenya | – | – | KM586429 | KM586601 | KM586515 | MK955193 | MK931209 | – |
| Angiosperms353 | Urticaceae | *Hesperocnide tenella* | Nuttall, L.W. | 227 | K | United States of America | 23920 | – | – | – | – | – | – | ERS5501733 |
| Sanger | Urticaceae | *Hesperocnide tenella* | B. Trusk | 188 | EA | USA | – | – | KC284967 | KC285019 | KC284993 | – | – | – |
| Sanger | Urticaceae | *Laportea aestuans* | Peterson | 7165 | US | Panama | – | – | KM586464 | KM586636 | – | – | – | – |
| Angiosperms353 | Urticaceae | *Laportea canadensis* | Bourdo Jr., E.A. | 29633 | K | – | – | – | – | – | – | – | – | ERS4414133 |
| Angiosperms353 | Urticaceae | *Laportea cuspidata* | Furuse, M. | 18489 | K | – | – | – | – | – | – | – | – | ERS4414135 |
| Angiosperms353 | Urticaceae | *Laportea floribunda* | Lowry II, P.P. | 4529 | K | – | – | – | – | – | – | – | – | ERS5503068 |
| Angiosperms353 | Urticaceae | *Laportea humblotii* | Schatz, G.E.; Rakotozafy, A.; D'Arcy W.; Randrianasolo, J. | 1510 | K | – | – | – | – | – | – | – | – | ERS5503069 |
| Sanger | Urticaceae | *Laportea interrupta* | Festo & Luke | 2502 | EA | Kenya | – | – | KM586445 | KM586617 | – | – | – | – |
| Sanger | Urticaceae | *Laportea peduncularis* | Weigend | 8713 | BSB | South Africa | – | – | KF559047 | KF558927 | – | – | – | – |
| Angiosperms353 | Urticaceae | *Laportea ruderalis* | Fosberg, F.R. | 55756 | K | – | – | – | – | – | – | – | – | ERR3672054 |
| Sanger | Urticaceae | *Laportea ruderalis* | Hunt | 6 | US | Micronesia | – | – | KM586461 | KM586633 | – | – | – | – |
| Angiosperms353 | Urticaceae | *Lecanthus peduncularis* | Johns, R.J. | 10035 | K | Indonesia | 23921 | – | – | – | – | – | – | ERS5501734 |
| Sanger | Urticaceae | *Lecanthus peduncularis* | Liuj | 10607 | KUN | China | – | KF137768 | KF137871 | KF138350 | KF138186 | KF138508 | KF138016 | – |
| Sanger | Urticaceae | *Lecanthus_petelotii* | Anon | 81066 | KUN | China | – | – | KF137873 | – | – | – | – | – |
| Sanger | Urticaceae | *Leucosyke australis* | J.I. Wheatley | 864 | K | Vanuatu | – | – | – | – | – | – | – | – |
| Angiosperms353 | Urticaceae | *Leucosyke capitellata* | Lugas, L. | 41 | K | Malaysia | 23930 | – | – | – | – | – | – | ERS4414208 |
| Sanger | Urticaceae | *Leucosyke quadrinervia* | WuZY | 2012438 | KUN | Taiwan | – | – | KF137875 | KF138354 | KF138190 | – | – | – |
| Angiosperms353 | Urticaceae | *Maoutia puya* | Eanghourt, Khou et al. | FFI-PA203 | K | Cambodia | 23932 | – | – | – | – | – | – | ERS5503002 |
| Sanger | Urticaceae | *Maoutia setosa* | WuZY | 2012453 | KUN | Taiwan | – | – | MH357903 | MH358290 | MH358097 | MH358211 | – | – |
| Angiosperms353 | Urticaceae | *Musanga cecropioides* | Maurin, O. | 4374 | K | – | 74208 | – | – | – | – | – | – | ERS5501755 |
| Sanger | Urticaceae | *Musanga cecropioides* | I. Friis et al. | 9427 | K | Ethiopia | – | – | – | – | – | – | – | – |
| Sanger | Urticaceae | *Myrianthus holstii* | D.B. Faushawe | 5150 | K | Zimbabwe | – | – | – | – | – | – | – | – |
| Angiosperms353 | Urticaceae | *Myrianthus preussii* | Bogner | 672 | K | Gabon | 23953 | – | – | – | – | – | – | ERS5501751 |
| Sanger | Urticaceae | *Myriocarpa cordifolia* | Monro 4630 | 4630 | BM | Panama | – | KF137770 | KF137877 | KF138357 | KF138193 | KF138512 | – | – |
| Angiosperms353 / Sanger | Urticaceae | *Myriocarpa obovata* | Monro & Penn | 6534 | K | Belize | – | KF137771 | KF137878 | KF138358 | – | KF138513 | KF138021 | ERS5502207 |
| Angiosperms353 | Urticaceae | *Nanocnide japonica* | Im, H.T.; et al. | 6680 | K | Japan | 23919 | – | – | – | – | – | – | ERS5501732 |

| Sanger | Urticaceae | *Nanocnide japonica* | Liuj | 10735 | KUN | China | – | KF137772 | KF137879 | KF138359 | KF138194 | KF138514 | KF138022 | – |
| --- | --- | --- | --- | --- | --- | --- | --- | --- | --- | --- | --- | --- | --- | --- |
| Sanger | Urticaceae | *Nanocnide lobata* | Liuj | 10799 | KUN | China | – | KF137773 | KF137881 | KF138361 | KF138196 | KF138516 | KF138024 | – |
| Sanger | Urticaceae | *Neodistemon indicus* | Larsen et al. | 2669 | K | Thailand | – | – | – | KF138363 | KF138198 | – | KF138026 | – |
| Angiosperms353 | Urticaceae | *Neraudia angulata* | Cowan, R.S. | 758 | K | United States of America | 23940 | – | – | – | – | – | – | ERS5501744 |
| Sanger | Urticaceae | *Neraudia angulata* | Je | 2012-1 | KUN | Hawaii, USA | – | – | MH357910 | MH358291 | MH358105 | MH358212 | MH358008 | – |
| Sanger | Urticaceae | *Neraudia kauaiensis* |  | 678003 | KUN | Hawaii, USA | – | – | KF137883 | KF138364 | KF138199 | – | KF138027 | – |
| Sanger | Urticaceae | *Neraudia melastomifolia* |  | 90861 | KUN | Hawaii, USA | – | – | KF137884 | KF138365 | KF138200 | KF138518 | KF138028 | – |
| Angiosperms353 / Sanger | Urticaceae | *Nothocnide mollissima* | Beaman, J.H. | 8882 | K | Malaysia | 23934 | – | KF137885 | KF138366 | KF138201 | KF138519 | – | ERS5501739 |
| Sanger | Urticaceae | *Obetia aldabrensis* | Renvoize | 1357 | US | Seychelles | – | – | KM586460 | KM586632 | – | – | – | – |
| Sanger | Urticaceae | *Obetia pinnatifida* | Greenway | 11371 | EA | Tanzania | – | – | KM586449 | KM586621 | – | – | – | – |
| Angiosperms353 | Urticaceae | *Obetia radula* | Greenway, P.J.; Kanuri | 11371 | K | Tanzania, the United Republic of | 23918 | – | – | – | – | – | – | ERS4414129 |
| Sanger | Urticaceae | *Obetia radula* | Deng | 700 | KUN | Kenya | – | – | KM586431 | KM586603 | – | – | – | – |
| Sanger | Urticaceae | *Obetia tenax* | Botha | Botha_9 | K | S Africa | – | – | KF137886 | KF138367 | – | – | – | – |
| Sanger | Urticaceae | *Oreocnide boniana* |  |  | KUN | Yunnan, China | – | – | – | – | MH358108 | – | – | – |
| Sanger | Urticaceae | *Oreocnide frutescens* | Liuj | 10623 | KUN | Yunnan, China | – | – | KF137887 | KF138368 | KF138203 | KF138521 | KF138029 | – |
| Sanger | Urticaceae | *Oreocnide frutescens subsp. occidentalis* | WuZY | 9260 | KUN | Yunnan, China | – | – | – | KF138370 | KF138205 | KF138523 | KF138031 | – |
| Sanger | Urticaceae | *Oreocnide pedunculata* | WuZY | 2012485 | KUN | Taiwan, China | – | – | MH357912 | MH358293 | MH358109 | MH358213 | MH358009 | – |
| Sanger | Urticaceae | *Oreocnide rubescens* | WuZY | 9234 | KUN | Yunnan, China | – | – | MH357913 | – | MH358111 | – | – | – |
| Sanger | Urticaceae | *Oreocnide tonkinensis var. discolor* |  |  | KUN | Yunnan, China | – | – | MH357914 | MH358295 | MH358112 | MH358215 | – | – |
| Angiosperms353 | Urticaceae | *Oreocnide trinervis* | Andau, D. | 2015 | K | Malaysia | 23935 | – | – | – | – | – | – | ERS5501740 |
| Sanger | Urticaceae | *Oreocnide trinervis* | WuZY | WuZY2012 474 | KUN | Taiwan, China | – | – | – | – | MH358113 | – | – | – |
|  |  |  |  |  |  | United Kingdom of Great Britain and Northern Ireland (the) |  |  |  |  |  |  |  |  |
| Angiosperms353 / Sanger | Urticaceae | *Parietaria judaica* | Fay, M.F. | 185 | K |  | 8071 | KF137779 | – | KF138371 | KF138206 | KF138524 | – | ERS5501728 |
| Sanger | Urticaceae | *Parietaria micrantha* | WuZY | 10373 | KUN | China | – | KF137780 | – | KF138372 | KF138207 | KF138525 | – | – |
| Sanger | Urticaceae | *Pellionia acutidentata* | Q. Shao & L. D. Duan | 62 | BM | China | – | – | KP858777 | – | – | – | – | – |
| Sanger | Urticaceae | *Pellionia brachyceras* | Wei Y.G. | 81 | IBK | Dahuadi, Guangxi, China | – | – | To submit | – | To submit | – | To submit | – |
| Sanger | Urticaceae | *Pellionia grijsii* | Y. H. Tseng | 1167 | TAI | China | – | – | KC420491 | – | – | – | – | – |
| Sanger | Urticaceae | *Pellionia heteroloba* | A. K. Monro & Y. G. Wei | 6459 | IBK | China | – | – | KP858806 | – | – | – | – | – |
| Sanger | Urticaceae | *Pellionia minima* | J. M. Hu | 1787 | TAI | Japan | – | – | KP858809 | – | – | – | – | – |
| Sanger | Urticaceae | *Pellionia radicans* |  | HSL113 | KUN | China | – | – | KF137891 | KF138375 | KF138210 | – | – | – |
| Sanger | Urticaceae | *Pellionia radicans* | Y. G. Wei | U | IBK | China | – | – | KP858811 | – | – | – | – | – |
| Sanger | Urticaceae | *Pellionia repens* | L. F. Fu & S. L. Huang FL0071 | FL0071 | IBK | China | – | – | KU161129 | – | – | – | – | – |
| Sanger | Urticaceae | *Pellionia retrohispida* | M. H. Li | 6 | BM | China | – | – | KP858808 | – | – | – | – | – |
| Sanger | Urticaceae | *Pellionia scabra* | Y. H. Tseng | 1224 | TAI | Taiwan | – | – | KC420492 | – | – | – | – | – |
| Sanger | Urticaceae | *Pellionia tsoongii* | WuZY | 9495 | KUN | China | – | – | KF137893 | KF138377 | KF138212 | – | – | – |
| Sanger | Urticaceae | *Pellionia viridis* |  | 5155 |  | Mt.OMI Westerm china | – | – | To submit | – | – | – | – | – |
| Sanger | Urticaceae | *Pellionia viridis* | H. G. Xu | 1995364 | MO | China | – | – | KP858805 | – | – | – | – | – |
| Angiosperms353 / Sanger | Urticaceae | *Phenax ballotifolius* | Wood, J.R.I.; et al. | 18048 | K | Bolivia (Plurinational State of) | 23942 | – | To submit | – | – | – | To submit | ERS5503003 |
| Sanger | Urticaceae | *Phenax hirtus* | C. R. Romero | COL000134 982 | COL | Colombia | – | – | – | – | – | – | MH151318 | – |

| Sanger | Urticaceae | *Phenax mexicanus* |  | XZB102 | E | Peru | – | – | MH357916 | MH358299 | – | MH358218 | – | – |
| --- | --- | --- | --- | --- | --- | --- | --- | --- | --- | --- | --- | --- | --- | --- |
| Sanger | Urticaceae | *Phenax sonneratii* |  | AM6399 | E | Bolivia | – | – | MH357917 | – | MH358117 | – | – | – |
| Sanger | Urticaceae | *Pilea alpina* |  |  | BM |  | – | – | DQ175543 | DQ179309 | – | – | – | – |
| Sanger | Urticaceae | *Pilea amplistipulata* | Shui Y.M. et. al. | 14319 | KUN | China | – | – | MT516340 | MT523095 | MT523051 | – | – | – |
| Sanger | Urticaceae | *Pilea angulata* | Fu L.F. et al. | FL0234 | IBK | China | – | – | MT516341 | MT523096 | MT523052 | – | – | – |
| Sanger | Urticaceae | *Pilea angustifolia* |  | 12-1838 | BM |  | – | – | DQ175556 | DQ179289 | – | – | – | – |
| Sanger | Urticaceae | *Pilea anisophylla* | Wen F. | WF182817- 10 | IBK | China | – | – | MT516342 | MT523097 | MT523053 | – | – | – |
| Sanger | Urticaceae | *Pilea aphrophila* |  |  | BM |  | – | – | DQ175589 | DQ179323 | – | – | – | – |
| Sanger | Urticaceae | *Pilea aquarum* | Wei Y.G. | 97 | IBK | China | – | – | MT516343 | MT523098 | MT523054 | – | – | – |
| Sanger | Urticaceae | *Pilea aquarum subsp. acutidentata* | Wen F. | WFLSH111 207 | IBK | China | – | – | MT516344 | MT523099 | MT523055 | – | – | – |
| Sanger | Urticaceae | *Pilea balansae* | Huang S.L. | HSL118-1 | IBK | Vietnam | – | – | MT516345 | MT523100 | – | – | – | – |
| Sanger | Urticaceae | *Pilea basicordata* |  | HSL012 | PE | China | – | – | DQ175614 | DQ179361 | – | – | – | – |
| Sanger | Urticaceae | *Pilea benguetensis* |  |  | BM | Phillipines | – | – | DQ175554 | DQ179337 | – | – | – | – |
| Sanger | Urticaceae | *Pilea boniana* | Qin et al. 3193 | 3193 | KUN | China | – | – | MT516347 | MT523102 | MT523057 | – | – | – |
| Angiosperms353 | Urticaceae | *Pilea cadierei* |  |  | K | – | 24019 | – | – | – | – | – | – | ERS4414131 |
| Sanger | Urticaceae | *Pilea cadierei* | Xin Z.B. | XZB102 | IBK | China | – | – | MT516348 | MT523103 | MT523058 | – | – | – |
| Sanger | Urticaceae | *Pilea cavaleriei* | LiDZ | 1080 | KUN | China | – | – | KF137895 | KF138380 | KF138214 | – | – | – |
| Sanger | Urticaceae | *Pilea ciliata* |  |  | BM | Jamaica | – | – | DQ175538 | DQ179300 | – | – | – | – |
| Sanger | Urticaceae | *Pilea clementis* |  |  | BM | Cuba | – | – | DQ175550 | DQ179310 | – | – | – | – |
| Sanger | Urticaceae | *Pilea consanguinea* |  | AM6809 | BM | Santo Domingo | – | – | DQ175539 | DQ179312 | – | – | – | – |
| Sanger | Urticaceae | *Pilea cordistipulata* | Huang S.L. | HSL140 | IBK | China | – | – | MT516350 | MT523105 | MT523060 | – | – | – |
| Sanger | Urticaceae | *Pilea costata* |  | WuZY- 09199 | BM | Peru | – | – | DQ175595 | DQ179290 | – | – | – | – |
| Sanger | Urticaceae | *Pilea daguensis* |  | WuZY- 09107 | BM | Mexico | – | – | DQ175567 | DQ179332 | – | – | – | – |
| Sanger | Urticaceae | *Pilea dauciodora* |  | HSL124 | BM | Mexico | – | – | DQ175562 | DQ176857 | – | – | – | – |
| Sanger | Urticaceae | *Pilea digitata* |  |  | MO | Panama | – | – | DQ175559 | DQ179326 | – | – | – | – |
| Sanger | Urticaceae | *Pilea dolichocarpa* | Monro A.K. | 6399 | IBK | China | – | – | MT516351 | MT523106 | MT523061 | – | – | – |
| Sanger | Urticaceae | *Pilea dominguensis* |  |  | BM | Santo Domingo | – | – | DQ175541 | DQ179313 | – | – | – | – |
| Sanger | Urticaceae | *Pilea ecboliophylla* |  |  | BM | Mexico | – | – | DQ175531 | DQ179292 | – | – | – | – |
| Sanger | Urticaceae | *Pilea elegantissima* |  | 12-1247 |  | China | – | – | MH357923 | MH358303 | MH358124 | – | – | – |
| Sanger | Urticaceae | *Pilea elliptilimba* | Huang S.L. | HSL113 | IBK | China | – | – | MT516352 | MT523107 | – | – | – | – |
| Sanger | Urticaceae | *Pilea foliosa* |  |  | BM | Peru | – | – | DQ175571 | DQ179291 | – | – | – | – |
| Sanger | Urticaceae | *Pilea forgetii* |  | HSL149 | BM | Panama | – | – | DQ175585 | DQ179333 | – | – | – | – |
| Sanger | Urticaceae | *Pilea forsythiana* |  | HSL099 | BM | Dominica | – | – | DQ175546 | DQ179311 | – | – | – | – |
| Sanger | Urticaceae | *Pilea fruticosa* |  |  | BM | Borneo | – | – | DQ175604 | DQ179353 | – | – | – | – |
| Sanger | Urticaceae | *Pilea glaberrima* |  | STET2663 | BM | Nepal | – | – | DQ175600 | DQ179352 | – | – | – | – |
| Sanger | Urticaceae | *Pilea gracilis* | Wei Y.G. | 39 | IBK | China | – | – | MT516353 | MT523108 | – | – | – | – |
| Sanger | Urticaceae | *Pilea grandifolia* |  | AM6818 | BM | Jamaica | – | – | DQ175551 | DQ179303 | – | – | – | – |
| Sanger | Urticaceae | *Pilea guizhouensis* | Monro A.K. | 6715 | IBK | China | – | – | MT516354 | MT523109 | MT523062 | – | – | – |
| Sanger | Urticaceae | *Pilea harrisii* |  |  | BM | Jamaica | – | – | DQ175537 | DQ179302 | – | – | – | – |
| Sanger | Urticaceae | *Pilea hexagona* | Sino-Vietnamese expidition | 775 | KUN | China | – | – | MT516356 | MT523111 | MT523064 | – | – | – |
| Sanger | Urticaceae | *Pilea hilliana* | liuzu | 2014 | KUN | China | – | – | MT516357 | MT523112 | MT523065 | – | – | – |
| Sanger | Urticaceae | *Pilea howelliana* | Wang Y.Z. | 4678 | KUN | China | – | – | MT516358 | MT523113 | MT523066 | – | – | – |
| Sanger | Urticaceae | *Pilea inaequalis* |  | HGX001 |  | Trinidad | – | – | DQ175552 | DQ179304 | – | – | – | – |
| Sanger | Urticaceae | *Pilea insolens* | FLPH Tibet Expedition | 12-1838 | IBK | China | – | – | MT516359 | MT523114 | MT523067 | – | – | – |
| Sanger | Urticaceae | *Pilea irrorata* |  |  | BM | Mexico | – | – | DQ175535 | DQ179294 | – | – | – | – |
| Sanger | Urticaceae | *Pilea japonica* | Huang S.L. | HSL012 | IBK | China | – | – | MT516360 | MT523115 | MT523068 | – | – | – |
| Sanger | Urticaceae | *Pilea krugii* |  | HSL116 | BM | Puerto Rico | – | – | DQ175581 | DQ179315 | – | – | – | – |

| Sanger | Urticaceae | *Pilea lapestris* |  | 726 | BM | Indonesia | – | – | DQ175598 | DQ179341 | – | – | – | – |
| --- | --- | --- | --- | --- | --- | --- | --- | --- | --- | --- | --- | --- | --- | --- |
| Sanger | Urticaceae | *Pilea lindeniana* |  |  | BM | Cuba | – | – | DQ175547 | DQ179314 | – | – | – | – |
| Angiosperms353 | Urticaceae | *Pilea longicaulis* | Monro, A.K. | 7590 | K | – | – | – | – | – | – | – | – | ERS5501984 |
| Sanger | Urticaceae | *Pilea longicaulis* |  |  | PE | China | – | – | DQ175611 | DQ179363 | – | – | – | – |
| Sanger | Urticaceae | *Pilea longicaulis var. erosa* | Monro A.K. | 6809 | IBK | China | – | – | MT516361 | MT523116 | – | – | – | – |
| Sanger | Urticaceae | *Pilea longipedunculata* | WuZY | 9199 | KUN | China | – | – | KF137897 | KF138382 | KF138216 | – | – | – |
| Sanger | Urticaceae | *Pilea martinii* | WuZY | 9107 | KUN | China | – | – | KF137898 | KF138383 | KF138217 | – | – | – |
| Sanger | Urticaceae | *Pilea melastomoides* | Huang S.L. | HSL124 | IBK | China | – | – | MT516363 | MT523118 | MT523070 | – | – | – |
| Sanger | Urticaceae | *Pilea mexicana* |  | AM6433 | BM | Panama | – | – | DQ175579 | DQ179278 | – | – | – | – |
| Sanger | Urticaceae | *Pilea microphylla* |  |  | BM | Brazil | – | – | MH357928 | MH358307 | MH358129 | – | – | – |
| Sanger | Urticaceae | *Pilea monilifera* |  |  |  | China | – | – | MK911055 | MK911077 | MK911100 | – | – | – |
| Sanger | Urticaceae | *Pilea multicellularis* | Tibet Expedition | 12-1247 | IBK | China | – | – | MT516365 | MT523120 | MT523072 | – | – | – |
| Sanger | Urticaceae | *Pilea nigrescens* |  |  | BM | Jamaica | – | – | DQ175582 | DQ179301 | – | – | – | – |
| Sanger | Urticaceae | *Pilea nonggangensis* | Huang S.L. | HSL149 | IBK | China | – | – | MT516366 | MT523121 | MT523073 | – | – | – |
| Sanger | Urticaceae | *Pilea notata* | Huang S.L. | HSL099 | IBK | China | – | – | MT516368 | MT523123 | MT523075 | – | – | – |
| Sanger | Urticaceae | *Pilea nummularifolia* |  |  | BM | Peru | – | – | DQ175588. | DQ179316 | – | – | – | – |
| Sanger | Urticaceae | *Pilea oxyodon* |  |  | KUN | China | – | – | KF137902 | KF138387 | KF138221 | – | – | – |
| Sanger | Urticaceae | *Pilea paniculigera* | Monro A.K. | 6818 | IBK | China | – | – | MT516369 | MT523124 | MT523076 | – | – | – |
| Sanger | Urticaceae | *Pilea pansamalana* |  | Wei047 | BM | Mexico | – | – | DQ175533 | DQ179296 | – | – | – | – |
| Sanger | Urticaceae | *Pilea pellionioides* | Hu G.X. | HGX001 | IBK | China | – | – | MT516370 | MT523125 | MT523077 | – | – | – |
| Sanger | Urticaceae | *Pilea pelonae* |  |  | BM | Dominican | – | – | DQ175540 | DQ179327 | – | – | – | – |
| Sanger | Urticaceae | *Pilea peltata* | Huang S.L. | HSL116 | IBK | China | – | – | MT516371 | MT523126 | MT523078 | – | – | – |
| Sanger | Urticaceae | *Pilea penninervis* | Wei Y.G. | 726 | IBK | China | – | – | MT516372 | MT523127 | MT523079 | – | – | – |
| Sanger | Urticaceae | *Pilea peperomiifolia* |  | 2104H | BM | Virgin Islands | – | – | DQ175569 | DQ179281 | – | – | – | – |
| Sanger | Urticaceae | *Pilea peperomioides* |  | WF150423- 33 | BM | cultivated in UK | – | – | DQ175605 | DQ179350 | – | – | – | – |
| Sanger | Urticaceae | *Pilea peploides* | Monro A.K. | 6433 | IBK | China | – | – | MT516373 | MT523128 | MT523080 | – | – | – |
| Sanger | Urticaceae | *Pilea peploides var. major* |  |  | BM | China | – | – | MH357931 | – | MH358132 | – | – | – |
| Sanger | Urticaceae | *Pilea pittieri* |  |  |  | Peru | – | – | DQ175560 | DQ179328 | – | – | – | – |
| Sanger | Urticaceae | *Pilea plataniflora* |  |  | BM | Japan | – | – | DQ175599 | DQ179349 | – | – | – | – |
| Sanger | Urticaceae | *Pilea pleuroneura* |  |  | BM | Guatemala | – | – | DQ175532 | DQ179297 | – | – | – | – |
| Sanger | Urticaceae | *Pilea pseudonotata* | Wei Y.G. | 47 | IBK | China | – | – | MT516375 | MT523130 | MT523082 | – | – | – |
| Sanger | Urticaceae | *Pilea pubescens* |  | EXLS-0272 | BM | Belize | – | – | DQ175558 | DQ179325 | – | – | – | – |
| Sanger | Urticaceae | *Pilea pumila* | Huang S.L. | 2104H | IBK | China | – | – | MT516376 | MT523131 | MT523083 | – | – | – |
| Sanger | Urticaceae | *Pilea racemiformis* | Wen F. | WF150423- 33 | IBK | China | – | – | MT516377 | MT523132 | MT523084 | – | – | – |
| Sanger | Urticaceae | *Pilea racemosa* |  |  | BM | China | – | – | DQ175602 | DQ179347 | – | – | – | – |
| Sanger | Urticaceae | *Pilea receptacularis* |  |  | PE | China | – | – | DQ175612 | DQ179362 | – | – | – | – |
| Sanger | Urticaceae | *Pilea rivularis* |  | 37636 | BM | Tanzania | – | – | DQ175606 | DQ179358 | – | – | – | – |
| Sanger | Urticaceae | *Pilea rufa* |  | WYG18051 9-02 | BM | Jamaica | – | – | DQ175578 | DQ179299 | – | – | – | – |
| Sanger | Urticaceae | *Pilea semisessilis* | Zhou Z.K. et al. | EXLS-0272 | KUN | China | – | – | MT516378 | MT523133 | MT523085 | – | – | – |
| Sanger | Urticaceae | *Pilea sinofasciata* |  |  | BM | China | – | – | KF137905 | KF138389 | KF138224 | – | – | – |
| Sanger | Urticaceae | *Pilea spathulifolia* |  |  | BM | Dominican Republic | – | – | DQ175570 | DQ179282 | – | – | – | – |
| Sanger | Urticaceae | *Pilea spicata* | Burkill H. | 37636 | K | China | – | – | MT516380 | MT523135 | MT523087 | – | – | – |
| Sanger | Urticaceae | *Pilea subcoriacea* | Wei Y.G. | 180519-02 | IBK | China | – | – | MT516381 | MT523136 | MT523088 | – | – | – |
| Sanger | Urticaceae | *Pilea succulenta* |  |  | BM | Cayman Islands | – | – | DQ175565 | DQ179280 | – | – | – | – |
| Sanger | Urticaceae | *Pilea swinglei* |  |  | PE | China | – | – | MH357933 | – | MH358134 | – | – | – |
| Sanger | Urticaceae | *Pilea ternifolia* |  |  | BM | Nepal | – | – | DQ175597 | DQ179346 | – | – | – | – |

| Sanger | Urticaceae | *Pilea tetraphylla* |  |  | BM | Madagascar | – | – | MH357934 | MH358310 | MH358135 | – | – | – |
| --- | --- | --- | --- | --- | --- | --- | --- | --- | --- | --- | --- | --- | --- | --- |
| Sanger | Urticaceae | *Pilea thymifolia* |  |  | BM | Peru | – | – | DQ175568 | DQ179283 | – | – | – | – |
| Sanger | Urticaceae | *Pilea tridentata* |  | AM6770 | BM | Mexico | – | – | DQ175536 | DQ179293 | – | – | – | – |
| Sanger | Urticaceae | *Pilea tripartita* |  | WF180821- 01 | BM | Panama | – | – | DQ175617 | DQ176859 | – | – | – | – |
| Sanger | Urticaceae | *Pilea tsiangiana* | Monro A.K. | 6770 | IBK | China | – | – | MT516382 | MT523137 | MT523089 | – | – | – |
| Sanger | Urticaceae | *Pilea umbrosa* | Wen F. | WF180821- 01 | IBK | China | – | – | MT516383 | MT523138 | MT523090 | – | – | – |
| Sanger | Urticaceae | *Pilea unciformis* | Huang S.L. | HSL132 | IBK | China | – | – | MT516384 | MT523139 | MT523091 | – | – | – |
| Sanger | Urticaceae | *Pilea villicaulis* | Shui et al. | 12871 | KUN | China | – | – | MT516385 | MT523140 | MT523092 | – | – | – |
| Sanger | Urticaceae | *Pilea virgata* |  | HSL132 | BM | Jamaica | – | – | DQ175548 | DQ179329 | – | – | – | – |
| Sanger | Urticaceae | *Pilea vulcanica* |  | 12871 | BM | Panama | – | – | DQ175563 | DQ179284 | – | – | – | – |
| Sanger | Urticaceae | *Pilea weddellii* |  |  | BM | Jamaica | – | – | DQ175545 | DQ179308 | – | – | – | – |
| Sanger | Urticaceae | *Pilea weimingii* | Lv R.D. | LRD001 | IBK | China | – | – | MT516386 | MT523141 | MT523093 | – | – | – |
| Sanger | Urticaceae | *Pipturus arborescens* |  | 11879 | KUN | Taiwan, China | – | – | KF137908 | KF138392 | KF138227 | KF138545 | – | – |
| Sanger | Urticaceae | *Pipturus argenteus* |  |  | E | Papua New Guinea | – | – | MH357935 | MH358311 | – | MH358236 | – | – |
| Sanger | Urticaceae | *Pipturus kauaiensis* |  | 90441 |  | Hawaii, USA | – | – | KF137910 | KF138394 | KF138229 | KF138546 | KF138051 | – |
| Angiosperms353 | Urticaceae | *Pipturus montanus* | Crayn, D.M.; Sennart, S. | 540 | K | – | – | – | – | – | – | – | – | ERS5503161 |
| Sanger | Urticaceae | *Pipturus ruber* |  | 8082 | KUN | Hawaii, USA | – | – | – | – | – | – | – | – |
| Angiosperms353 | Urticaceae | *Poikilospermum acuminatum* | Risdale, C.E. et al. | 1268 |  |  | – | – | – | – | – | – | – | – |
| Angiosperms353 | Urticaceae | *Poikilospermum amboinense* | Takeuchi, W.; Ama, D. | 16391 |  |  | – | – | – | – | – | – | – | – |
| Sanger | Urticaceae | *Poikilospermum cordifolium* | Sinclair&Kadim | 10358 | E | Malaysia | – | – | To submit | – | – | – | – | – |
| Angiosperms353 | Urticaceae | *Poikilospermum cordifolium* | DeWilde et al. | SAN 143991 |  |  | – | – | – | – | – | – | – | – |
| Angiosperms353 | Urticaceae | *Poikilospermum erectum* | Risdale, C.E. | SMHI 482 |  |  | – | – | – | – | – | – | – | – |
| Angiosperms353 | Urticaceae | *Poikilospermum inaequale* | Takeuchi, W. et al. | 19531 |  |  | – | – | – | – | – | – | – | – |
| Sanger | Urticaceae | *Poikilospermum lanceolatum* | WuZY | 9235 | KUN | China | – | KF137786 | KF137912 | KF138396 | KF138231 | KF138548 | KF138053 | – |
| Angiosperms353 | Urticaceae | *Poikilospermum lanceolatum* | Middleton, D.J. et al. | 1744 |  |  | – | – | – | – | – | – | – | – |
| Sanger | Urticaceae | *Poikilospermum lanceolatum* | Wu | 9235 | KUN | China | – | – | KF137912 | KF138396 | – | – | – | – |
| Angiosperms353 | Urticaceae | *Poikilospermum microstachys* | Niyomdham, C. et al. | 1027 |  |  | – | – | – | – | – | – | – | – |
| Angiosperms353 | Urticaceae | *Poikilospermum naucleiflorum* | Chantaranothai, P. | 1215 |  |  | – | – | – | – | – | – | – | – |
| Angiosperms353 | Urticaceae | *Poikilospermum nobile* | Johns, R.J. | 8786 |  |  | – | – | – | – | – | – | – | – |
| Sanger | Urticaceae | *Poikilospermum scabrinervium* | Wilkie | 94166 | E | Indonesia | – | – | To submit | – | – | – | – | – |
| Angiosperms353 | Urticaceae | *Poikilospermum scabrinervium* | Dewol et al. | SAN 124644 |  |  | – | – | – | – | – | – | – | – |
| Angiosperms353 | Urticaceae | *Poikilospermum scortechinii* | T&P | 476 |  |  | – | – | – | – | – | – | – | – |
| Sanger | Urticaceae | *Poikilospermum sp. 1* | Sun | 13176 | KUN | Laos | – | – | KM586453 | KM586625 | – | – | – | – |
| Sanger | Urticaceae | *Poikilospermum suaveolens* | WuZY | 9160 | KUN | China | – | – | KF137914 | KF138398 | KF138233 | KF138550 | KF138054 | – |
| Angiosperms353 | Urticaceae | *Poikilospermum suaveolens* | Radin, J. et al. | SAN 133007 |  |  | – | – | – | – | – | – | – | – |
| Sanger | Urticaceae | *Poikilospermum suaveolens* |  |  | KUN | China | – | – | KF137913 | KF138397 | – | – | – | – |
| Angiosperms353 | Urticaceae | *Poikilospermum tangaum* | Postar; Geoffray | SAN 145768 |  |  | – | – | – | – | – | – | – | – |
| Sanger | Urticaceae | *Pouzolzia australis* | Hadiah | 393 | NSW | Australia: South Pacific | – | – | – | AY208723 | AY208700 | – | – | – |
| Sanger | Urticaceae | *Pouzolzia calophylla* |  |  |  | Xizang, China | – | – | KF137915 | KF138399 | KF138234 | KF138551 | – | – |
| Sanger | Urticaceae | *Pouzolzia elegans var. elegans* | L0942456 | L0942456 | KUN | Taiwan, China | – | – | – | – | MH358140 | – | – | – |
| Sanger | Urticaceae | *Pouzolzia guineensis* | Bidgood et al. | 3008 | K | Tanzania | – | – | – | KF138400 | KF138235 | KF138552 | KF138055 | – |
| Sanger | Urticaceae | *Pouzolzia mixta* | Lovett & Congdon | 2945 | K | Tanzania | – | – | KF137916 | KF138401 | KF138236 | KF138553 | – | – |
| Sanger | Urticaceae | *Pouzolzia poeppigiana* |  | HSL140 | E | Peru | – | – | MH357938 | MH358315 | MH358141 | – | – | – |

| Sanger | Urticaceae | *Pouzolzia rugulosa* | LiDZ | 1071 | KUN | Nepal, Kathmandu | – | – | KF137817 | KF138288 | KF138125 | KF138449 | KF137960 | – |
| --- | --- | --- | --- | --- | --- | --- | --- | --- | --- | --- | --- | --- | --- | --- |
| Angiosperms353 | Urticaceae | *Pouzolzia sanguinea* | Sambuling, S. | 449 | K | Malaysia | 23922 | – | – | – | – | – | – | ERS5501735 |
| Sanger | Urticaceae | *Pouzolzia sanguinea* | WuZY | 9483 |  | Guangxi, China | – | – | KF137918 | KF138403 | KF138238 | KF138555 | KF138057 | – |
| Sanger | Urticaceae | *Pouzolzia sanguinea var. elegans* |  | Wei039 |  | Xizang, China | – | – | KF137917 | KF138402 | KF138237 | KF138554 | KF138056 | – |
| Sanger | Urticaceae | *Pouzolzia sp.* | RC | 1682 |  | Panchthar, Nepal | – | – | KF137919 | KF138404 | KF138239 | KF138556 | KF138058 | – |
| Angiosperms353 | Urticaceae | *Pouzolzia zeylanica* | Sibil, J. | 83 | K | Malaysia | 23923 | – | – | – | – | – | – | ERS5501736 |
| Sanger | Urticaceae | *Pouzolzia zeylanica* | WuZY | 10167 |  | Yunnan, China | – | – | KF137920 | KF138405 | KF138240 | KF138557 | KF138059 | – |
| Sanger | Urticaceae | *Procris archboldiana* | A. C. Smith | 5987 | K | Fiji | – | – | KP858785 | – | – | – | – | – |
| Sanger | Urticaceae | *Procris crenata* | Y. H. Tseng | 1170 | TAI | China | – | – | KP858782 | – | – | – | – | – |
| Sanger | Urticaceae | *Procris crenata* | Shi | 15115 | A | China | – | – | KP858783 | – | – | – | – | – |
| Sanger | Urticaceae | *Procris frutescens* | Takeuchi | 8800 | A | Papua New Guinea | – | – | KP858781 | – | – | – | – | – |
| Sanger | Urticaceae | *Procris montana* | R. O. Gardenr | 5955 | MO | Australia | – | – | KP858786 | – | – | – | – | – |
| Angiosperms353 | Urticaceae | *Procris wightiana* | Monro, A.K.; Fu, L.F. | 7603 | K | – | – | – | – | – | – | – | – | ERS5502206 |
| Angiosperms353 | Urticaceae | *Rousselia humilis* | Ekman, E.L. | 4973 | K | Haiti | 23945 | – | – | – | – | – | – | ERS5501747 |
| Sanger | Urticaceae | *Rousselia humilis* | Howard | 6273 | US | Cuba | – | – | – | KM586645 | KM586559 | – | – | – |
| Sanger | Urticaceae | *Rousselia humilis* | Haroslav & Holman | 435 | BM | Cuba | – | – | – | – | – | – | – | – |
| Angiosperms353 | Urticaceae | *Sarcochlamys pulcherrima* | Grierson, A.J.C.; Long, D.G. | 1525 | K | Bhutan | 23939 | – | – | – | – | – | – | ERS5501743 |
| Sanger | Urticaceae | *Sarcochlamys pulcherrima* |  |  | KUN | China | – | KF137790 | KF137924 | KF138409 | KF138244 | KF138561 | – | – |
|  |  |  |  |  |  | United Kingdom of Great Britain and Northern Ireland (the) |  |  |  |  |  |  |  |  |
| Angiosperms353 | Urticaceae | *Soleirolia soleirolii* | Sheahan, M-C. | 17 | K |  | 8154 | – | – | – | – | – | – | ERS4414200 |
| Sanger | Urticaceae | *Soleirolia soleirolii* | Monro A.K. | s.n. | BM | Cultivated | – | – | KF137926 | KF138411 | KF138246 | KF138563 | KF138063 | – |
| Angiosperms353 | Urticaceae | *Touchardia latifolia* | Stone, B.C. | 3626 | K | United States of America | 23938 | – | – | – | – | – | – | ERS4414130 |
| Sanger | Urticaceae | *Touchardia latifolia* | Jeffrey | 201101 | KUN | Hawaii | – | – | KF137927 | KF138412 | – | – | – | – |
| Sanger | Urticaceae | *Urera altissima* | Lliully | 460 | K | Bolivia | – | – | To submit | – | – | – | – | – |
| Sanger | Urticaceae | *Urera aurantiaca* | Loza | 63 | K | Bolivia | – | – | To submit | To submit | – | – | – | – |
| Angiosperms353 | Urticaceae | *Urera baccifera* | Goes, S.P.; et al. | 20285  /G150694 | K | – | – | – | – | – | – | – | – | ERS4414134 |
| Sanger | Urticaceae | *Urera baccifera* | Cayola | 2530 | BM | Bolivia | – | – | To submit | To submit | – | – | – | – |
| Sanger | Urticaceae | *Urera batesii* | Carvalho | 3412 | K | Equatorial Guinea | – | – | KF971186 | KF971219 | – | – | – | – |
| Sanger | Urticaceae | *Urera cameroonensis* | Leeuenberg | 7187 | K |  | – | – | To submit | – | – | – | – | – |
| Angiosperms353 | Urticaceae | *Urera caracasana* | Monro, A.K. | 6840 | K | – | – | – | – | – | – | – | – | ERS4414132 |
| Sanger | Urticaceae | *Urera caracasana* | Wood | 8834 | K | Bolivia | – | – | KF137929.1 | KF138415 | – | – | – | – |
| Sanger | Urticaceae | *Urera cordifolia* | Carvalho | 3046 | K | CM | – | – | To submit | – | – | – | – | – |
| Sanger | Urticaceae | *Urera cordifolia* | Sunderland | 1190 | K | CM | – | – | To submit | – | – | – | – | – |
| Sanger | Urticaceae | *Urera elata* | Lewis | 2224 | US | Panama | – | – | KM58647 | KM586642 | – | – | – | – |
| Sanger | Urticaceae | *Urera fenestrata* | Monro A.K. | 5452 | K | Costa Rica | – | – | To submit | To submit | – | – | – | – |
| Sanger | Urticaceae | *Urera fischeri* | Luke & Luke | 7087 | K | KE | – | – | To submit | – | – | – | – | – |
| Sanger | Urticaceae | *Urera fischeri* | Faden & Beentje | 8522 | EA | Kenya | – | – | KM586427 | KM586599 | – | – | – | – |
| Sanger | Urticaceae | *Urera glabra* | s.n. | 100694 | KUN | Hawaii | – | – | KF1379930 | KF138416 | – | – | – | – |
| Sanger | Urticaceae | *Urera glabriuscula* | Calonico | 21101 | BM | Mexico | – | – | To submit | – | – | – | – | – |
| Angiosperms353 | Urticaceae | *Urera hypselodendron* | Faden, R.B. et al. | 85200 | K | Kenya | 23913 | – | – | – | – | – | – | ERS4414137 |
| Sanger | Urticaceae | *Urera hypselodendron* | Beenlije | 3257 | EA | Kenya | – | – | KM586430 | KM586602 | – | – | – | – |
| Sanger | Urticaceae | *Urera keayi* | Leeuwenberg | 4474 | K | CI | – | – | To submit | – | – | – | – | – |
| Sanger | Urticaceae | *Urera killipiana* | Serviu | 372 | BM | Mexico | – | – | To submit | To submit | – | – | – | – |

| Angiosperms353 | Urticaceae | *Urera laciniata* | Morawetz, W.; Wallnofer, B. | M13-29985 | K | – | – | – | – | – | – | – | – | ERS4414126 |
| --- | --- | --- | --- | --- | --- | --- | --- | --- | --- | --- | --- | --- | --- | --- |
| Sanger | Urticaceae | *Urera laciniata* | Araujo | 3016 | BM | Bolivia | – | – | To submit | To submit | – | – | – | – |
| Sanger | Urticaceae | *Urera lianiformis* | Solano | 6825 | BM | Costa Rica | – | – | KF138570 | KF138418 | – | – | – | – |
| Sanger | Urticaceae | *Urera mannii* | Lowe | 1736 | K | NG | – | – | To submit | – | – | – | – | – |
| Sanger | Urticaceae | *Urera oblongifolia* | Morton & Jarr | 3517 | K | SL | – | – | To submit | – | – | – | – | – |
| Sanger | Urticaceae | *Urera obovata* | Thijssen | 40 | K | CI | – | – | To submit | – | – | – | – | – |
| Sanger | Urticaceae | *Urera pacifica* | Steinman | 3265 | BM | Mexico | – | – | To submit | To submit | – | – | – | – |
| Sanger | Urticaceae | *Urera repens* | Eiminjeze & Oguntayo | 72749 | K | NG | – | – | To submit | – | – | – | – | – |
| Sanger | Urticaceae | *Urera rigida* | Adam | 25511 | K | LR | – | – | To submit | – | – | – | – | – |
| Sanger | Urticaceae | *Urera rigida* | Breteler | Breteler | K | LR | – | – | To submit | – | – | – | – | – |
| Sanger | Urticaceae | *Urera robusta* | Adams | 4823 | K | GH | – | – | To submit | – | – | – | – | – |
| Angiosperms353 | Urticaceae | *Urera sandwicensis* | Bulmer, C. | 379 | K | United States of America | 23914 | – | – | – | – | – | – | – |
| Sanger | Urticaceae | *Urera sansibarica* | Luke | 11527 | EA | Tanzania | – | – | KM586428 | KM586600 | – | – | – | – |
| Sanger | Urticaceae | *Urera simplex* | Monro A.K. | 5102 | K | Panama | – | – | To submit | To submit | – | – | – | – |
| Sanger | Urticaceae | *Urera spaerophyllya* | Humbert | 3055 | K | MG | – | – | To submit | – | – | – | – | – |
| Sanger | Urticaceae | *Urera thonneri* | Breteler et al. | 2378 | K | CM | – | – | To submit | – | – | – | – | – |
| Sanger | Urticaceae | *Urera trinervis* | Friis | 3920 | C | Ethiopia | – | – | KF137932 | KF138421 | – | – | – | – |
| Sanger | Urticaceae | *Urtica angustifolia* |  |  | KUN | China | – | KF137796 | KF137933 | KF138421 | KF138256 | KF138573 | KF138067 | – |
| Sanger | Urticaceae | *Urtica aquatica* | M. Weigend | 7478 | BSB | Peru | – | – | KF971214 | KF971247 | – | – | – | – |
| Sanger | Urticaceae | *Urtica ardens* |  | 81152 | KUN | China | – | – | KF137934 | KF138422 | KF138257 | KF138574 | – | – |
| Sanger | Urticaceae | *Urtica atrichocaulis* | WuZY | 10358 | KUN | China | – | – | KF137935 | KF138423 | KF13825 | KF138575 | KF138068 | – |
| Sanger | Urticaceae | *Urtica atrovirens* |  |  | E | Italy | – | – | MH357956 | – | MH358161 | MH358244 | – | – |
| Sanger | Urticaceae | *Urtica cannabina* | LZH | 2012 | KUN | China | – | – | MH357957 | MH358322 | MH358162 | MH358245 | MH358029 | – |
| Sanger | Urticaceae | *Urtica dioica subsp. dioica* | BROWP | 135 | KUN | UK | – | – | KF137936 | KF138424 | KF138259 | KF138576 | – | – |
| Sanger | Urticaceae | *Urtica echinata* |  |  | E | Peru | – | – | MH357961 | MH358325 | MH358166 | MH358248 | – | – |
| Sanger | Urticaceae | *Urtica fissa* | WuZY | 10378 | KUN | China | – | – | KF137937 | KF138425 | KF138260 | KF138577 | KF138069 | – |
| Sanger | Urticaceae | *Urtica flabellata* | M. Weigend et al. | 7728 | BSB | Peru | – | – | KF558908 | KF559028 | – | – | – | – |
| Sanger | Urticaceae | *Urtica hyperborea* |  | 81200 | KUN | China | – | – | KF137939 | KF138427 | KF138262 | KF13857 | KF138071 | – |
| Sanger | Urticaceae | *Urtica kioviensis* |  |  | E | Austria | – | – | MH357963 | MH358326 | MH358168 | MH358250 | MH358032 | – |
| Sanger | Urticaceae | *Urtica laetevirens* | Mawenzhang | 2011 | KUN | Canada | – | – | – | MH358327 | MH358169 | MH358251 | – | – |
| Sanger | Urticaceae | *Urtica magellanica* |  |  | E | Chile | – | – | MH357964 | – | MH358170 | MH358252 | – | – |
| Sanger | Urticaceae | *Urtica mairei* | WuZY | 9354 | KUN | China | – | KF137797 | KF137940 | KF138428 | KF138263 | KF138580 | – | – |
| Sanger | Urticaceae | *Urtica massaica* | J.M. Kimen & al. | KARI42/02 | EA | Kenya | – | – | KM586438 | KM586610 | KM586524 | – | – | – |
| Sanger | Urticaceae | *Urtica membranacea* |  |  | E | Greece | – | – | MH357968 | MH358329 | MH358174 | MH358255 | – | – |
| Sanger | Urticaceae | *Urtica sp. 1* | Lixinhui | 1102 | KUN | Kenya | – | – | KF137941 | KF138429 | KF138264 | KF138581 | KF138072 | – |
| Sanger | Urticaceae | *Urtica taiwaniana* | H. Sun | 11345 | KUN | Taiwan | – | – | KM586420 | KM586592 | KM586506 | – | – | – |
| Sanger | Urticaceae | *Urtica thunbergiana* | H. Sun | 11881 | KUN | China | – | – | KM586421 | KM586593 | KM586507 | – | – | – |
| Sanger | Urticaceae | *Urtica triangularis subsp. pinnatifida* |  | 80860 | KUN | China | – | – | KF137943 | KF138431 | KF138266 | KF138583 | KF138073 | – |
|  |  |  |  |  |  | United Kingdom of Great Britain and Northern Ireland  (the) |  |  |  |  |  |  |  |  |
| Angiosperms353 | Urticaceae | *Urtica urens* | Fay, M.F. | 173 | K |  | 8160 | – | – | – | – | – | – | ERS4414127 |
| Sanger | Urticaceae | *Urtica zayuensis* | WuZY | 10361 | KUN | China | – | – | KF137945 | KF138433 | KF138268 | KF138585 | KF138075 | – |
| Angiosperms353 | Urticaceae | *Zhengyia shennongensis* | Deng, T.; Zhang, D.G.; Sun, H. | 2295 | K | – | – | – | – | – | – | – | – | ERS5501754 |
| Sanger | Urticaceae | *Zhengyiia shennongensis* | SNJ Exped. | 2.0111E+10 | KUN | China | – | – | KC284949 | KC285001 | KC284975 | – | – | – |

| **Sequence data** | **Family** | **Taxon** | **Voucher** | **Collector No** | **Herbarium** | **Country** | **RBGKew BankId** | **18S** | **ITS** | **trnLF** | **rbcL** | **rpll4-rps8- infA-rpl36** | **matK** | **Angiosperms3 53 (A353)** |
| --- | --- | --- | --- | --- | --- | --- | --- | --- | --- | --- | --- | --- | --- | --- |
| Angiosperms353 | Cannabaceae | *Trema orientale* | Reeves, G.& | 37 | K | South Africa | 27834 | – | – | – | – | – | – | ERS8701987 |
|  |  |  | Weiblen, G.D.; Montgomery, R.; |  |  | – |  |  |  |  |  |  |  |  |
| Angiosperms353 | Moraceae | *Antiaropsis decipiens* | Isua, B.; Molem, K. | 1865 | K |  | – | – | – | – | – | – | – | ERS4414199 |
| Angiosperms353 | Moraceae | *Batocarpus orinocensis* | Palacios, W. | 3265 | K | – | – | – | – | – | – | – | – | ERS4414228 |
|  |  |  |  |  |  | Korea (the |  |  |  |  |  |  |  |  |
| Angiosperms353 | Moraceae | *Broussonetia kazinoki* | Chase, M.W. | 17827 | K | Republic of) | 17657 | – | – | – | – | – | – | ERS4414202 |
| Angiosperms353 | Moraceae | *Ficus sagittifolia* | Chase, M.W. | 19852 | K | Ivoryt Coast | 19722 | – | – | – | – | – | – | ERS4414205 |
|  |  |  | Wurdack, K.J.; Redden, K.; |  |  | – |  |  |  |  |  |  |  |  |
|  |  |  | Rodriguez, A.; Perry, C.; James, |  |  |  | | | | | | | | |
| Angiosperms353 | Moraceae | *Maquira guianensis* | H.; Simon, H.; Ragnauth, P. | 4570 | K | – – – – – – – To submit | | | | | | | | |
| Angiosperms353 | Moraceae | *Milicia africana* | Williamson, L. | 187 | K | – – – – – – – – ERS4414198 | | | | | | | | |
| Angiosperms353 | Moraceae | *Parartocarpus venenosa* | Renvoize, S.A.; Wilmot-Dear, M.; Saenz, A. | 1518 | K | – | – | – | – | – | – | – | – | ERS5501666 |
| Sanger | Urticaceae | *Achudemia javanica* | Robinson H.C. and Kloss C.B. | 1914 | SING | Indonesia | – | – | MT516339 | MT523094 | MT523050 | – | – | – |
| Sanger | Urticaceae | *Archiboehmeria atrata* | WuZY | 9469 | KUN | China, Guangxi | – | – | KF137798 | KF138269 | KF138106 | KF138434 | KF137946 | – |
| Angiosperms353 | Urticaceae | *Archiboehmeria atrata* | Hiep, N.T. et al. | 189 | K | Viet Nam | 23937 | – | – | – | – | – | – | ERS5501742 |
| Angiosperms353 / Sanger | Urticaceae | *Astrothalamus reticulatus* | Argent, G. et al. | 981987 | K |  | 23941 | – | KF137800 | KF138271 | KF138108 | – | To submit | ERS5501745 |
| Sanger | Urticaceae | *Australina flaccida* | Friis, I. et al. | 12293 | K | Ethiopia | – | KF137745 | KF137801 | KF138272 | KF138109 | KF138436 | MH357978 | – |
| Angiosperms353 | Urticaceae | *Australina pusilla* | Raven, P.H.; et al. | 25912 | K | Australia | 23949 | – | – | – | – | – | – | ERS5501750 |
| Sanger | Urticaceae | *Boehmeria aspera* | H. J. Sarrazola | 937 | HUA | Colombia | – | – | – | MH151324 | – | – | MH151315 | – |
| Sanger | Urticaceae | *Boehmeria bullata* | H. J. Sarrazola | CHG-46 | HUA | Colombia | – | – | – | – | – | – | MH151320 | – |
| Angiosperms353 / Sanger | Urticaceae | *Boehmeria burgerania* | Monro A.K. | 6842 | K | Costa Rica | – | – | To submit | – | – | – | – | ERS5503004 |
| Sanger | Urticaceae | *Boehmeria caudata* |  |  | E | Peru | – | – | – | MH358260 | MH358040 | – | – | – |
|  |  |  |  |  |  | Bolivia |  |  |  |  |  |  |  |  |
|  |  |  |  |  |  | (Plurinational State |  |  |  |  |  |  |  |  |
| Angiosperms353 | Urticaceae | *Boehmeria caudata* | Wood, J.R.I. | 18657 | K | of) | 24013 | – | – | – | – | – | – | ERS5501752 |
| Sanger | Urticaceae | *Boehmeria celtidifolia* | H. J. Sarrazola | 957 | HUA | Colombia | – | – | – | MH151327 | – | – | MH151323 | – |
| Sanger | Urticaceae | *Boehmeria clidemioides* | Nie | 4243 | KUN | China | – | – | KM586402 | KM586574 | KM586488 | MK955173 | MK931191 | – |
| Sanger | Urticaceae | *Boehmeria clidemioides var. umbrosa* | WuZY | 10336 | KUN | China, Yunna | – | – | KF137822 | KF138293 | KF138130 | KF138453 | KF137965 | – |
| Sanger | Urticaceae | *Boehmeria cylindrica* | Abbott | 18035 | FLAS | USA? | – | – | – | – | KJ773314 | – | KJ772586 | – |
| Sanger | Urticaceae | *Boehmeria densiflora* | WuZY | 2012450 | KUN | Taiwan | – | – | KF137806 | KF138277 | KF138114 | – | KF137951 | – |
| Sanger | Urticaceae | *Boehmeria depauperata* |  |  | KUN | Yunnan, China | – | – | KF137807 | KF138278 | KF138115 | KF138440 | – | – |
| Sanger | Urticaceae | *Boehmeria grandis* | Morden 1120 | 1120 | BISH | Hawaii | – | – | – | – | AF500354 | – | – | – |
| Sanger | Urticaceae | *Boehmeria holosericea* | TKMVP | 873 | ?? | Korea | – | – | KT119556 | – | – | – | – | – |
| Sanger | Urticaceae | *Boehmeria japonica* |  | 100024 | KUN |  | – | – | KF137808 | KF138279 | KF138116 | – | – | – |
|  |  |  |  |  | Herbarium of | China? |  |  |  |  |  |  |  |  |
| Sanger | Urticaceae | *Boehmeria japonica var. silvestrii* | Li | 26 | Jiujiang University |  | – | – | FJ750380 | FJ750411 | – | – | – | – |
| Sanger | Urticaceae | *Boehmeria japonica var. tenera* | D. Tao | 84 | ?? | China? | – | – | MK911052 | MK911074 | MK911097 | – | MK931192 | – |
| Sanger | Urticaceae | *Boehmeria nivea* | Liuj | 10645 | KUN | China, Fujian | – | – | KF137815 | KF138286 | KF138123 | KF138447 | KF137958 | – |
| Angiosperms353 | Urticaceae | *Boehmeria nivea* | Hu, S.Y. | 8113 | K | China | 24011 | – | – | – | – | – | – | ERS4414210 |
| Sanger | Urticaceae | *Boehmeria nivea var. tenacissima* | Liuj | 10679 | KUN | China, Zhejiang | – | – | KF137814 | KF138285 | KF138122 | KF138446 | KF137957 | – |
| Sanger | Urticaceae | *Boehmeria pavonii* | H. J. Sarrazola | 933 | HUA | Colombia | – | – | – | MH151326 | – | – | MH151319 | – |
| Sanger | Urticaceae | *Boehmeria penduliflora* | WuZY | 9460 | KUN | China, Guangxi | – | – | KF137816 | KF138287 | KF138124 | KF138448 | KF137959 | – |
|  |  |  |  |  | Herbarium of | China? |  |  |  |  |  |  |  |  |
| Sanger | Urticaceae | *Boehmeria pilosiuscula* | Li | 17 | Jiujiang University |  | – | – | FJ750372 | FJ750422 | – | – | – | – |

| Sanger | Urticaceae | *Boehmeria platanifolia* |  | AM6715 | NA | South Korea | – | – | – | – | KM218340 | – | – | – |
| --- | --- | --- | --- | --- | --- | --- | --- | --- | --- | --- | --- | --- | --- | --- |
|  |  |  |  |  | Herbarium of Jiujiang University | China? |  |  |  |  |  |  |  |  |
| Sanger | Urticaceae | *Boehmeria polystachya* | Li | 29 |  |  | – | – | FJ750376 | FJ750421 | – | – | – | – |
|  |  |  |  |  | Herbarium of Jiujiang University | China? |  |  |  |  |  |  |  |  |
| Sanger | Urticaceae | *Boehmeria pseudotricuspis* | Li | 12 |  |  | – | – | FJ750375 | FJ750400 | – | – | – | – |
| Sanger | Urticaceae | *Boehmeria ramiflora* | L0942407 | L0942407 | NHN | Jamaica | – | – | MH357855 | – | MH358042 | – | – | – |
|  |  |  |  |  | Herbarium of Jiujiang University | China? |  |  |  |  |  |  |  |  |
| Sanger | Urticaceae | *Boehmeria siamensis* | Li | 28 |  |  | – | – | FJ750374 | FJ750428 | – | – | – | – |
| Sanger | Urticaceae | *Boehmeria sieboldiana* |  | LiDZ1080 | E | Japan | – | – | MH357859 | – | MH358045 | MH358182 | – | – |
| Sanger | Urticaceae | *Boehmeria sp.* | RC | 1555 | KUN | Nepal, Mayagdi | – | – | KF137818 | KF138289 | KF138126 | KF138450 | KF137961 | – |
| Sanger | Urticaceae | *Boehmeria sp. 1* | Nie | 4249 | KUN | China? | – | – | – | KM586573 | KM586487 | – | – | – |
| Sanger | Urticaceae | *Boehmeria sp.2* |  |  | KUN | Dominica | – | – | MH357861 | MH358261 | MH358049 | – | MH357981 | – |
|  |  |  |  |  | Ramie Repository, Huazhong Agricultural University, Wuhan, China | China |  |  |  |  |  |  |  |  |
| Sanger | Urticaceae | *Boehmeria splitgerbera* | HZAURS | D9 |  |  | – | – | – | HQ380834 | – | – | – | – |
| Sanger | Urticaceae | *Boehmeria tsaratananensis* | Razanajtovo et al. | MHR 001 | G | Madagascar | – | – | – | – | – | – | – | – |
| Sanger | Urticaceae | *Boehmeria ulmifolia* | H. J. Sarrazola | 929 | HUA | Colombia | – | – | – | MH151325 | – | – | MH151317 | – |
| Sanger | Urticaceae | *Boehmeria virgata subsp. macrophylla* | WuZY | 9196 | KUN | China? | – | – | KF137811 | KF138282 | KF138119 | KF138443 | KF137954 | – |
|  |  |  |  |  | Herbarium of Jiujiang University | China? |  |  |  |  |  |  |  |  |
| Sanger | Urticaceae | *Boehmeria virgata var. densiglomerata* | Li | 24 |  |  | – | – | FJ750378 | FJ750417 | – | – | – | – |
| Sanger | Urticaceae | *Boehmeria virgata var. macrostachya* | HZAURS | X3 | ?? | China? | – | – | – | HQ380837 | – | – | – | – |
| Sanger | Urticaceae | *Boehmeria virgata var. rotundifolia* | Liuj | 10629 | KUN | China, Yunnan | – | – | KF137812 | KF138283 | KF138120 | KF138444 | KF137955 | – |
| Sanger | Urticaceae | *Boehmeria virgata var. scabrella* | WuZY | 9468 | KUN | China, Guangxi | – | – | KF137813 | KF138284 | KF138121 | KF138445 | KF137956 | – |
|  |  |  |  |  | Herbarium of Jiujiang University | China? |  |  |  |  |  |  |  |  |
| Sanger | Urticaceae | *Boehmeria virgata var. strigosa* | Li | 21 |  |  | – | – | FJ750383 | FJ750418 | – | – | – | – |
| Sanger | Urticaceae | *Boehmeria virgata var. tomentosa* | WuZY | 9011 | KUN | China, Yunnan | – | – | KF137820 | KF138291 | KF138128 | KF138452 | KF137963 | – |
|  |  |  |  |  | Herbarium of Jiujiang University | China? |  |  |  |  |  |  |  |  |
| Sanger | Urticaceae | *Boehmeria zollingeriana* | Li | 8 |  |  | – | – | FJ750366 | FJ750424 | – | – | – | – |
| Sanger | Urticaceae | *Boehmeria zollingeriana var. blinii* | LiDZ | 1084 | KUN | China, Guizhou | – | – | KF137824 | KF138295 | KF138132 | – | KF137966 | – |
| Sanger | Urticaceae | *Cecropia angustifolia* | Monro 4424 | 4424 | BM | Panama | – | – | – | – | – | – | – | – |
| Angiosperms353 / Sanger | Urticaceae | *Cecropia ficifolia* | Berg, C.C. et al. | 18418 | K | Brazil | 23955 | – | KF137825 | KF138296 | KF138133 | – | – | ERS4414209 |
| Sanger | Urticaceae | *Cecropia glazioviana* | Bruno | 16 | UFP | Brazil | – | – | MH357864 | MH358262 | MH358053 | – | – | – |
| Sanger | Urticaceae | *Cecropia hololeura* | Bruno | 15 | UFP | Brazil | – | – | – | MH358263 | MH358054 | – | MH357985 | – |
| Sanger | Urticaceae | *Cecropia obtusifolia* | Monro 3767 | 3767 | BM | El Salvador | – | – | – | KF138297 | KF138134 | KF138455 | KF137967 | – |
| Sanger | Urticaceae | *Cecropia pachystachya* | Bruno | 17 | UFP | Brazil | – | – | MH357866 | MH358265 | MH358056 | MH358187 | – | – |
| Angiosperms353 | Urticaceae | *Chamabainia cuspidata* | Zhing-tao, W. | 870226 | K | – | – | – | – | – | – | – | – | ERS5501757 |
| Sanger | Urticaceae | *Chamabainia cuspidata* | WuZY | 10086 | KUN | China, Yunnan | – | – | KF137827 | KF138299 | KF138136 | KF138457 | KF137969 | – |
| Sanger | Urticaceae | *Coussapoa glaberrima* | Br | 2015 | KUN | Brazil | – | – | – | MH358268 | MH358060 | – | MH357987 | – |
| Sanger | Urticaceae | *Coussapoa parvifolia* | A.K. Monro et al. | 6833 | BM | Costa Rica | – | – | – | KF138301 | – | KF138459 | – | – |
| Angiosperms353 | Urticaceae | *Coussapoa villosa* | Pennington, T.D. | 10630 | K | – | – | – | – | – | – | – | – | ERS5501756 |
| Sanger | Urticaceae | *Cypholophus heterophyllus* |  | 6032 | NHN | Fiji | – | – | – | MH358269 | – | – | – | – |
| Sanger | Urticaceae | *Cypholophus macrocephalus* |  |  | NHN | Vanuatu | – | – | MH357871 | MH358270 | MH358061 | MH358191 | MH357988 | – |
| Angiosperms353 | Urticaceae | *Cypholophus montanus* | Argent, G. | 535 | K | Indonesia | 23927 | – | – | – | – | – | – | ERS5501737 |
| Sanger | Urticaceae | *Cypholophus montanus* | L0942484 | L0942484 | NHN | Papua, Indonesia | – | – | MH357873 | – | MH358063 | – | – | – |

| Sanger | Urticaceae | *Cypholophus sp.* | L0406219 | L0406219 | NHN | Sulawesi, Indonesia | – | – | MH357874 | MH358273 | MH358065 | MH358192 | MH357991 | – |
| --- | --- | --- | --- | --- | --- | --- | --- | --- | --- | --- | --- | --- | --- | --- |
| Sanger | Urticaceae | *Cypholophus sp. 1* | L0792369 | L0792369 | NHN | Irian Jaya, Indonesia | – | – | – | – | MH358064 | – | MH357990 | – |
| Sanger | Urticaceae | *Cypholophus sp. 2* | L0792372 | L0792372 | NHN | Bougainville, Papua New Guinea | – | – | – | – | MH358066 | – | – | – |
| Sanger | Urticaceae | *Debregeasia elliptica* | WuZY | 10061 | KUN | China, Yunnan | – | – | KF137830 | KF138303 | KF138139 | KF138461 | KF137972 | – |
| Angiosperms353 | Urticaceae | *Debregeasia longifolia* | Duaneh, J. | 91 | K | Malaysia | 23936 | – | – | – | – | – | – | ERS5501741 |
| Sanger | Urticaceae | *Debregeasia longifolia* | WuZY | 9471 | KUN | China, Guangxi | – | – | KF137831 | KF138304 | KF138140 | KF138462 | KF137973 | – |
| Sanger | Urticaceae | *Debregeasia orientalis* |  | 81482 | KUN | China, Xizang | – | – | KF137834 | KF138307 | KF138143 | KF138465 | KF137976 | – |
| Sanger | Urticaceae | *Debregeasia saeneb* |  | 81107 | KUN | China, Xizang | – | – | KF137835 | KF138308 | KF138144 | KF138466 | KF137977 | – |
| Sanger | Urticaceae | *Debregeasia squamata* | WuZY | 9204 | KUN | Yunnan, China | – | – | KF137837 | KF138310 | KF138146 | KF138468 | KF137979 | – |
| Sanger | Urticaceae | *Dendrocnide excelsa* |  |  | E | Australia | – | – | – | MH358274 | – | – | MH357992 | – |
| Sanger | Urticaceae | *Dendrocnide meyeniana* | Yi 20111184 | 20111184 | KUN | China | – | KF137838 | – | KF138311 | KF138147 | KF138469 | KF137980 | – |
| Sanger | Urticaceae | *Dendrocnide sinuata* | WuZY | 9238 | KUN | China | – | KF137754 | KF137839 | KF138312 | KF138148 | KF138470 | KF137981 | – |
| Sanger | Urticaceae | *Dendrocnide sp.* | WuZY | 9035 | KUN | China | – | – | KF137840 | KF138313 | KF138149 | KF138471 | KF137982 | – |
| Angiosperms353 | Urticaceae | *Dendrocnide stimulans* | Beaman, J.H. | 10369 | K | Malaysia | 23908 | – | – | – | – | – | – | ERS4414128 |
| Sanger | Urticaceae | *Dendrocnide urentissima* | WuZY | 9211 | KUN | China | – | – | KF137841 | KF138314 | KF138150 | KF138472 | – | – |
| Angiosperms353 | Urticaceae | *Didymodoxa caffra* | Friis, I.; Gilbert, M.G.; Vollesen, K. | 3532 | K | – | – | – | – | – | – | – | – | ERS5502184 |
| Sanger | Urticaceae | *Didymodoxa caffra* | Abdallah, R. et al. | 96/92 | K | Ethiopia? | – | KF137755 | – | KF138315 | KF138151 | KF138473 | – | – |
| Angiosperms353 | Urticaceae | *Discocnide mexicana* | Manriquez I., G.; Esquivel Hernadez, K.B. | 6554 | K | – | – | – | – | – | – | – | – | ERS5501982 |
| Sanger | Urticaceae | *Discocnide mexicana* | Gereau *et al* . | 2205 | BM | Mexico | – | – | KF137842 | DQ179369 | KF138152 | KF138474 | KF137983 | – |
| Sanger | Urticaceae | *Droguetia ambigua* | Styles & Styles | 2426 | K | South Africa | – | KF137756 | KF137843 | KF138317 | AM235161 | KF138475 | – | – |
| Angiosperms353 | Urticaceae | *Droguetia iners* | Poulsen, A.D.; et al. | 1000 | K | Uganda | 23947 | – | – | – | – | – | – | ERS5501749 |
| Sanger | Urticaceae | *Droguetia iners* | Liuj | 10621 | KUN | China? | – | KF137757 | KF137844 | KF138318 | KF138154 | KF138476 | KF137984 | – |
|  |  |  |  |  |  | Bagvio. Beugoerc Prov.Lozon.3/1913 |  |  |  |  |  |  |  |  |
| Sanger | Urticaceae | *Elatostema polypodioides* |  | 1119052 | BM |  | – | – | – | – | – | – | – | – |
| Sanger | Urticaceae | *Elatostema acuminatum* | FRI | 29035 | K | Malagsia 22th mile Ginting→Simpsh road | – | – | – | – | – | – | – | – |
| Sanger | Urticaceae | *Elatostema acuteserratum* | Y. H. Tseng | 1121 | TAI | Taiwan | – | – | KP858860 | – | – | – | – | – |
| Sanger | Urticaceae | *Elatostema agusanense* |  | 38445 | BM | Phillipines.Mr.Bulu san,Sorsogon Prov,Luzon | – | – | – | – | – | – | – | – |
| Sanger | Urticaceae | *Elatostema albopilosoides* | Y. G. Wei & F. Wen | 1083 | IBK | China | – | – | KP858885 | – | – | – | – | – |
| Sanger | Urticaceae | *Elatostema androstachyum* | Y. G. Wei | g120 | IBK | China | – | – | KP858871 | – | – | – | – | – |
| Sanger | Urticaceae | *Elatostema angustum* |  | 13788 | BM | Phillipines Mt.Canumag,Rizal Prov.Luzon | – | – | To submit | – | – | – | – | – |
| Sanger | Urticaceae | *Elatostema angwesnm* |  | 5529 |  |  | – | – | – | – | – | – | – | – |
| Sanger | Urticaceae | *Elatostema apoeuse* |  | 11793 | BM | Phillipines.Lor? 6/7-1917 | – | – | – | – | – | – | – | – |
| Sanger | Urticaceae | *Elatostema asterocephalum* | Y. G. Wei | 623 | IBK | China | – | – | KP858878 | – | – | – | – | – |

| Sanger | Urticaceae | *Elatostema atroviride* |  | 36 |  | Sanceng Caue,Qiaotou Village Jialiang Toum,Libo County,Guizhou province | – | – | – | – | – | – | – | – |
| --- | --- | --- | --- | --- | --- | --- | --- | --- | --- | --- | --- | --- | --- | --- |
| Sanger | Urticaceae | *Elatostema attenuatoides* | Wei Y.G. | 76 | IBK | Dahuadi, Guangxi, China | – | – | – | – | To submit | – | To submit | – |
|  |  |  |  |  |  | Vietnam Cao Bang on lower Slqw lelow bamboo zow  on raks |  |  |  |  |  |  |  |  |
| Sanger | Urticaceae | *Elatostema attenuatoides* |  | 4336 | K |  | – | – | To submit | – | – | – | – | – |
|  |  |  |  |  |  | Phillipines Cabadbaraw,Mr.Ur danera,Agosan Prov,Mindanao |  |  |  |  |  |  |  |  |
| Sanger | Urticaceae | *Elatostema auronii* |  | 13894 | BM |  | – | – | – | – | – | – | – | – |
| Sanger | Urticaceae | *Elatostema backeri* | HNK | 2724 |  | Chu Village,Ta Sua, Bac Yen Son La,Vietnam | – | – | – | – | – | – | – | – |
| Sanger | Urticaceae | *Elatostema balansae* | A. K. Monro & Y. G. Wei | 6466 | IBK | China | – | – | KP858847 | – | – | – | – | – |
| Sanger | Urticaceae | *Elatostema banahaense* | C. I. Peng | 23770 | HAST | Philippines | – | – | KP858815 | – | – | – | – | – |
| Sanger | Urticaceae | *Elatostema banahaense* |  | 3680 | K | Thailand. Khao Choug | – | – | – | – | – | – | – | – |
| Sanger | Urticaceae | *Elatostema belense* |  | 11317 | BM | Phillipines.Baraan Prov.Lozon.  Mr.Mariveles 3/1911 | – | – | To submit | – | – | – | – | – |
| Sanger | Urticaceae | *Elatostema beugoereuje* |  | 14274 | BM | New Guinea | – | – | To submit | – | – | – | – | – |
| Sanger | Urticaceae | *Elatostema binatum* |  | 092 | IBK | Banbi, Longzhou County, Guangxi, China | – | – | To submit | – | To submit | – | To submit | – |
| Sanger | Urticaceae | *Elatostema binatum* | A. K. Monro & Y. G. Wei | 6663 | IBK | China | – | – | KP858895 | – | – | – | – | – |
| Sanger | Urticaceae | *Elatostema blechnoides* |  | 6085 | BM | New Guinea.Westem Highlands,Distntv, nr Tomba Village,S.of Mr.Hagen range | – | – | To submit | – | – | – | – | – |
| Sanger | Urticaceae | *Elatostema brachyodontum* | A. K. Monro & Y. G. Wei | 6724 | IBK | China | – | – | KP858888 | – | – | – | – | – |
| Sanger | Urticaceae | *Elatostema bsungletense* |  | 14274 |  |  | – | – | To submit | – | – | – | – | – |
| Sanger | Urticaceae | *Elatostema bulbiferum* |  | 33967 | K | Thailand.Kanchana bun;Erawan | – | – | To submit | – | – | – | – | – |
| Sanger | Urticaceae | *Elatostema bullatum* |  | 10928 | K | Borneo.Sabah.Rana u District nt Posing Hor Spningsi | – | – | – | – | – | – | – | – |
| Sanger | Urticaceae | *Elatostema bwlloifetum* |  | 4315 |  |  | – | – | To submit | – | – | – | – | – |
| Sanger | Urticaceae | *Elatostema calcareum* | R. C. M. Gregor | 432 | BM | Mariana Islands | – | – | KP858817 | – | – | – | – | – |

| Sanger | Urticaceae | *Elatostema calciferum* |  | 83 |  | Yeji Tuo,Limi Town,Yongshun County Hunan province | – | – | – | – | – | – | – | – |
| --- | --- | --- | --- | --- | --- | --- | --- | --- | --- | --- | --- | --- | --- | --- |
| Sanger | Urticaceae | *Elatostema catarctum* |  | 2002131 |  | Jialiang,Libo county,Guizhou province | – | – | To submit | – | – | – | – | – |
| Sanger | Urticaceae | *Elatostema celingense* | FLF | 37 | IBK | Lihu, Nandan County, Guangxi, China | – | – | To submit | – | To submit | – | To submit | – |
| Sanger | Urticaceae | *Elatostema circulosum* |  | 5547 |  | Phillipines.Mr.Mali nao Albng Prov.Lozon. | – | – | – | – | – | – | – | – |
| Sanger | Urticaceae | *Elatostema clemeusii* |  | 20788 | K | Malagsia Sarawak,Kapir,Upp er,Nejang River | – | – | To submit | – | – | – | – | – |
| Sanger | Urticaceae | *Elatostema coriaceifolium* |  | 61 |  | Shuirao village,Libo county Guizhou province | – | – | To submit | – | – | – | – | – |
|  |  |  |  |  |  | Xiao Xi National Natural Reserve,Yongshun County.Hunan province |  |  |  |  |  |  |  |  |
| Sanger | Urticaceae | *Elatostema cuspidatum* |  | 72 |  |  | – | – | To submit | – | – | – | – | – |
| Sanger | Urticaceae | *Elatostema cwpidatum* |  | 8850312 |  | E.Nepal,Koshi Zone,Sankhuwa sabhaDistr,Around Tashi Gaun(Tashigaom)( 2160-2340m) | – | – | – | – | – | – | – | – |
| Sanger | Urticaceae | *Elatostema cyrtandrifolium* | Y. H. Tseng | 1139 | TAI | Taiwan | – | – | KP858844 | – | – | – | – | – |
| Sanger | Urticaceae | *Elatostema dactgiccephcwm* |  | 23678 |  | Gaoagong shan | – | – | To submit | – | – | – | – | – |
| Sanger | Urticaceae | *Elatostema delicarum* |  | 10343 | BM | Phillipines Todaga Mr.Apo Davao District Mindano 9/1909 | – | – | To submit | – | – | – | – | – |
|  |  |  |  |  |  | The 4th Group of Shuichun Village Yuping Town,Libo County,Guizhou |  |  |  |  |  |  |  |  |
| Sanger | Urticaceae | *Elatostema discolor* |  | 59 |  |  | – | – | To submit | – | – | – | – | – |
| Sanger | Urticaceae | *Elatostema dissectum* | A. J. C. Grierson & D. G. Long | 393 | K | Bhutan | – | – | KP858832 | – | – | – | – | – |
| Sanger | Urticaceae | *Elatostema edule* | Y. H. Tseng | 1119 | TAI | Taiwan | – | – | KP858818 | – | – | – | – | – |
| Sanger | Urticaceae | *Elatostema ellipticum* | FKW | 7951 | K | India | – | – | – | – | – | – | – | – |
| Sanger | Urticaceae | *Elatostema fengshanense* | A. K. Monro & Y. G. Wei | 6651 | IBK | China | – | – | KP858848 | – | – | – | – | – |
| Sanger | Urticaceae | *Elatostema ficoides* |  | 1949 |  | Central Nepal No.1 west | – | – | To submit | – | – | – | – | – |

| Sanger | Urticaceae | *Elatostema filicinum* |  | 0006249871 | BM | New Euinea,Camp  I, | – | – | – | – | – | – | – | – |
| --- | --- | --- | --- | --- | --- | --- | --- | --- | --- | --- | --- | --- | --- | --- |
| Sanger | Urticaceae | *Elatostema garrettii* | P. Suvarnakoses | 38824 | K | Thailand | – | – | KP858849 | – | – | – | – | – |
| Sanger | Urticaceae | *Elatostema gibbsiae* |  | 11312 | K | Sabah.Ranau Disrncr. Mr Kinabalo.Liwago river vallege1700- 1800m. | – | – | – | – | – | – | – | – |
| Sanger | Urticaceae | *Elatostema glabrutum* |  | 16325 | BM | Phillipines.Mr.Bulu san,Sorsogon Prov,Luzon | – | – | To submit | – | – | – | – | – |
| Sanger | Urticaceae | *Elatostema glaucescens* |  | 18397 | BM | Phillipines.Mr.Polis | – | – | – | – | – | – | – | – |
| Sanger | Urticaceae | *Elatostema glochidiodes* |  | 2002121 |  | YongKang,Libo County,Guizhou province Lin-Dong | – | – | To submit | – | – | – | – | – |
| Sanger | Urticaceae | *Elatostema goudotianum* | S. Malcomber | 1723 | BM | Madagascar | – | – | KP858798 | – | – | – | – | – |
| Sanger | Urticaceae | *Elatostema grande* | I. R. Telfor | 10363 | K | Australia | – | – | KP858821 | – | – | – | – | – |
| Sanger | Urticaceae | *Elatostema grandidentatum* | A. J. C. Grierson & D. G. Long | 1999 | K | Bhutan | – | – | KP858833 | – | – | – | – | – |
| Sanger | Urticaceae | *Elatostema grandifolium* | K. L. Rechinger | 1780 | BM | Samoa | – | – | KP858823 | – | – | – | – | – |
| Sanger | Urticaceae | *Elatostema gyrocephalum* | Y. G. Wei | 7101 | IBK | China | – | – | KP858886 | – | – | – | – | – |
| Sanger | Urticaceae | *Elatostema hechiense* | A. K. Monro & Y. G. Wei | 6507 | IBK | China | – | – | KP858875 | – | – | – | – | – |
| Sanger | Urticaceae | *Elatostema hezhouense* | A. K. Monro & Y. G. Wei | 6820 | IBK | China | – | – | KP858881 | – | – | – | – | – |
| Sanger | Urticaceae | *Elatostema hirtellipedunculatum* | Y. H. Tseng | 1123 | TAI | Taiwan | – | – | KP858863 | – | – | – | – | – |
| Sanger | Urticaceae | *Elatostema holophgllum* |  | 23546 | BM | Sorsogou Pror,Luzon,Phillipi nes | – | – | To submit | – | – | – | – | – |
| Sanger | Urticaceae | *Elatostema hookerianum* | Y. G. Wei | 7 | IBK | China | – | – | KP858904 | – | – | – | – | – |
| Sanger | Urticaceae | *Elatostema hypoglaucum* | Y. H. Tseng | 1190 | TAI | Taiwan | – | – | KP858891 | – | – | – | – | – |
| Sanger | Urticaceae | *Elatostema ichangense* |  | 67 |  | Aojiahu Village,Lmoyixi Town Guzhang County,Hunan province | – | – | To submit | – | – | – | – | – |
| Sanger | Urticaceae | *Elatostema incisum* | G. E. Schatx et al. | 3459 | MO | Madagascar | – | – | KP858799 | – | – | – | – | – |
| Sanger | Urticaceae | *Elatostema inregnfolium* |  | 1359 | K | New Euinea.Morobe District.Aseki patrol area near Wengomanga(via Oiwa) | – | – | To submit | – | – | – | – | – |
| Sanger | Urticaceae | *Elatostema insulare* | A. C. Smith | 8145 | K | Fiji | – | – | KP858839 | – | – | – | – | – |
| Sanger | Urticaceae | *Elatostema integrifolium* | E. S. Brown | 1484 | BM | Solomon Islands | – | – | KP858852 | – | – | – | – | – |
| Sanger | Urticaceae | *Elatostema involucratum* | Tsugaru | 30727 | A | Japan | – | – | KP858866 | – | – | – | – | – |
| Sanger | Urticaceae | *Elatostema japonicum* | G. Murata | 72870 | KYO | Japan | – | – | KP858867 | – | – | – | – | – |

| Sanger | Urticaceae | *Elatostema jilicavle* |  | 7627 | BM | Phillipines. Dumagvere. Cuernos MH.Negros Oriemral Prov. Negror Island. 6/1908 | – | – | – | – | – | – | – | – |
| --- | --- | --- | --- | --- | --- | --- | --- | --- | --- | --- | --- | --- | --- | --- |
|  |  |  |  |  |  | Malagsia.Sabah,Kot a Belud District S of Sagap on NW side of Mt.Kinabalu |  |  |  |  |  |  |  |  |
| Sanger | Urticaceae | *Elatostema kabagense* |  | 9783 | K |  | – | – | To submit | – | – | – | – | – |
| Sanger | Urticaceae | *Elatostema kraemeri* | K. L. Rechinger | 1548 | BM | Samoa | – | – | KP858824 | – | – | – | – | – |
| Sanger | Urticaceae | *Elatostema laetevirens* | S. Tsugarul & T. Takahashi | 25903 | MO | Japan | – | – | KP858893 | – | – | – | – | – |
| Sanger | Urticaceae | *Elatostema laevissimum* | Y. G. Wei | 6393 | IBK | China | – | – | KP858861 | – | – | – | – | – |
| Sanger | Urticaceae | *Elatostema lagunense* |  | 17529 | BM | Phillipines.Los Baüos.Mr.Maguilin g,Laguna Prov,Luzon | – | – | To submit | – | – | – | – | – |
|  |  |  |  |  |  | Indonena,W Sumbawa,Mr.Barula nteh,trailfrom Batuctulang Pusu |  |  |  |  |  |  |  |  |
| Sanger | Urticaceae | *Elatostema lanafohum* |  | 2014 | BM |  | – | – | – | – | – | – | – | – |
|  |  |  |  |  |  | Indonesia.W.Sumba wa,Mr Batulantehtrail fnom Batudulang to Pwsu |  |  |  |  |  |  |  |  |
| Sanger | Urticaceae | *Elatostema lancifolium* |  | 4678 | K |  | – | – | – | – | – | – | – | – |
|  |  |  |  |  |  | Malagsia Sarawak Eat Upper Rejang river Bcre of Mr.Majan low elevnhvn |  |  |  |  |  |  |  |  |
| Sanger | Urticaceae | *Elatostema laxiflorum* |  | 20790 | K |  | – | – | – | – | – | – | – | – |
| Sanger | Urticaceae | *Elatostema lineare* | J & M. S. Clemens | 50589 | BM | Malaysia | – | – | KP858835 | – | – | – | – | – |
| Sanger | Urticaceae | *Elatostema lineolatum* | Y. H. Tseng | 1108 | TAI | Taiwan | – | – | KP858855 | – | – | – | – | – |
| Sanger | Urticaceae | *Elatostema lithoneurum* | J & M. S. Clemens | s.n. | BM | Malaysia | – | – | KP858827 | – | – | – | – | – |
|  |  |  |  |  |  | Cao Bang,Tra Tinh District Quoc Toan Municipality;Trang Hem(Lakes  District) |  |  |  |  |  |  |  |  |
| Sanger | Urticaceae | *Elatostema longibracteatum* |  | 4288 | K |  | – | – | To submit | – | – | – | – | – |
| Sanger | Urticaceae | *Elatostema longistipulum* | A. K. Monro & Y. G. Wei | 6462 | IBK | China | – | – | KP858856 | – | – | – | – | – |
| Sanger | Urticaceae | *Elatostema lungzhouense* | A. K. Monro & Y. G. Wei | 6786 | IBK | China | – | – | KP858868 | – | – | – | – | – |
| Sanger | Urticaceae | *Elatostema lutescens* | C. B. Robinson | 14004 | BM | Philippines | – | – | KP858816 | – | – | – | – | – |
| Sanger | Urticaceae | *Elatostema macintyrei* | A. K. Monro & Y. G. Wei | 6743 | IBK | China | – | – | KP858850 | – | – | – | – | – |

| Sanger | Urticaceae | *Elatostema macnphllium* |  | 001119057 |  | New Euinea,A.F.R.Wolla sta Expeditio Camp V16 | – | – | To submit | – | – | – | – | – |
| --- | --- | --- | --- | --- | --- | --- | --- | --- | --- | --- | --- | --- | --- | --- |
| Sanger | Urticaceae | *Elatostema madagascariense* | Richard Razakamalala et al. | 3199 | MO | Africa | – | – | KP858800 | – | – | – | – | – |
| Sanger | Urticaceae | *Elatostema malacotrichum* | Y. G. Wei | 13 | IBK | China | – | – | KP858876 | – | – | – | – | – |
| Sanger | Urticaceae | *Elatostema megacephalum* |  | 1368 | K | Chingmai,Doi Ka | – | – | To submit | – | – | – | – | – |
| Sanger | Urticaceae | *Elatostema microcephalanthum* | Y. H. Tseng | 1100 | TAI | Taiwan | – | – | KP858874 | – | – | – | – | – |
| Sanger | Urticaceae | *Elatostema molle* |  | 262 | K | Peninsular:Narathi  wat Bacho | – | – | – | – | – | – | – | – |
| Sanger | Urticaceae | *Elatostema monandrum* | F. Miyamoto et al. | 9588281 | MO | Nepal | – | – | KP858905 | – | – | – | – | – |
| Sanger | Urticaceae | *Elatostema monticola* | H. A. Osmaston | 1952 | BM | Congo | – | – | KP858803 | – | – | – | – | – |
| Sanger | Urticaceae | *Elatostema morobense* | R. J. Johns | 9209A | K | Indonesia | – | – | KP858820 | – | – | – | – | – |
| Sanger | Urticaceae | *Elatostema multicaule* | A. K. Monro & Y. G. Wei | 6748 | IBK | China | – | – | KP858884 | – | – | – | – | – |
| Sanger | Urticaceae | *Elatostema myrtillus* | Y. G. Wei | 44937 | IBK | China | – | – | KP858880 | – | – | – | – | – |
| Sanger | Urticaceae | *Elatostema nacuturn* |  | 9338 |  | Aalung,Guizhou Y.Tsiang | – | – | To submit | – | – | – | – | – |
| Sanger | Urticaceae | *Elatostema nalkerae* |  | 20916 | K | Laos.Tawrieng,Che ng Kwang | – | – | To submit | – | – | – | – | – |
|  |  |  |  |  |  | Baibi Cave,Qiaotoy Village Jialiang Town,Libo County Guizhou province |  |  |  |  |  |  |  |  |
| Sanger | Urticaceae | *Elatostema nanchuanense* |  | 38 |  |  | – | – | To submit | – | – | – | – | – |
| Sanger | Urticaceae | *Elatostema nasutum* | A. K. Monro & Y. G. Wei | 6401 | IBK | China | – | – | KP858902 | – | – | – | – | – |
| Sanger | Urticaceae | *Elatostema nrangii* |  | 775 |  |  | – | – | To submit | – | – | – | – | – |
| Sanger | Urticaceae | *Elatostema oblongifolium* | A. K. Monro & Y. G. Wei | 6713 | IBK | China | – | – | KP858897 | – | – | – | – | – |
| Sanger | Urticaceae | *Elatostema obtusum* | Y. H. Tseng | 1191 | TAI | Taiwan | – | – | KP858900 | – | – | – | – | – |
| Sanger | Urticaceae | *Elatostema obtusum* | Bouffordet al | al.27415 | A | China | – | – | KP858901 | – | – | – | – | – |
|  |  |  |  |  |  | Indonesia.Sumarm, Dëlëag Singkoet,N of Bërasvagl,Karo Plareau |  |  |  |  |  |  |  |  |
| Sanger | Urticaceae | *Elatostema oglvnnum* |  | 8571 | K |  | – | – | – | – | – | – | – | – |
|  |  |  |  |  |  | Phillipines Luzon.Sorrogon Mr Bulusan about lake Agangag on stores |  |  |  |  |  |  |  |  |
| Sanger | Urticaceae | *Elatostema oimulans* |  | 9654 | K |  | – | – | – | – | – | – | – | – |
| Sanger | Urticaceae | *Elatostema oligophlebium* | Y. G. Wei | g075 | IBK | China | – | – | KP858877 | – | – | – | – | – |
| Sanger | Urticaceae | *Elatostema orienlau* |  | 17156 |  |  | – | – | – | – | – | – | – | – |
| Sanger | Urticaceae | *Elatostema orientale* | C. F. M. Synnerton | 784 | BM | Zimbabwe | – | – | KP858802 | – | – | – | – | – |
| Sanger | Urticaceae | *Elatostema paivaeanum* | Charles Doumenge | 456 | MO | Africa | – | – | KP858801 | – | – | – | – | – |
| Sanger | Urticaceae | *Elatostema paracuminatw* |  | 21016 | K | pu Bia Laos | – | – | – | – | – | – | – | – |

|  |  |  |  |  |  | Baibi care,Qiaotou Village,Jialiang Town,Libo county,Guizhou  province |  |  |  |  |  |  |  |  |
| --- | --- | --- | --- | --- | --- | --- | --- | --- | --- | --- | --- | --- | --- | --- |
| Sanger | Urticaceae | *Elatostema parvum* |  | 41 |  |  | – | – | To submit | – | – | – | – | – |
| Sanger | Urticaceae | *Elatostema parvum* | Y. H. Tseng | 1182 | TAI | Taiwan | – | – | KP858794 | – | – | – | – | – |
| Sanger | Urticaceae | *Elatostema pedicillatum* |  | 31355 | BM | Borneo Mr Kinabalu | – | – | To submit | – | – | – | – | – |
|  |  |  |  |  |  | New Euinea.Indouesra. Mimika Regeng,PF Freenort Indonessa Concesne area:main road 34 mile W side of Aikwa bridge |  |  |  |  |  |  |  |  |
| Sanger | Urticaceae | *Elatostema peltijolium* |  | 290 | K |  | – | – | To submit | – | – | – | – | – |
| Sanger | Urticaceae | *Elatostema penibukanense* | J. H. Beaman | 10466 | K | Malaysia | – | – | KP858854 | – | – | – | – | – |
| Sanger | Urticaceae | *Elatostema peniwkanense* |  | 29737 | BM | Borneo Mr Kinabalu ilau Basin | – | – | – | – | – | – | – | – |
| Sanger | Urticaceae | *Elatostema pennineive* |  | 2430 | K | Biunei Temburong. Batu Apoi;Bukir Galagas | – | – | – | – | – | – | – | – |
|  |  |  |  |  |  | Lang Son Húu Lung District Húu Lien Municipa city Húu Lien proctected Area  between Peo Dat |  |  |  |  |  |  |  |  |
| Sanger | Urticaceae | *Elatostema pergameneom* |  | 4121 | K |  | – | – | – | – | – | – | – | – |
| Sanger | Urticaceae | *Elatostema pergameneum* |  | 07298 | IBK | Sanlian, Longzhou County, Guangxi, China | – | – | To submit | – | To submit | – | To submit | – |
| Sanger | Urticaceae | *Elatostema perlongfolium* |  | 16408 | BM | Phillipines Irobin.Mf.Bulasan.S orsogon Prov.Luzon | – | – | – | – | – | – | – | – |
| Sanger | Urticaceae | *Elatostema phillipinense* |  | 6907 | K | Phillipines.Negros. Canlaon Volcauo | – | – | To submit | – | – | – | – | – |
| Sanger | Urticaceae | *Elatostema pianmaense* |  | 24077 |  | Gaoagong shan | – | – | To submit | – | – | – | – | – |
| Sanger | Urticaceae | *Elatostema pinnatum* | J & M. S. Clemens | s.n. | BM | Malaysia | – | – | KP858836 | – | – | – | – | – |
| Sanger | Urticaceae | *Elatostema platyphyllum* | Y. H. Tseng | 1158 | TAI | Taiwan | – | – | KC420490 | – | – | – | – | – |
| Sanger | Urticaceae | *Elatostema podohglum* |  | 19714 | BM | Phillipines  ,Mr.Polos Ifugo Sub prov,Luzoni | – | – | To submit | – | – | – | – | – |

|  |  |  |  |  |  | Huangcao ping,Yaolu Town Libo  County,Guizhou |  |  |  |  |  |  |  |  |
| --- | --- | --- | --- | --- | --- | --- | --- | --- | --- | --- | --- | --- | --- | --- |
| Sanger | Urticaceae | *Elatostema pseudissectum* |  | 28 |  |  | – | – | To submit | – | – | – | – | – |
| Sanger | Urticaceae | *Elatostema pusillum* | R. Bedi | 931 | K | Bhutan | – | – | KP858906 | – | – | – | – | – |
| Sanger | Urticaceae | *Elatostema pycndontum* |  | 39 |  | W.T.Wang Baibi Cave Qiaotou Village.Jialiang Town,Libo County.Guizhou province | – | – | To submit | – | – | – | – | – |
| Sanger | Urticaceae | *Elatostema ramosum* | AM | 6800 | IBK | Banbi, Longzhou County, Guangxi, China | – | – | To submit | – | To submit | – | To submit | – |
| Sanger | Urticaceae | *Elatostema reticulatum* | P. I. Forster | 7564 | K | Australia | – | – | KP858834 | – | – | – | – | – |
| Angiosperms353 | Urticaceae | *Elatostema retrohirtum* | Monro, A.K.; et al. | 7601 | K | – | – | – | – | – | – | – | – | ERS4414235 |
| Sanger | Urticaceae | *Elatostema retrohirtum* |  | 236 |  | Doi Chuangdao,Siam,Ta iland 1921.1.4 1500 1800m | – | – | To submit | – | – | – | – | – |
| Sanger | Urticaceae | *Elatostema rubrosvipulatum* |  | 11322 | K | Sabah Nanav Dishnr,Mr Kinabalu Liwago Riov | – | – | – | – | – | – | – | – |
| Sanger | Urticaceae | *Elatostema rugosum* | Hooker | s.n. | BM | New Zealand | – | – | – | – | – | – | – | – |
| Sanger | Urticaceae | *Elatostema rupestre* |  | 245 |  | Chuia Chuli | – | – | – | – | – | – | – | – |
| Sanger | Urticaceae | *Elatostema salvinioides* |  | 4215 | K | Northern Thailand Chiangmai Doi chingdao | – | – | – | – | – | – | – | – |
| Sanger | Urticaceae | *Elatostema samoense* | K. L. Rechinger | s.n. | BM | Samoa | – | – | KP858826 | – | – | – | – | – |
|  |  |  |  |  |  | New Euinea.Morobe Distncr 5m Sw of Wagau Mr.Shungol |  |  |  |  |  |  |  |  |
| Sanger | Urticaceae | *Elatostema schroeteri* |  | 12474 | K |  | – | – | – | – | – | – | – | – |
| Sanger | Urticaceae | *Elatostema serra* | W. Takeuchi et al. | 19898 | MO | Papua New Guinea | – | – | KP858837 | – | – | – | – | – |
| Sanger | Urticaceae | *Elatostema sessile* |  |  | K | Indowesra.Jara | – | – | To submit | – | – | – | – | – |
| Sanger | Urticaceae | *Elatostema setulosum* | Y. G. Wei | g124 | IBK | China | – | – | KP858872 | – | – | – | – | – |
| Sanger | Urticaceae | *Elatostema sinense* | Q. Shao & L. D. Duan | 76 | BM | China | – | – | KP858795 | – | – | – | – | – |
| Sanger | Urticaceae | *Elatostema sinense* | Q. Shao & L. D. Duan | 1 | BM, PE | China | – | – | KP858796 | – | – | – | – | – |
| Sanger | Urticaceae | *Elatostema sinense* | T. T. Chen | s.n. | ? | China | – | – | KP858797 | – | – | – | – | – |
| Sanger | Urticaceae | *Elatostema sinopurpureum* | Y. G. Wei & F. Wen | 1081 | IBK | China | – | – | KP858869 | – | – | – | – | – |
|  |  |  |  |  |  | Phillipines.Irosin Mt.Bulusan,Sorsog on Prov,Luzon |  |  |  |  |  |  |  |  |
| Sanger | Urticaceae | *Elatostema sorsogoncnse* |  | 16322 | BM |  | – | – | To submit | – | – | – | – | – |

| Sanger | Urticaceae | *Elatostema sp.* |  | 143234 | K | Phillipines Luzon.Laguna Pioviuce,Mr. Makiling | – | – | – | – | – | – | – | – |
| --- | --- | --- | --- | --- | --- | --- | --- | --- | --- | --- | --- | --- | --- | --- |
| Sanger | Urticaceae | *Elatostema sterma* |  | 959 |  | Hendcian, | – | – | To submit | – | – | – | – | – |
|  |  |  |  |  |  | Xiao Xi National Natural Reserve,Yongshun County.Hunan  province |  |  |  |  |  |  |  |  |
| Sanger | Urticaceae | *Elatostema stewardii* |  | 77 |  |  | – | – | To submit | – | – | – | – | – |
| Sanger | Urticaceae | *Elatostema strictum* | K. L. Rechinger | 388 | BM | Samoa | – | – | KP858825 | – | – | – | – | – |
| Sanger | Urticaceae | *Elatostema strigillosum* | C. I. Peng | 23775 | HAST | Philippines | – | – | KP858814 | – | – | – | – | – |
| Sanger | Urticaceae | *Elatostema subcoriaceum* | Y. H. Tseng | 1127 | TAI | Taiwan | – | – | KP858894 | – | – | – | – | – |
| Sanger | Urticaceae | *Elatostema sublineare* | A. K. Monro & Y. G. Wei | 6686 | IBK | China | – | – | KP858882 | – | – | – | – | – |
| Sanger | Urticaceae | *Elatostema surigaeense* |  | 9857 | BM | Phillipines Mt.Banajao,Laguna Prov,Luzon | – | – | To submit | – | – | – | – | – |
| Sanger | Urticaceae | *Elatostema suzukii* | Yasuda | 5170 | KYO | Japan | – | – | KP858865 | – | – | – | – | – |
| Sanger | Urticaceae | *Elatostema tenuicaudatum* | Y. G. Wei & F. Wen | 1050 | IBK | China | – | – | KP858862 | – | – | – | – | – |
| Sanger | Urticaceae | *Elatostema tenuinerve* | A. K. Monro & Y. G. Wei | 6502 | IBK | China | – | – | KP858896 | – | – | – | – | – |
| Sanger | Urticaceae | *Elatostema thalictroides* | J. H. Beaman | 7650 | MO | Malaysia | – | – | KP858838 | – | – | – | – | – |
| Sanger | Urticaceae | *Elatostema tianeense* | A. K. Monro & Y. G. Wei | 6491 | IBK | China | – | – | KP858879 | – | – | – | – | – |
|  |  |  |  |  |  | Papua New Euinea W.Highlands Prov,Bismarck range,Mt.Oibo |  |  |  |  |  |  |  |  |
| Sanger | Urticaceae | *Elatostema tndens* |  | 10475 | K |  | – | – | – | – | – | – | – | – |
| Sanger | Urticaceae | *Elatostema toidesaustrale* | A. C. Smith | 8131 | K | Fiji | – | – | KP858790 | – | – | – | – | – |
| Sanger | Urticaceae | *Elatostema toidesfilicoides* | H. B. R. Parham | 235a | BM | Fiji | – | – | KP858789 | – | – | – | – | – |
| Sanger | Urticaceae | *Elatostema toidesfruticulosum* | L. S. Brass | 2607 | BM | Solomon Island | – | – | KP858788 | – | – | – | – | – |
| Sanger | Urticaceae | *Elatostema toideslonchophyllum* | J. & M. S. Clemens | 20816 | K | Malaysia | – | – | KP858793 | – | – | – | – | – |
| Sanger | Urticaceae | *Elatostema toidesvariolaminosum var. latum* | Rena George | S38983 | K | Malaysia | – | – | KP858791 | – | – | – | – | – |
| Sanger | Urticaceae | *Elatostema villosum* | Y. H. Tseng | 1275 | TAI | Taiwan | – | – | KP858812 | – | – | – | – | – |
| Sanger | Urticaceae | *Elatostema viridescens* |  | 15371 | BM | Phillipines.Irosin.Pr ov.Sorsogon.Mr.Bul usan.Luzon Island | – | – | To submit | – | – | – | – | – |
| Sanger | Urticaceae | *Elatostema vittatum* | J. H. Beaman | 10976 | K | Malaysia | – | – | KP858792 | – | – | – | – | – |
| Sanger | Urticaceae | *Elatostema welwitschii* | Welwitsch | 6270 | BM | Angola | – | – | KP858841 | – | – | – | – | – |
| Sanger | Urticaceae | *Elatostema xanthophyllum* | Y. G. Wei & V.T. Do | VMN_CN2 4 | IBK | Vietnam | – | – | KP858890 | – | – | – | – | – |
| Sanger | Urticaceae | *Elatostema yachense* | A. K. Monro & Y. G. Wei | 6725 | IBK | China | – | – | KP858883 | – | – | – | – | – |
| Sanger | Urticaceae | *Elatostema yakushimense* | Murata et al. | 15490 | A | Japan | – | – | KP858892 | – | – | – | – | – |
| Sanger | Urticaceae | *Elatostema yaoshanense* | F. Wen | 070509A | IBK | China | – | – | KP858903 | – | – | – | – | – |
| Angiosperms353 | Urticaceae | *Elatostematoides vittatum* | Beaman, J.H. | 10976 | K | – | – | – | – | – | – | – | – | ERS5503005 |
| Angiosperms353 / Sanger | Urticaceae | *Forsskaolea angustifolia* | Chase, M.W. | 16132 | K | Spain | 15957 | KF137762 | KF137861 | KF138334 | KF138170 | KF138493 | – | ERS5501729 |
| Sanger | Urticaceae | *Forsskaolea candida* | unknown collector | 3193 | US | Namibia | – | – | KM586475 | KM586647 | KM586561 | MK955181 | – | – |
| Angiosperms353 | Urticaceae | *Gesnouinia arborea* | Wildpret, W.; et al. | s.n. | K | Spain | 23944 | – | – | – | – | – | – | ERS5501746 |
| Sanger | Urticaceae | *Gesnouinia arborea* | C. Evrard | 12088 | BM | Spain, Canary Islands | – | – | KF137862 | DQ179372 | KF138172 | KF138494 | KF138000 | – |
| Angiosperms353 | Urticaceae | *Gibbsia insignis* | Sands, M.J.S. | 7149 | K | Indonesia | 23931 | – | – | – | – | – | – | ERS5501738 |

| Angiosperms353 | Urticaceae | *Girardinia diversifolia* | Abdallah, R.; et al. | 9689 | K | Tanzania, the United Republic of | 23917 | – | – | – | – | – | – | ERS5501731 |
| --- | --- | --- | --- | --- | --- | --- | --- | --- | --- | --- | --- | --- | --- | --- |
| Sanger | Urticaceae | *Girardinia diversifolia subsp. diversifolia* | Liuj | 10776 | KUN | China | – | KF137763 | KF137863 | KF138337 | KF138174 | KF138496 | KF138001 | – |
| Sanger | Urticaceae | *Girardinia diversifolia subsp. suborbiculata* | WuZY | 10011 | KUN | China | – | – | – | KF138338 | KF138175 | KF138497 | KF138003 | – |
| Sanger | Urticaceae | *Girardinia diversifolia subsp. triloba* | WuZY | 10229 | KUN | China | – | – | – | KF138341 | KF138178 | KF138500 | KF138006 | – |
| Angiosperms353 | Urticaceae | *Gonostegia hirta* | Monro, A.K.; Fu, L.F. | 7504 | K | – | – | – | – | – | – | – | – | ERS5502208 |
| Sanger | Urticaceae | *Gonostegia hirta* | WuZY | 9478 | KUN | Guangxi, China | – | – | KF137865 | KF138342 | KF138179 | – | KF138007 | – |
| Sanger | Urticaceae | *Gonostegia parvifolia* | Liuj | 10649 | KUN | Fujian, China | – | – | KF137867 | KF138344 | KF138181 | KF138502 | KF138009 | – |
| Angiosperms353 | Urticaceae | *Gyrotaenia myriocarpa* | Ekman, E.L. | 7356 | K | Haiti | 23915 | – | – | – | – | – | – | ERS5501730 |
| Sanger | Urticaceae | *Gyrotaenia myriocarpa* | P. AcevedoRdgz. | 12971 | US | Dominican Republic | – | – | KM586472 | KM586644 | KM586558 | – | – | – |
| Angiosperms353 | Urticaceae | *Haroldiella rapaensis* | Perlman | 18115 | K | – | – | – | – | – | – | – | – | ERS5503008 |
| Sanger | Urticaceae | *Hemistylus boehmerioides* | Schimpff | 445 | G | Ecuador | – | – | – | – | – | – | – | – |
| Angiosperms353 / Sanger | Urticaceae | *Hemistylus macrostachya* | Pittier | 11788 | K | Venezuela (Bolivarian Republic of) | 23946 | – | KF137868 | KF138346 | KF138182 | KF138503 | – | ERS5501748 |
| Sanger | Urticaceae | *Hesperocnide sandwicensis* | J.B. Gillett | 16765 | EA | Kenya | – | – | KM586429 | KM586601 | KM586515 | MK955193 | MK931209 | – |
| Angiosperms353 | Urticaceae | *Hesperocnide tenella* | Nuttall, L.W. | 227 | K | United States of  America | 23920 | – | – | – | – | – | – | ERS5501733 |
| Sanger | Urticaceae | *Hesperocnide tenella* | B. Trusk | 188 | EA | USA | – | – | KC284967 | KC285019 | KC284993 | – | – | – |
| Sanger | Urticaceae | *Laportea aestuans* | Peterson | 7165 | US | Panama | – | – | KM586464 | KM586636 | – | – | – | – |
| Angiosperms353 | Urticaceae | *Laportea canadensis* | Bourdo Jr., E.A. | 29633 | K | – | – | – | – | – | – | – | – | ERS4414133 |
| Angiosperms353 | Urticaceae | *Laportea cuspidata* | Furuse, M. | 18489 | K | – | – | – | – | – | – | – | – | ERS4414135 |
| Angiosperms353 | Urticaceae | *Laportea floribunda* | Lowry II, P.P. | 4529 | K | – | – | – | – | – | – | – | – | ERS5503068 |
| Angiosperms353 | Urticaceae | *Laportea humblotii* | Schatz, G.E.; Rakotozafy, A.; D'Arcy W.; Randrianasolo, J. | 1510 | K | – | – | – | – | – | – | – | – | ERS5503069 |
| Sanger | Urticaceae | *Laportea interrupta* | Festo & Luke | 2502 | EA | Kenya | – | – | KM586445 | KM586617 | – | – | – | – |
| Sanger | Urticaceae | *Laportea peduncularis* | Weigend | 8713 | BSB | South Africa | – | – | KF559047 | KF558927 | – | – | – | – |
| Angiosperms353 | Urticaceae | *Laportea ruderalis* | Fosberg, F.R. | 55756 | K | – | – | – | – | – | – | – | – | ERR3672054 |
| Sanger | Urticaceae | *Laportea ruderalis* | Hunt | 6 | US | Micronesia | – | – | KM586461 | KM586633 | – | – | – | – |
| Angiosperms353 | Urticaceae | *Lecanthus peduncularis* | Johns, R.J. | 10035 | K | Indonesia | 23921 | – | – | – | – | – | – | ERS5501734 |
| Sanger | Urticaceae | *Lecanthus peduncularis* | Liuj | 10607 | KUN | China | – | KF137768 | KF137871 | KF138350 | KF138186 | KF138508 | KF138016 | – |
| Sanger | Urticaceae | *Lecanthus_petelotii* | Anon | 81066 | KUN | China | – | – | KF137873 | – | – | – | – | – |
| Sanger | Urticaceae | *Leucosyke australis* | J.I. Wheatley | 864 | K | Vanuatu | – | – | – | – | – | – | – | – |
| Angiosperms353 | Urticaceae | *Leucosyke capitellata* | Lugas, L. | 41 | K | Malaysia | 23930 | – | – | – | – | – | – | ERS4414208 |
| Sanger | Urticaceae | *Leucosyke quadrinervia* | WuZY | 2012438 | KUN | Taiwan | – | – | KF137875 | KF138354 | KF138190 | – | – | – |
| Angiosperms353 | Urticaceae | *Maoutia puya* | Eanghourt, Khou et al. | FFI-PA203 | K | Cambodia | 23932 | – | – | – | – | – | – | ERS5503002 |
| Sanger | Urticaceae | *Maoutia setosa* | WuZY | 2012453 | KUN | Taiwan | – | – | MH357903 | MH358290 | MH358097 | MH358211 | – | – |
| Angiosperms353 | Urticaceae | *Musanga cecropioides* | Maurin, O. | 4374 | K | – | 74208 | – | – | – | – | – | – | ERS5501755 |
| Sanger | Urticaceae | *Musanga cecropioides* | I. Friis et al. | 9427 | K | Ethiopia | – | – | – | – | – | – | – | – |
| Sanger | Urticaceae | *Myrianthus holstii* | D.B. Faushawe | 5150 | K | Zimbabwe | – | – | – | – | – | – | – | – |
| Angiosperms353 | Urticaceae | *Myrianthus preussii* | Bogner | 672 | K | Gabon | 23953 | – | – | – | – | – | – | ERS5501751 |
| Sanger | Urticaceae | *Myriocarpa cordifolia* | Monro 4630 | 4630 | BM | Panama | – | KF137770 | KF137877 | KF138357 | KF138193 | KF138512 | – | – |
| Angiosperms353 / Sanger | Urticaceae | *Myriocarpa obovata* | Monro & Penn | 6534 | K | Belize | – | KF137771 | KF137878 | KF138358 | – | KF138513 | KF138021 | ERS5502207 |
| Angiosperms353 | Urticaceae | *Nanocnide japonica* | Im, H.T.; et al. | 6680 | K | Japan | 23919 | – | – | – | – | – | – | ERS5501732 |
| Sanger | Urticaceae | *Nanocnide japonica* | Liuj | 10735 | KUN | China | – | KF137772 | KF137879 | KF138359 | KF138194 | KF138514 | KF138022 | – |
| Sanger | Urticaceae | *Nanocnide lobata* | Liuj | 10799 | KUN | China | – | KF137773 | KF137881 | KF138361 | KF138196 | KF138516 | KF138024 | – |
| Sanger | Urticaceae | *Neodistemon indicus* | Larsen et al. | 2669 | K | Thailand | – | – | – | KF138363 | KF138198 | – | KF138026 | – |
| Angiosperms353 | Urticaceae | *Neraudia angulata* | Cowan, R.S. | 758 | K | United States of America | 23940 | – | – | – | – | – | – | ERS5501744 |
| Sanger | Urticaceae | *Neraudia angulata* | Je | 2012-1 | KUN | Hawaii, USA | – | – | MH357910 | MH358291 | MH358105 | MH358212 | MH358008 | – |

| Sanger | Urticaceae | *Neraudia kauaiensis* |  | 678003 | KUN | Hawaii, USA | – | – | KF137883 | KF138364 | KF138199 | – | KF138027 | – |
| --- | --- | --- | --- | --- | --- | --- | --- | --- | --- | --- | --- | --- | --- | --- |
| Sanger | Urticaceae | *Neraudia melastomifolia* |  | 90861 | KUN | Hawaii, USA | – | – | KF137884 | KF138365 | KF138200 | KF138518 | KF138028 | – |
| Angiosperms353 / Sanger | Urticaceae | *Nothocnide mollissima* | Beaman, J.H. | 8882 | K | Malaysia | 23934 | – | KF137885 | KF138366 | KF138201 | KF138519 | – | ERS5501739 |
| Sanger | Urticaceae | *Obetia aldabrensis* | Renvoize | 1357 | US | Seychelles | – | – | KM586460 | KM586632 | – | – | – | – |
| Sanger | Urticaceae | *Obetia pinnatifida* | Greenway | 11371 | EA | Tanzania | – | – | KM586449 | KM586621 | – | – | – | – |
| Angiosperms353 | Urticaceae | *Obetia radula* | Greenway, P.J.; Kanuri | 11371 | K | Tanzania, the United Republic of | 23918 | – | – | – | – | – | – | ERS4414129 |
| Sanger | Urticaceae | *Obetia radula* | Deng | 700 | KUN | Kenya | – | – | KM586431 | KM586603 | – | – | – | – |
| Sanger | Urticaceae | *Obetia tenax* | Botha | Botha_9 | K | S Africa | – | – | KF137886 | KF138367 | – | – | – | – |
| Sanger | Urticaceae | *Oreocnide boniana* |  |  | KUN | Yunnan, China | – | – | – | – | MH358108 | – | – | – |
| Sanger | Urticaceae | *Oreocnide frutescens* | Liuj | 10623 | KUN | Yunnan, China | – | – | KF137887 | KF138368 | KF138203 | KF138521 | KF138029 | – |
| Sanger | Urticaceae | *Oreocnide frutescens subsp. occidentalis* | WuZY | 9260 | KUN | Yunnan, China | – | – | – | KF138370 | KF138205 | KF138523 | KF138031 | – |
| Sanger | Urticaceae | *Oreocnide pedunculata* | WuZY | 2012485 | KUN | Taiwan, China | – | – | MH357912 | MH358293 | MH358109 | MH358213 | MH358009 | – |
| Sanger | Urticaceae | *Oreocnide rubescens* | WuZY | 9234 | KUN | Yunnan, China | – | – | MH357913 | – | MH358111 | – | – | – |
| Sanger | Urticaceae | *Oreocnide tonkinensis var. discolor* |  |  | KUN | Yunnan, China | – | – | MH357914 | MH358295 | MH358112 | MH358215 | – | – |
| Angiosperms353 | Urticaceae | *Oreocnide trinervis* | Andau, D. | 2015 | K | Malaysia | 23935 | – | – | – | – | – | – | ERS5501740 |
| Sanger | Urticaceae | *Oreocnide trinervis* | WuZY | WuZY2012 474 | KUN | Taiwan, China | – | – | – | – | MH358113 | – | – | – |
|  |  |  |  |  |  | United Kingdom of Great Britain and Northern Ireland  (the) |  |  |  |  |  |  |  |  |
| Angiosperms353 / Sanger | Urticaceae | *Parietaria judaica* | Fay, M.F. | 185 | K |  | 8071 | KF137779 | – | KF138371 | KF138206 | KF138524 | – | ERS5501728 |
| Sanger | Urticaceae | *Parietaria micrantha* | WuZY | 10373 | KUN | China | – | KF137780 | – | KF138372 | KF138207 | KF138525 | – | – |
| Sanger | Urticaceae | *Pellionia acutidentata* | Q. Shao & L. D. Duan | 62 | BM | China | – | – | KP858777 | – | – | – | – | – |
| Sanger | Urticaceae | *Pellionia brachyceras* | Wei Y.G. | 81 | IBK | Dahuadi, Guangxi,  China | – | – | To submit | – | To submit | – | To submit | – |
| Sanger | Urticaceae | *Pellionia grijsii* | Y. H. Tseng | 1167 | TAI | China | – | – | KC420491 | – | – | – | – | – |
| Sanger | Urticaceae | *Pellionia heteroloba* | A. K. Monro & Y. G. Wei | 6459 | IBK | China | – | – | KP858806 | – | – | – | – | – |
| Sanger | Urticaceae | *Pellionia minima* | J. M. Hu | 1787 | TAI | Japan | – | – | KP858809 | – | – | – | – | – |
| Sanger | Urticaceae | *Pellionia radicans* |  | HSL113 | KUN | China | – | – | KF137891 | KF138375 | KF138210 | – | – | – |
| Sanger | Urticaceae | *Pellionia radicans* | Y. G. Wei | U | IBK | China | – | – | KP858811 | – | – | – | – | – |
| Sanger | Urticaceae | *Pellionia repens* | L. F. Fu & S. L. Huang FL0071 | FL0071 | IBK | China | – | – | KU161129 | – | – | – | – | – |
| Sanger | Urticaceae | *Pellionia retrohispida* | M. H. Li | 6 | BM | China | – | – | KP858808 | – | – | – | – | – |
| Sanger | Urticaceae | *Pellionia scabra* | Y. H. Tseng | 1224 | TAI | Taiwan | – | – | KC420492 | – | – | – | – | – |
| Sanger | Urticaceae | *Pellionia tsoongii* | WuZY | 9495 | KUN | China | – | – | KF137893 | KF138377 | KF138212 | – | – | – |
| Sanger | Urticaceae | *Pellionia viridis* |  | 5155 |  | Mt.OMI Westerm  china | – | – | To submit | – | – | – | – | – |
| Sanger | Urticaceae | *Pellionia viridis* | H. G. Xu | 1995364 | MO | China | – | – | KP858805 | – | – | – | – | – |
| Angiosperms353 / Sanger | Urticaceae | *Phenax ballotifolius* | Wood, J.R.I.; et al. | 18048 | K | Bolivia (Plurinational State  of) | 23942 | – | To submit | – | – | – | To submit | ERS5503003 |
| Sanger | Urticaceae | *Phenax hirtus* | C. R. Romero | COL000134 982 | COL | Colombia | – | – | – | – | – | – | MH151318 | – |
| Sanger | Urticaceae | *Phenax mexicanus* |  | XZB102 | E | Peru | – | – | MH357916 | MH358299 | – | MH358218 | – | – |
| Sanger | Urticaceae | *Phenax sonneratii* |  | AM6399 | E | Bolivia | – | – | MH357917 | – | MH358117 | – | – | – |
| Sanger | Urticaceae | *Pilea alpina* |  |  | BM |  | – | – | DQ175543 | DQ179309 | – | – | – | – |
| Sanger | Urticaceae | *Pilea amplistipulata* | Shui Y.M. et. al. | 14319 | KUN | China | – | – | MT516340 | MT523095 | MT523051 | – | – | – |
| Sanger | Urticaceae | *Pilea angulata* | Fu L.F. et al. | FL0234 | IBK | China | – | – | MT516341 | MT523096 | MT523052 | – | – | – |
| Sanger | Urticaceae | *Pilea angustifolia* |  | 12-1838 | BM |  | – | – | DQ175556 | DQ179289 | – | – | – | – |

| Sanger | Urticaceae | *Pilea anisophylla* | Wen F. | WF182817-  10 | IBK | China | – | – | MT516342 | MT523097 | MT523053 | – | – | – |
| --- | --- | --- | --- | --- | --- | --- | --- | --- | --- | --- | --- | --- | --- | --- |
| Sanger | Urticaceae | *Pilea aphrophila* |  |  | BM |  | – | – | DQ175589 | DQ179323 | – | – | – | – |
| Sanger | Urticaceae | *Pilea aquarum* | Wei Y.G. | 97 | IBK | China | – | – | MT516343 | MT523098 | MT523054 | – | – | – |
| Sanger | Urticaceae | *Pilea aquarum subsp. acutidentata* | Wen F. | WFLSH111 207 | IBK | China | – | – | MT516344 | MT523099 | MT523055 | – | – | – |
| Sanger | Urticaceae | *Pilea balansae* | Huang S.L. | HSL118-1 | IBK | Vietnam | – | – | MT516345 | MT523100 | – | – | – | – |
| Sanger | Urticaceae | *Pilea basicordata* |  | HSL012 | PE | China | – | – | DQ175614 | DQ179361 | – | – | – | – |
| Sanger | Urticaceae | *Pilea benguetensis* |  |  | BM | Phillipines | – | – | DQ175554 | DQ179337 | – | – | – | – |
| Sanger | Urticaceae | *Pilea boniana* | Qin et al. 3193 | 3193 | KUN | China | – | – | MT516347 | MT523102 | MT523057 | – | – | – |
| Angiosperms353 | Urticaceae | *Pilea cadierei* |  |  | K | – | 24019 | – | – | – | – | – | – | ERS4414131 |
| Sanger | Urticaceae | *Pilea cadierei* | Xin Z.B. | XZB102 | IBK | China | – | – | MT516348 | MT523103 | MT523058 | – | – | – |
| Sanger | Urticaceae | *Pilea cavaleriei* | LiDZ | 1080 | KUN | China | – | – | KF137895 | KF138380 | KF138214 | – | – | – |
| Sanger | Urticaceae | *Pilea ciliata* |  |  | BM | Jamaica | – | – | DQ175538 | DQ179300 | – | – | – | – |
| Sanger | Urticaceae | *Pilea clementis* |  |  | BM | Cuba | – | – | DQ175550 | DQ179310 | – | – | – | – |
| Sanger | Urticaceae | *Pilea consanguinea* |  | AM6809 | BM | Santo Domingo | – | – | DQ175539 | DQ179312 | – | – | – | – |
| Sanger | Urticaceae | *Pilea cordistipulata* | Huang S.L. | HSL140 | IBK | China | – | – | MT516350 | MT523105 | MT523060 | – | – | – |
| Sanger | Urticaceae | *Pilea costata* |  | WuZY- 09199 | BM | Peru | – | – | DQ175595 | DQ179290 | – | – | – | – |
| Sanger | Urticaceae | *Pilea daguensis* |  | WuZY- 09107 | BM | Mexico | – | – | DQ175567 | DQ179332 | – | – | – | – |
| Sanger | Urticaceae | *Pilea dauciodora* |  | HSL124 | BM | Mexico | – | – | DQ175562 | DQ176857 | – | – | – | – |
| Sanger | Urticaceae | *Pilea digitata* |  |  | MO | Panama | – | – | DQ175559 | DQ179326 | – | – | – | – |
| Sanger | Urticaceae | *Pilea dolichocarpa* | Monro A.K. | 6399 | IBK | China | – | – | MT516351 | MT523106 | MT523061 | – | – | – |
| Sanger | Urticaceae | *Pilea dominguensis* |  |  | BM | Santo Domingo | – | – | DQ175541 | DQ179313 | – | – | – | – |
| Sanger | Urticaceae | *Pilea ecboliophylla* |  |  | BM | Mexico | – | – | DQ175531 | DQ179292 | – | – | – | – |
| Sanger | Urticaceae | *Pilea elegantissima* |  | 12-1247 |  | China | – | – | MH357923 | MH358303 | MH358124 | – | – | – |
| Sanger | Urticaceae | *Pilea elliptilimba* | Huang S.L. | HSL113 | IBK | China | – | – | MT516352 | MT523107 | – | – | – | – |
| Sanger | Urticaceae | *Pilea foliosa* |  |  | BM | Peru | – | – | DQ175571 | DQ179291 | – | – | – | – |
| Sanger | Urticaceae | *Pilea forgetii* |  | HSL149 | BM | Panama | – | – | DQ175585 | DQ179333 | – | – | – | – |
| Sanger | Urticaceae | *Pilea forsythiana* |  | HSL099 | BM | Dominica | – | – | DQ175546 | DQ179311 | – | – | – | – |
| Sanger | Urticaceae | *Pilea fruticosa* |  |  | BM | Borneo | – | – | DQ175604 | DQ179353 | – | – | – | – |
| Sanger | Urticaceae | *Pilea glaberrima* |  | STET2663 | BM | Nepal | – | – | DQ175600 | DQ179352 | – | – | – | – |
| Sanger | Urticaceae | *Pilea gracilis* | Wei Y.G. | 39 | IBK | China | – | – | MT516353 | MT523108 | – | – | – | – |
| Sanger | Urticaceae | *Pilea grandifolia* |  | AM6818 | BM | Jamaica | – | – | DQ175551 | DQ179303 | – | – | – | – |
| Sanger | Urticaceae | *Pilea guizhouensis* | Monro A.K. | 6715 | IBK | China | – | – | MT516354 | MT523109 | MT523062 | – | – | – |
| Sanger | Urticaceae | *Pilea harrisii* |  |  | BM | Jamaica | – | – | DQ175537 | DQ179302 | – | – | – | – |
| Sanger | Urticaceae | *Pilea hexagona* | Sino-Vietnamese expidition | 775 | KUN | China | – | – | MT516356 | MT523111 | MT523064 | – | – | – |
| Sanger | Urticaceae | *Pilea hilliana* | liuzu | 2014 | KUN | China | – | – | MT516357 | MT523112 | MT523065 | – | – | – |
| Sanger | Urticaceae | *Pilea howelliana* | Wang Y.Z. | 4678 | KUN | China | – | – | MT516358 | MT523113 | MT523066 | – | – | – |
| Sanger | Urticaceae | *Pilea inaequalis* |  | HGX001 |  | Trinidad | – | – | DQ175552 | DQ179304 | – | – | – | – |
| Sanger | Urticaceae | *Pilea insolens* | FLPH Tibet Expedition | 12-1838 | IBK | China | – | – | MT516359 | MT523114 | MT523067 | – | – | – |
| Sanger | Urticaceae | *Pilea irrorata* |  |  | BM | Mexico | – | – | DQ175535 | DQ179294 | – | – | – | – |
| Sanger | Urticaceae | *Pilea japonica* | Huang S.L. | HSL012 | IBK | China | – | – | MT516360 | MT523115 | MT523068 | – | – | – |
| Sanger | Urticaceae | *Pilea krugii* |  | HSL116 | BM | Puerto Rico | – | – | DQ175581 | DQ179315 | – | – | – | – |
| Sanger | Urticaceae | *Pilea lapestris* |  | 726 | BM | Indonesia | – | – | DQ175598 | DQ179341 | – | – | – | – |
| Sanger | Urticaceae | *Pilea lindeniana* |  |  | BM | Cuba | – | – | DQ175547 | DQ179314 | – | – | – | – |
| Angiosperms353 | Urticaceae | *Pilea longicaulis* | Monro, A.K. | 7590 | K | – | – | – | – | – | – | – | – | ERS5501984 |
| Sanger | Urticaceae | *Pilea longicaulis* |  |  | PE | China | – | – | DQ175611 | DQ179363 | – | – | – | – |
| Sanger | Urticaceae | *Pilea longicaulis var. erosa* | Monro A.K. | 6809 | IBK | China | – | – | MT516361 | MT523116 | – | – | – | – |
| Sanger | Urticaceae | *Pilea longipedunculata* | WuZY | 9199 | KUN | China | – | – | KF137897 | KF138382 | KF138216 | – | – | – |
| Sanger | Urticaceae | *Pilea martinii* | WuZY | 9107 | KUN | China | – | – | KF137898 | KF138383 | KF138217 | – | – | – |

| Sanger | Urticaceae | *Pilea melastomoides* | Huang S.L. | HSL124 | IBK | China | – | – | MT516363 | MT523118 | MT523070 | – | – | – |
| --- | --- | --- | --- | --- | --- | --- | --- | --- | --- | --- | --- | --- | --- | --- |
| Sanger | Urticaceae | *Pilea mexicana* |  | AM6433 | BM | Panama | – | – | DQ175579 | DQ179278 | – | – | – | – |
| Sanger | Urticaceae | *Pilea microphylla* |  |  | BM | Brazil | – | – | MH357928 | MH358307 | MH358129 | – | – | – |
| Sanger | Urticaceae | *Pilea monilifera* |  |  |  | China | – | – | MK911055 | MK911077 | MK911100 | – | – | – |
| Sanger | Urticaceae | *Pilea multicellularis* | Tibet Expedition | 12-1247 | IBK | China | – | – | MT516365 | MT523120 | MT523072 | – | – | – |
| Sanger | Urticaceae | *Pilea nigrescens* |  |  | BM | Jamaica | – | – | DQ175582 | DQ179301 | – | – | – | – |
| Sanger | Urticaceae | *Pilea nonggangensis* | Huang S.L. | HSL149 | IBK | China | – | – | MT516366 | MT523121 | MT523073 | – | – | – |
| Sanger | Urticaceae | *Pilea notata* | Huang S.L. | HSL099 | IBK | China | – | – | MT516368 | MT523123 | MT523075 | – | – | – |
| Sanger | Urticaceae | *Pilea nummularifolia* |  |  | BM | Peru | – | – | DQ175588. | DQ179316 | – | – | – | – |
| Sanger | Urticaceae | *Pilea oxyodon* |  |  | KUN | China | – | – | KF137902 | KF138387 | KF138221 | – | – | – |
| Sanger | Urticaceae | *Pilea paniculigera* | Monro A.K. | 6818 | IBK | China | – | – | MT516369 | MT523124 | MT523076 | – | – | – |
| Sanger | Urticaceae | *Pilea pansamalana* |  | Wei047 | BM | Mexico | – | – | DQ175533 | DQ179296 | – | – | – | – |
| Sanger | Urticaceae | *Pilea pellionioides* | Hu G.X. | HGX001 | IBK | China | – | – | MT516370 | MT523125 | MT523077 | – | – | – |
| Sanger | Urticaceae | *Pilea pelonae* |  |  | BM | Dominican | – | – | DQ175540 | DQ179327 | – | – | – | – |
| Sanger | Urticaceae | *Pilea peltata* | Huang S.L. | HSL116 | IBK | China | – | – | MT516371 | MT523126 | MT523078 | – | – | – |
| Sanger | Urticaceae | *Pilea penninervis* | Wei Y.G. | 726 | IBK | China | – | – | MT516372 | MT523127 | MT523079 | – | – | – |
| Sanger | Urticaceae | *Pilea peperomiifolia* |  | 2104H | BM | Virgin Islands | – | – | DQ175569 | DQ179281 | – | – | – | – |
| Sanger | Urticaceae | *Pilea peperomioides* |  | WF150423- 33 | BM | cultivated in UK | – | – | DQ175605 | DQ179350 | – | – | – | – |
| Sanger | Urticaceae | *Pilea peploides* | Monro A.K. | 6433 | IBK | China | – | – | MT516373 | MT523128 | MT523080 | – | – | – |
| Sanger | Urticaceae | *Pilea peploides var. major* |  |  | BM | China | – | – | MH357931 | – | MH358132 | – | – | – |
| Sanger | Urticaceae | *Pilea pittieri* |  |  |  | Peru | – | – | DQ175560 | DQ179328 | – | – | – | – |
| Sanger | Urticaceae | *Pilea plataniflora* |  |  | BM | Japan | – | – | DQ175599 | DQ179349 | – | – | – | – |
| Sanger | Urticaceae | *Pilea pleuroneura* |  |  | BM | Guatemala | – | – | DQ175532 | DQ179297 | – | – | – | – |
| Sanger | Urticaceae | *Pilea pseudonotata* | Wei Y.G. | 47 | IBK | China | – | – | MT516375 | MT523130 | MT523082 | – | – | – |
| Sanger | Urticaceae | *Pilea pubescens* |  | EXLS-0272 | BM | Belize | – | – | DQ175558 | DQ179325 | – | – | – | – |
| Sanger | Urticaceae | *Pilea pumila* | Huang S.L. | 2104H | IBK | China | – | – | MT516376 | MT523131 | MT523083 | – | – | – |
| Sanger | Urticaceae | *Pilea racemiformis* | Wen F. | WF150423- 33 | IBK | China | – | – | MT516377 | MT523132 | MT523084 | – | – | – |
| Sanger | Urticaceae | *Pilea racemosa* |  |  | BM | China | – | – | DQ175602 | DQ179347 | – | – | – | – |
| Sanger | Urticaceae | *Pilea receptacularis* |  |  | PE | China | – | – | DQ175612 | DQ179362 | – | – | – | – |
| Sanger | Urticaceae | *Pilea rivularis* |  | 37636 | BM | Tanzania | – | – | DQ175606 | DQ179358 | – | – | – | – |
| Sanger | Urticaceae | *Pilea rufa* |  | WYG18051 9-02 | BM | Jamaica | – | – | DQ175578 | DQ179299 | – | – | – | – |
| Sanger | Urticaceae | *Pilea semisessilis* | Zhou Z.K. et al. | EXLS-0272 | KUN | China | – | – | MT516378 | MT523133 | MT523085 | – | – | – |
| Sanger | Urticaceae | *Pilea sinofasciata* |  |  | BM | China | – | – | KF137905 | KF138389 | KF138224 | – | – | – |
| Sanger | Urticaceae | *Pilea spathulifolia* |  |  | BM | Dominican  Republic | – | – | DQ175570 | DQ179282 | – | – | – | – |
| Sanger | Urticaceae | *Pilea spicata* | Burkill H. | 37636 | K | China | – | – | MT516380 | MT523135 | MT523087 | – | – | – |
| Sanger | Urticaceae | *Pilea subcoriacea* | Wei Y.G. | 180519-02 | IBK | China | – | – | MT516381 | MT523136 | MT523088 | – | – | – |
| Sanger | Urticaceae | *Pilea succulenta* |  |  | BM | Cayman Islands | – | – | DQ175565 | DQ179280 | – | – | – | – |
| Sanger | Urticaceae | *Pilea swinglei* |  |  | PE | China | – | – | MH357933 | – | MH358134 | – | – | – |
| Sanger | Urticaceae | *Pilea ternifolia* |  |  | BM | Nepal | – | – | DQ175597 | DQ179346 | – | – | – | – |
| Sanger | Urticaceae | *Pilea tetraphylla* |  |  | BM | Madagascar | – | – | MH357934 | MH358310 | MH358135 | – | – | – |
| Sanger | Urticaceae | *Pilea thymifolia* |  |  | BM | Peru | – | – | DQ175568 | DQ179283 | – | – | – | – |
| Sanger | Urticaceae | *Pilea tridentata* |  | AM6770 | BM | Mexico | – | – | DQ175536 | DQ179293 | – | – | – | – |
| Sanger | Urticaceae | *Pilea tripartita* |  | WF180821- 01 | BM | Panama | – | – | DQ175617 | DQ176859 | – | – | – | – |
| Sanger | Urticaceae | *Pilea tsiangiana* | Monro A.K. | 6770 | IBK | China | – | – | MT516382 | MT523137 | MT523089 | – | – | – |
| Sanger | Urticaceae | *Pilea umbrosa* | Wen F. | WF180821- 01 | IBK | China | – | – | MT516383 | MT523138 | MT523090 | – | – | – |

| Sanger | Urticaceae | *Pilea unciformis* | Huang S.L. | HSL132 | IBK | China | – | – | MT516384 | MT523139 | MT523091 | – | – | – |
| --- | --- | --- | --- | --- | --- | --- | --- | --- | --- | --- | --- | --- | --- | --- |
| Sanger | Urticaceae | *Pilea villicaulis* | Shui et al. | 12871 | KUN | China | – | – | MT516385 | MT523140 | MT523092 | – | – | – |
| Sanger | Urticaceae | *Pilea virgata* |  | HSL132 | BM | Jamaica | – | – | DQ175548 | DQ179329 | – | – | – | – |
| Sanger | Urticaceae | *Pilea vulcanica* |  | 12871 | BM | Panama | – | – | DQ175563 | DQ179284 | – | – | – | – |
| Sanger | Urticaceae | *Pilea weddellii* |  |  | BM | Jamaica | – | – | DQ175545 | DQ179308 | – | – | – | – |
| Sanger | Urticaceae | *Pilea weimingii* | Lv R.D. | LRD001 | IBK | China | – | – | MT516386 | MT523141 | MT523093 | – | – | – |
| Sanger | Urticaceae | *Pipturus arborescens* |  | 11879 | KUN | Taiwan, China | – | – | KF137908 | KF138392 | KF138227 | KF138545 | – | – |
| Sanger | Urticaceae | *Pipturus argenteus* |  |  | E | Papua New Guinea | – | – | MH357935 | MH358311 | – | MH358236 | – | – |
| Sanger | Urticaceae | *Pipturus kauaiensis* |  | 90441 |  | Hawaii, USA | – | – | KF137910 | KF138394 | KF138229 | KF138546 | KF138051 | – |
| Angiosperms353 | Urticaceae | *Pipturus montanus* | Crayn, D.M.; Sennart, S. | 540 | K | – | – | – | – | – | – | – | – | ERS5503161 |
| Sanger | Urticaceae | *Pipturus ruber* |  | 8082 | KUN | Hawaii, USA | – | – | – | – | – | – | – | – |
| Angiosperms353 | Urticaceae | *Poikilospermum acuminatum* | Risdale, C.E. et al. | 1268 |  |  | – | – | – | – | – | – | – | – |
| Angiosperms353 | Urticaceae | *Poikilospermum amboinense* | Takeuchi, W.; Ama, D. | 16391 |  |  | – | – | – | – | – | – | – | – |
| Sanger | Urticaceae | *Poikilospermum cordifolium* | Sinclair&Kadim | 10358 | E | Malaysia | – | – | To submit | – | – | – | – | – |
| Angiosperms353 | Urticaceae | *Poikilospermum cordifolium* | DeWilde et al. | SAN 143991 |  |  | – | – | – | – | – | – | – | – |
| Angiosperms353 | Urticaceae | *Poikilospermum erectum* | Risdale, C.E. | SMHI 482 |  |  | – | – | – | – | – | – | – | – |
| Angiosperms353 | Urticaceae | *Poikilospermum inaequale* | Takeuchi, W. et al. | 19531 |  |  | – | – | – | – | – | – | – | – |
| Sanger | Urticaceae | *Poikilospermum lanceolatum* | WuZY | 9235 | KUN | China | – | KF137786 | KF137912 | KF138396 | KF138231 | KF138548 | KF138053 | – |
| Angiosperms353 | Urticaceae | *Poikilospermum lanceolatum* | Middleton, D.J. et al. | 1744 |  |  | – | – | – | – | – | – | – | – |
| Sanger | Urticaceae | *Poikilospermum lanceolatum* | Wu | 9235 | KUN | China | – | – | KF137912 | KF138396 | – | – | – | – |
| Angiosperms353 | Urticaceae | *Poikilospermum microstachys* | Niyomdham, C. et al. | 1027 |  |  | – | – | – | – | – | – | – | – |
| Angiosperms353 | Urticaceae | *Poikilospermum naucleiflorum* | Chantaranothai, P. | 1215 |  |  | – | – | – | – | – | – | – | – |
| Angiosperms353 | Urticaceae | *Poikilospermum nobile* | Johns, R.J. | 8786 |  |  | – | – | – | – | – | – | – | – |
| Sanger | Urticaceae | *Poikilospermum scabrinervium* | Wilkie | 94166 | E | Indonesia | – | – | To submit | – | – | – | – | – |
| Angiosperms353 | Urticaceae | *Poikilospermum scabrinervium* | Dewol et al. | SAN 124644 |  |  | – | – | – | – | – | – | – | – |
| Angiosperms353 | Urticaceae | *Poikilospermum scortechinii* | T&P | 476 |  |  | – | – | – | – | – | – | – | – |
| Sanger | Urticaceae | *Poikilospermum sp. 1* | Sun | 13176 | KUN | Laos | – | – | KM586453 | KM586625 | – | – | – | – |
| Sanger | Urticaceae | *Poikilospermum suaveolens* | WuZY | 9160 | KUN | China | – | – | KF137914 | KF138398 | KF138233 | KF138550 | KF138054 | – |
| Angiosperms353 | Urticaceae | *Poikilospermum suaveolens* | Radin, J. et al. | SAN 133007 |  |  | – | – | – | – | – | – | – | – |
| Sanger | Urticaceae | *Poikilospermum suaveolens* |  |  | KUN | China | – | – | KF137913 | KF138397 | – | – | – | – |
| Angiosperms353 | Urticaceae | *Poikilospermum tangaum* | Postar; Geoffray | SAN 145768 |  |  | – | – | – | – | – | – | – | – |
| Sanger | Urticaceae | *Pouzolzia australis* | Hadiah | 393 | NSW | Australia: South  Pacific | – | – | – | AY208723 | AY208700 | – | – | – |
| Sanger | Urticaceae | *Pouzolzia calophylla* |  |  |  | Xizang, China | – | – | KF137915 | KF138399 | KF138234 | KF138551 | – | – |
| Sanger | Urticaceae | *Pouzolzia elegans var. elegans* | L0942456 | L0942456 | KUN | Taiwan, China | – | – | – | – | MH358140 | – | – | – |
| Sanger | Urticaceae | *Pouzolzia guineensis* | Bidgood et al. | 3008 | K | Tanzania | – | – | – | KF138400 | KF138235 | KF138552 | KF138055 | – |
| Sanger | Urticaceae | *Pouzolzia mixta* | Lovett & Congdon | 2945 | K | Tanzania | – | – | KF137916 | KF138401 | KF138236 | KF138553 | – | – |
| Sanger | Urticaceae | *Pouzolzia poeppigiana* |  | HSL140 | E | Peru | – | – | MH357938 | MH358315 | MH358141 | – | – | – |
| Sanger | Urticaceae | *Pouzolzia rugulosa* | LiDZ | 1071 | KUN | Nepal, Kathmandu | – | – | KF137817 | KF138288 | KF138125 | KF138449 | KF137960 | – |
| Angiosperms353 | Urticaceae | *Pouzolzia sanguinea* | Sambuling, S. | 449 | K | Malaysia | 23922 | – | – | – | – | – | – | ERS5501735 |
| Sanger | Urticaceae | *Pouzolzia sanguinea* | WuZY | 9483 |  | Guangxi, China | – | – | KF137918 | KF138403 | KF138238 | KF138555 | KF138057 | – |
| Sanger | Urticaceae | *Pouzolzia sanguinea var. elegans* |  | Wei039 |  | Xizang, China | – | – | KF137917 | KF138402 | KF138237 | KF138554 | KF138056 | – |
| Sanger | Urticaceae | *Pouzolzia sp.* | RC | 1682 |  | Panchthar, Nepal | – | – | KF137919 | KF138404 | KF138239 | KF138556 | KF138058 | – |
| Angiosperms353 | Urticaceae | *Pouzolzia zeylanica* | Sibil, J. | 83 | K | Malaysia | 23923 | – | – | – | – | – | – | ERS5501736 |
| Sanger | Urticaceae | *Pouzolzia zeylanica* | WuZY | 10167 |  | Yunnan, China | – | – | KF137920 | KF138405 | KF138240 | KF138557 | KF138059 | – |
| Sanger | Urticaceae | *Procris archboldiana* | A. C. Smith | 5987 | K | Fiji | – | – | KP858785 | – | – | – | – | – |

| Sanger | Urticaceae | *Procris crenata* | Y. H. Tseng | 1170 | TAI | China | – | – | KP858782 | – | – | – | – | – |
| --- | --- | --- | --- | --- | --- | --- | --- | --- | --- | --- | --- | --- | --- | --- |
| Sanger | Urticaceae | *Procris crenata* | Shi | 15115 | A | China | – | – | KP858783 | – | – | – | – | – |
| Sanger | Urticaceae | *Procris frutescens* | Takeuchi | 8800 | A | Papua New Guinea | – | – | KP858781 | – | – | – | – | – |
| Sanger | Urticaceae | *Procris montana* | R. O. Gardenr | 5955 | MO | Australia | – | – | KP858786 | – | – | – | – | – |
| Angiosperms353 | Urticaceae | *Procris wightiana* | Monro, A.K.; Fu, L.F. | 7603 | K | – | – | – | – | – | – | – | – | ERS5502206 |
| Angiosperms353 | Urticaceae | *Rousselia humilis* | Ekman, E.L. | 4973 | K | Haiti | 23945 | – | – | – | – | – | – | ERS5501747 |
| Sanger | Urticaceae | *Rousselia humilis* | Howard | 6273 | US | Cuba | – | – | – | KM586645 | KM586559 | – | – | – |
| Sanger | Urticaceae | *Rousselia humilis* | Haroslav & Holman | 435 | BM | Cuba | – | – | – | – | – | – | – | – |
| Angiosperms353 | Urticaceae | *Sarcochlamys pulcherrima* | Grierson, A.J.C.; Long, D.G. | 1525 | K | Bhutan | 23939 | – | – | – | – | – | – | ERS5501743 |
| Sanger | Urticaceae | *Sarcochlamys pulcherrima* |  |  | KUN | China | – | KF137790 | KF137924 | KF138409 | KF138244 | KF138561 | – | – |
|  |  |  |  |  |  | United Kingdom of Great Britain and Northern Ireland (the) |  |  |  |  |  |  |  |  |
| Angiosperms353 | Urticaceae | *Soleirolia soleirolii* | Sheahan, M-C. | 17 | K |  | 8154 | – | – | – | – | – | – | ERS4414200 |
| Sanger | Urticaceae | *Soleirolia soleirolii* | Monro A.K. | s.n. | BM | Cultivated | – | – | KF137926 | KF138411 | KF138246 | KF138563 | KF138063 | – |
| Angiosperms353 | Urticaceae | *Touchardia latifolia* | Stone, B.C. | 3626 | K | United States of America | 23938 | – | – | – | – | – | – | ERS4414130 |
| Sanger | Urticaceae | *Touchardia latifolia* | Jeffrey | 201101 | KUN | Hawaii | – | – | KF137927 | KF138412 | – | – | – | – |
| Sanger | Urticaceae | *Urera altissima* | Lliully | 460 | K | Bolivia | – | – | To submit | – | – | – | – | – |
| Sanger | Urticaceae | *Urera aurantiaca* | Loza | 63 | K | Bolivia | – | – | To submit | To submit | – | – | – | – |
| Angiosperms353 | Urticaceae | *Urera baccifera* | Goes, S.P.; et al. | 20285  /G150694 | K | – | – | – | – | – | – | – | – | ERS4414134 |
| Sanger | Urticaceae | *Urera baccifera* | Cayola | 2530 | BM | Bolivia | – | – | To submit | To submit | – | – | – | – |
| Sanger | Urticaceae | *Urera batesii* | Carvalho | 3412 | K | Equatorial Guinea | – | – | KF971186 | KF971219 | – | – | – | – |
| Sanger | Urticaceae | *Urera cameroonensis* | Leeuenberg | 7187 | K |  | – | – | To submit | – | – | – | – | – |
| Angiosperms353 | Urticaceae | *Urera caracasana* | Monro, A.K. | 6840 | K | – | – | – | – | – | – | – | – | ERS4414132 |
| Sanger | Urticaceae | *Urera caracasana* | Wood | 8834 | K | Bolivia | – | – | KF137929.1 | KF138415 | – | – | – | – |
| Sanger | Urticaceae | *Urera cordifolia* | Carvalho | 3046 | K | CM | – | – | To submit | – | – | – | – | – |
| Sanger | Urticaceae | *Urera cordifolia* | Sunderland | 1190 | K | CM | – | – | To submit | – | – | – | – | – |
| Sanger | Urticaceae | *Urera elata* | Lewis | 2224 | US | Panama | – | – | KM58647 | KM586642 | – | – | – | – |
| Sanger | Urticaceae | *Urera fenestrata* | Monro A.K. | 5452 | K | Costa Rica | – | – | To submit | To submit | – | – | – | – |
| Sanger | Urticaceae | *Urera fischeri* | Luke & Luke | 7087 | K | KE | – | – | To submit | – | – | – | – | – |
| Sanger | Urticaceae | *Urera fischeri* | Faden & Beentje | 8522 | EA | Kenya | – | – | KM586427 | KM586599 | – | – | – | – |
| Sanger | Urticaceae | *Urera glabra* | s.n. | 100694 | KUN | Hawaii | – | – | KF1379930 | KF138416 | – | – | – | – |
| Sanger | Urticaceae | *Urera glabriuscula* | Calonico | 21101 | BM | Mexico | – | – | To submit | – | – | – | – | – |
| Angiosperms353 | Urticaceae | *Urera hypselodendron* | Faden, R.B. et al. | 85200 | K | Kenya | 23913 | – | – | – | – | – | – | ERS4414137 |
| Sanger | Urticaceae | *Urera hypselodendron* | Beenlije | 3257 | EA | Kenya | – | – | KM586430 | KM586602 | – | – | – | – |
| Sanger | Urticaceae | *Urera keayi* | Leeuwenberg | 4474 | K | CI | – | – | To submit | – | – | – | – | – |
| Sanger | Urticaceae | *Urera killipiana* | Serviu | 372 | BM | Mexico | – | – | To submit | To submit | – | – | – | – |
| Angiosperms353 | Urticaceae | *Urera laciniata* | Morawetz, W.; Wallnofer, B. | M13-29985 | K | – | – | – | – | – | – | – | – | ERS4414126 |
| Sanger | Urticaceae | *Urera laciniata* | Araujo | 3016 | BM | Bolivia | – | – | To submit | To submit | – | – | – | – |
| Sanger | Urticaceae | *Urera lianiformis* | Solano | 6825 | BM | Costa Rica | – | – | KF138570 | KF138418 | – | – | – | – |
| Sanger | Urticaceae | *Urera mannii* | Lowe | 1736 | K | NG | – | – | To submit | – | – | – | – | – |
| Sanger | Urticaceae | *Urera oblongifolia* | Morton & Jarr | 3517 | K | SL | – | – | To submit | – | – | – | – | – |
| Sanger | Urticaceae | *Urera obovata* | Thijssen | 40 | K | CI | – | – | To submit | – | – | – | – | – |
| Sanger | Urticaceae | *Urera pacifica* | Steinman | 3265 | BM | Mexico | – | – | To submit | To submit | – | – | – | – |
| Sanger | Urticaceae | *Urera repens* | Eiminjeze & Oguntayo | 72749 | K | NG | – | – | To submit | – | – | – | – | – |
| Sanger | Urticaceae | *Urera rigida* | Adam | 25511 | K | LR | – | – | To submit | – | – | – | – | – |
| Sanger | Urticaceae | *Urera rigida* | Breteler | Breteler | K | LR | – | – | To submit | – | – | – | – | – |

| Sanger | Urticaceae | *Urera robusta* | Adams | 4823 | K | GH | – | – | To submit | – | – | – | – | – |
| --- | --- | --- | --- | --- | --- | --- | --- | --- | --- | --- | --- | --- | --- | --- |
| Angiosperms353 | Urticaceae | *Urera sandwicensis* | Bulmer, C. | 379 | K | United States of America | 23914 | – | – | – | – | – | – | – |
| Sanger | Urticaceae | *Urera sansibarica* | Luke | 11527 | EA | Tanzania | – | – | KM586428 | KM586600 | – | – | – | – |
| Sanger | Urticaceae | *Urera simplex* | Monro A.K. | 5102 | K | Panama | – | – | To submit | To submit | – | – | – | – |
| Sanger | Urticaceae | *Urera spaerophyllya* | Humbert | 3055 | K | MG | – | – | To submit | – | – | – | – | – |
| Sanger | Urticaceae | *Urera thonneri* | Breteler et al. | 2378 | K | CM | – | – | To submit | – | – | – | – | – |
| Sanger | Urticaceae | *Urera trinervis* | Friis | 3920 | C | Ethiopia | – | – | KF137932 | KF138421 | – | – | – | – |
| Sanger | Urticaceae | *Urtica angustifolia* |  |  | KUN | China | – | KF137796 | KF137933 | KF138421 | KF138256 | KF138573 | KF138067 | – |
| Sanger | Urticaceae | *Urtica aquatica* | M. Weigend | 7478 | BSB | Peru | – | – | KF971214 | KF971247 | – | – | – | – |
| Sanger | Urticaceae | *Urtica ardens* |  | 81152 | KUN | China | – | – | KF137934 | KF138422 | KF138257 | KF138574 | – | – |
| Sanger | Urticaceae | *Urtica atrichocaulis* | WuZY | 10358 | KUN | China | – | – | KF137935 | KF138423 | KF13825 | KF138575 | KF138068 | – |
| Sanger | Urticaceae | *Urtica atrovirens* |  |  | E | Italy | – | – | MH357956 | – | MH358161 | MH358244 | – | – |
| Sanger | Urticaceae | *Urtica cannabina* | LZH | 2012 | KUN | China | – | – | MH357957 | MH358322 | MH358162 | MH358245 | MH358029 | – |
| Sanger | Urticaceae | *Urtica dioica subsp. dioica* | BROWP | 135 | KUN | UK | – | – | KF137936 | KF138424 | KF138259 | KF138576 | – | – |
| Sanger | Urticaceae | *Urtica echinata* |  |  | E | Peru | – | – | MH357961 | MH358325 | MH358166 | MH358248 | – | – |
| Sanger | Urticaceae | *Urtica fissa* | WuZY | 10378 | KUN | China | – | – | KF137937 | KF138425 | KF138260 | KF138577 | KF138069 | – |
| Sanger | Urticaceae | *Urtica flabellata* | M. Weigend et al. | 7728 | BSB | Peru | – | – | KF558908 | KF559028 | – | – | – | – |
| Sanger | Urticaceae | *Urtica hyperborea* |  | 81200 | KUN | China | – | – | KF137939 | KF138427 | KF138262 | KF13857 | KF138071 | – |
| Sanger | Urticaceae | *Urtica kioviensis* |  |  | E | Austria | – | – | MH357963 | MH358326 | MH358168 | MH358250 | MH358032 | – |
| Sanger | Urticaceae | *Urtica laetevirens* | Mawenzhang | 2011 | KUN | Canada | – | – | – | MH358327 | MH358169 | MH358251 | – | – |
| Sanger | Urticaceae | *Urtica magellanica* |  |  | E | Chile | – | – | MH357964 | – | MH358170 | MH358252 | – | – |
| Sanger | Urticaceae | *Urtica mairei* | WuZY | 9354 | KUN | China | – | KF137797 | KF137940 | KF138428 | KF138263 | KF138580 | – | – |
| Sanger | Urticaceae | *Urtica massaica* | J.M. Kimen & al. | KARI42/02 | EA | Kenya | – | – | KM586438 | KM586610 | KM586524 | – | – | – |
| Sanger | Urticaceae | *Urtica membranacea* |  |  | E | Greece | – | – | MH357968 | MH358329 | MH358174 | MH358255 | – | – |
| Sanger | Urticaceae | *Urtica sp. 1* | Lixinhui | 1102 | KUN | Kenya | – | – | KF137941 | KF138429 | KF138264 | KF138581 | KF138072 | – |
| Sanger | Urticaceae | *Urtica taiwaniana* | H. Sun | 11345 | KUN | Taiwan | – | – | KM586420 | KM586592 | KM586506 | – | – | – |
| Sanger | Urticaceae | *Urtica thunbergiana* | H. Sun | 11881 | KUN | China | – | – | KM586421 | KM586593 | KM586507 | – | – | – |
| Sanger | Urticaceae | *Urtica triangularis subsp. pinnatifida* |  | 80860 | KUN | China | – | – | KF137943 | KF138431 | KF138266 | KF138583 | KF138073 | – |
|  |  |  |  |  |  | United Kingdom of Great Britain and Northern Ireland  (the) |  |  |  |  |  |  |  |  |
| Angiosperms353 | Urticaceae | *Urtica urens* | Fay, M.F. | 173 | K |  | 8160 | – | – | – | – | – | – | ERS4414127 |
| Sanger | Urticaceae | *Urtica zayuensis* | WuZY | 10361 | KUN | China | – | – | KF137945 | KF138433 | KF138268 | KF138585 | KF138075 | – |
| Angiosperms353 | Urticaceae | *Zhengyia shennongensis* | Deng, T.; Zhang, D.G.; Sun, H. | 2295 | K | – | – | – | – | – | – | – | – | ERS5501754 |
| Sanger | Urticaceae | *Zhengyiia shennongensis* | SNJ Exped. | 2.0111E+10 | KUN | China | – | – | KC284949 | KC285001 | KC284975 | – | – | – |
